# Annelid adult cell type diversity and their pluripotent cellular origins

**DOI:** 10.1101/2023.04.25.537979

**Authors:** Patricia Álvarez-Campos, Helena García-Castro, Elena Emili, Alberto Pérez-Posada, David A. Salamanca-Díaz, Vincent Mason, Bria Metzger, Alexandra E. Bely, Nathan Kenny, B. Duygu Özpolat, Jordi Solana

## Abstract

Annelids are a broadly distributed, highly diverse, economically and environmentally important group of animals. Most species can regenerate missing body parts, and many are able to reproduce asexually. Therefore, many annelids can generate all adult cell types in adult stages. However, the putative adult stem cell populations involved in these processes, as well as the diversity of adult cell types generated by them, are still unknown. Here, we recover 75,218 single cell transcriptomes of *Pristina leidyi*, a highly regenerative and asexually-reproducing freshwater annelid. We characterise all major annelid adult cell types, and validate many of our observations by HCR *in situ* hybridisation. Our results uncover complex patterns of regionally expressed genes in the annelid gut, as well as neuronal, muscle and epidermal specific genes. We also characterise annelid-specific cell types such as the chaetal sacs and *globin*+ cells, and novel cell types of enigmatic affinity, including a *vigilin*+ cell type, a *lumbrokinase*+ cell type, and a diverse set of metabolic cells. Moreover, we characterise transcription factors and gene networks that are expressed specifically in these populations. Finally, we uncover a broadly abundant cluster of putative stem cells with a pluripotent signature. This population expresses well-known stem cell markers such as *vasa, piwi* and *nanos* homologues, but also shows heterogeneous expression of differentiated cell markers and their transcription factors. In these *piwi*+ cells, we also find conserved expression of pluripotency regulators, including multiple chromatin remodelling and epigenetic factors. Finally, lineage reconstruction analyses reveal the existence of differentiation trajectories from *piwi*+ cells to diverse adult types. Our data reveal the cell type diversity of adult annelids for the first time and serve as a resource for studying annelid cell types and their evolution. On the other hand, our characterisation of a *piwi+* cell population with a pluripotent stem cell signature will serve as a platform for the study of annelid stem cells and their role in regeneration.

## Introduction

Most annelid species can regenerate at least some body parts and continuously add new body segments from a posterior growth zone throughout their lives. Many are also capable of asexual reproduction by fragmentation or fission. Therefore, many annelids can generate and regenerate all adult cell types from pieces of the adult body (Bely, 2006, 2014). The cellular and molecular mechanisms of these adult cell differentiation processes are still poorly understood. Most annelids have active proliferation zones that produce new segments and tissues throughout their lives from the tail end and in the fission zones. Cell populations that express conserved stem cell markers such as *piwi* have been detected in these proliferative zones (Özpolat and Bely, 2015, 2016). However, whether these are stem cells and the extent of their developmental potency is unknown. To understand how annelids continuously produce new differentiated cells during posterior growth, asexual fission and regeneration, it is key to elucidate how many cell types are present in adult annelids, and to reconstruct their adult differentiation trajectories.

Tracing developmental cell lineages is remarkably difficult in adult animal models without well-developed transgenesis. Single cell transcriptomics (scRNA-seq) has emerged as a powerful tool to study the cellular composition – the *cell type atlas* – of multicellular organisms (Tanay and Sebe-Pedros, 2021). But, importantly, scRNA-seq has also fuelled the development of lineage reconstruction algorithms (Tritschler et al., 2019). These algorithms order cells in their differentiation trajectory, revealing the genetic changes that underlie the transition from stem cell to differentiated cell types. Making use of this powerful approach, differentiation trajectories have been described in adult cell type differentiation models such as planarians (Plass et al., 2018), cnidarians (Siebert et al., 2019), sponges (Musser et al., 2021), and amphibians (Gerber et al., 2018; Lust et al., 2022).

Cell type atlases of embryonic and larval annelids have previously been generated (Achim et al., 2018; Sur and Meyer, 2021; Vergara et al., 2021). However, despite the multiplication of single cell atlas studies in diverse metazoan species, annelid adult cell types and their differentiation trajectories are still uncharacterised. *Pristina leidyi* (hereafter referred to as *Pristina*) is a convenient laboratory model annelid to address these questions (Bely, 2022; Zattara and Bely, 2011). It grows very rapidly in culture conditions by asexual reproduction, using a mechanism called paratomic fission, in which the worm starts forming and differentiating new head and tail segments from within a single body segment, producing a chain of worms (Zattara and Bely, 2011). Eventually these clones separate and become distinct individuals. Thus, these worms are constantly generating all body parts and therefore all adult cell types. Three different zones of intense proliferation have been described in adult *Pristina* worms by S-phase cell EdU/BrdU labelling, located in the anterior end, the posterior end and the fission zones (Zattara and Bely, 2011, 2013). These areas also contain large numbers of *piwi*+, *nanos*+ and *vasa*+ cells (Özpolat and Bely, 2015, 2016). This molecular signature has been associated with the stem cells of very diverse invertebrates (Gehrke and Srivastava, 2016; Juliano et al., 2010; Lai and Aboobaker, 2018; Solana, 2013). The transcriptome of these cells has been profiled in some organisms, giving insight into their expression patterns and their heterogeneity, which reflects their developmental potency. For instance, within the stem cell pool in planarians there are stem cells that coexpress *piwi* with transcription factors characteristic of differentiated cell types. However, in annelids, the transcriptional profiles of *piwi+* cells and their differentiated counterparts are still unknown.

Here we used scRNA-seq to profile the adult cell type atlas of *Pristina* and reconstruct differentiation trajectories. We characterised all major adult cell types and uncovered an abundant *piwi*+ cell cluster with a clear stem cell signature. We reconstructed *piwi*+ cell differentiation trajectories to diverse cell types, a signature of pluripotency. We also showed that this population is heterogeneous, indicating the presence of committed stem cells. Finally, we characterised the molecular signature of annelid *piwi*+ cells at both the transcriptional and epigenetic level, and revealed their transcriptional program composed of RNA binding proteins, cell cycle control, DNA repair mechanisms, and chromatin regulators. Our data show that adult cell type differentiation in *Pristina* is underlied by a *piwi+* cell population with a pluripotent stem cell signature.

### A cell type atlas of the annelid *Pristina leidyi*

We first obtained a new transcriptome from adult *Pristina* individuals (mixed stages, mRNA) using Iso-Seq on the PacBio Sequel II platform. From 1,546,939 CCS reads our high quality data spans 111,961 full transcripts, with a mean length of 3,305 bp. This data was combined with previously published transcriptomic resources (Nyberg et al., 2012) using EvidentialGene (Gilbert, 2019), resulting in a transcriptomic resource containing 37,263 transcripts, which cover 96.3% of the metazoan BUSCO cassette. These were further flattened by taking only the primary transcript (discarding EvidentialGene “alternate” sequences) resulting in a transcriptomic resource comprising 29,807 contigs. We annotated these transcripts using eggNOG (Cantalapiedra et al., 2021) against the metazoan database and performing Diamond Blast (Buchfink et al., 2021) against a local version of the *nr* database (Supplementary File 1).

We obtained cell dissociations of adult mixed populations of *Pristina* containing intact organisms in all fissioning stages (Figure 1A). We used ACME dissociation, a recently developed protocol that fixes the cells early during the dissociation process (Garcia-Castro et al., 2021). ACME produces cell suspensions that are fixed and permeabilised, and are therefore ideal for performing SPLiT-seq (Rosenberg et al., 2018) or other single cell transcriptomic methods based on combinatorial barcoding. These methods tend to give less gene and UMI content per cell, but are in turn cost effective and allow obtaining tens of thousands of cell profiles at a low cost, maximising the detection of rarer cell types. We performed three independent SPLiT-seq experiments (Figure 1A) and sequenced them using the Illumina NovaSeq 6000 platform at 2x 150 bp read length. We processed these reads with our SPLiT-seq pipeline (Garcia-Castro et al., 2021; Rosenberg et al., 2018) and obtained a total of 80,387 cell profiles above 50 genes per cell. We aimed at identifying doublets using Scrublet (Wolock et al., 2019) and Solo (Bernstein et al., 2020), obtaining 2,870 and 2,554 cell barcodes respectively (Supplementary Figure 1A). The overlap of both methods was 458 cells, a significant (p = 0.006, above expected p = 0.001) but moderate result that indicates that doublet identification tools have a low level of agreement on such a complex dataset. We then examined whether doublets have a large influence on the overall quality of the data, for instance by creating cell clusters dominated by doublets. We processed the dataset containing doublets and performed Leiden clustering in 4 different resolutions (Supplementary Figure 1B) ranging from 46 to 88 clusters. This revealed that even at high clustering resolutions there are no clusters dominated by Scrublet and/or Solo doublets (Supplementary Figure 1C), showing that Scrublet and Solo-detected doublets are not a major source of clustering formation.

**Figure 1:**
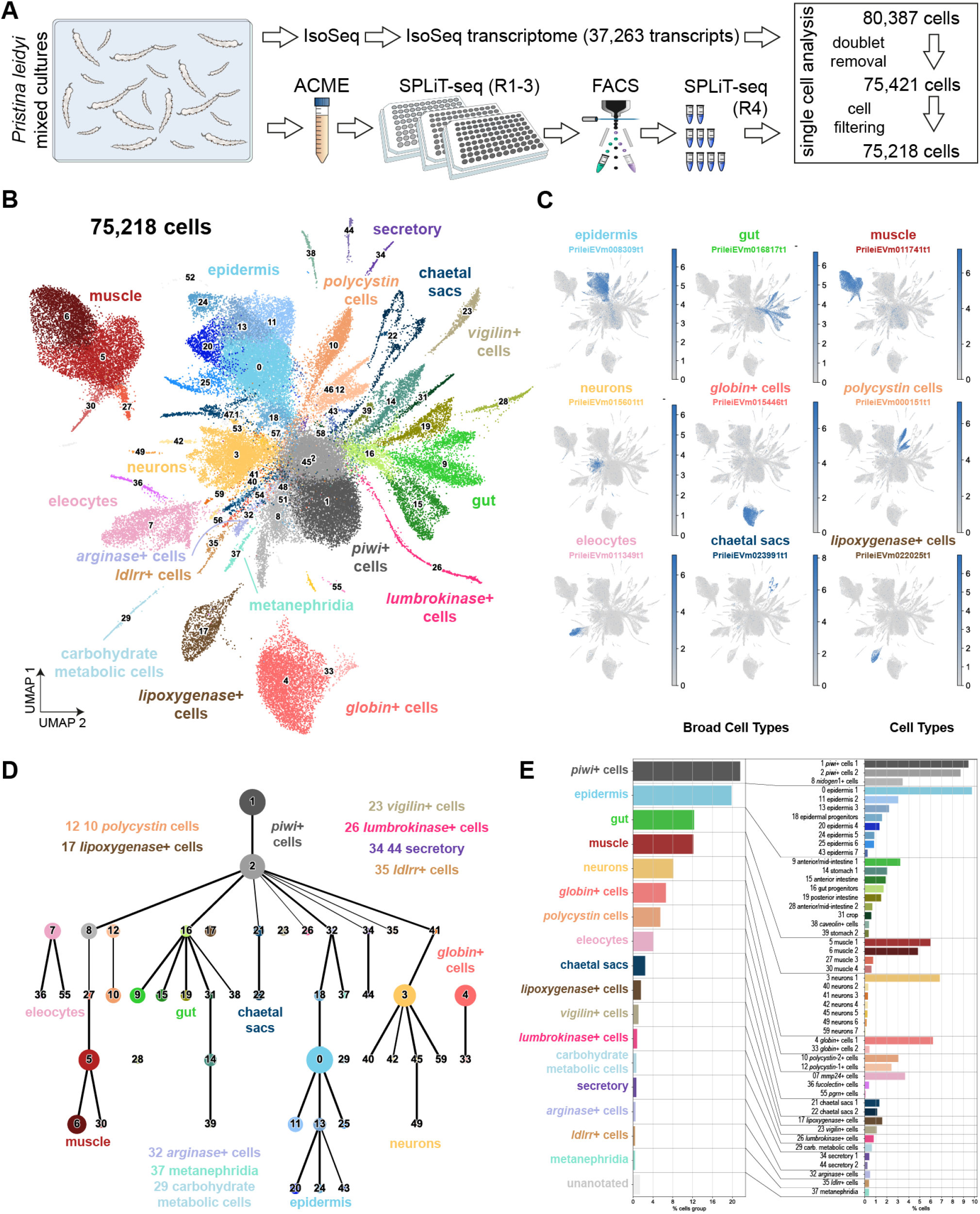
Single cell atlas of adult *Pristina leidyi* annelids. **A**: Experimental workflow. **B**: UMAP visualisation of the 75,218-cell *Pristina leidyi* single-cell transcriptomic cell atlas with clusters coloured according to their cell cluster classification. **C**: UMAP feature plots of markers of the major broad types, including epidermis, gut, muscle, neurons, *globin*+ cells, *polycystin* cells, eleocytes, chaetal sacs and *lipoxygenase*+ cells. **D**: Lineage reconstruction abstracted graph (PAGA) showing the most probable path connecting the clusters. Each node corresponds to the cell clusters identified with the leiden algorithm. The size of nodes is proportional to the amount of cells in the cluster, and the thickness of the edges is proportional to the connectivity probabilities. **E**: Cell cluster percentage at the broad cell cluster and the cell cluster levels.

We removed both doublet lists from the dataset, eliminating 4,966 cells (6.1%), a value conservatively above the ∼3% doublet expectation of our SPLiT-seq experiments. We explored the preprocessing parameter space with this 75,421-cell dataset, revealing that most conditions generated 40-60 reproducible clusters at resolution 1, but that a large number of Principal Components (PCs) and Highly Variable Genes (HVG) were needed to capture the full structure of the dataset (Supplementary File 2). We then processed the dataset eliminating the top expressed gene (PrileiEVm0194901t1), since this made up about 60% of our counts, likely a product of SPLiT-seq’s library amplification process. We further eliminated cells above 700 genes and 900 raw counts as likely doublets, obtaining a final dataset of 75,218 cells. We processed this dataset with 18,000 HVGsm, 45 neighbours and 105 PCs, obtaining a mean of 110 genes per cell and 397 counts per cell (Supplementary Figure 2AB). Despite this relatively low gene and UMI content, cell clustering with the Leiden algorithm (resolution 1.5) allowed us to robustly identify 60 cell clusters (Figure 1B, Supplementary Figure 2C-D, Supplementary Figure 3A) that are reproducible across parameter conditions (Supplementary File 2), and have highly specific markers (Figure 1C, Supplementary Figure 2E. We calculated marker genes for each cluster using the Wilcoxon and the Logistic Regression methods (Supplementary Files 3 and 4), which showed a high but not complete overlap. In some clusters, one of the two methods performed better than the other, but no method performed best in all clusters (Supplementary Figure 2F). We report the overlapping markers (Supplementary File 5). We noticed that some small clusters (ranging from 174 to 41 cells, 0.2% and 0.05% of the dataset respectively) shared the same UMAP space as clusters 1 and 2 (Supplementary Figure 3B) and expressed genes characteristic of these cell populations. These small clusters either represent subsets of the clusters 1 and 2 population or leftover doublets unidentified by Scrublet or Solo, and were flagged as “unannotated”. The total amount of unannotated cells is 1,048 (1.39% of the total dataset), a very small number that is within the order of magnitude of expected doublets (∼3%).

**Figure 2:**
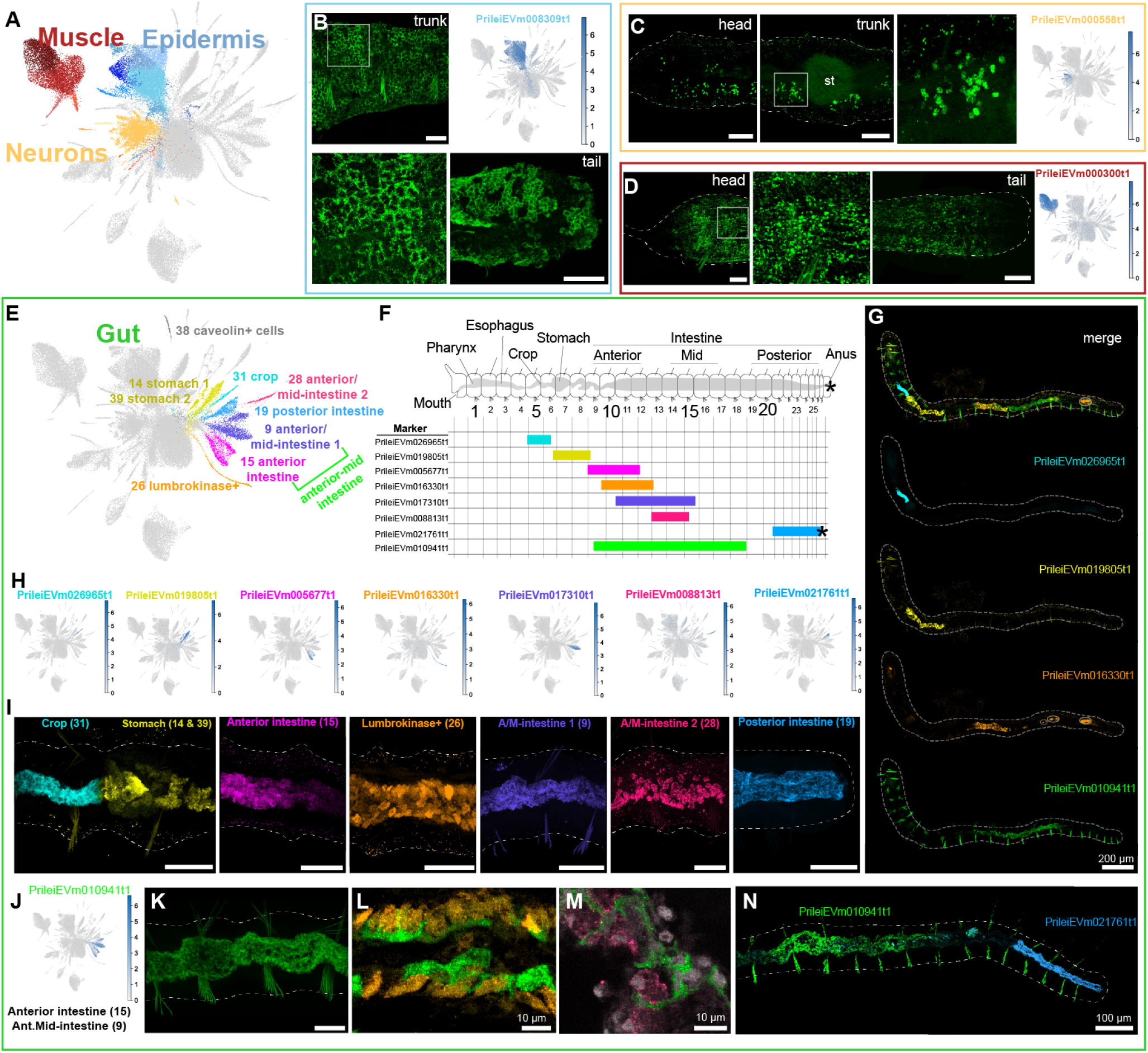
*Pristina leidyi* epidermis, muscle, neuron and gut organisation. **A:** UMAP visualisation highlighting Epidermis (blue), Muscle (red), and Neuron (yellow) clusters. **B:** Hybridization Chain Reaction (HCR) and UMAP feature plot of expression of epidermis marker PrileiEVm008309t1, showing extensive signal in the epidermal cells across the body, including trunk and tail regions. The bottom left panel is a close up of the top left panel. **C:** HCR and UMAP feature plot of expression of the neuronal marker PrileiEVm000558t1, showing groups of neuronal cell bodies across the worm’s body, including the head and the stomach region (trunk). The right microscopy panel is a close-up from the middle microscopy panel. **D:** HCR and UMAP feature plot of expression of the muscle marker PrileiEVm000300t1, showing expression along the worm, including the head and the tail regions shown as examples. The middle microscopy panel is a close-up from the left microscopy panel, evidencing muscle fibres. **E:** UMAP visualisation highlighting gut and associated clusters, including cell clusters 9, 14, 15, 19, 26, 28, 31 and 39. The colour code matches the colours in the microscopy images. **F:** General distribution of the marker expression representing each gut region along the worm. For detailed analyses, see Supplementary Figure 7. **G:** Example HCR sample showing 4 different markers simultaneously, but in distinct regions of the gut (blue, yellow, orange, green). Dashed line indicates the outline of the worm. Circles indicate background signal in the gut. **H:** UMAP feature plot of expression of diverse gut cluster markers. **I:** HCR expression of diverse gut cluster markers. In all images anterior is to the left, posterior is to the right. All images are lateral views. Note the strict border between the crop and stomach, where there is no co-expression of the markers. **J:** UMAP feature plot of expression of anterior and mid intestine marker PrileiEVm010941t1, with expression in cell clusters 9 and 15. **K:** HCR expression of PrileiEVm010941t1. **L:** HCR expression of PrileiEVm010941t1 (green) and *lumbrokinase*+ cell marker PrileiEVm016330t1 (orange), showing non overlapping expression in the same gut region. **M:** HCR expression of PrileiEVm010941t1 (green) and anterior/mid-intestine marker PrileiEVm008813t1 (pink), showing non overlapping expression in the same gut region. **N:** HCR expression of PrileiEVm010941t1 and posterior intestine marker PrileiEVm021761t1, showing non overlapping expression in distinct gut regions.In all panels, anterior is left, dorsal is up (unless otherwise noted). Tail in B is ventrolateral. All scale bars are 50 μm unless noted on the figure.

**Figure 3:**
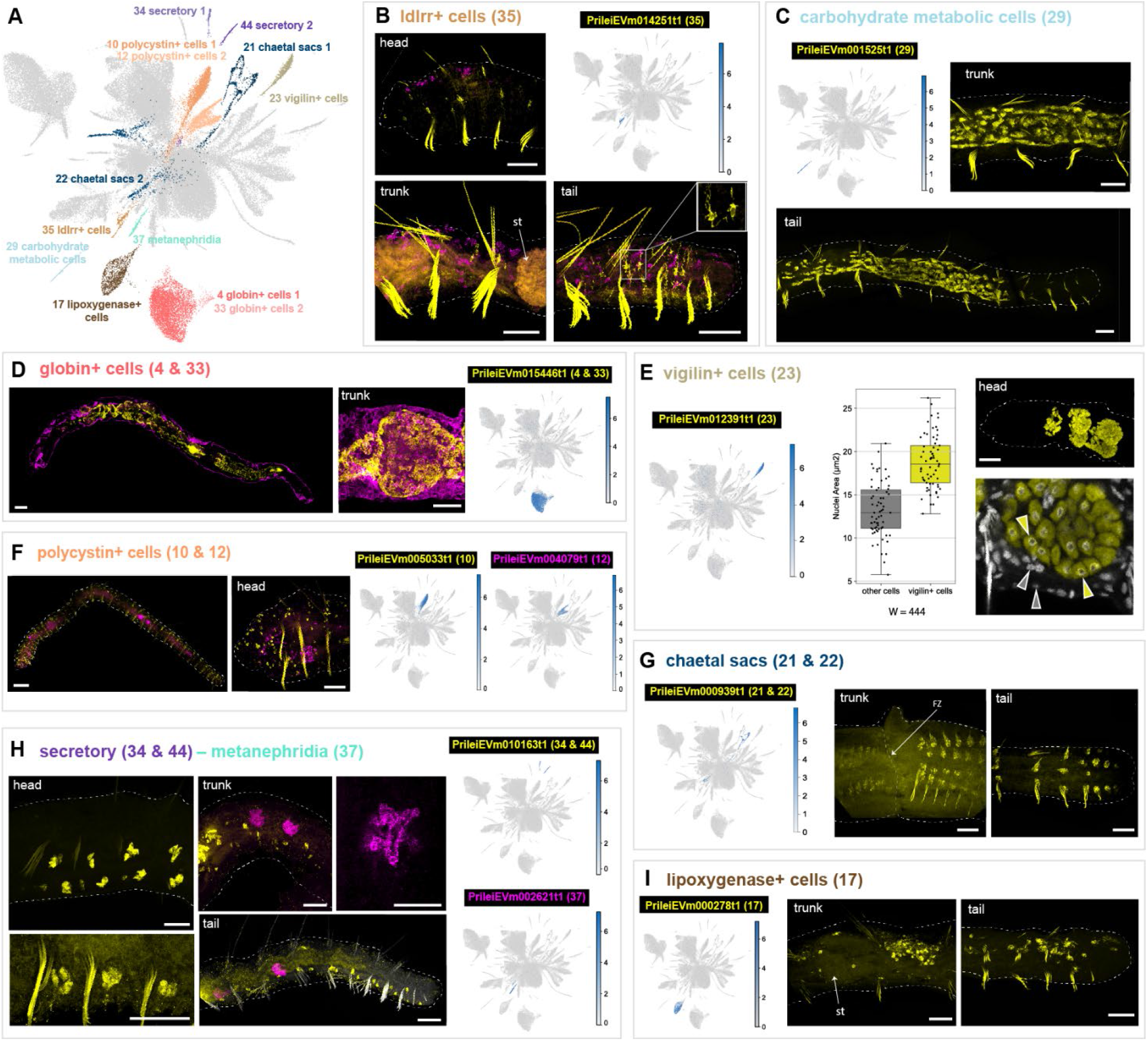
Annelid specific and novel cell types. **A:** UMAP visualisation highlighting annelid specific and novel cell types. **B:** HCR and UMAP feature plot of expression of the ldlrr+ cell marker (cluster 35) PrileiEVm014251t1, showing signal throughout the whole animal body (head, trunk and tail). Detail of the extensions of *ldlrr*+ cells is shown in the inset of the tail picture. Magenta counterstaining corresponds to eleocytes and *nidogen*+ cell marker (clusters 7 and 36) PrileiEVm005681t1. **C:** HCR and UMAP feature plot of *lipoxygenase*+ cell marker (cluster 17) PrileiEVm000278t1, showing cell expression in the trunk, around the stomach (st), and posterior parts of the animals. **D:** HCR and UMAP feature plot of *globin*+ cell marker (clusters 4 and 33) PrileiEVm015446t1, showing expression around the animal’s gut. Magenta staining corresponds to epidermal marker PrileiEVm008309t1. **E:** HCR and UMAP feature plot of *vigilin*+ cell markers (cluster 23) PrileiEVm012391t1, in the anterior part of the animal, together with the quantification of cell nuclei area. **F:** HCR and UMAP feature plots of *polycystin* cell markers (clusters 10 and 12) PrileiEVm005033t1 and PrileiEVm004079t1, showing the expression of *polycystin2*+ cells (yellow) segmentally repeated throughout the body wall of the animal and *polycystin1*+ (magenta) only in the anterior region. **G:** HCR and UMAP feature plots of chaetal sacs markers (cluster 21 and 22) PrileiEVm000939t1, showing the expression in the fission zone (FZ) and in the tail of the animal. **H:** HCR and UMAP feature plots of secretory (cluster 34 and 44) and metanephridia (clusters 37) markers PrileiEVm010163t1 and PrileiEVm002621t1. Expression of secretory cells are segmentally repeated mostly ventrally all along the whole body of the animal and metanephridia in some specific segments in midbody and posterior regions. **I:** HCR and the UMAP feature plot of carbohydrate metabolic cells marker (cluster 29) PrileiEVm001525t1, showing extensive signal in the posterior end of the animal. In all panels, anterior is left, dorsal is up (unless otherwise noted). Head in H is ventrolateral. All scale bars are 50 μm.

We then used PAGA (Wolf et al., 2019) to reconstruct differentiation trajectories. When we performed PAGA on the full dataset, small unannotated clusters connected other groups of clusters with unrelated markers (Supplementary Figure 4), reinforcing the idea that they could represent doublets.

**Figure 4:**
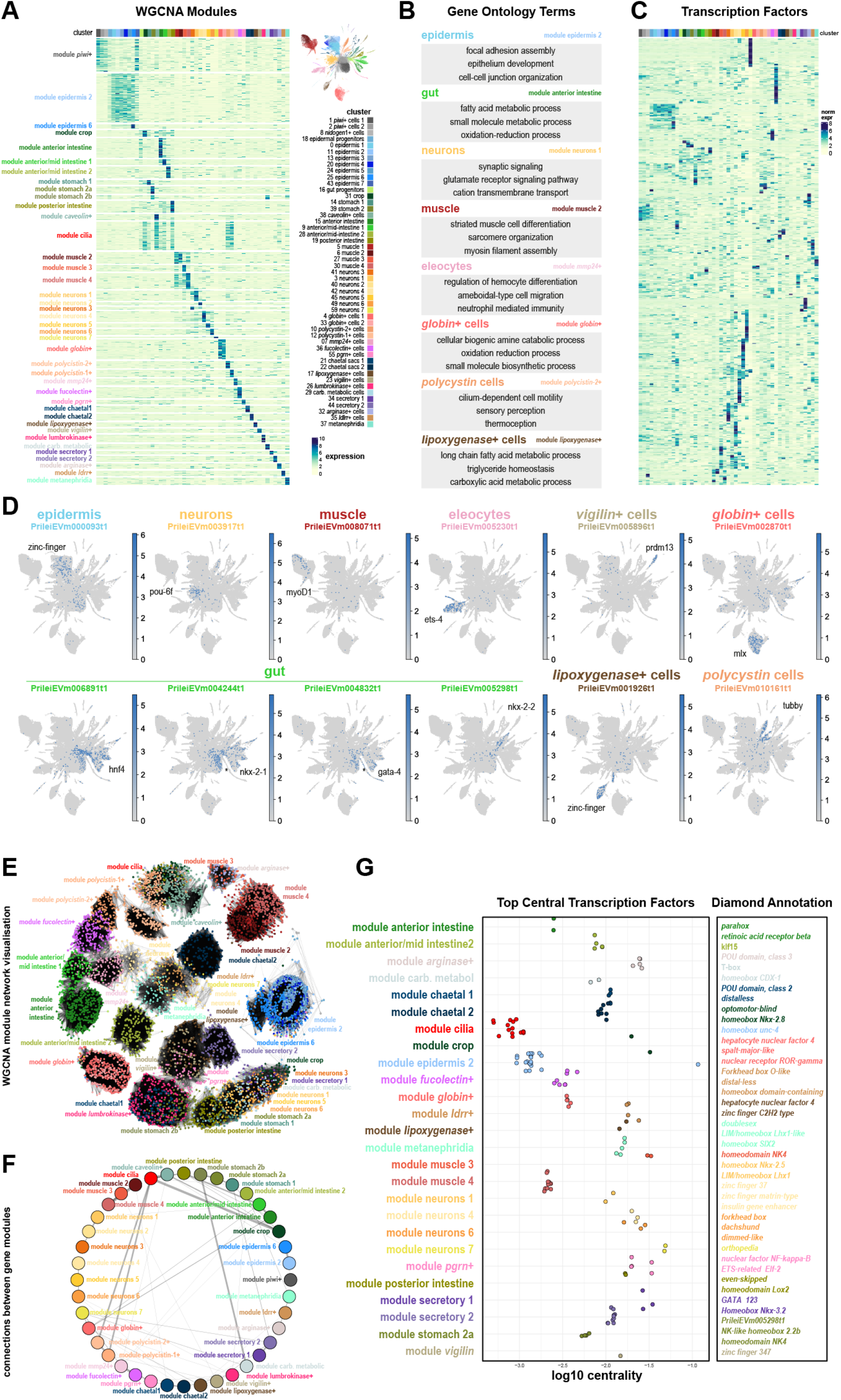
The transcriptional landscape of annelid cell type differentiation. **A:** Expression heatmap of 10,796 genes over 40 WGCNA modules of expression (rows), sorted by cluster expression (columns). **B:** Summary of Gene Ontology terms associated with example modules. **C:** Expression heatmap of 958 TFs (rows) sorted by cluster expression (columns). **D:** UMAP feature plots of TFs associated with individual cell types or broad types. The asterisk points to key differences between TFs. **E:** Network visualisation of WGCNA modules. In this visualisation, each gene is represented as a dot, coloured according to its cluster of highest expression, and edges represent gene coexpression. **F:** Module network visualisation summarising coexpression values between different modules, showing associations between different modules. **G:** Stripplot showing the top central TFs identified in WGCNA modules, and their annotations.

We then performed PAGA using only annotated clusters (Figure 1D). This lineage reconstruction allowed us to classify the broad cell types (Figure 1E). We also performed a co-occurrence analysis of cell type clusters (Levy et al., 2021), using the gene expression data of highly variable genes, summed at the cell cluster level. This analysis broadly confirmed our cluster groups (Supplementary Figure 5). We annotated individual cell types and group identities by considering their gene markers within the context of the published annelid cell type literature, the lineage reconstruction and the HCR characterization (Figure 1E, Supplementary Note 1).

**Figure 5:**
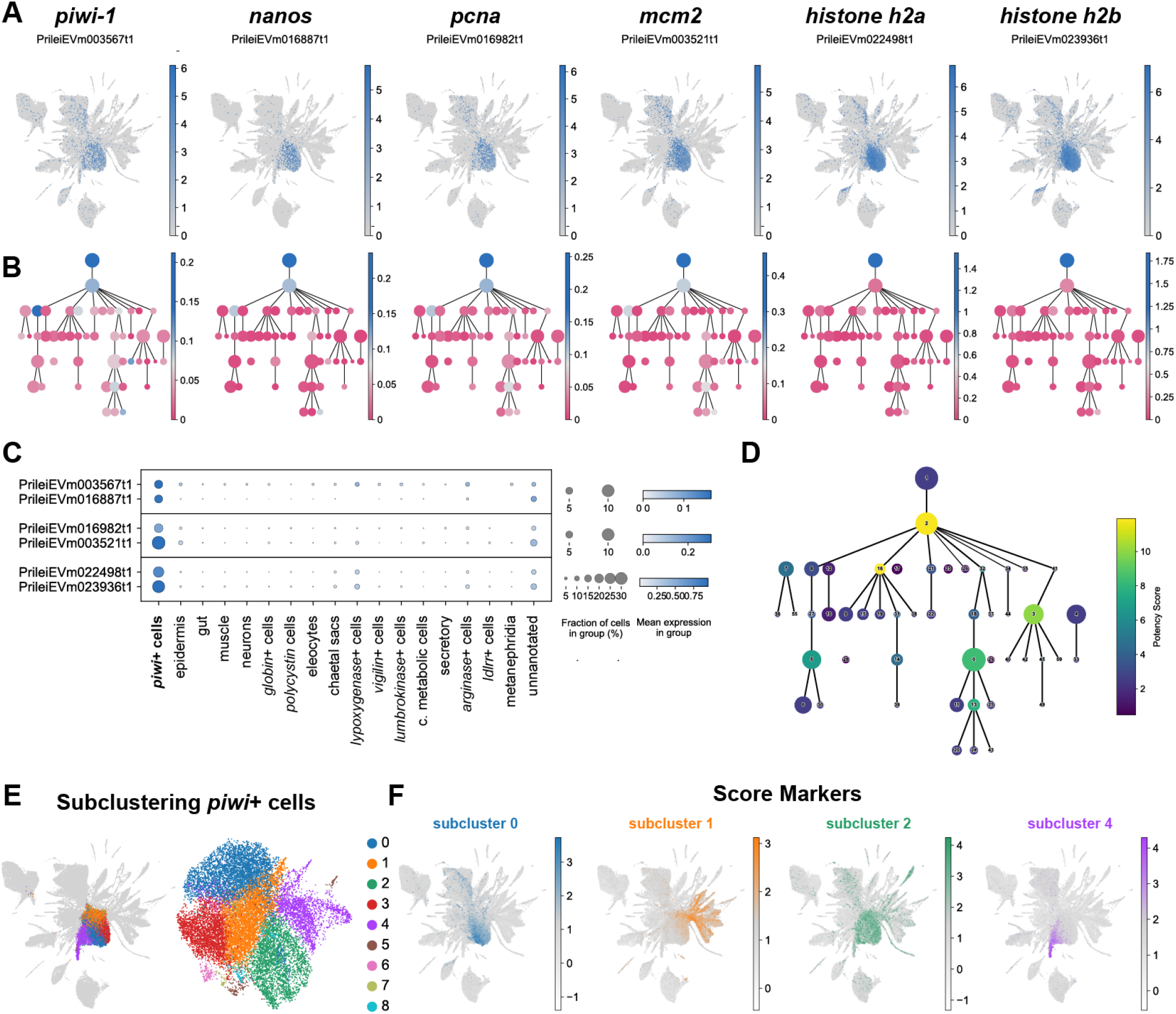
Pluripotent stem cell signature of *Pristina piwi*+ cells. **A:** UMAP feature plots of stem cell and proliferation markers, including *piwi, nanos, pcna, mcm2, histone h2a*, and *histone h2b*. **B:** PAGA feature plots of stem cell markers. The graph nodes represent the individual cell clusters and the colour intensity, from dark blue (high) to darkish pink (low), represents the expression of each marker. **C:** Dot Plot showing the expression of stem cell markers in broad cell types. The colour intensity of the dots represents the mean expression and the size of the dot represents the fraction of cells expressing the marker. Due to the highest expression of histone genes each pair is presented in an individual panel, with its own maximum colour intensity and size. **D:** PAGA plot coloured according to potency score, ranging from dark blue (low) to yellow (high). **E:** UMAP visualisation of the 16,247 cells of *piwi*+ clusters 1, 2 and 8, in their original UMAP embedding (left) and their subclustering UMAP embedding (right) with clusters coloured according to their cell subcluster classification. **F:** UMAP score plots of markers of *piwi+* subclusters 0, 1, 2 and 4, showing differential expression in gut, epidermis, *lumbrokinase*+, *vigilin*+ and *nidogen*+ cell clusters.

### HCR *in situ* hybridisation validates epidermal, muscular and neuronal cellular identities in *Pristina leidyi*

We developed a multiplexed HCR *in situ* hybridization protocol for *Pristina* and validated most cluster identities using specific cluster markers (Supplementary File 6, Supplementary Figure 6). First, we characterised major cell types such as epidermis, neurons, and muscle (Figure 2A). We characterised the epidermis based on the expression of PrileiEVm008309t1. This marker was found all across the outer body wall and along the entire length of the worm’s body (Figure 2B). Neural populations were defined based on the expression of *synaptotagmin* (PrileiEVm012030t1) and validated by *in situ* expression of PrileiEVm000558t1, a broad neuronal marker. We found staining anteriorly in the head and in ventral clusters of neurons across the body, reminiscent of previously-published immunostainings for neurons (Zattara and Bely, 2011, 2015) (Figure 2C). Finally, we characterised muscle clusters based on their high expression of muscle markers (e.g. *myosin, tropomyosin, troponin*). The *in situ* hybridization of one of these markers, the myosin heavy chain homolog gene PrileiEVm000300t1, revealed longitudinal muscle fibres extending along the surface of the animal (Figure 2D).

**Figure 6:**
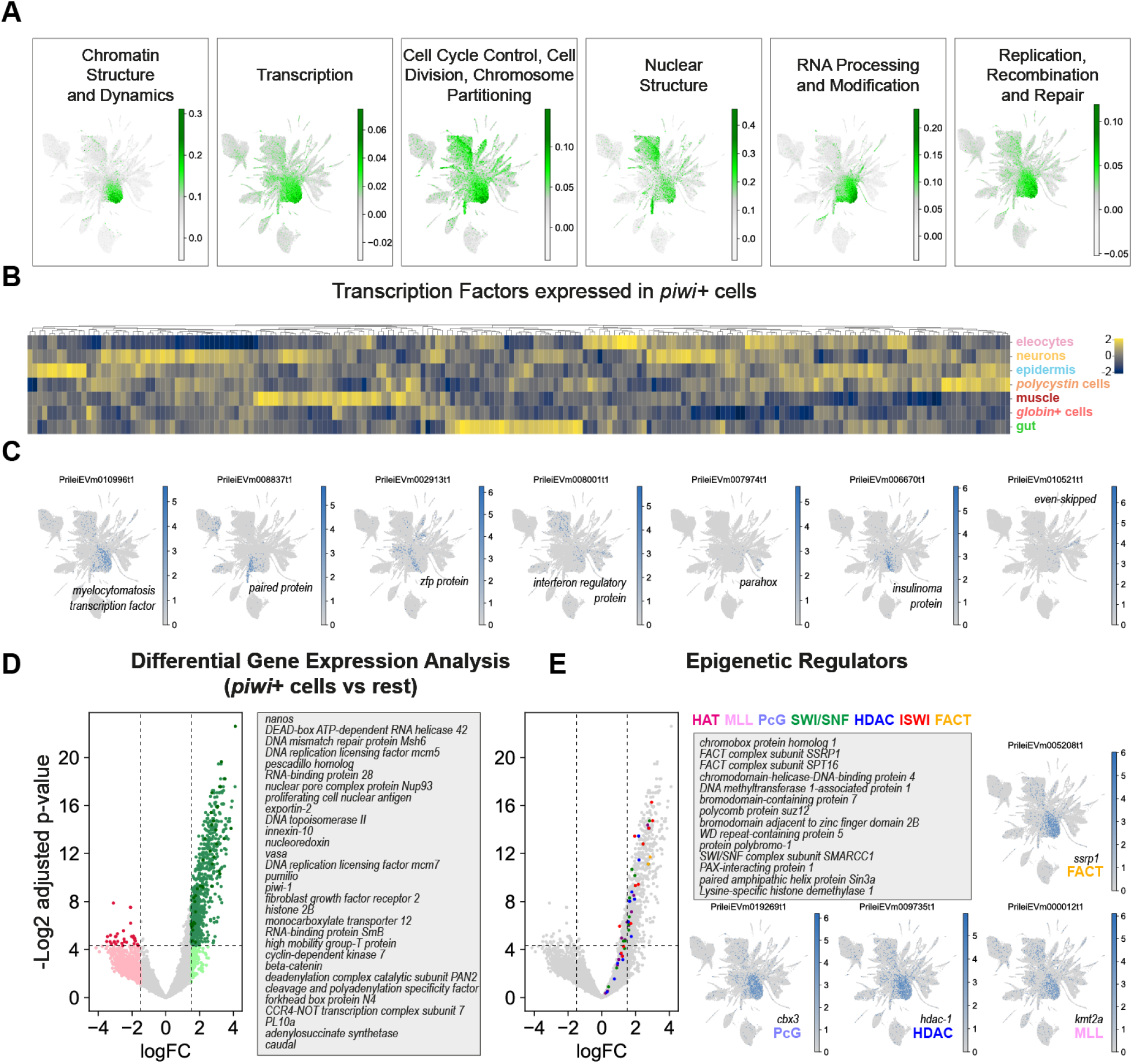
Transcriptomic profile of annelid *piwi*+ cells. **A:** UMAP visualisation of scored gene expression of COGs in *piwi*+ cells. **B:** Expression heatmap of 200 top TFs expressed in *piwi*+ cells and their expression in the main broad cell types, showing several clusters of TFs expressed in both *piwi*+ cells and one or more broad type. **C:** UMAP feature plots of example TFs coexpressed in *piwi*+ cells and other broad types. **D:** Limma differential gene expression analysis of *piwi*+ cells (clusters 1, 2 and 8) against all other cell types, with annotation lines for the examples indicated in dark green. **E:** Detail of annotated epigenetic regulators and their expression enriched in *piwi*+ cells, and example UMAP visualisations of representative epigenetic factors.

### High antero-posterior regionalisation of the *Pristina leidyi* gut

We identified 10 gut and gut-associated cell clusters (Figure 2E), and visualised the localization of their markers using HCR (Figure 2F-N). These analyses revealed that *Pristina* has a complex gut organisation with specific molecular regions and cell types along the entire antero-posterior axis. Some of these regions were restricted to as few as 2 segments, such as the crop region (cluster 31) which always occurred in segments 5-7 (Figure 2F-I; Supplementary Figure 7). Some gut markers exhibited consistent and sharp borders. In all samples analysed, crop and stomach (clusters 14 and 39) always had a sharp border with cells at this boundary expressing either the crop marker or the stomach marker, but never both (Figure 2H’). Similarly, the most posterior gut marker (PrileiEVm021761t1) was always expressed up until the anus, largely coincident with a region with long cilia in the posterior intestine. In contrast, some markers were expressed in broadly the same regions of the gut, but their cellular expression did not overlap (Figure 2L, M), indicating the presence of distinct cell types in those regions. Among them, we found a cell cluster with high expression of lumbrokinase enzymes (cluster 26, Figure 2I), identifying the cell type that produces this previously described fibrinolytic enzyme (Altaf et al., 2021). The expression of intestine markers along the anterior-posterior axis tended to be proportional to the worm’s overall length, suggesting that these gut regions expand proportionally as the worms grow longer (Figure 2F, Supplementary Figure 7). These results show that our single cell data resolves the complex gut organisation of *Pristina*, with distinct molecular regions along the anterior-posterior axis and several regionally-specific cell types.

### Single cell transcriptomics reveals a wealth of annelid cell types and novel cell types

We then aimed to characterise the remaining set of clusters (Figure 3A). We identified novel cell types as well as annelid cell types. Among the novel cell types, we identified a population of *ldlrr*+ cells (cluster 35), which are distributed throughout the animal (Figure 3B) and have a morphology with numerous extensions (Figure 3B, inset), reminiscent of astrocytes. Furthermore *ldlrr+* cells express PrileiEVm006872t1, a homolog of the intermediate filament gliarin (Xu et al., 1999). On the other hand, cluster 29 corresponded to cells located in the posterior gut, up to 3-4 segments before the tail end. This population was called carbohydrate metabolic cells due to the expression of Krebs cycle and mitochondrial enzymes (Figure 3C). We did not find previous descriptions of these populations in the annelid literature and therefore consider these novel cell types.

We also found annelid types that likely represent cell types previously described in annelids at the morphological or molecular level. For instance, we found a prominent cell population (clusters 4 and 33) that expressed several extracellular globins (Supplementary Files 3-4). These transcripts have been characterised in the annelid species *Platynereis dumerilii* (Song et al., 2020), where they are expressed in transverse trunk vessels and parapodial vessels. In contrast, we found that *globin*+ cells in *Pristina* occupy large areas in the vicinity of the gut (Figure 3D). Then, we identified a cluster (23) marked by the expression of *vigilin*, an RNA-binding protein important for chromosome stability and cell ploidy (Cheng and Jansen, 2017). For instance, the *Drosophila vigilin* homologue, DDP1, is found in polytene chromosomes (Cortes et al., 1999). *Pristina vigilin*+ cells are located in three large bulbs in the head segments of the worm (Figure 3E). Similar cells have been described morphologically as pharyngeal glands in some other *Pristina* species, as well as in several oligochaetes (Collado and Schmelz, 2000; Stephenson, 1922). Interestingly, this cluster showed a higher number of RNA counts per cell (Supplementary Figure 2D). We wondered if this was a technical artefact or a biological observation instead, with *vigilin*+ cells being larger polyploid cells, for instance. We quantified the cell nuclei area of *vigilin*+ cells and determined that their size is significantly larger than that of other cells (Figure 3E). Furthermore, we found a transcript encoding a *mucin* gene in the marker list. Together, these data suggest that *vigilin*+ cells are polyploid cells that function as salivary glands.

We then examined two prominent and abundant (3.1% and 2.4%) clusters marked by *polycystin* genes, a family of genes associated with cilia (Esarte Palomero et al., 2023). We found that *polycystin2*+ cells (cluster 10) were segmentally repeated in the body wall of the worm (Figure 3F), likely corresponding to sensory cells equipped with ciliary tufts (Gelder, 1984; Giere and Rhode, 1987). In contrast, *polycystin1*+ cells (cluster 12) were enriched in the head segments (Figure 3F). We also found that clusters 21 and 22 corresponded to the chaetal sacs (Figure 3G), which were marked by the expression of a transcript encoding a chitin synthase protein (PrileiEVm000573t1). Clusters 34 and 44 corresponded to segmentally repeated cells all along the body of the animal, with a likely secretory function (Figure 3H), based on the expression of a conotoxin protein (PrileiEVm010163t1). Cluster 37 corresponded to the metanephridia with a clear tubular structure (Figure 3H). Finally, *lipoxygenase*+ cells (cluster 17) were characterised by the expression of numerous lipoxygenase enzymes (Supplementary Files 3-4). These fatty acid-peroxidising enzymes have been implicated in a range of immune, signalling and metabolic functions (Mashima and Okuyama, 2015). *Lipoxygenase*+ cells are large cells distributed throughout the AP axis of the animal (Figure 3I), and could correspond to the previously described chloragocytes (Schenk and Hoeger, 2020). Altogether, these observations identified several annelid cell types such as the chaetal sacs, the *vigilin*+ cells, the *lipoxygenase*+ cells, the *polycystin* or the *globin*+ cells, but also revealed previously unknown cell types such as the *ldlrr*+ cells and the carbohydrate metabolic cells, with function and homologies that are yet to be explored. Thus, our single cell dataset reveals new biological insights into blood-related cell types, metabolic cell types and polyploidisation, opening up numerous research avenues for annelid researchers and for the investigation of the evolution of cell types.

### The transcriptional landscape of annelid adult cell differentiation

We then investigated the specific gene expression patterns of each *Pristina* cell type. Given the low UMI and gene counts of our combinatorial single cell dataset, we used a pseudobulk approach, aggregating raw reads coming from all cells in each cluster. This allowed us to quantify a mean of 11,117 genes per cluster (Supplementary Figure 8A). We then used Weighted Gene Coexpression Network Analysis (WGCNA) to identify genes with correlated expression patterns. We identified 10,796 genes distributed over forty modules of specific gene expression, broadly corresponding to most cell clusters identified (Figure 4A, Supplementary File 7). We used Gene Ontology analysis to extract biologically relevant terms for each cell type (Figure 4B, Supplementary File 8). One module corresponded to genes expressed in several cell types but enriched in cilia-related GO terms (Figure 4A, Supplementary File 8). To assess the potential regulatory layer underlying this transcriptional landscape, we focused our attention on Transcription Factors (TFs). We annotated 958 *Pristina* TFs (see Methods, Supplementary File 9, Supplementary Figure 8B-E) and identified cell type-specific expression of dozens of TFs (Figure 4C), including well-known markers or regulators of several cell types, such as a *pou-6f* gene in neurons and a *myoD* gene in muscle (Figure 4D). This included rich regulatory detail, for instance in the gut, with *hnf4* and *nkx-2-1* TFs broadly expressed in gut clusters, but excluded from *lumbrokinase*+ cells, and a *gata-4* TF with similar expression, but including the*lumbrokinase*+ cells (Figure 4D, asterisk). This analysis allowed us to obtain insight for the first time into our annelid specific and novel cell types, identifying TFs specific to eleocytes, *vigilin*+ cells, *globin*+ cells, *lypoxygenase*+ cells and *polycystin* cells. Next, we used graph analysis to visualise *Pristina* WGCNA modules as a network, and identified several connected components that reliably match the WGCNA modules and roughly recapitulate cell type-specific gene expression (Figure 4E). This allowed us to explore the relationships between gene modules by computing the number of cross-connections between pairs of modules. This highlighted connections between cilia, esophagus and *polycystin* cells suggesting the presence of cilia in such cell types (Figure 4F), among other connections. Lastly, we were also able to detect and calculate the centrality of the different TFs present within each sub-network, revealing further putative TF regulators of each differentiated cell type (Figure 4G), including multiple homeobox, forkhead and zinc-finger TFs among others. Overall, our analysis reveals the transcriptomic landscape of annelid adult cell type differentiation and showcases the power of ACME and SPLiT-Seq to unravel the TF expression underlying cell type differentiation in uncharacterised species.

### *Piwi*+ cells are abundant and heterogeneous and have a pluripotent stem cell signature

Next we focused on identifying and characterising putative stem cell populations in *Pristina. Piwi*+ cells have been described previously in this species (Özpolat and Bely, 2015; Özpolat et al., 2016) but their transcriptional profiles, cellular properties and differentiation capacities remain largely unknown. We found that the central clusters 1, 2 and 8 highly expressed *piwi-1* and *nanos* (Figure 5A, left panels). These clusters constitute 21.6 % of our dataset (Figure 1E), indicating that *piwi*+ cells are an abundant cell type in *Pristina*. The representation of *piwi*+ cells in our 3 independent SPLiT-seq experiments ranged from 13.0% to 33.8%. This indicates that the percentage of *piwi*+ cells is highly variable, potentially reflecting differences in the average nutritional state (and therefore growth and fission states) of worms in our three experiments. We then analysed the expression of the proliferation markers *pcna* and *mcm2*, as well as histones *h2a* and *h2b*. These genes were very highly expressed in central clusters 1 and 2 (Figure 5A, right panels). Moreover, our PAGA analysis revealed that most differentiated cell types were connected to *piwi*+ cells by differentiation trajectories (Figure 1D, Figure 5B), including epidermis, muscle and gut, suggesting these cells are a pluripotent population. While we observed expression of proliferation markers in other clusters (Figure 5A-C), clusters 1, 2 and 8 concentrate most of their expression (ranging from 70.0% to 82.1% of all reads mapped to these features), indicating that *piwi*+ cells are the major proliferative cell type in *Pristina*. To model the developmental potency of *Pristina piwi*+ cells we calculated the potency score (Plass et al., 2018). This graph analysis metric evaluates the normalised degree of each node of the abstracted PAGA graph as an estimation of the number of differentiation trajectories that connect to it. The highest potency score in our abstracted graph was attained by *piwi*+ cell cluster 2 (Figure 5D), strongly suggesting that *piwi+* cells are pluripotent stem cells. Clusters 16, 0, 3 and 13 also attained high potency scores, as they were connected by the PAGA analysis to several gut, neuronal, and epidermal clusters, reinforcing the notion that these are progenitors of their differentiated types (Figure 5D). Pluripotent cells in other organisms have been shown to be heterogeneous (Hulett et al., 2022; Martinez Arias and Brickman, 2011; Messmer et al., 2019; Mohammed et al., 2017; van Wolfswinkel et al., 2014), consisting of mixtures of cells that co-express stem cell markers and transcripts that are characteristic of the cell types that they will differentiate into. To elucidate if *Pristina piwi*+ cells are heterogeneous we performed a subclustering analysis (Figure 5E) and scored the markers obtained in this analysis (Figure 5F). *Piwi*+ subclusters contained markers of several differentiated types, including gut and epidermal cells (Figure 5F, Supplementary Figure 9). A further subcluster showed expression in the *nidogen*+ cells, which are connected by PAGA with muscle. These results show that *piwi*+ cells are heterogeneous and co-express stem cell markers plus markers of the several cell types that they may differentiate into. Altogether, these analyses showed that *piwi*+ cells in *Pristina* are a heterogeneous cell population with a pluripotent stem cell signature.

### Chromatin regulators are conserved markers of annelid *piwi*+ cells

We then sought to understand the transcriptomic profile of *Pristina piwi*+ cells. We first annotated Clusters of Orthologous Groups (COGs) (Cantalapiedra et al., 2021; Galperin et al., 2015) across the species transcriptome, and scored their expression in the single cell dataset. We found that *piwi*+ cells were enriched in COGs related to chromatin, transcription, cell cycle, nuclear structure, RNA biology and DNA repair (Figure 6A, Supplementary File 10). This analysis allowed us to look for the first time at the transcriptomic features of *piwi+* cells in annelids. We then sought to understand their transcriptional regulation by identifying their highly expressed TFs (see Methods). Interestingly, a high proportion of TFs highly expressed in *piwi+* cells were also highly expressed in one or more differentiated cell type groups (Figure 6B, Supplementary File 11). Examples of these included TFs expressed in *piwi+* cells and other cell types such as *vigilin*+ cells, muscle, *polycystin* cells, gut and epidermis (Figure 6C, Supplementary File 12). This finding is highly consistent with the specialised or lineage committed stem cell concept and strongly suggests that these TFs are those that prime and regulate differentiation to their correspondent cell types. We then used limma (see Methods) to obtain the full transcriptional profile of *piwi*+ cells and identified a list of 735 significantly enriched transcripts (t-test with empirical Bayesian moderation of standard errors, false discovery rate by Benjamini-Hochberg, p-value < 0.05, logFC>2, Figure 6D, Supplementary File 13). Notably, this list included stem cell regulators such as *piwi, vasa, nanos* and *pumilio*, known to be expressed in pluripotent stem cells across the animal tree of life, as well as in germ cells (Gehrke and Srivastava, 2016; Juliano et al., 2010; Lai and Aboobaker, 2018; Solana, 2013). Also notable in the *Pristina piwi*+ cell transcriptome were cell cycle regulators, DNA repair proteins and purine synthesis enzymes, also consistent with other pluripotent stem cell transcriptomic profiles (Alie et al., 2015; Labbe et al., 2012; Önal et al., 2012; Solana et al., 2012). A very prominent feature of *Pristina piwi*+ cells was the expression of epigenetic regulators and/or chromatin remodelers. To corroborate this feature, we used BLAST to search for homologs of the most important chromatin remodelling complex components, including the HAT, MLL, PcG, SWI/SNF, HDAC, ISWI, and FACT complexes (Marakulina et al., 2023; Medvedeva et al., 2015). We identified 156 *Pristina* transcripts encoding these (Supplementary File 14), and found them all enriched in *piwi*+ cells (Figure 6E). Similar to human and planarian pluripotent cells (Dattani et al., 2019; Labbe et al., 2012; Önal et al., 2012), this shows that epigenetic regulation is a conserved feature of animal pluripotent cells. Taken together, our data strongly suggest a model where post-transcriptional and epigenetic regulators control stem cell maintenance and pluripotency, and a panoply of TFs prime these to differentiate into multiple cell types.

## Discussion

In this study, we report a new transcriptome and single cell atlas of adult *Pristina leidyi*, an annelid species capable of extensive adult cell type generation and regeneration. Our datasets provide an unprecedented perspective on adult cell type differentiation in annelids and their pluripotent cellular sources. The adult cell type atlas of *Pristina* reveals the cellular identities that make up adult annelids. We uncover ∼50 distinct cell clusters and validate many of them using a newly developed multiplexed HCR *in situ* approach. Therefore, our data show that ACME and SPLiT-seq are robust approaches for cell atlas studies, in spite of the relatively low UMI content. The capacity of cost-effectively profiling tens of thousands of fixed cells, combined with single cell analysis and pseudobulk approaches, allows deriving rich transcriptional and regulatory profiles.

Our data reveal well-known cell types such as epidermis and muscle, a complex organisation of the annelid gut, as well as multiple annelid-specific cell types and novel cell types. We studied their distribution patterns along the body as well their transcriptional and regulatory profiles, including gene expression modules and transcription factors. These new cell types offer key novel information to the field of cell type evolution, a field that has been reinvigorated by single cell transcriptomics. For instance, we found a *vigilin*+ cell type that expresses mucins and is localised in the head region, strongly indicating that these are *Pristina* salivary glands. Interestingly, *vigilin* has been implicated in polyploidisation events (Cheng and Jansen, 2017) and we show that *vigilin*+ nuclei have larger sizes, consistent with a plausible polyploidisation. This resemblance suggests a deep homology of salivary glands in arthropods and annelids. Alternatively, convergent evolution could have driven the co-option of similar mechanisms for a similar function. Cell types such as the *globin*+ cells and the eleocytes could be representatives of blood types related to haemocytes in other species and vertebrate blood cells. On the other hand, cell populations such as the *ldlrr*+ cells and the carbohydrate metabolic cells have no known homologue cell types in other groups. Future studies will focus on transcriptomic comparisons of these cell types to elucidate their evolution.

The differentiation of the majority of these cell types can be reconstructed from the *piwi*+ cell population in *Pristina*, which shows hallmarks of pluripotency. First, it expresses conserved RNA-binding proteins such as *vasa, nanos, pumilio* and *piwi*. These transcripts have been found in pluripotent stem cells in sponges, cnidarians, acoels, planarians, colonial ascidians and other organisms, as well as the germ line of most animals (Gehrke and Srivastava, 2016; Juliano et al., 2010; Lai and Aboobaker, 2018; Solana, 2013). Second, differentiation trajectories from *piwi*+ cells to a broad collection of cell types can be reconstructed using lineage reconstruction algorithms (Tritschler et al., 2019; Wolf et al., 2019). These exploit the presence of cells captured along their differentiation process, with transcriptomes intermediate between those of stem cells and differentiated cells. The concept of germ layers is key to the definition of pluripotency, but it is difficult to apply to asexually reproducing animals, where all cell types are differentiated from adult populations rather than embryonic germ layers. We therefore apply the pluripotency definition based on the reconstructions to broadly different cell types, including epidermis, muscle and gut, known to originate from distinct embryonic germ layers in annelids (Ackermann et al., 2005; Goto et al., 1999; Meyer et al., 2010; Özpolat et al., 2017; Smith and Weisblat, 1994; Weisblat and Shankland, 1985). Third, the *piwi+* cell cluster is heterogeneous and includes subpopulations that express stem cell markers and markers of differentiation to broad cell type groups or individual types. This is consistent with the idea of lineage committed stem cells that have already started their differentiation process (Hulett et al., 2022; Martinez Arias and Brickman, 2011; Messmer et al., 2019; Mohammed et al., 2017; van Wolfswinkel et al., 2014). Our analysis reveals rich regulatory information, including dozens of transcription factors that are expressed in *piwi+* cells and in a given set of differentiated types. Fourth, our analysis uncovers a prominent expression of epigenetic regulators and chromatin remodelers in *piwi*+ cells. This is a signature of pluripotency in human (Gaspar-Maia et al., 2011; Schlesinger and Meshorer, 2019) and planarian stem cells (Önal et al., 2012; Solana et al., 2012), but is still understudied in other models. Our data reveals that this is a hallmark of pluripotency in annelids. Importantly, *piwi*+ cells concentrate most of the expression of cell cycle related transcripts but we cannot rule out that other cell types are able to undergo cell division. For instance, some epidermal clusters also express cell proliferation markers and histones. The expression of epigenetic regulators is however very restricted to *piwi*+ cells and therefore can be considered a hallmark of pluripotency. Altogether, our study reveals that a *piwi*+ cell population with a pluripotent stem cell signature underlies adult cell type generation in annelids.

## Methods summary

### *Pristina leidyi* culture and maintenance

*Pristina leidyi* culture was originally obtained from Carolina Biological Supply (Bely and Wray, 2001). Specimens were cultured in 1% filtered artificial seawater, and fed with 0.03g/L of dried spirulina powder every 2 weeks. Under these conditions, worms reproduce continuously by paratomic fission.

### Iso-Seq

*Pristina leidyi* of mixed conditions, including fissioning animals, were used. Total RNA was extracted using Trizol. Total RNA was provided to the Earlham Institute Genomics Pipelines Group, Norwich, UK, using a PacBio Sequel II SMRT cell, and sequenced (8M, v2, 30hr Movie). Iso-Seq3 analysis was performed by the provider. A total of 3,932,103 CCS reads were captured across the samples on the cell, with 1,546,939 assigned to *Pristina leidyi*. These were classified and clustered, resulting in 54,350 high-quality isoforms. The output of Iso-Seq was combined with a previous *Pristina leidyi* transcriptome (Nyberg et al., 2012).

### ACME dissociation

ACME was performed as previously described (Garcia-Castro et al., 2021) with some modifications, detailed in the Supplementary Methods. For each sample, we added ∼120 *Pristina leidyi* worms at mixed stages (including fissioning animals) to a 15 mL Falcon tube. Depending on the batch, we used animals at different starvation conditions: 12 days (library 12), 4 days (library 21) or 7 days (library 30).

### SPLiT-seq

SPLiT-seq was performed as previously described (Garcia-Castro et al., 2021) with some modifications, detailed in the Supplementary Methods. Sequencing was provided by Novogene (China). A total of 124,349,078 (12_1), 135,900,060 (12_2), 410,765,606 (21_1), 833,784,688 (21_2), 807,486,658 (21_3), 643,285,668 (30_2), 627,640,824 (30_3), 711,569,074 (30_4), 725,038,254 (30_5) reads were sequenced. The data was mapped using our Iso-Seq transcriptome as a reference. The Iso-Seq transcriptome of *Pristina leidyi* assembled as described above was created to have a reference database for read mapping. The SPLiT-seq read processing pipeline was performed as previously described (Garcia-Castro et al., 2021). A detailed description is available in the Supplementary Methods.

### Data processing and analysis

We used Scrublet (Wolock et al., 2019) and Solo 0.1 (Bernstein et al., 2020) to identify potential doublets. We then seed to optimise the analysis parameters by iteratively running the analysis with different minimum genes counts, maximum number of genes, maximum number of counts, number of top highly variable genes, number of neighbours, number of principal components, and leiden clustering resolution arguments. We processed the final dataset with conditions learned from our parameter space exploration. For the PAGA analysis we removed unannotated clusters since preliminary analyses indicated that these small clusters interfere with lineage reconstruction. For CPM calculation, raw counts were extracted with a custom Python script (see project repository). For the Co-ocurrence analysis, we used the function ‘treeFromEnsembleClustering() from the code provided by Levy and collaborators (Levy et al., 2021). We performed the WGCNA (Langfelder and Horvath, 2008) using a subset of the CPM table genes with CV > 1 and softPower 5 estimated after visualising the Scale-Free Topology Model Fit. Differential Gene Expression Analysis was performed using the edgeR (Robinson et al., 2010) and limma (Ritchie et al., 2015) R packages, and the pseudo bulk count matrix Full code and documentation is available at the project repository.

### Transcription factor annotation and analysis

The resulting TransDecoder-translated proteome of *Pristina leidyi* was queried for evidence of Transcription Factor (TF) homology using (i) InterProScan (Jones et al., 2014), PANTHER (Thomas et al., 2022), and (ii) SUPERFAMILY (Gough et al., 2001; Pandurangan et al., 2019) domain databases with standard parameters, (iii) using BLAST reciprocal best hits (Moreno-Hagelsieb and Latimer, 2008)against swissprot transcription factors (UniProt, 2023), and (iv) using OrthoFinder (Emms and Kelly, 2019) with standard parameters against a set of model organisms. For Transcription factor analysis, the CPM table was subset to retrieve the *Pristina leidyi* TFs.

Full code and documentation is available at the project repository.

### HCR *in situ* hybridization and confocal imaging

For Hybridization Chain Reaction (HCR), previously published protocols (Kuehn et al., 2022) were used with mainly modifications for *Pristina leidyi* fixation and 1st day of the protocol, based on the species colorimetric *in situ* hybridization protocols (Özpolat and Bely, 2015). The entire protocol can be accessed in https://github.com/BDuyguOzpolat/Pristina_leidyi-protocols. Confocal imaging was carried out using Zeiss LSM710 and LSM780 microscopes at the microscopy facility at Marine Biological Laboratory. For each set of HCRs, control tubes were included.

## Availability of Data and Materials

The datasets supporting the conclusions of this article are available in:

Code: https://github.com/scbe-lab/pristina-cell-type-atlas

GEO: GSE230505

## Acknowledgments

We thank Robert Hedley and Vasiliki Tsioligka at the Flow Cytometry Facility at the Dunn School of Pathology (University of Oxford), the MBL Imaging Facility, and Ryan Null with HCR probe design assistance. We thank Maria Rossello for discussions about the transcriptional landscape analysis and the DGE analysis.

## Authors’ Contributions

PA-C, HG-C, JS and BDO conceived the study and designed the experiments. PA-C, HG-C and EE generated cell dissociations and performed single cell transcriptomic experiments using *Pristina leidyi*, assisted by VM. HG-C, BM and BDO generated HCR *in situ* hybridisation data. NJK performed bioinformatic experiments on the *Pristina leidyi* transcriptome and initial bioinformatic single cell analyses. AP-P performed bioinformatic analyses on the transcriptional landscape of *Pristina leidyi*. DAS-D performed bioinformatic single cell analyses. JS performed bioinformatic single cell analyses and *Pristina leidyi piwi*+ population transcriptomic analyses. AEB contributed to the interpretation of the single cell analysis data. JS, BDO, PA-C and HG-C wrote the manuscript and generated the figures, with contributions from all other authors. All authors read and approved the final version of the manuscript.

## References

Achim, K., Eling, N., et al., (2018). Whole-Body Single-Cell Sequencing Reveals Transcriptional Domains in the Annelid Larval Body. Mol Biol Evol 35, 1047–1062.

Ackermann, C., Dorresteijn, A., et al., (2005). Clonal domains in postlarval Platynereis dumerilii (Annelida: Polychaeta). J Morphol 266, 258–280.

Alie, A., Hayashi, T., et al., (2015). The ancestral gene repertoire of animal stem cells. Proc Natl Acad Sci U S A 112, E7093–7100.

Altaf, F., Wu, S., et al., (2021). Role of Fibrinolytic Enzymes in Anti-Thrombosis Therapy. Front Mol Biosci 8, 680397.

Bely, A.E., (2006). Distribution of segment regeneration ability in the Annelida. Integr Comp Biol 46, 508–518.

Bely, A.E., (2014). Early events in annelid regeneration: a cellular perspective. Integr Comp Biol 54, 688–699.

Bely, A.E., (2022). Journey beyond the embryo: The beauty of Pristina and naidine annelids for studying regeneration and agametic reproduction. Curr Top Dev Biol 147, 469–495.

Bernstein, N.J., Fong, N.L., et al., (2020). Solo: Doublet Identification in Single-Cell RNA-Seq via Semi-Supervised Deep Learning. Cell Syst 11, 95–101 e105.

Cantalapiedra, C.P., Hernandez-Plaza, A., et al., (2021). eggNOG-mapper v2: Functional Annotation, Orthology Assignments, and Domain Prediction at the Metagenomic Scale. Mol Biol Evol 38, 5825–5829.

Cheng, M.H., Jansen, R.P., (2017). A jack of all trades: the RNA-binding protein vigilin. Wiley Interdiscip Rev RNA 8.

Collado, R., Schmelz, R.M., (2000). Pristina silvicola and Pristina terrena spp. nov., two new soil-dwelling species of Naididae (Oligochaeta, Annelida) from the tropical rain forest near Manaus, Brazil, with comments on the genus Pristinella. Journal of Zoology 251, 509–516.

Cortes, A., Huertas, D., et al., (1999). DDP1, a single-stranded nucleic acid-binding protein of Drosophila, associates with pericentric heterochromatin and is functionally homologous to the yeast Scp160p, which is involved in the control of cell ploidy. EMBO J 18, 3820–3833.

Dattani, A., Sridhar, D., et al., (2019). Planarian flatworms as a new model system for understanding the epigenetic regulation of stem cell pluripotency and differentiation. Semin Cell Dev Biol 87, 79–94.

Emms, D.M., Kelly, S., (2019). OrthoFinder: phylogenetic orthology inference for comparative genomics. Genome Biol 20, 238.

Esarte Palomero, O., Larmore, M., et al., (2023). Polycystin Channel Complexes. Annu Rev Physiol 85, 425–448.

Galperin, M.Y., Makarova, K.S., et al., (2015). Expanded microbial genome coverage and improved protein family annotation in the COG database. Nucleic Acids Res 43, D261–269.

Garcia-Castro, H., Kenny, N.J., et al., (2021). ACME dissociation: a versatile cell fixation-dissociation method for single-cell transcriptomics. Genome Biol 22, 89.

Gaspar-Maia, A., Alajem, A., et al., (2011). Open chromatin in pluripotency and reprogramming. Nat Rev Mol Cell Biol 12, 36–47.

Gehrke, A.R., Srivastava, M., (2016). Neoblasts and the evolution of whole-body regeneration. Curr Opin Genet Dev 40, 131–137.

Gelder, S.R., (1984). Diet and histophysiology of the alimentary canal of Lumbricillus lineatus (Oligochaeta, Enchytraeidae). Hydrobiologia 115, 71–81.

Gerber, T., Murawala, P., et al., (2018). Single-cell analysis uncovers convergence of cell identities during axolotl limb regeneration. Science 362, eaaq0681.

Giere, O., Rhode, B., (1987). Anatomy and ultrastructure of the marine oligochaete Tubificoides benedii (Tubificidae), with emphasis on its epidermis-cuticle-complex. Hydrobiologia 155, 159–159.

Gilbert, D.G., (2019). Longest protein, longest transcript or most expression, for accurate gene reconstruction of transcriptomes? bioRxiv, 829184.

Goto, A., Kitamura, K., et al., (1999). Cell fate analysis of teloblasts in the Tubifex embryo by intracellular injection of HRP. Dev Growth Differ 41, 703–713.

Gough, J., Karplus, K., et al., (2001). Assignment of homology to genome sequences using a library of hidden Markov models that represent all proteins of known structure. J Mol Biol 313, 903–919.

Hulett, R.E., Kimura, J.O., et al., (2022). Acoel single-cell atlas reveals expression dynamics and heterogeneity of a pluripotent stem cell population. bioRxiv, 2022.2002.2010.479464.

Jones, P., Binns, D., et al., (2014). InterProScan 5: genome-scale protein function classification. Bioinformatics 30, 1236–1240.

Juliano, C.E., Swartz, S.Z., et al., (2010). A conserved germline multipotency program. Development 137, 4113–4126.

Kuehn, E., Clausen, D.S., et al., (2022). Segment number threshold determines juvenile onset of germline cluster expansion in Platynereis dumerilii. J Exp Zool B Mol Dev Evol 338, 225–240.

Labbe, R.M., Irimia, M., et al., (2012). A comparative transcriptomic analysis reveals conserved features of stem cell pluripotency in planarians and mammals. Stem Cells 30, 1734–1745.

Lai, A.G., Aboobaker, A.A., (2018). EvoRegen in animals: Time to uncover deep conservation or convergence of adult stem cell evolution and regenerative processes. Dev Biol 433, 118–131.

Langfelder, P., Horvath, S., (2008). WGCNA: an R package for weighted correlation network analysis. BMC Bioinformatics 9, 559.

Levy, S., Elek, A., et al., (2021). A stony coral cell atlas illuminates the molecular and cellular basis of coral symbiosis, calcification, and immunity. Cell 184, 2973–2987 e2918.

Lust, K., Maynard, A., et al., (2022). Single-cell analyses of axolotl telencephalon organization, neurogenesis, and regeneration. Science 377, eabp9262.

Marakulina, D., Vorontsov, I.E., et al., (2023). EpiFactors 2022: expansion and enhancement of a curated database of human epigenetic factors and complexes. Nucleic Acids Res 51, D564–D570.

Martinez Arias, A., Brickman, J.M., (2011). Gene expression heterogeneities in embryonic stem cell populations: origin and function. Curr Opin Cell Biol 23, 650–656.

Mashima, R., Okuyama, T., (2015). The role of lipoxygenases in pathophysiology; new insights and future perspectives. Redox Biol 6, 297–310.

Medvedeva, Y.A., Lennartsson, A., et al., (2015). EpiFactors: a comprehensive database of human epigenetic factors and complexes. Database (Oxford) 2015, bav067.

Messmer, T., von Meyenn, F., et al., (2019). Transcriptional Heterogeneity in Naive and Primed Human Pluripotent Stem Cells at Single-Cell Resolution. Cell Rep 26, 815–824 e814.

Meyer, N.P., Boyle, M.J., et al., (2010). A comprehensive fate map by intracellular injection of identified blastomeres in the marine polychaete Capitella teleta. Evodevo 1, 8.

Mohammed, H., Hernando-Herraez, I., et al., (2017). Single-Cell Landscape of Transcriptional Heterogeneity and Cell Fate Decisions during Mouse Early Gastrulation. Cell Rep 20, 1215–1228.

Moreno-Hagelsieb, G., Latimer, K., (2008). Choosing BLAST options for better detection of orthologs as reciprocal best hits. Bioinformatics 24, 319–324.

Musser, J.M., Schippers, K.J., et al., (2021). Profiling cellular diversity in sponges informs animal cell type and nervous system evolution. Science 374, 717–723.

Nyberg, K.G., Conte, M.A., et al., (2012). Transcriptome characterization via 454 pyrosequencing of the annelid Pristina leidyi, an emerging model for studying the evolution of regeneration. BMC Genomics 13, 287.

Önal, P., Grun, D., et al., (2012). Gene expression of pluripotency determinants is conserved between mammalian and planarian stem cells. EMBO J 31, 2755–2769.

Özpolat, B.D., Bely, A.E., (2015). Gonad establishment during asexual reproduction in the annelid Pristina leidyi. Dev Biol 405, 123–136.

Özpolat, B.D., Bely, A.E., (2016). Developmental and molecular biology of annelid regeneration: a comparative review of recent studies. Curr Opin Genet Dev 40, 144–153.

Özpolat, B.D., Handberg-Thorsager, M., et al., (2017). Cell lineage and cell cycling analyses of the 4d micromere using live imaging in the marine annelid Platynereis dumerilii. Elife 6.

Özpolat, B.D., Sloane, E.S., et al., (2016). Plasticity and regeneration of gonads in the annelid Pristina leidyi. Evodevo 7, 22.

Pandurangan, A.P., Stahlhacke, J., et al., (2019). The SUPERFAMILY 2.0 database: a significant proteome update and a new webserver. Nucleic Acids Res 47, D490–D494.

Plass, M., Solana, J., et al., (2018). Cell type atlas and lineage tree of a whole complex animal by single-cell transcriptomics. Science 360, eaaq1723.

Ritchie, M.E., Phipson, B., et al., (2015). limma powers differential expression analyses for RNA-sequencing and microarray studies. Nucleic Acids Res 43, e47.

Robinson, M.D., McCarthy, D.J., et al., (2010). edgeR: a Bioconductor package for differential expression analysis of digital gene expression data. Bioinformatics 26, 139–140.

Rosenberg, A.B., Roco, C.M., et al., (2018). Single-cell profiling of the developing mouse brain and spinal cord with split-pool barcoding. Science 360, 176–182.

Schenk, S., Hoeger, U., (2020). Annelid Coelomic Fluid Proteins. Subcell Biochem 94, 1–34.

Schlesinger, S., Meshorer, E., (2019). Open Chromatin, Epigenetic Plasticity, and Nuclear Organization in Pluripotency. Dev Cell 48, 135–150.

Siebert, S., Farrell, J.A., et al., (2019). Stem cell differentiation trajectories in Hydra resolved at single-cell resolution. Science 365.

Smith, C.M., Weisblat, D.A., (1994). Micromere fate maps in leech embryos: lineage-specific differences in rates of cell proliferation. Development 120, 3427–3438.

Solana, J., (2013). Closing the circle of germline and stem cells: the Primordial Stem Cell hypothesis. Evodevo 4, 2.

Solana, J., Kao, D., et al., (2012). Defining the molecular profile of planarian pluripotent stem cells using a combinatorial RNAseq, RNA interference and irradiation approach. Genome Biol 13, R19.

Song, S., Starunov, V., et al., (2020). Globins in the marine annelid Platynereis dumerilii shed new light on hemoglobin evolution in bilaterians. BMC Evol Biol 20, 165.

Stephenson, J., (1922). XII.—On the Septal and Pharyngeal Glands of the Microdrili (Oligochæta). Earth and Environmental Science Transactions of The Royal Society of Edinburgh 53, 241–264.

Sur, A., Meyer, N.P., (2021). Resolving Transcriptional States and Predicting Lineages in the Annelid Capitella teleta Using Single-Cell RNAseq. Frontiers in Ecology and Evolution 8.

Tanay, A., Sebe-Pedros, A., (2021). Evolutionary cell type mapping with single-cell genomics. Trends Genet 37, 919–932.

Thomas, P.D., Ebert, D., et al., (2022). PANTHER: Making genome-scale phylogenetics accessible to all. Protein Sci 31, 8–22.

Tritschler, S., Buttner, M., et al., (2019). Concepts and limitations for learning developmental trajectories from single cell genomics. Development 146.

UniProt, C., (2023). UniProt: the Universal Protein Knowledgebase in 2023. Nucleic Acids Res 51, D523–D531.

van Wolfswinkel, J.C., Wagner, D.E., et al., (2014). Single-cell analysis reveals functionally distinct classes within the planarian stem cell compartment. Cell Stem Cell 15, 326–339.

Vergara, H.M., Pape, C., et al., (2021). Whole-body integration of gene expression and single-cell morphology. Cell 184, 4819–4837 e4822.

Weisblat, D.A., Shankland, M., (1985). Cell lineage and segmentation in the leech. Philos Trans R Soc Lond B Biol Sci 312, 39–56.

Wolf, F.A., Hamey, F.K., et al., (2019). PAGA: graph abstraction reconciles clustering with trajectory inference through a topology preserving map of single cells. Genome Biol 20, 59.

Wolock, S.L., Lopez, R., et al., (2019). Scrublet: Computational Identification of Cell Doublets in Single-Cell Transcriptomic Data. Cell Syst 8, 281–291 e289.

Xu, Y., Bolton, B., et al., (1999). Gliarin and macrolin, two novel intermediate filament proteins specifically expressed in sets and subsets of glial cells in leech central nervous system. J Neurobiol 40, 244–253.

Zattara, E.E., Bely, A.E., (2011). Evolution of a novel developmental trajectory: fission is distinct from regeneration in the annelid Pristina leidyi. Evol Dev 13, 80–95.

Zattara, E.E., Bely, A.E., (2013). Investment choices in post-embryonic development: quantifying interactions among growth, regeneration, and asexual reproduction in the annelid Pristina leidyi. J Exp Zool B Mol Dev Evol 320, 471–488.

Zattara, E.E., Bely, A.E., (2015). Fine taxonomic sampling of nervous systems within Naididae (Annelida: Clitellata) reveals evolutionary lability and revised homologies of annelid neural components. Front Zool 12, 8.

