## Supplement for "Annelid adult cell type diversity and their pluripotent cellular origins"

### Supplementary Note
### Supplementary Figure Legends
### Supplementary File Legends
### Supplementary Methods
### Supplementary References

#### Supplementary Note 1: naming and grouping of the clusters

To characterise the cell types represented by each of our cell clusters we analysed and discussed the markers of each cluster, their Diamond BLAST annotations, and put these into the context of the available annelid and single cell literature. Broad groups of cell types were obtained by leveraging the information of the PAGA analysis and the co-occurrence matrix.

***Piwi*+ cells**: We found that the expression of *piwi-1* concentrated in clusters 1 and 2, but also in cluster 8. A number of unannotated clusters were mixed with clusters 1 and 2 in the UMAP space and also expressed *piwi-1*.

**Epidermis**: We identified epidermal cells based on the analysis of the expression of the markers PrileiEVm008309t1 and PrileiEVm008287t1, encoding intermediate filament proteins.

**Gut**: We found this group of cell clusters connected to cluster 16, suggesting that cluster 16 contains gut progenitors. We identified and named the different regions of the *Pristina leidyi* gut based on the analysis of the expression of the markers PrileiEVm010132t1 (unannotated transcript), PrileiEVm010941t1 (unannotated transcript), PrileiEVm000199t1 (encoding a rootletin protein), PrileiEVm017310t1 (unannotated transcript), PrileiEVm019805t1 (unannotated transcript), PrileiEVm005677t1 (encoding a long-chain-fatty-acid-CoA ligase protein), PrileiEVm021761t1 (unannotated transcript), PrileiEVm001383t1 (encoding a von Willebrand factor D and EGF domain-containing protein), PrileiEVm026965t1 (unannotated transcript) and PrileiEVm020317t1 (unannotated transcript). We found that Cluster 38 was only connected to the group of gut cell clusters below the threshold and we named it after the marker PrileiEVm022227t1 (encoding a caveolin protein), but kept it in the broad gut group.

**Muscle**: We found that clusters 5, 6, 27 and 30 highly expressed components of the muscle sarcomere, including myosin heavy chain (PrileiEVm000300t1), myosin light chain (PrileiEVm019310t1, PrileiEVm025675t1, PrileiEVm020932t1), tropomyosin (PrileiEVm016172t1, PrileiEVm015909t1), troponin (PrileiEVm014978t1, PrileiEVm014978t1, PrileiEVm018738t1) and titin (PrileiEVm000447t1). Cluster 27 contains cells that are mixed with Clusters 8, 1 and 2, is connected by PAGA analysis to cluster 8, and likely represents a progenitor state.

**Neurons**: We identified the neuronal populations based on the expression of synaptotagmin (PrileiEVm012030t1) and other neuronal specific transcripts such as PrileiEVm000558t1, which *in-situ* expression marks the Ventral Nerve Cord (VNC). Those markers were highly expressed in cluster 3, 40, 41, 42, 45, 49 and 59. Cluster 49 also expressed a homolog of arrestin protein-domain (PrileiEVm014226t1) suggesting a possible identity for this cluster as photoreceptor neurons.

***Globin*+ cells**: We found that clusters 4 and 33 highly expressed extracellular globins, including PrileiEVm020672t1, PrileiEVm018061t1, PrileiEVm020599t1, PrileiEVm020147t1.

***Polycystin* cells**: We named this cluster group based on the expression of *polycystin*-2 (PrileiEVm005033t1) and *polycystin*-1+ (PrileiEVm004079t1) homologue genes.

**Eleocytes**: We identified this cluster based on the expression of a vitellogenin homologue (PrileiEVm000002t1) which was also expressed in *globin*+ cells. The 3 clusters interconnected in the PAGA graph were named after markers *mmp24* (PrileiEVm009033t1), *fucolectin* (PrileiEVm007557t1) and *pgrn* (PrileiEVm018088t1).

**Chaetal sacs**: We identified the chaetal sacs based on the expression of PrileiEVm000939t1 (encoding a kunitz-type protease inhibitor protein), PrileiEVm007502t1 (encoding an intermediate filament protein) and PrileiEVm000573t1 (encoding a chitin synthase protein).

***Lipoxygenase*+ cells**: We named this cluster based on the expression of several *lipoxygenase* transcripts, including PrileiEVm000278t1, PrileiEVm008087t1 and PrileiEVm008285t1.

***Vigilin*+ cells**: We named this cluster after the expression of the RNA binding protein *vigilin* (PrileiEVm001339t1).

***Lumbrokinase*+ cells**: We found that cluster 26 highly expressed several homologues of the lumbrokinase enzymes, including PrileiEVm016387t1, PrileiEVm016330t1, PrileiEVm016446t1 and PrileiEVm016633t1.

**Carbohydrate metabolic cells**: We named this cluster based on the expression of several transcripts annotated as enzymes involved in carbohydrate metabolism, including a fructose-bisphosphate aldolase (PrileiEVm013994t1), a phosphoenolpyruvate carboxykinase (PrileiEVm026130t1), a malate dehydrogenase (PrileiEVm014911t1), as well as mitochondrial enzymes such as a glutamate dehydrogenase (PrileiEVm008904t1), a pyruvate carboxylase (PrileiEVm001525t1), and a mitochondrial malate dehydrogenase (PrileiEVm026983t1).

**Secretory**: We proposed cluster 34 and 44 as secretory clusters based on the expression of a homolog of the conotoxin protein (PrileiEVm010163t1). The *in-situ* for this transcript marks a cluster of large cells segmentally repeating on the ventral side, after each chaetae bundle but not overlapping with the chaetae themselves. Also, some single cells in the posterior growth zone seem to be marked. These clusters also appear to express the synaptotagmin transcript (PrileiEVm012030t1) suggesting a possible neuro-secretory function.

***Arginase*+ cells**: We named this cluster group based on the expression of an arginase homologue, PrileiEVm015527t1

***Ldlrr*+ cells**: We named this cluster group based on the expression of a low-density lipoprotein receptor homologue, PrileiEVm010669t1.

**Metanephridia**: We identified metanephridia cells based on the analysis of expression data for the marker PrileiEVm002621t1.

### Supplementary Figure Legends

#### Figure Supplementary 1: Doublet identification with Solo and Scrublet

A – UMAP visualisation of the 80,387 cells prior to doublet classification. Cells classified as doublets by Solo and Scrublet are highlighted in pink and orange respectively, and cells classified as doublets by both are highlighted in red.

B – UMAP visualisation of the 80,387 cell dataset after clustering in 4 different resolutions.

C – Percentage of cell classified as doublets by Solo, Scrublet and both methods in each cell cluster across the 4 different resolutions

#### Figure Supplementary 2: *Pristina leidyi* cell type atlas metrics

A – Scatter plot of number of counts per cell vs number of genes per cell.

B – UMAP visualisation of 75,218-cell *Pristina leidyi* single-cell transcriptomic cell atlas with cells coloured according to the experiment of origin, named Lib 12, Lib 21 and Lib 30.

C – Violin plots showing the distribution of genes detected per cell in each cluster.

D – Violin plots showing the distribution of UMI counts detected per cell in each cluster.

E – Dot Plot showing the expression of the top 2 markers detected by the Wilcoxon method in each cluster. Clusters are sorted according to hierarchical clustering. The values plotted in each dot are the fraction of cells in each cluster that express each marker (dot size) and the score of the marker (dot colour)

F – Score distribution of the top 20 markers of each cluster detected by the Wilcoxon method (left) and the logistic regression method (right)

#### Figure Supplementary 3: *Pristina leidyi* cell type atlas cell clusters and broad cell clusters

A – UMAP visualisation of the 75,218-cell *Pristina leidyi* single-cell transcriptomic cell atlas with clusters coloured according to their cell type classification, subdivided by broad clusters.

B – UMAP visualisation of the 75,218-cell *Pristina leidyi* single-cell transcriptomic cell atlas with unannotated clusters highlighted.

#### Figure Supplementary 4: Lineage reconstruction with unannotated clusters

Lineage reconstruction abstracted graph showing the most probable path connecting the clusters. Each node corresponds to the cell clusters identified with the leiden algorithm. The size of nodes is proportional to the amount of cells in the cluster, and the thickness of the edges is proportional to the connectivity probabilities. This analysis includes the unannotated clusters (1,048 cells, 1.39% of the total dataset).

#### Figure Supplementary 5: Co-ocurrence analysis

Cell cluster co-ocurrence matrix showing similarities between related cell types based on gene expression correlation.

#### Figure Supplementary 6: Negative controls

For all HCR experiments, controls that included only hairpin amplifiers but no probes were carried out, imaged using the same imaging conditions, and images were processed with the same settings as the HCR samples.

A – Panel shows maximum projections of Z-stacks in the head region of *Pristina* for a control and an HCR sample, when the same imaging and image processing conditions are applied. The head region typically did not have a lot of background or autofluorescence, except for chaetae, which typically show autofluorescence.

B – Single focal plane images of the gut region in HCR and control samples. Stomach region typically showed some background signal (hollow arrowheads) while the real signal was generally easy to distinguish (solid arrowheads) especially when signal-to-noise ratio was high. In the third row, background is shown when levels are adjusted to maximum in Fiji, but for the same control sample, if the image processing is done at the same levels as the HCR samples (second row), there is very little background.

C – Panel shows maximum projections of Z-stacks in the tail region for a control and an HCR sample, when the same imaging and image processing conditions are applied. When amplifiers with Alexa-546 fluorophore were used, we typically observed some background in the epidermal layer (hollow arrowheads), but the actual HCR signal was still possible to distinguish (solid arrowheads) (also see panel D).

D – Additional examples of autofluorescence and background signals. When the Alexa-647 fluorophore was used, gut content showed a high background (pink), therefore we avoided imaging the areas where there was high gut content. Nevertheless, the real signal was still possible to distinguish in these samples (solid arrowheads). On the right, we show another example of epidermal background (hollow arrowheads) versus actual signal (solid arrowheads) in the head region in different samples.

#### Figure Supplementary 7: Gut variations

Detailed analyses showing all the samples analysed for the markers of gut clusters. Star denotes the tail end of the worm (anus). When there is a number next to the star, the number indicates the total number of segments for that particular worm (which was not possible to fit into the schematic because of length). 3 or 4 gut markers were tested in each set of worms, allowing analyses of multiple markers in a single sample. Each sample is indicated on the left (e.g. "Worm 01). Bars show the extent of expression observed for each marker.

#### Figure Supplementary 8: Pseudobulk Transcription Factor Analysis

A – Barplot showing the number of genes quantified with more than 5 counts per million (cpm) per cluster in a pseudobulk analysis, showing that most clusters have > 4000 genes quantified, with a mean of 11,117 genes per cluster.

B – Barplot showing the number of TFs per TF type.

C – Boxplot showing the coefficient of variation distribution for each TF type.

D – Barplot showing the top TF classes based on the number of instances where gene expression of a given TF class was found to be significant in explaining differences between cell clusters (ANOVA, Tukey test comparison of means).

E – (Left) Barplot showing predominance of different TF classes in the transcriptomic profile of different cell clusters, measured as number of CPMs per gene and per class. Each color represents a transcription factor class. (Right) clustering of cell clusters based on similarities in the transcriptomic profile of TF classes.

#### Figure Supplementary 9:

UMAP visualisation of *piwi*+ subclusters in the 75,218 and the 16,247 datasets containing all cells and only *piwi*+ cell clusters, respectively, and score UMAP plots of markers of *piwi*+ subclusters. Markers were calculated using the Wilcoxon and the Logistic Regression methods.

### Supplementary File Legends

#### Supplementary File 1: EggNOG and Diamond Blast Annotation

#### Supplementary File 2: Preprocessing parameter space exploration

UMAP visualisation of the 80,387 cells preprocessed with different parameters for minimum gene counts (30-100), maximum number of genes (300-1000), maximum number of counts (500-1200), number of top highly variable genes (4,000-22,000), number of neighbors (15-85), and number of principal components (15-145). For each parameter, UMAP feature plots of diagnostic markers PrileiEVm023936t1 (piwi+ cells), PrileiEVm008309t1 (epidermis), PrileiEVm011741t1 (muscle), PrileiEVm021316t1 (gut), PrileiEVm022250t1 (gut), PrileiEVm000325t1 (neurons), PrileiEVm013699t1 (eleocytes), and PrileiEVm020595t1 (unnanotated).

#### Supplementary File 3: Cluster Wilcoxon markers and Diamond Blast annotations

#### Supplementary File 4: Cluster Logistic Regression markers and Diamond Blast annotations

#### Supplementary File 5: *Pristina leidyi* cell cluster markers

UMAP visualisation of 75,218-cell *Pristina leidyi* single-cell transcriptomic cell atlas. In the top left panel of each page each cluster is coloured. In the remaining plots, UMAP feature plots of markers of the top 8 common markers among the Wilcoxon and the Logistic Regression methods. If there are less than 8 common markers the plots only display the UMAP visualisation without any feature gene expression.

**Supplementary File 6: HCR *in situ* probes**

**Supplementary File 7: WGCNA modules**

**Supplementary File 8: WGCNA modules GO terms**

**Supplementary File 9: Transcription Factor annotations**

**Supplementary File 10:*piwi*+ cells COGs**

**Supplementary File 11: Top 200 TFs expressed in *piwi*+ cells**

**Supplementary File 12: Top 200 TFs expressed in *piwi*+ cells UMAP visualisation**

**Supplementary File 13: Transcriptomic profile of *piwi*+ cells and Diamond Blast annotations**

**Supplementary File 14: Annotation of *Pristina* epigenetic factor homologues**

### Supplementary Methods

#### *Pristina leidyi* culture and maintenance

*Pristina leidyi* culture was originally obtained from Carolina Biological Supply (Bely and Wray, 2001). Specimens were cultured both in plastic boxes and fish tanks with 1 L and 50 L of 1% filtered artificial seawater, respectively. Water was changed every week and animals were fed with 0.03g/L of dried spirulina powder every 2 weeks. Under these conditions, worms reproduce continuously by paratomic fission (Bely and Wray, 2001).

#### Iso-seq

Approximately 100 *Pristina leidyi* of mixed conditions, including fissioning animals, were manually picked out of culture using a glass Pasteur pipette. These were placed into a single 1.5 mL Eppendorf tube, and spun on a low speed benchtop centrifuge to pellet. The supernatant was removed. Total RNA was extracted from the pelleted worms using the Trizol method and the standard manufacturer protocol. The quality of this was assessed using a Nanodrop, giving a concentration of 1083.7 ng/uL, an A260/A280 ratio of 2.01 and an A260/A230 ratio of 2.03. Quality was further assessed using a Bioanalyzer (Agilent), although a RIN value was not calculated due to the difference in profile commonly observed in annelid RNA samples. Total RNA was provided to the Earlham Institute Genomics Pipelines Group, Norwich, UK, and (after QC to confirm quality) was used as the basis of PacBio Iso-Seq Express Template Preparation (v2) library construction. This sample, along with 3 others, was loaded onto a PacBio Sequel II SMRT cell, and sequenced (8M, v2, 30hr Movie).

Iso-Seq3 analysis was performed by the provider. A total of 3,932,103 CCS reads were captured across the samples on the cell, with 1,546,939 assigned to *Pristina leidyi.* These were classified and clustered, resulting in 54,350 high-quality isoforms.

#### Sequence concatenation and redundancy removal

The sequences gained from Iso-Seq sequencing analysis were combined with sequences derived from previous analysis of the *Pristina leidyi* transcriptome (Nyberg et al., 2012). First the isotigs from the Nyberg et al dataset were concatenated with the isoform sequences derived from Iso-Seq analysis. Redundancy was removed from these reads using the EvidentialGene (Gilbert, 2019) tr2aacds4.pl approach (March 2020 v4 version) with settings -cdnaseq -NCPU 8 -MAXMEM 16000 -logfile, keeping only a single sequence representative per locus with the best evidence score. Transdecoder v5.5 was then used to predict the protein coding regions of transcripts (LongOrfs -m 25, Predict --single_best_only).

#### Diamond Blast annotation

We implemented diamond v2.0.8.146 (Buchfink et al., 2021) to provide an initial putative identity to orthologs present in our reference transcriptome. This software performed a blastx search against the whole downloaded database with default settings and organized the results into a table with the settings --salltitles -b8 -c1 -p8 --outfmt 6 qseqid sseqid pident evalue stitle.

#### eggNOG annotation

The assembled transcriptome of *Pristina leidyi* was transformed to protein sequence using TransDecoder ( https://github.com/TransDecoder/TransDecoder/wiki); first, we ran `TransDecoder.LongOrfs` with standard parameters; second, we ran hmmscan vs Pfam database and BLAST vs Swissprot database, with parameters:`-max_target_seqs 1 -evalue 1e-5` and default parameters respectively, to gather supporting evidence for coding transcripts; third, we ran `TransDecoder.Predict` with parameters `--retain_pfam_hits pfam.domtblout --retain_blastp_hits blastp.outfmt6 --single_best_only`. The resulting translated transcriptome (hereafter referred to as proteome) was queried using EggNOG mapper (Cantalapiedra et al., 2021) with the parameters: `-m diamond --sensmode sensitive --target_orthologs all --go_evidence non-electronic` against the EggNOG metazoa database. From the EggNOG output, GO term, functional category COG, and gene name association files, were generated using custom bash code. Full code is available at the project repository.

#### ACME dissociation

ACME was performed as previously described (Garcia-Castro et al., 2021) with some modifications. For each sample, we added ~120 *Pristina leidyi* worms at mixed stages (including fissioning animals) to a 15 mL Falcon tube (~100 uL of biomass volume). Depending on the batch, we used animals at different starvation conditions: 12 days (library 12), 4 days (library 21) or 7 days (library 30). We removed most culture water and added 300 uL of NAC solution per tube. NAC solution was freshly prepared by diluting N-acetyl cysteine powder in 1x PBS buffer to a 7.5% w/v. The 1x PBS buffer was made from a nuclease-free 10x PBS stock solution. We flicked samples in NAC for 30’’, and added 10 mL of ACME solution per tube immediately after. The ACME solution was prepared fresh using 6.5 mL of nuclease-free commercial H2O, 1.5 mL of methanol, 1 mL of acetic acid and 1 mL of glycerol per sample. Samples were incubated in ACME for 35 min, at room temperature, in a rocking table (40-45 rpm). To help dissociation, tubes were manually shaken every 10 min. After incubation, samples were pipetted up and down to complete dissociation. From this point, samples were kept on ice to prevent RNA degradation. With cells still on ACME, we filtered through 50 µm strainers (CellTrics) into new 15 mL Falcon tubes. Samples were centrifuged at 1000 g for 6 min (4°C) to remove ACME, and pellets were resuspended in 8 mL of 1x PBS 1% BSA fresh buffer. We centrifuged again at 1000 g for 6 min (4°C) and discarded the supernatant. Pellets were resuspended in 900 uL of 1x PBS 1% BSA fresh buffer and transferred to 1.5 mL Eppendorf tubes. To cryopreserve cells, we added 100 uL of DMSO per sample and stored at -80°C.

#### SPLiT-seq

SPLiT-seq was performed as previously described (Garcia-Castro et al., 2021) with the following modifications:

Cell count

Cryopreserved ACME-dissociated cells were thawed and centrifuged twice at 1000 g for 6 min (4°C) to remove the DMSO. Pellets were resuspended in 250 uL of 1x PBS 1% BSA fresh buffer. For each sample, we prepared a separate 1:3 dilution with 50 uL of cells and 100 uL of buffer. Dilutions were stained for 15 min, at RT, with 0.2 uL of DRAQ5 (5 mM stock solution, Bioscience) and 0.6 uL of Concanavalin-A conjugated with AlexaFluor 488 (1 mg/mL stock solution, Invitrogen). The remaining undiluted samples were kept at 4°C. Cell count was performed on the stained dilutions by flow cytometry. From this, we calculated the concentration on the main samples and diluted them to a final working concentration of 625-1,250 events/uL.

Round 1 of Barcoding: Reverse Transcription

The Round 1 plate was loaded with 8 uL/well of Round 1 barcodes, 8 uL/well of cells at a concentration of 625-1,250 events/uL (5,000-10,000 events per well) and 8 uL/well of the following RT mix: 4μL of 5x Maxima RT Buffer (Thermo Scientific), 0.375 μL of Superase-In RNAse inhibitor (20 U/μL, Invitrogen), 1 μL of 10 mM/each dNTPs (NEB), 0.625 μL of nuclease-free H2O and 2 μL of Maxima H Minus RT (200 U/μL, Thermo Scientific). In library 30, we also added 10% w/v of PEG 8000 to the RT mix.The reverse transcription reaction ran in a thermocycler for 35 min at 50ºC. After incubation, reactions were pooled in a 15 mL Falcon tube. We added 10% Triton X-100 to the cells, to a final concentration of 0.1%, and centrifuged at 1200 g for 6 min. Cells were resuspended in 2 mL of NEB buffer 3.1 (NEB) with 20 uL of Superase-In RNase Inhibitor (20 U/uL, Invitrogen).

Round 2 of Barcoding: Ligation 1

The ligation mix was prepared with 500 μL of 10x T4 Ligase Buffer (NEB), 100 μL of T4 DNA ligase (400 U/μL, NEB), 100 μL of 1x PBS 1% BSA buffer, and 1340 μL of nuclease-free water. For library 30, we additionally added 10% w/v of PEG 8000 to the ligation mix.

Round 3 of Barcoding: Ligation 2

Pooled cells from Round 2 were mixed with 150 μL of T4 DNA ligase (400 U/μL, NEB). The Round 3 plate was loaded with 55 μL/well of this mix.

Washing

After last blocking, we pooled cells in a 15 mL Falcon tube and added 10% Triton-X 100 to a final concentration of 0.1%. Cells were centrifuged at 1200 g for 6 min (4°C). The supernatant was discarded and the pellet was resuspended in 4.04 mL of washing buffer (4 mL of 1x PBS and 40 µL of 10% Triton X-100). Cells were centrifuged again, resuspended in 800 uL of 1x PBS 1% BSA buffer, and split in two 1.5 mL Epp tubes (400 uL/each). These samples were stored at -80°C in 10% DMSO.

FACS

FACS was performed in the middle of the SPLiT-seq protocol. We thawed previously barcoded samples, added 2 µL of 10% Triton X-100 per tube, and centrifuged at 1200 g for 6 min (4ºC) to eliminate the DMSO. Supernatants were carefully discarded, and pellets were resuspended in 500 µL of 1x PBS 1% BSA buffer. We added another 2 uL of 10% Triton X-100 per tube and repeated centrifugation in the same conditions. Final pellets were resuspended in 400 µL of 1x PBS 1% BSA buffer and stained with 0.5 µL of DRAQ5 (5 mM stock solution, Bioscience) and 1 µL of Concanavalin-A conjugated with AlexaFluor 488 (1 mg/mL stock solution, Invitrogen). Stained cells were incubated for 45 min, on ice, in a dark box. Cells were sorted using a BD FACS Aria III (BD Biosciences) set in 4-ways Purify Mode and 45 Psi of pressure, with an 85-um nozzle. DRAQ5 and Concanavalin-A positive singlets were sorted in sub-libraries of 9000-25,000 cells, collected directly into 50 uL of 2x Lysis Buffer. FACS time was about 1.5 hours per batch.

Cell lysis

The sorted sub-libraries were adjusted to a volume of 100 uL, when necessary, using 1x PBS 1% BSA buffer. We added 10 µL of Proteinase K (20 mg/mL) to each sub-library and incubated for 2 h at 55ºC. After incubation, lysates were frozen at -80ºC.

Template Switch

The Template Switch mix was prepared using 44 µL of 5x Maxima RT Buffer (Thermo Scientific), 44 µL of 20% Ficoll PM 400 (Sigma Aldrich), 22 µL of 10 mM/each dNTPs (NEB), 5.5 µL of Superase-In RNAse inhibitor (20 U/µL, Invitrogen), 5.5 µL of TSO primer (100 µM), 11 µL of Maxima H Minus RT (200 U/µL, Thermo Scientific), 0.022g (10% w/v) of PEG 8000 (only for libraries 21 and 30), and up to 220 µL of nuclease-free water per sample.

PCR Amplification

Samples were amplified for 5 cycles of PCR and 10-11 cycles of qPCR.

Size selection

We purified qPCR reactions by two consecutive rounds of SPRI size selection at ratios of 0.8x and 0.7x. After the first 0.8x size selection, the eluted volume (20 uL) was adjusted to 100 uL using nuclease-free water. Final fragment distributions and concentrations were assessed by running a

High Sensitivity DNA bioanalyzer (Agilent 2100) and a Qubit dsDNA High Sensitivity Assay (Thermo Fisher), respectively, according to the manufacturer's protocols.

Tagmentation

Tagmentation was performed using the Nextera XT DNA Library Preparation Kit (Illumina). We prepared the tagmentation reactions by mixing 5 µL of cDNA (1 ng in total), 10 µL of Tagment DNA Buffer (TD) and 5 µL of Amplicon Tagment Mix (ATM). Reactions were incubated in a preheated thermocycler for 5 min at 55ºC. Samples were placed on ice immediately after incubation. To stop tagmentation, we added 5 µL of Neutralize Tagment Buffer (NT), mixed well, and incubated at room temperature for 5 min.

Round 4 of Barcoding: PCR

We prepared a separate reaction mix for each sub-library, containing 22 µL of tagmented cDNA, 15 µL of Nextera PCR Master Mix (Nextera XT DNA Library Preparation Kit), 1 µL of P5_oligo (10 µM) and 1 µL of a Round 4 barcode (10 µM). We used different barcodes for each sub-library. The PCR reaction ran as follows: 72ºC (3 min); 95ºC (30 s); 12 cycles of 95ºC (10 s), 55ºC (30 s) and 72ºC (30 s); and 72ºC (5 min). PCR samples were purified by two subsequent rounds of SPRI size selection (0.7x and 0.6x). Fragment distribution was assessed running a High Sensitivity DNA bioanalyzer (Agilent 2100) and final concentrations were quantified using a Qubit dsDNA High Sensitivity Assay (Thermo Fisher).

#### SPLiT-seq read processing

SPLiTseq reads were provided by Novogene (China). A total of 124,349,078 (12_1), 135,900,060 (12_2), 410,765,606 (21_1), 833,784,688 (21_2), 807,486,658 (21_3), 643,285,668 (30_2), 627,640,824 (30_3), 711,569,074 (30_4), 725,038,254 (30_5) reads were sequenced. These were assayed for QC purposes using FastQC (https://www.bioinformatics.babraham.ac.uk/projects/fastqc/, v0.11.9, 2019) and residual adaptor sequence, low-quality, and short reads were observed. CutAdapt v2.8 (Martin, 2011) was used to trim read 1 (transcripts) and read 2 (UMI and barcodes) sequences. The following settings: cutadapt -j 4 -m 60 -q 10 -b AGATCGGAAGAG were run for read 1. To trim read 2, settings: cutadapt -j 4 -m 94 --trim-n -q 10 -b CTGTCTCTTATA were used. To confirm barcodes were correctly in position, and not affected by indels, read 2 sequences were checked for "phase" using grep, with known flanking sequence as a search. Reads were retained when UMI and UBC barcodes were in the correct location. Finally, pairfq makepairs v 0.17 (https://github.com/sestaton/Pairfq) was used to retain correctly paired, complete reads. These were fed into SPLiTseq toolbox (https://github.com/RebekkaWegmann/splitseq_toolbox v 1.0) for further analysis.

The Iso-seq transcriptome of *Pristina leidyi* assembled as described above was created to have a reference database for read mapping. We then used Dropseq_tools-2.3.0 (https://github.com/broadinstitute/Drop-seq/releases/tag/v2.3.0) to process the generated GTF file and create a sequence dictionary, a refFlat, a reduced GTF and the corresponding interval files. We generated a reference index using STAR-2.7.3a (Dobin et al., 2013) with the parameters --sjdbOverhang 99 --genomeSAindexNbases 13 --genomeChrBinNbits 14. Each of the sub-libraries was processed separately and properly combined later in the analysis. The SPLiTseq toolbox (https://github.com/RebekkaWegmann/splitseq_toolbox) which envelops algorithms from Drop-seq_tools-2.3.0, was used to retrieve, correct and label the barcodes with a hamming distance ≤1. Mapping to the reference transcriptome used STAR-2.7.3a (https://github.com/alexdobin/STAR/releases/tag/2.7.3a) with --quantMode GeneCounts and all other default settings with the exception of --outFilterMultimapNmax 10 to retain and analyse reads which mapped up to ten different loci in the reference. We implemented Picard v2.21.1-SNAPSHOT (https://github.com/broadinstitute/picard) to re-order, merge, align and tag reads for each sub-library with the SortSam and MergeBamAlignment features. We implemented sequentially the features Drop-seq_tools-2.3.0 TagReadWithInterval and TagReadWithGeneFunction to create expression matrices of each library with the feature of Drop-seq_tools-2.3.0 DigitalExpression with the settings: READ_MQ=0, EDIT_DISTANCE=1, MIN_NUM_GENES_PER_CELL=50, and LOCUS_FUNCTION_LIST=INTRONIC. These matrices together with the gene models and raw reads are uploaded to GEO under the accession code GSE230505.

#### Doublet identification and analysis

We used Scrublet (Wolock et al., 2019) to identify potential doublets. We used the implementation in the Scanpy package (Wolf et al., 2018), with the 3 different experiments as “batch keys” and an empirically optimised threshold of 0.14. With these conditions, Scrublet classified as doublets 2,870 of the 80,387 cell barcodes. To independently identify doublets, we implemented a deep learning model with Solo 0.1 (Bernstein et al., 2020). We train the model with default settings except for a maximum number of 400 epochs. After subsetting the calculated doublet scores per cell, we filtered by the top putative doublets (>1.5). Full code implemented is available at the project repository.

We then preprocessed this dataset containing doublets to analyse their effects in cell clusters. This dataset contains 80,387 cells, of which 2,870 and 2,554 cells were considered doublets by Scrublet and Solo respectively with a 458 overlap. The processing eliminated genes with high counts using sc.pp.filter_genes with max_counts = 1000000. Then we calculated metrics using sc.pp.calculate_qc_metrics, sliced the matrix genes_by_counts < 700 and total_counts < 900, and normalised the matrix using sc.pp.normalize_total with a target_sum=1e4. We selected high variable genes using sc.pp.highly_variable_genes with n_top_genes = 18000, and sliced the matrix to contain only those genes, storing the raw in an adata.raw object. We then scaled the matrix with sc.pp.scale, performed pca with sc.tl.pca, constructed a kNN graph with sc.pp.neighbors, with 45 neighbours and 105 principal components, and calculated a UMAP visualisation with sc.tl.umap. We then plotted doublet cells identified by scrublet, solo and both in this visualisation. To determine if these doublets were major contributors to cell clusters, we run a clustering algorithm using sc.tl.leiden with resolution parameters 1, 2, 3 and 4. These gave respectively 47, 70, 83 and 89. We then calculated the proportions of doublets in each cluster using pandas and plotted them using matplotlib.

#### Parameter space optimisation

We optimised the parameter space iteratively running a custom function that processes the dataset accepting different arguments (minimum genes counts, maximum number of genes, maximum number of counts, number of top highly variable genes, number of neighbours, number of principal components, and leiden clustering resolution) and saves a figure report. The figure report includes a number of informative genes identified from preliminary analyses of the dataset because of their specific but also relatively complex expression pattern (PrileiEVm023936t1, PrileiEVm008309t1, PrileiEVm011741t1, PrileiEVm021316t1, PrileiEVm022250t1, PrileiEVm000325t1, PrileiEVm013699t1, PrileiEVm020595t1), as well as the UMAP visualisation and the number of clusters obtained. This function was run on the 75,421 cell dataset with the doublets excluded. We sequentially run iterations of this function trying the following values: minimum genes counts (30, 40, 50, 60, 70, 80, 90, 100), maximum number of genes (300, 400, 500, 600, 700, 800, 900, 1000), maximum number of counts (500, 600, 700, 800, 900, 1000, 1100, 1200), number of top highly variable genes (4000, 6000, 8000, 10000, 12000, 14000, 18000, 22000), number of neighbours (15, 25, 35, 45, 55, 65, 75, 85), number of principal components (15, 25, 45, 65, 85, 105, 125, 145), with the other parameters in each iteration remaining fixed in standard values (50, 700, 900, 18000, 45, 105, 1 respectively). We examined the result of each run to visually inspect the complexity of the cluster visualisation and the number of clusters obtained.

#### Single cell transcriptomic analysis

We processed the final dataset with conditions optimised from our parameter space exploration. We started this processing with the matrix of 75,421 cells after doublet exclusion. The processing eliminated genes with high counts using sc.pp.filter_genes with max_counts = 1000000. We calculated metrics using sc.pp.calculate_qc_metrics, sliced the matrix genes_by_counts < 700 and total_counts < 900. This step eliminated 203 cells, giving us our final dataset of 75,218 cells. We normalised the matrix using sc.pp.normalize_total with a target_sum=1e4. We selected high variable genes using sc.pp.highly_variable_genes with n_top_genes = 18000, and sliced the matrix to contain only those genes, storing the raw in an adata.raw object. We then scaled the matrix with sc.pp.scale, performed pca with sc.tl.pca, constructed a kNN graph with sc.pp.neighbors, with 45 neighbours and 105 principal components, and calculated a UMAP visualisation with sc.tl.umap (min_dist=0.5, spread = 1, alpha = 1, gamma = 1.0). We run the Leiden clustering algorithm using sc.tl.leiden with resolutions 0.5, 1, 1.5 and 2, which gave 34, 50, 60 and 70 clusters respectively. We calculated marker genes for each cluster using sc.tl.rank_genes_groups, using the clusters of obtained with all 4 resolution parameters, and using both the Wilcoxon (method='wilcoxon') and the Logistic Regression (method='logreg') We selected resolution 1.5 for further downstream analyses.

#### PAGA

For the PAGA analysis we removed unannotated clusters. Preliminary analyses indicated that these small clusters interfere with the PAGA analysis. The expression of *piwi* in them is relatively high, suggesting that they could be subpopulations of *piwi*+ cells, but they also had specific markers, suggesting that they contain differentiated types. Our interpretation of these clusters is that they are rare cell types that, at this resolution, are clustered together with their progenitor including *piwi*+ cells. The presence of these confounds the PAGA analysis. Alternatively, they could represent leftover doublets. Altogether they are a small number of cells. To identify these clusters we calculated the mean of each transcript from the adata.X object and ranked the expression of stem cell genes by obtaining the average mean expression of PrileiEVm016887t1, PrileiEVm004300t1, PrileiEVm003567t1, PrileiEVm016982t1, and PrileiEVm003521t1. This generated a rank of clusters that contained *piwi*+ cells including clusters 1, 2 and 8 (with 7103, 6557 and 2587 cells) but also contained smaller clusters with ~2 orders of magnitude fewer cells, including clusters 51, 57, 58, 48, 43, 53, 52, 50, 47 (with 85, 50, 41, 153, 191, 74, 77, 117 and 154 cells). We decided to leave unnanotated clusters with ranked expression > 0.0500 and fewer than 175 cells, which gave us the final list of clusters 46, 47, 48, 50, 51, 52, 53, 54, 56, 57, and 58.

We then performed a PAGA analysis with and without these clusters. We selected a random cell from cluster 1 as roon using adata.uns['iroot'] = np.flatnonzero(adata.obs[clusteringlayer] == '1')[0] We then used the Scanpy implementation of Difussion Pseudotime, using sc.tl.dpt(adata, n_branchings=1). We then run sc.tl.paga on the selected clusters of resolution 1.5. Our PAGA plot is generated with sc.pl.paga(adata, threshold=0.25, solid_edges='connectivities_tree', root=1, layout='rt', node_size_scale=2, node_size_power=0.9, max_edge_width=3, fontsize=20). The Potency Score was plotted using sc.pl.paga with similar parameters and passing color = 'degree_solid', cmap = 'viridis' arguments to the function.

#### CPM calculation

Raw counts were extracted with a custom Python script (see project repository) that slices the raw unprocessed matrix to contain only the cells that are present in the processed matrix. The cluster information is transferred from the processed matrix to the unprocessed matrix using a pandas script. Then the sum of all counts for each gene in each cluster is obtained using numpy on the matrix. The resulting raw summed counts dataset was normalised by pseudobulk “library size” using the

`DESeqDataSetFromMatrix()` function with parameter `design = ~ condition` and the `counts()` function with parameter `normalised = TRUE` from the package DESeq2 (Love et al., 2014).

#### Co-occurrence analysis

Cell type co-occurrence analysis was performed using the function `treeFromEnsembleClustering()` from the code provided by Levy and collaborators (Levy et al., 2021) using parameters: `h = c(0.75,0.95), clustering_algorithm = "hclust", clustering_method = "average", cor_method = "pearson", p = 0.1, n = 1000, bootstrap=FALSE`. Briefly, we performed 1000 iterations of cross-cell type Pearson correlation using 90% downsampling of highly variable genes (FC > 1.5) followed by hierarchical clustering of cell types. Co-occurring pairs of cell types across iterations are quantified to generate a co-occurrence matrix that is hierarchically clustered to generate the cell type tree.

#### Transcription factor annotation

The resulting TransDecoder-translated proteome of *Pristina* was queried for evidence of Transcription Factor (TF) homology using (i) InterProScan (Jones et al., 2014) against the Pfam (Mistry et al., 2021), PANTHER (Thomas et al., 2022), and (ii) SUPERFAMILY (Gough et al., 2001; Pandurangan et al., 2019) domain databases with standard parameters, (iii) using BLAST reciprocal best hits (Moreno-Hagelsieb and Latimer, 2008) against swissprot transcription factors (UniProt, 2023), and (iv) using OrthoFinder (Emms and Kelly, 2019) with standard parameters against a set of model organisms (Human, Zebrafish, Mouse, Drosophila) with well annotated transcription factor databases (following AnimalTFDB v3.0) (Hu et al., 2019). For the latter, a given Pristina gene was counted as TF if at least another TF gene from any of the species belonged to the same orthogroup as the Pristina gene. The different sources of evidence were pooled together and we kept those Pristina genes with at least two independent sources of TF evidence. Every TF gene was assigned a class based on their sources of evidence.

#### Transcription factor analysis

The CPM table was subset to retrieve the *Pristina* TFs, and gene expression across cell types was scaled and visualised using the ComplexHeatmap package (Gu et al., 2016). To analyse the TFs at the class level, for a given class X, we calculated the median and average coefficient of variation (CV) of class X across cell types, the number of genes pertaining to class X, and the cumulative number, average, and median counts of class X. We visualised the relationship between CV and number of genes using the base and ggplot2 packages (https://ggplot2.tidyverse.org/) in R v4.0.3 (https://www.R-project.org/)

We did a multivariate analysis two-way ANOVA to detect differences of TF expression between cell clusters, TF classes, and the interaction of the two. TF counts were aggregated at the broad cell cluster level and we kept only those TFs from classes with four or more annotated genes. The

ANOVA was run using aov(), followed by Tukey comparison of means using TukeyHSD(). The most prominent classes explaining differences across cell clusters were retrieved by quantifying and sorting the results of the Tukey test.

To represent these differences visually, we calculated the expression prominence of each TF class (the sum of counts per gene). For a given TF class X, we defined the prominence of class X across cell clusters as the addition of the counts of all genes of class X in each cluster, divided by the number of genes of class X expressed at each cluster. The resulting matrix was normalised and visualised using a custom ggplot2 wrapper function in R v4.0.3.

#### WGCNA analysis

We ran WGCNA (Langfelder and Horvath, 2008) using a subset of the CPM table genes with CV > 1 and softPower 5 estimated after visualising the Scale-Free Topology Model Fit. Adjacency and Topological Overlapped (TOM) matrices were calculated using standard parameters. For dynamic cutting of the tree, we chose 100 genes as minimum module size. Provided the discrete expression of gene modules, these were named and recolored manually following a similar criterion than when naming cell clusters. The resulting classification in modules was used to reorder the expression dataset, and the dataset was represented for visualisation using ComplexHeatmap (Gu et al., 2016).

To calculate the association between TF classes and modules, we calculated the connectivity of each TF gene to each module eigengene. For a given TF class X, we quantified the number of genes of class X with a connectivity equal or higher than 0.5 to each module eigengene. The resulting matrix was normalised and represented using the package ComplexHeatmap.

WGCNA graphs were constructed using the TOM matrix and pruning from sparse interactions using an arbitrary low threshold of connectedness (> 0.01). A subset of the resulting graph (>0.4) (hereafter ".4 graph") was used for exploratory analysis using the igraph package (Csárdi and Nepusz, 2006) and the Kamada-Kawai layout algorithm (Kamada and Kawai, 1989) with parameters `maxiter = 100 * NUM_GENES_GRAPH , kkconst = NUM_GENES_GRAPH`, where NUM_GENES_GRAPH is the number of genes present in the .4 graph. Connected component membership was calculated using the function components() from the igraph package, and its percent of agreement with the WGCNA module membership was calculated using the adjusted Rand Index implementation adjustedRandIndex() from the package mclust (Scrucca et al., 2016). The .4 graph was subdivided into subgraphs corresponding to the connected components using a custom wrapper function that implements the induced_subgraph() function of the igraph package. Centrality of the TFs belonging to each separate sub-graph was calculated using the closeness() function from the igraph package in a custom wrapper function, and visualised using ggplot2.

We used a less stringent subset of the .01 graph (> 0.2, rather than 0.4) to analyse cross-module connections. Using a custom wrapper function, a 'gene x module' matrix was constructed counting how many genes from each module are direct neighbours to a given gene x, and normalised by dividing the number of connections of gene x to each module by the size of the module that gene x is part of. These numbers were later aggregated at the module level to retrieve the number of normalised cross-connections between modules. The resulting matrix was transformed into a graph using graph_from_adjacency_matrix() from igraph with parameters `mode = "upper", weighted = TRUE, diag = FALSE`, and the number of cross-connections was used for edge size to highlight the largest amounts of cross-connections.

#### Limma analysis

Differential Gene Expression Analysis was performed using the edgeR (Robinson et al., 2010) and limma (Ritchie et al., 2015) R packages, and the pseudo bulk count matrix. Briefly, we made a distinction between piwi-positive and piwi-negative cell clusters in order to retrieve the genes that are differentially expressed in piwi-positive cells. A DGE object was created using the counts table and a sample information table with the aforementioned distinction, as well as a model matrix. The dataset was filtered using the filterByExpr() function from edgeR, and normalised using the voom() method from limma. Linear modelling was done using the lmFit() function with the model matrix (all piwi-positive vs non-piwi-positive), and statistics were calculated with the eBayes() function. The results were plotted using the EnhancedVolcano (https://github.com/kevinblighe/EnhancedVolcano) and ggplot2 packages.

#### Gene Ontology analysis

Gene Ontology (GO) analyses were performed using the R package topGO (Alexa et al., 2006) and the `elim` method using a custom wrapper function. GO terms with less than three significantly annotated genes were discarded. Unless otherwise specified, we chose the totality of *Pristina* genes as the gene universe population to compare against.

#### *Piwi*+ cell transcription factor analysis

For this analysis we used raw counts extracted at the broad cell type group and normalised them as described above. Then, the relative enrichment of expression in each broad cell type group was calculated by subtracting the log cpm (with a pseudocount) of each cell type from the mean log cpm (with a pseudocount) of the remaining broad types. We then filtered this table to contain only TFs and extracted those with the higher coefficients of variation (cv >1). We used this table to sort the top 200 TFs with higher levels of enrichment (log ratios) in *piwi*+ cells compared to all other cell types.

#### Epigenetic factor analysis

We extracted lists of epigenetic factor components from https://epifactors.autosome.org/ (Marakulina et al., 2023; Medvedeva et al., 2015), containing human protein sequences. We then blasted those against the translated *Pristina* transcriptome using tblastn. We manually curated the selection of top hits for each epigenetic factor, and annotated those that are annotated as members of more than one epigenetic regulation complex (Supplementary File 14).

#### HCR *in situ* hybridization

For Hybridization Chain Reaction (HCR), previously published protocols (Kuehn et al., 2022) were used with mainly modifications for *Pristina leidyi* fixation and 1st day of the protocol, based on the species colorimetric in situ hybridization protocols (Özpolat and Bely, 2015). Specifically, samples were fixed in 4% PFA for 40-45 minutes, dehydration/rehydration steps in methanol were skipped, and after washes in 1x PBSt, HCR protocol was carried out on the same day. Day 1 of the original colorimetric *in situ* hybridization protocol (which includes pronase digestion, acetylation, and post-fixation) was found to be essential for successful *Pristina leidyi* HCR results. The entire protocol can be accessed in https://github.com/BDuyguOzpolat/Pristina_leidyi-protocols

Selection of markers and designing probesets: For each cell cluster, top expression markers with coding sequence length of 700 bp or longer were listed (for compatibility with HCR probe design). Probesets were designed for 1 or 2 of these markers per cluster using the Özpolat Lab algorithm (https://github.com/rwnull/insitu_probe_generator) (Kuehn et al., 2022). The sequences used for probe design were confirmed to be in 5' to 3' orientation using https://web.expasy.org/translate. For each probeset, the lower probe pair limit was 11 and the upper limit was 34 pairs. Complete list and sequences of probesets, along with the associated initiator information can be found in Supplementary File 6.

Hairpin amplifiers were ordered from Molecular Instruments (Choi et al., 2018). For all HCR experiments a combination of the following hairpin-fluorophore conjugations were used: B1-546, B2-488, B3-647, B4-594, B3-594, B4-647, B4-488.

#### Confocal imaging

Confocal imaging was carried out using Zeiss LSM710 and LSM780 microscopes at the microscopy facility at Marine Biological Laboratory. For each set of HCRs, control tubes were included. Controls did not have any probes, but had hairpins, in order to assess the unspecific background signal (Supplementary Figure 6). Image analyses and editing were carried out in Fiji (Schindelin et al., 2012), panels and schematics were prepared using Adobe Illustrator.

#### Nuclei area quantification

For comparison of *vigilin*+ cell nuclei size with the other cell types in the area, we used the nuclear staining in confocal Z-stacks, and measured the area for each nucleus using Fiji (Schindelin et al., 2012). 3 different worm samples were used for measurements. Samples were imaged as z-stacks, and the nuclei to be measured were picked from 5 focal planes across the stack. At each focal plane 5 nuclei for *vigilin*+ cells and 5 nuclei from the nearby cells that are negative for *vigilin* were measured (25 nuclei each group, 50 nuclei per sample). The R Wilcoxon rank sum test (wilcox.test) was used for statistical analyses using R to compare the two groups.

#### Subclustering *piwi*+ clusters

We selected *piwi*+ cells by selecting cells in clusters 1, 2 and 8, including 16,247 cells, and we reanalysed them alone from the raw unprocessed matrix. We calculated metrics using sc.pp.calculate_qc_metrics, and normalised the matrix using sc.pp.normalize_total with a target_sum=1e4. We selected high variable genes using sc.pp.highly_variable_genes with n_top_genes = 18000, and sliced the matrix to contain only those genes, storing the raw in an adata.raw object. We then scaled the matrix with sc.pp.scale, performed pca with sc.tl.pca, constructed a kNN graph with sc.pp.neighbors, with 35 neighbours and 25 principal components, and calculated a UMAP visualisation with sc.tl.umap (min_dist=0.5, spread = 1, alpha = 1, gamma = 1.0). We run the Leiden clustering algorithm using sc.tl.leiden with resolutions 0.4, which gives 10 clusters. We calculated marker genes for each cluster using sc.tl.rank_genes_groups using both the Wilcoxon (method='wilcoxon') and the Logistic Regression (method='logreg').

#### Scores

To calculate gene scores we used the Scanpy function sc.tl.score_genes with a control size equal to the length of the gene list and a number of bins equal to 25.

### Supplementary References

**Alexa, A., Rahnenfuhrer, J.*, et al.***, (2006). Improved scoring of functional groups from gene expression data by decorrelating GO graph structure. *Bioinformatics* **22**, 1600-1607.

**Bely, A.E., Wray, G.A.**, (2001). Evolution of regeneration and fission in annelids: insights from engrailed- and orthodenticle-class gene expression. *Development* **128**, 2781-2791.

**Bernstein, N.J., Fong, N.L.*, et al.***, (2020). Solo: Doublet Identification in Single-Cell RNA-Seq via Semi-Supervised Deep Learning. *Cell Syst* **11**, 95-101 e105.

**Buchfink, B., Reuter, K.*, et al.***, (2021). Sensitive protein alignments at tree-of-life scale using DIAMOND. *Nat Methods* **18**, 366-368.

**Cantalapiedra, C.P., Hernandez-Plaza, A.*, et al.***, (2021). eggNOG-mapper v2: Functional Annotation, Orthology Assignments, and Domain Prediction at the Metagenomic Scale. *Mol Biol Evol* **38**, 5825-5829.

**Choi, H.M.T., Schwarzkopf, M.*, et al.***, (2018). Third-generation in situ hybridization chain reaction: multiplexed, quantitative, sensitive, versatile, robust. *Development* **145**.

**Csárdi, G., Nepusz, T.**, (2006). The igraph software package for complex network research.

**Dobin, A., Davis, C.A.*, et al.***, (2013). STAR: ultrafast universal RNA-seq aligner. *Bioinformatics* **29**, 15-21.

**Emms, D.M., Kelly, S.**, (2019). OrthoFinder: phylogenetic orthology inference for comparative genomics. *Genome Biol* **20**, 238.

**Garcia-Castro, H., Kenny, N.J.*, et al.***, (2021). ACME dissociation: a versatile cell fixation-dissociation method for single-cell transcriptomics. *Genome Biol* **22**, 89.

**Gilbert, D.G.**, (2019). Longest protein, longest transcript or most expression, for accurate gene reconstruction of transcriptomes? *bioRxiv*, 829184.

**Gough, J., Karplus, K.*, et al.***, (2001). Assignment of homology to genome sequences using a library of hidden Markov models that represent all proteins of known structure. *J Mol Biol* **313**, 903-919.

**Gu, Z., Eils, R.*, et al.***, (2016). Complex heatmaps reveal patterns and correlations in multidimensional genomic data. *Bioinformatics* **32**, 2847-2849.

**Hu, H., Miao, Y.R.*, et al.***, (2019). AnimalTFDB 3.0: a comprehensive resource for annotation and prediction of animal transcription factors. *Nucleic Acids Res* **47**, D33-D38.

**Jones, P., Binns, D.*, et al.***, (2014). InterProScan 5: genome-scale protein function classification. *Bioinformatics* **30**, 1236-1240.

**Kamada, T., Kawai, S.**, (1989). An algorithm for drawing general undirected graphs. *Information Processing Letters* **31**, 7-15.

**Kuehn, E., Clausen, D.S.*, et al.***, (2022). Segment number threshold determines juvenile onset of germline cluster expansion in Platynereis dumerilii. *J Exp Zool B Mol Dev Evol* **338**, 225-240.

**Langfelder, P., Horvath, S.**, (2008). WGCNA: an R package for weighted correlation network analysis. *BMC Bioinformatics* **9**, 559.

**Levy, S., Elek, A.*, et al.***, (2021). A stony coral cell atlas illuminates the molecular and cellular basis of coral symbiosis, calcification, and immunity. *Cell* **184**, 2973-2987 e2918.

**Love, M.I., Huber, W.*, et al.***, (2014). Moderated estimation of fold change and dispersion for RNA-seq data with DESeq2. *Genome Biol* **15**, 550.

**Marakulina, D., Vorontsov, I.E.*, et al.***, (2023). EpiFactors 2022: expansion and enhancement of a curated database of human epigenetic factors and complexes. *Nucleic Acids Res* **51**, D564-D570.

**Martin, M.**, (2011). Cutadapt removes adapter sequences from high-throughput sequencing reads. *EMBnet.journal* **17**, 3.

**Medvedeva, Y.A., Lennartsson, A.*, et al.***, (2015). EpiFactors: a comprehensive database of human epigenetic factors and complexes. *Database (Oxford)* **2015**, bav067.

**Mistry, J., Chuguransky, S.*, et al.***, (2021). Pfam: The protein families database in 2021. *Nucleic Acids Res* **49**, D412-D419.

**Moreno-Hagelsieb, G., Latimer, K.**, (2008). Choosing BLAST options for better detection of orthologs as reciprocal best hits. *Bioinformatics* **24**, 319-324.

**Nyberg, K.G., Conte, M.A.*, et al.***, (2012). Transcriptome characterization via 454 pyrosequencing of the annelid Pristina leidyi, an emerging model for studying the evolution of regeneration. *BMC Genomics* **13**, 287.

**Özpolat, B.D., Bely, A.E.**, (2015). Gonad establishment during asexual reproduction in the annelid Pristina leidyi. *Dev Biol* **405**, 123-136.

**Pandurangan, A.P., Stahlhacke, J.*, et al.***, (2019). The SUPERFAMILY 2.0 database: a significant proteome update and a new webserver. *Nucleic Acids Res* **47**, D490-D494.

**Ritchie, M.E., Phipson, B.*, et al.***, (2015). limma powers differential expression analyses for RNA-sequencing and microarray studies. *Nucleic Acids Res* **43**, e47.

**Robinson, M.D., McCarthy, D.J.*, et al.***, (2010). edgeR: a Bioconductor package for differential expression analysis of digital gene expression data. *Bioinformatics* **26**, 139-140.

**Schindelin, J., Arganda-Carreras, I.*, et al.***, (2012). Fiji: an open-source platform for biological-image analysis. *Nat Methods* **9**, 676-682.

**Scrucca, L., Fop, M.*, et al.***, (2016). mclust 5: Clustering, Classification and Density Estimation Using Gaussian Finite Mixture Models. *The R journal* **8 1**, 289-317.

**Thomas, P.D., Ebert, D.*, et al.***, (2022). PANTHER: Making genome-scale phylogenetics accessible to all. *Protein Sci* **31**, 8-22.

**UniProt, C.**, (2023). UniProt: the Universal Protein Knowledgebase in 2023. *Nucleic Acids Res* **51**, D523-D531.

**Wolf, F.A., Angerer, P.*, et al.***, (2018). SCANPY: large-scale single-cell gene expression data analysis. *Genome Biol* **19**, 15.

**Wolock, S.L., Lopez, R.*, et al.***, (2019). Scrublet: Computational Identification of Cell Doublets in Single-Cell Transcriptomic Data. *Cell Syst* **8**, 281-291 e289.
