## Supplementary figures and images for "Annelid adult cell type diversity and their pluripotent cellular origins"

### Supp File 12.pdf

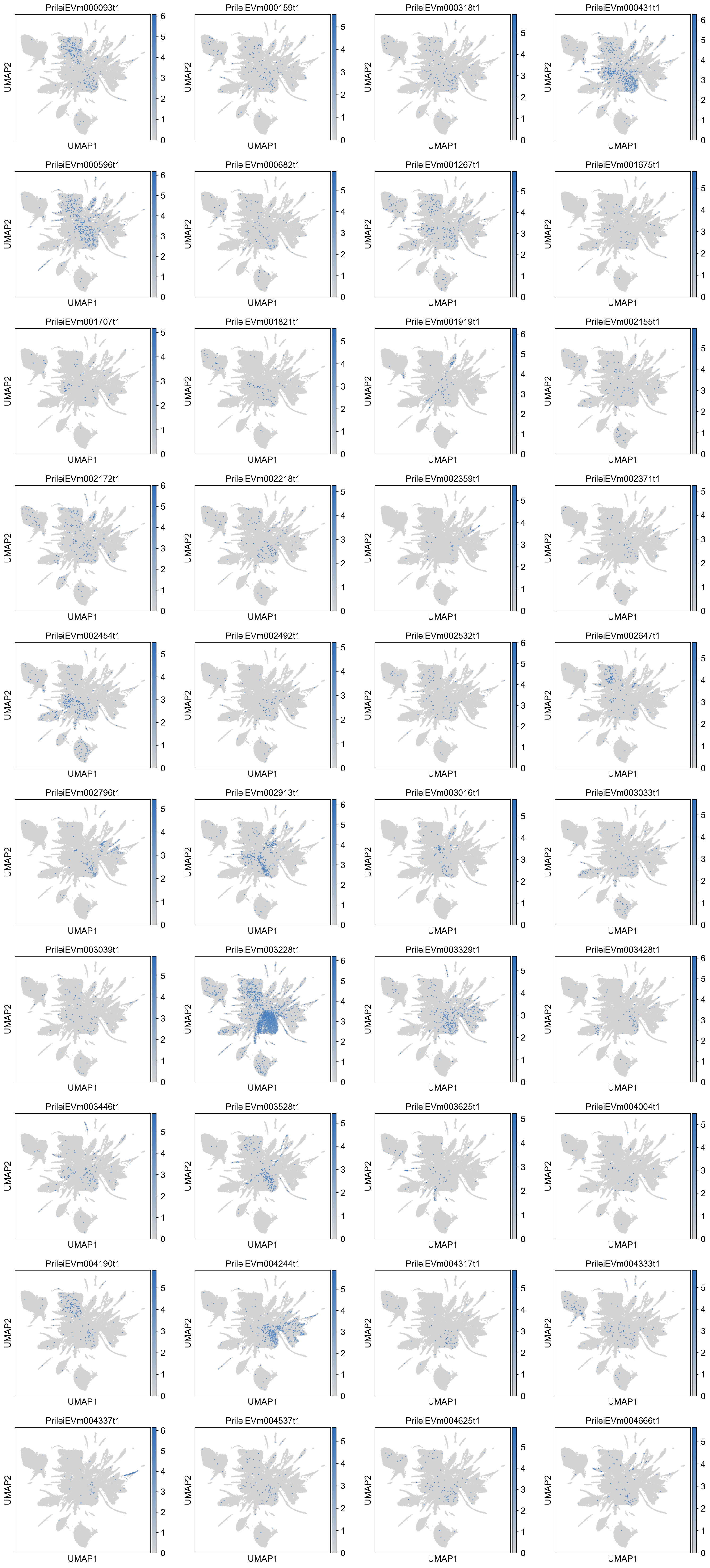

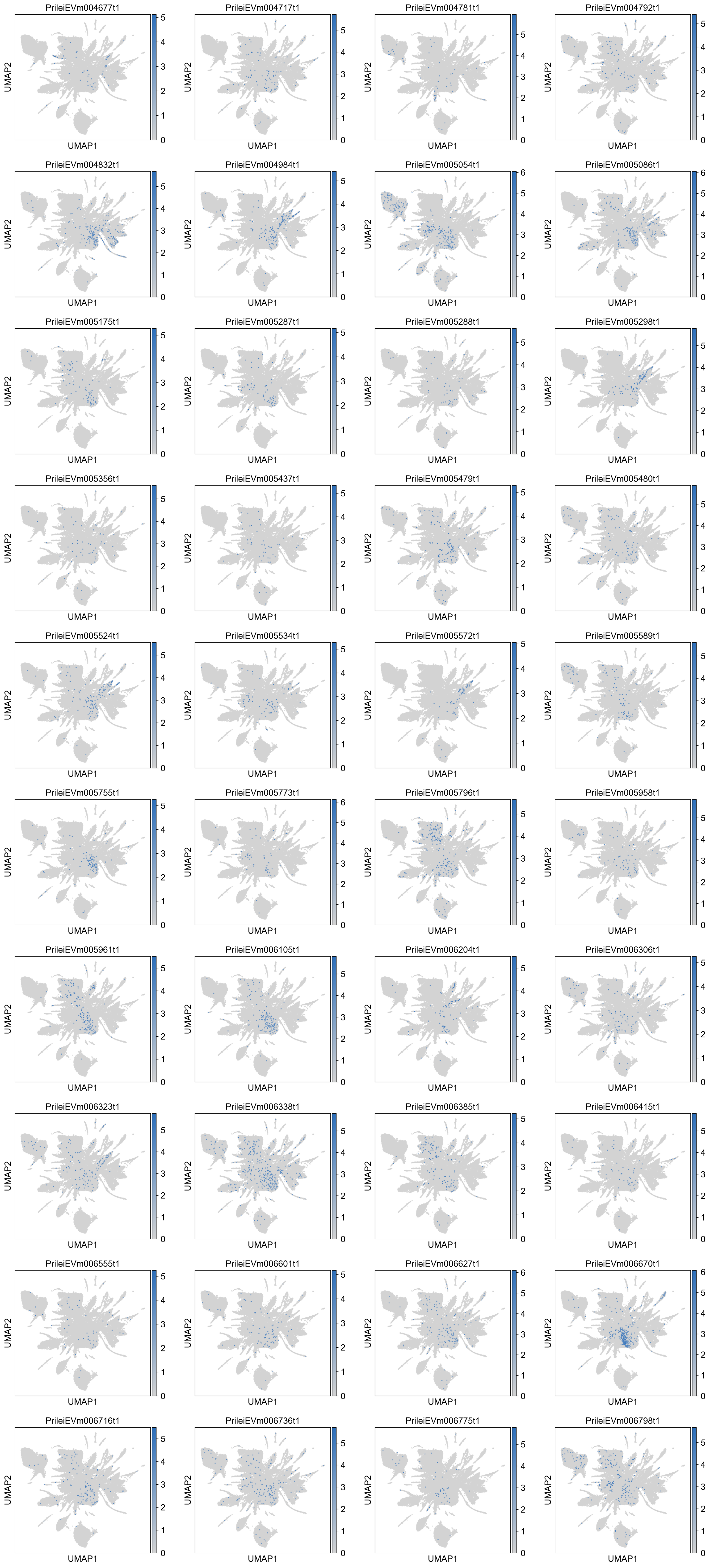

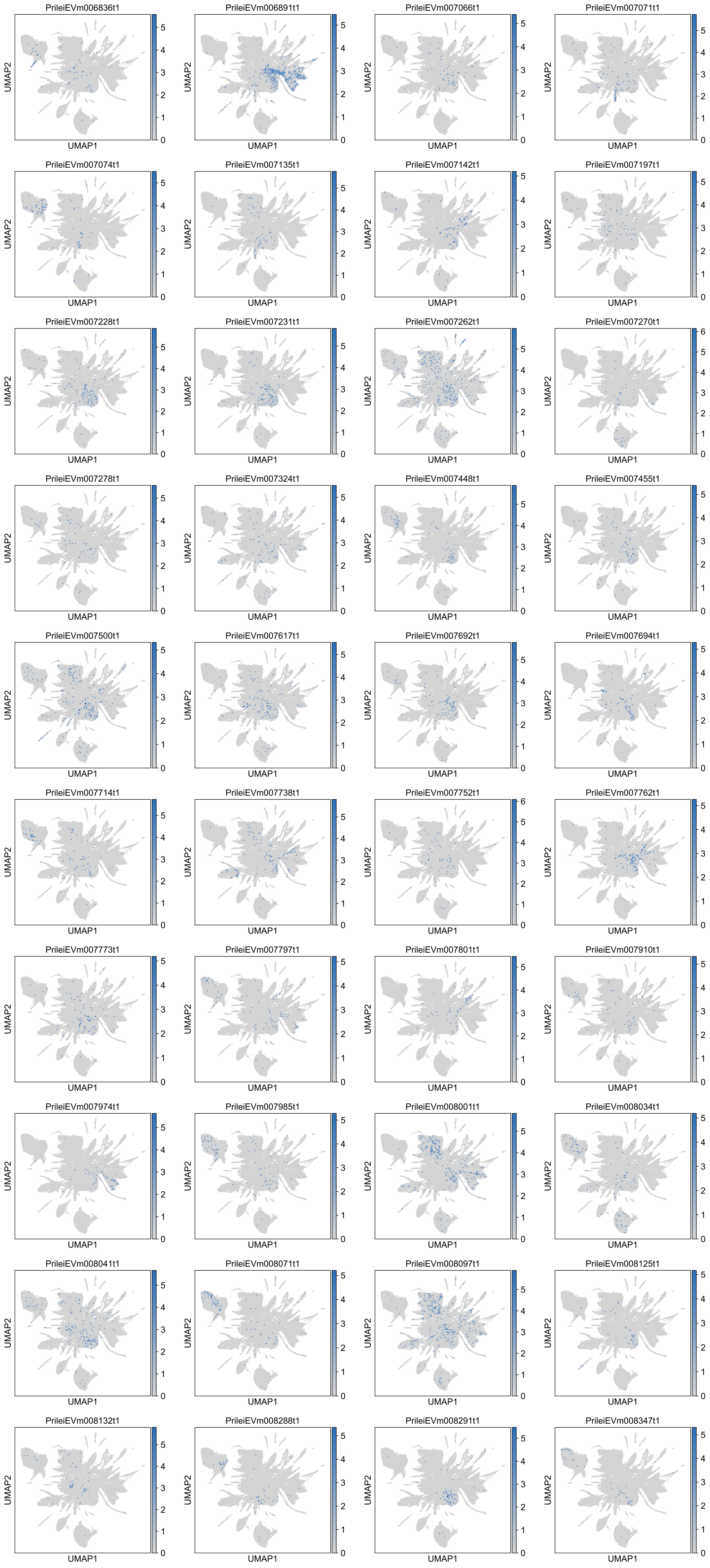

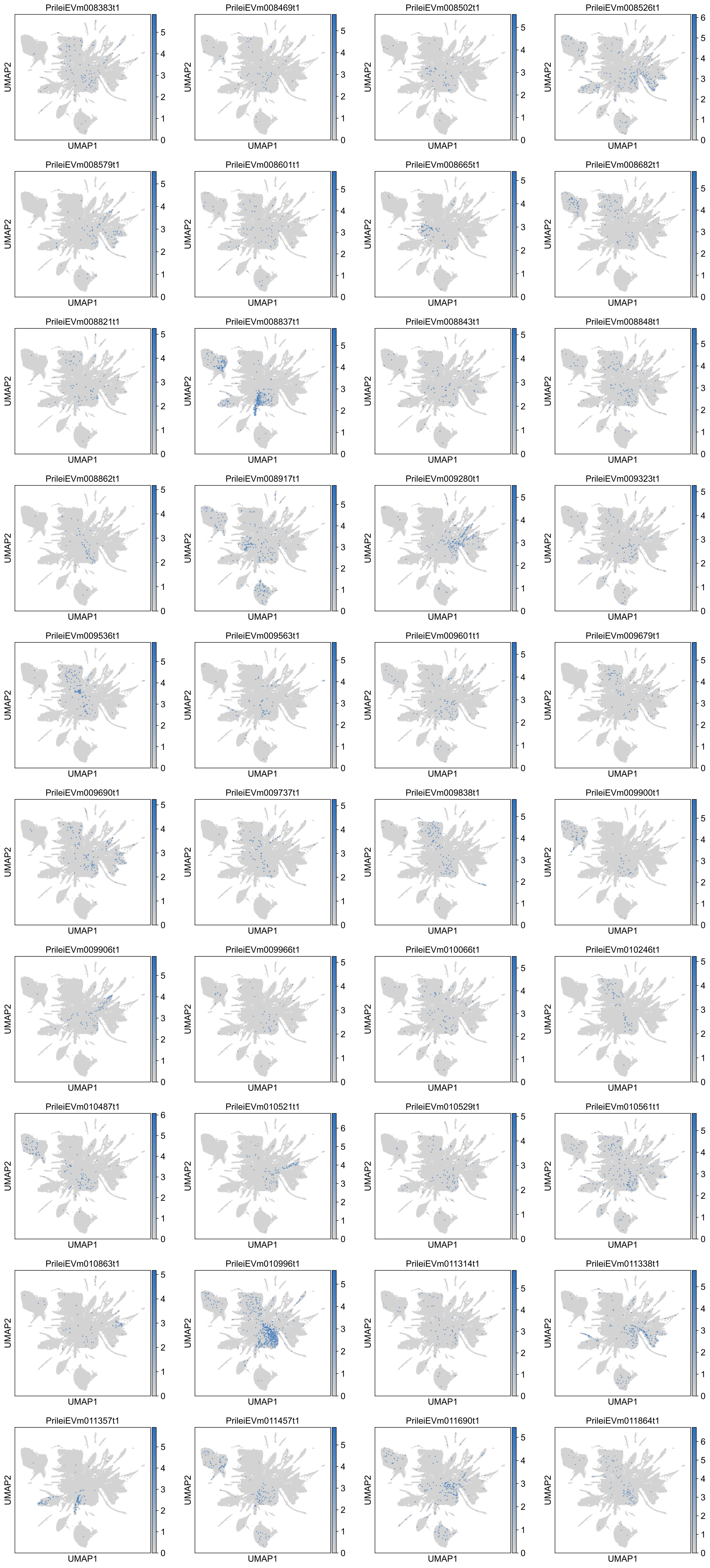

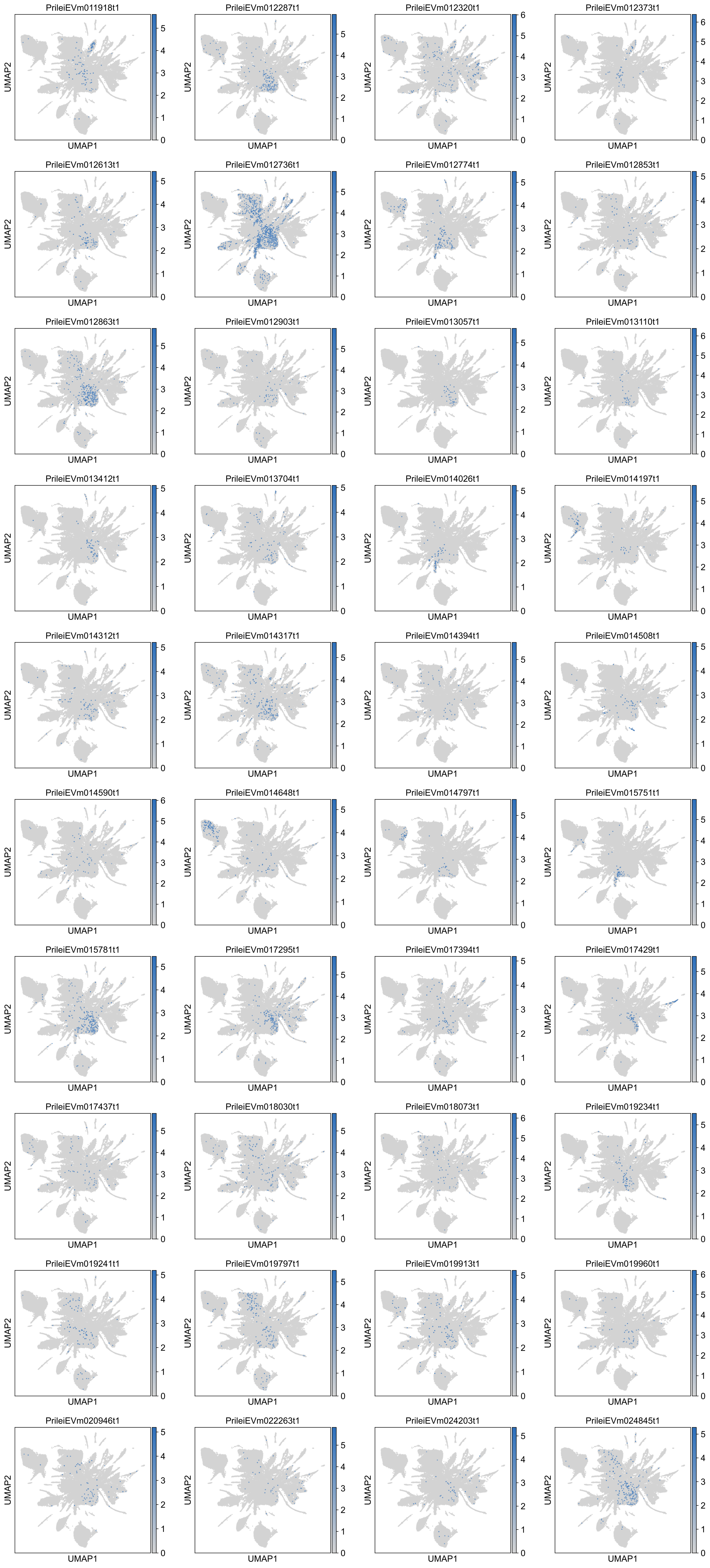

### Supplementary Figures

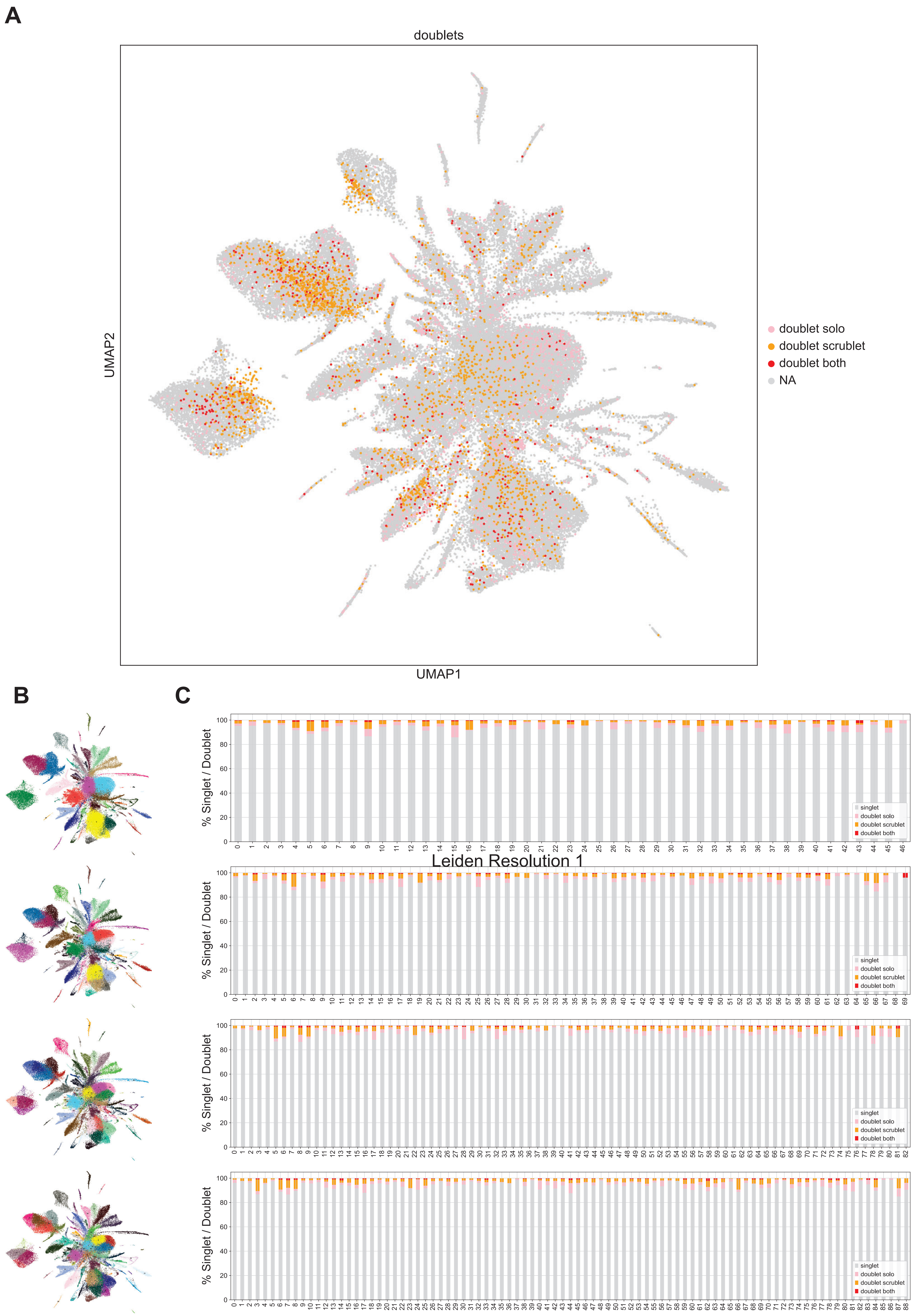

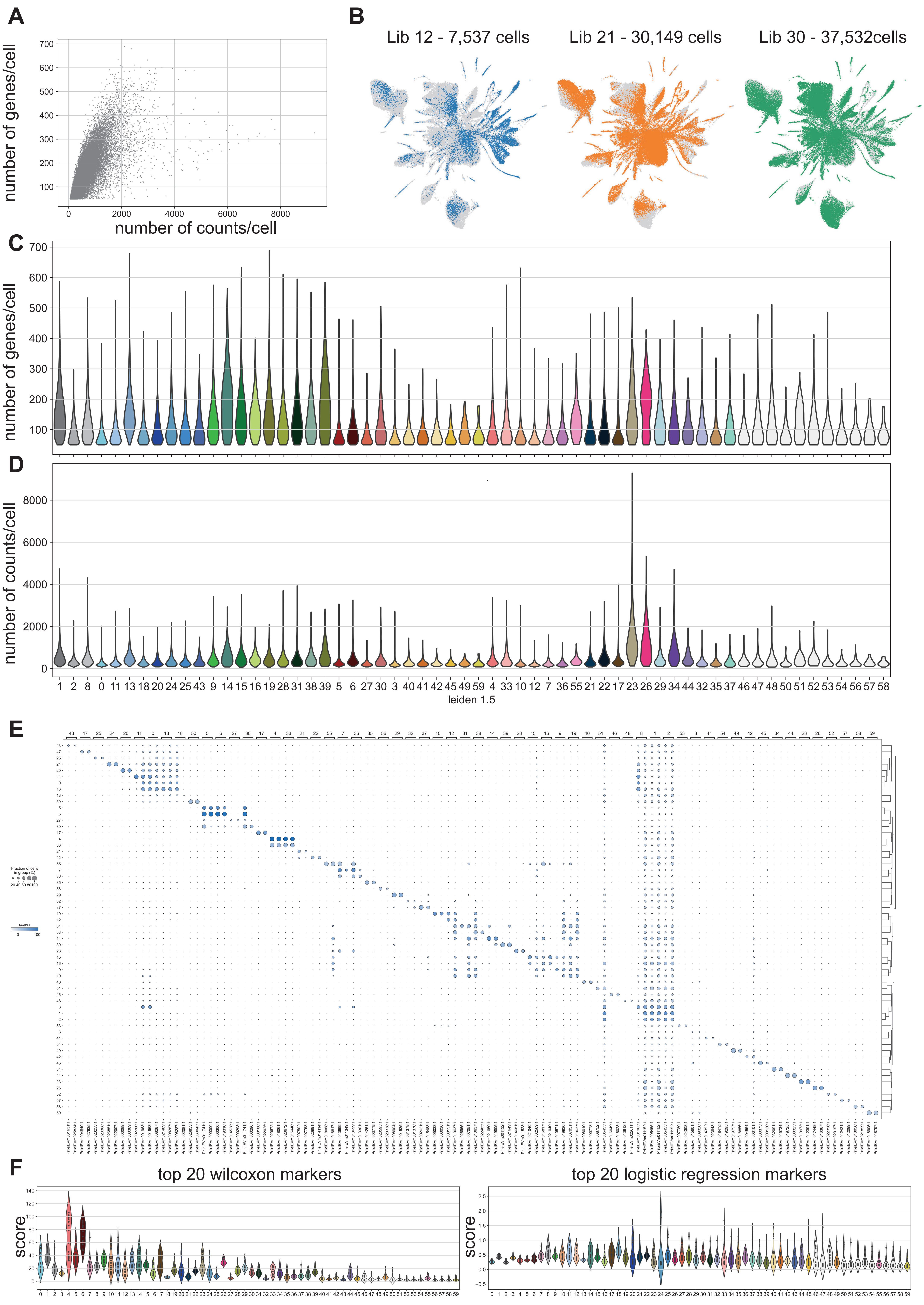

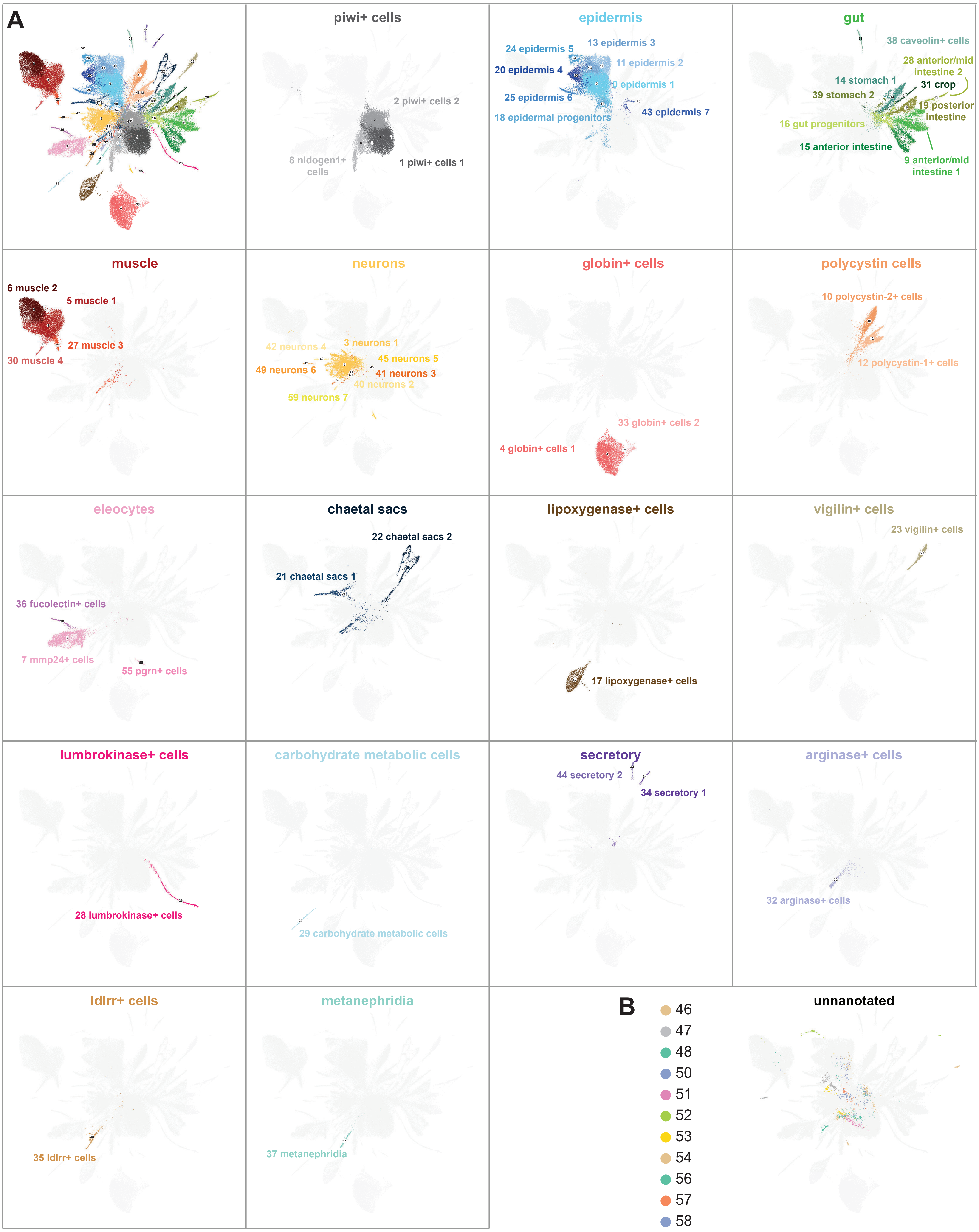

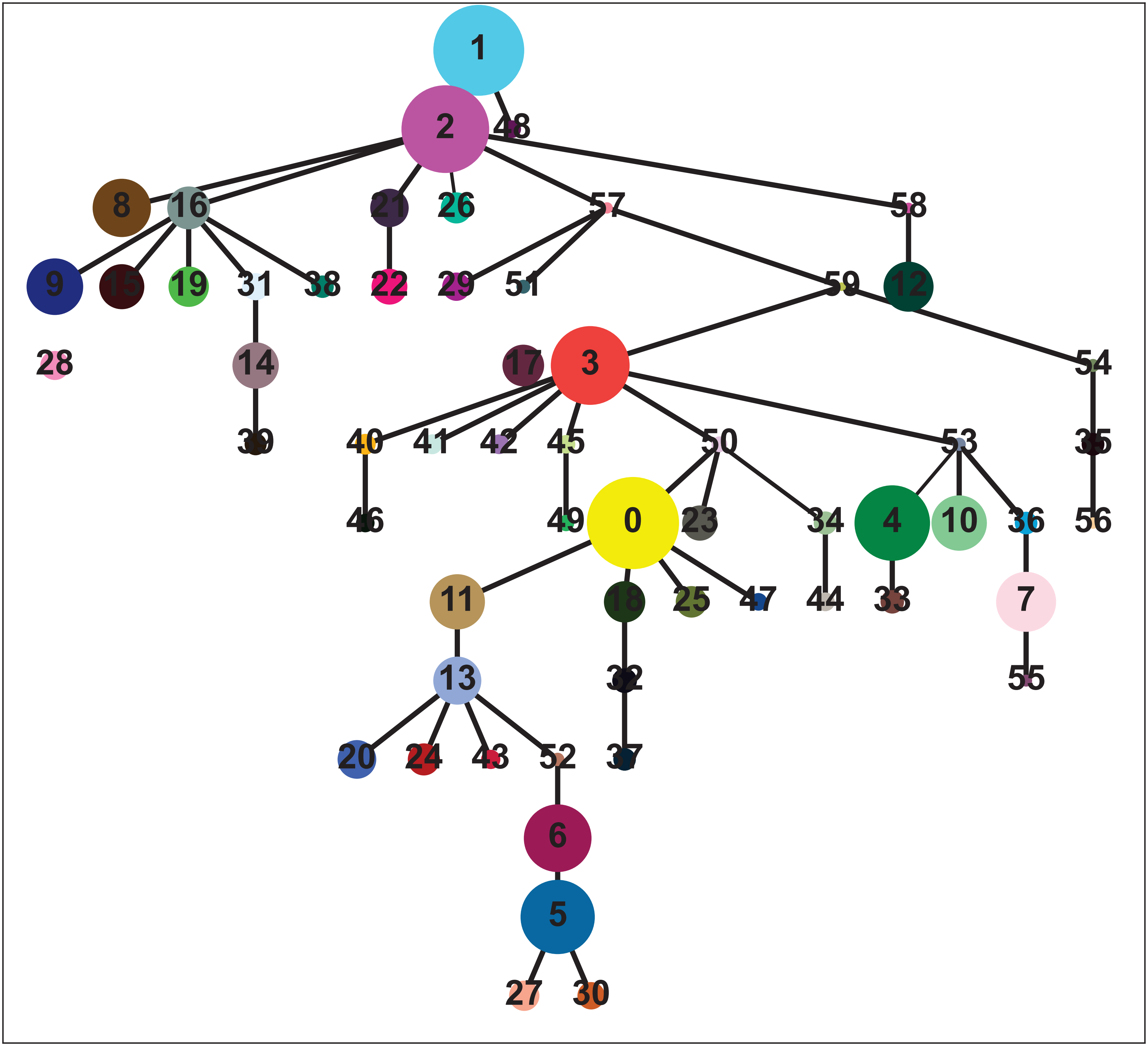

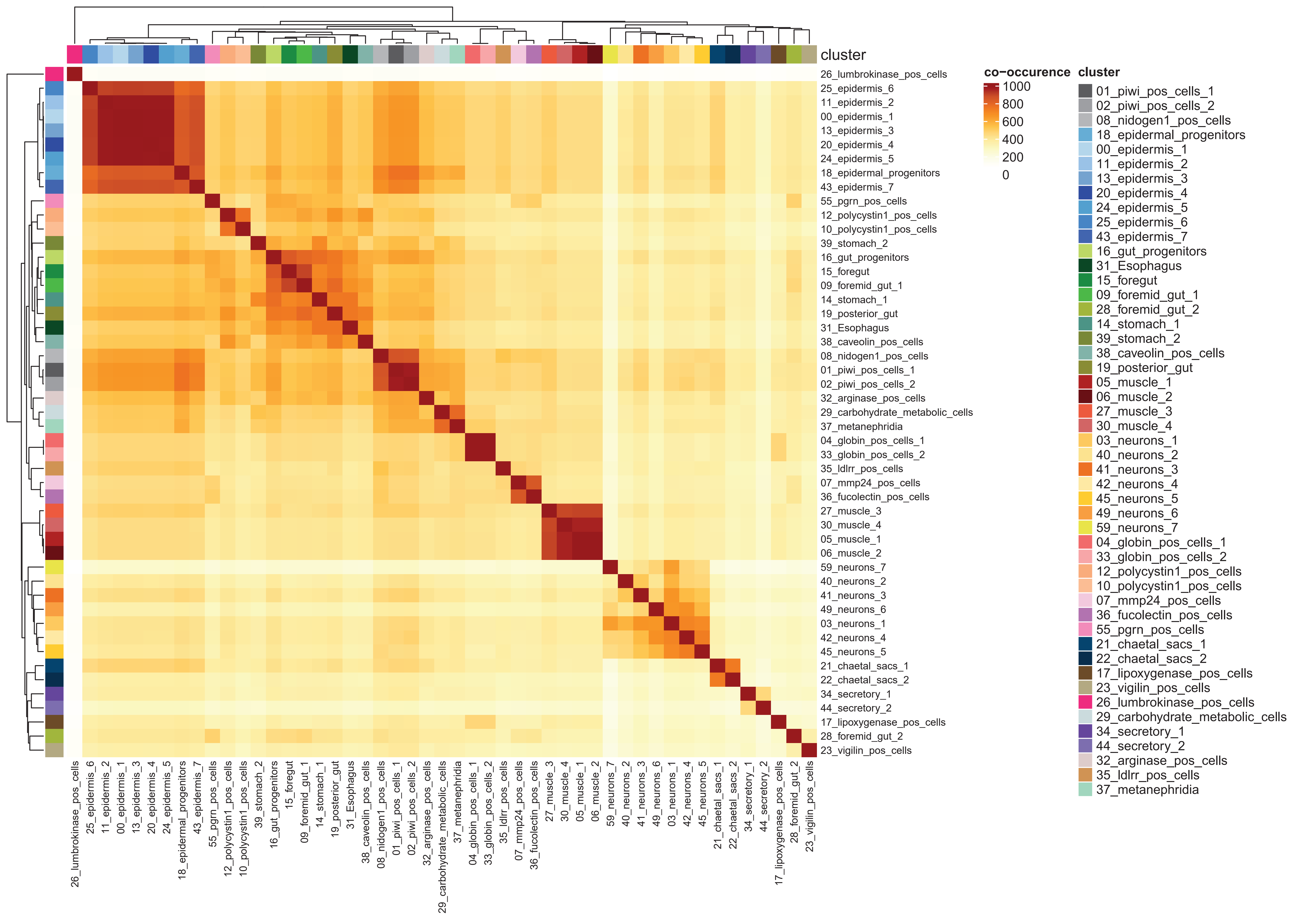

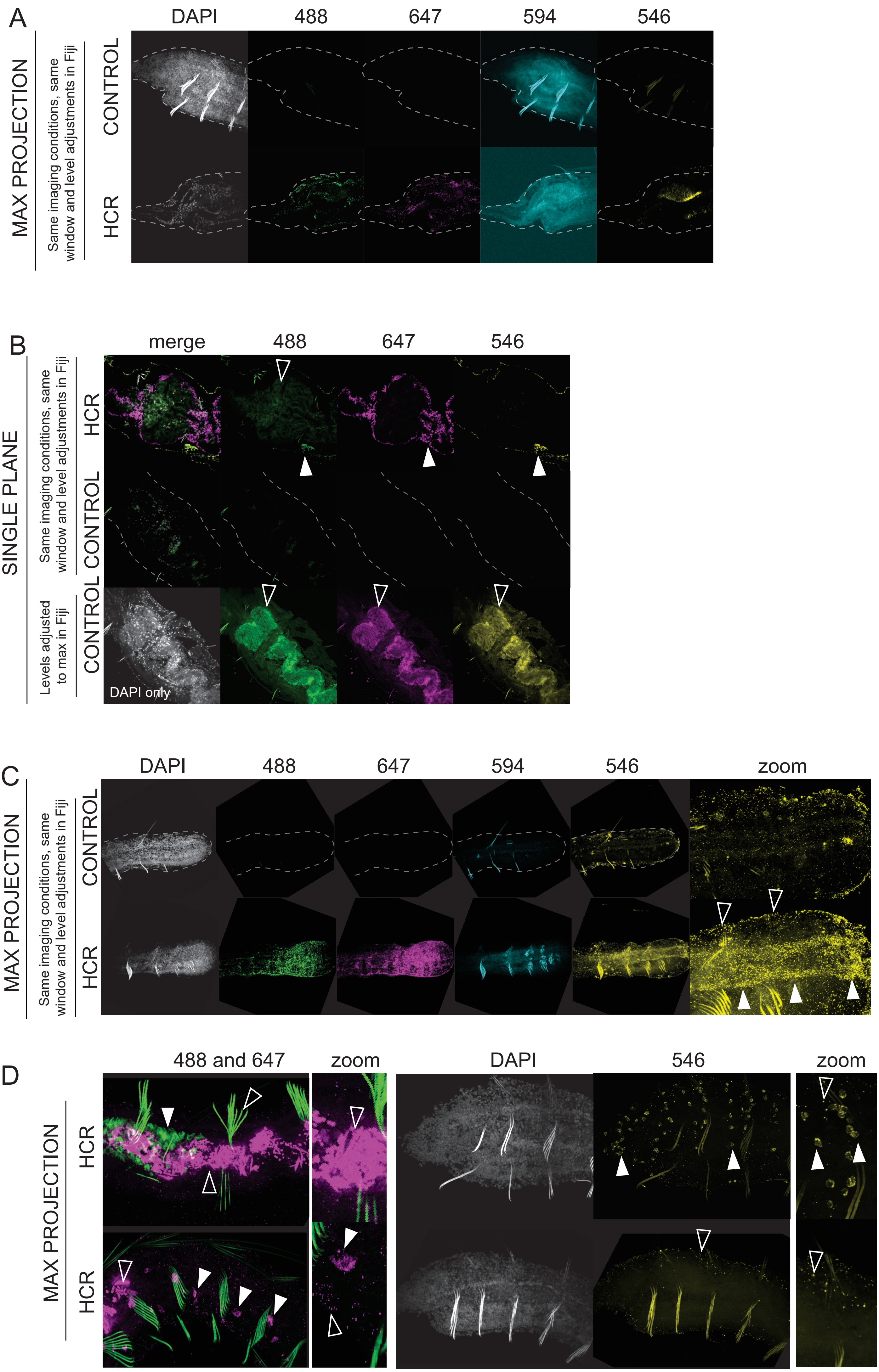

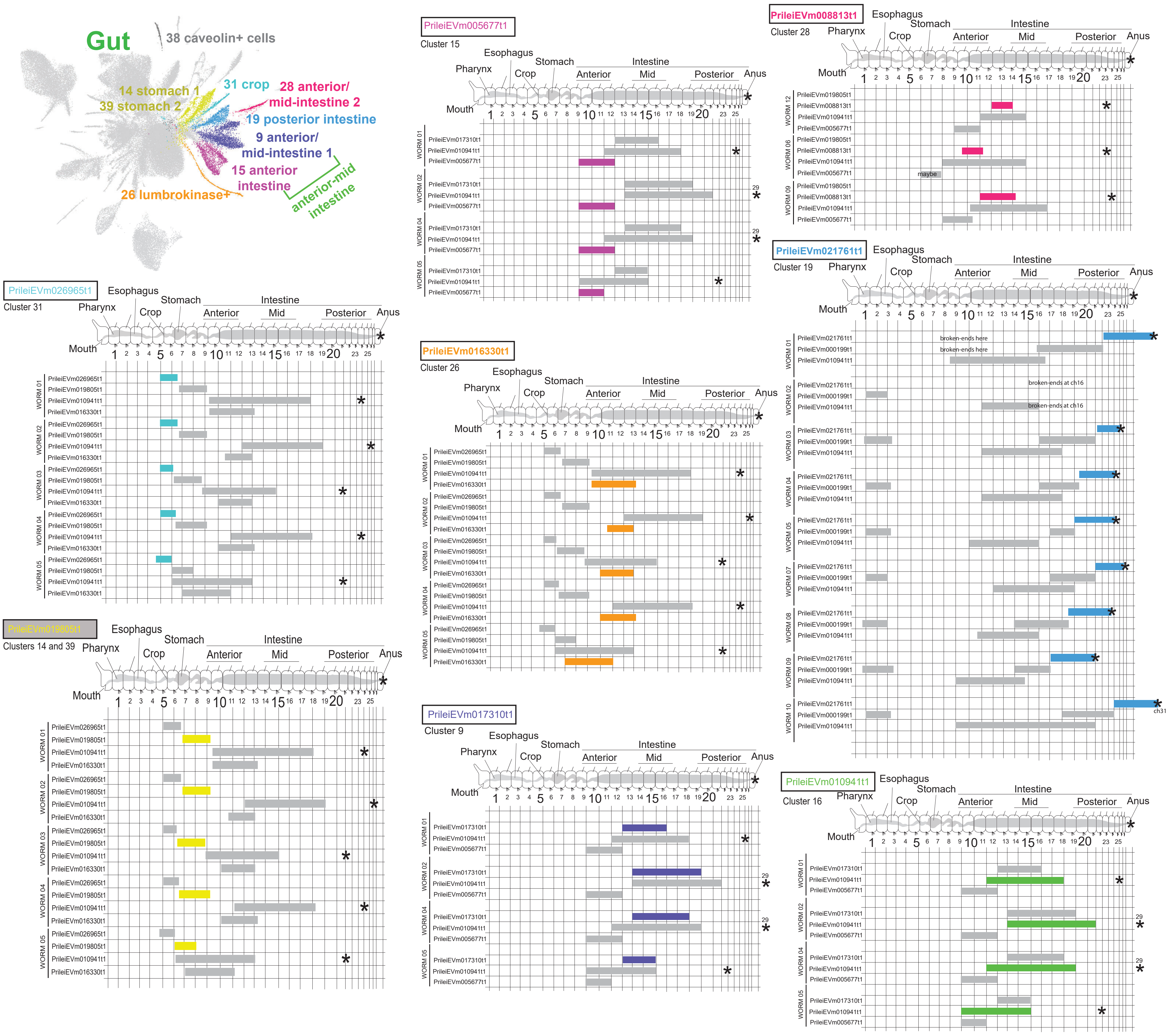

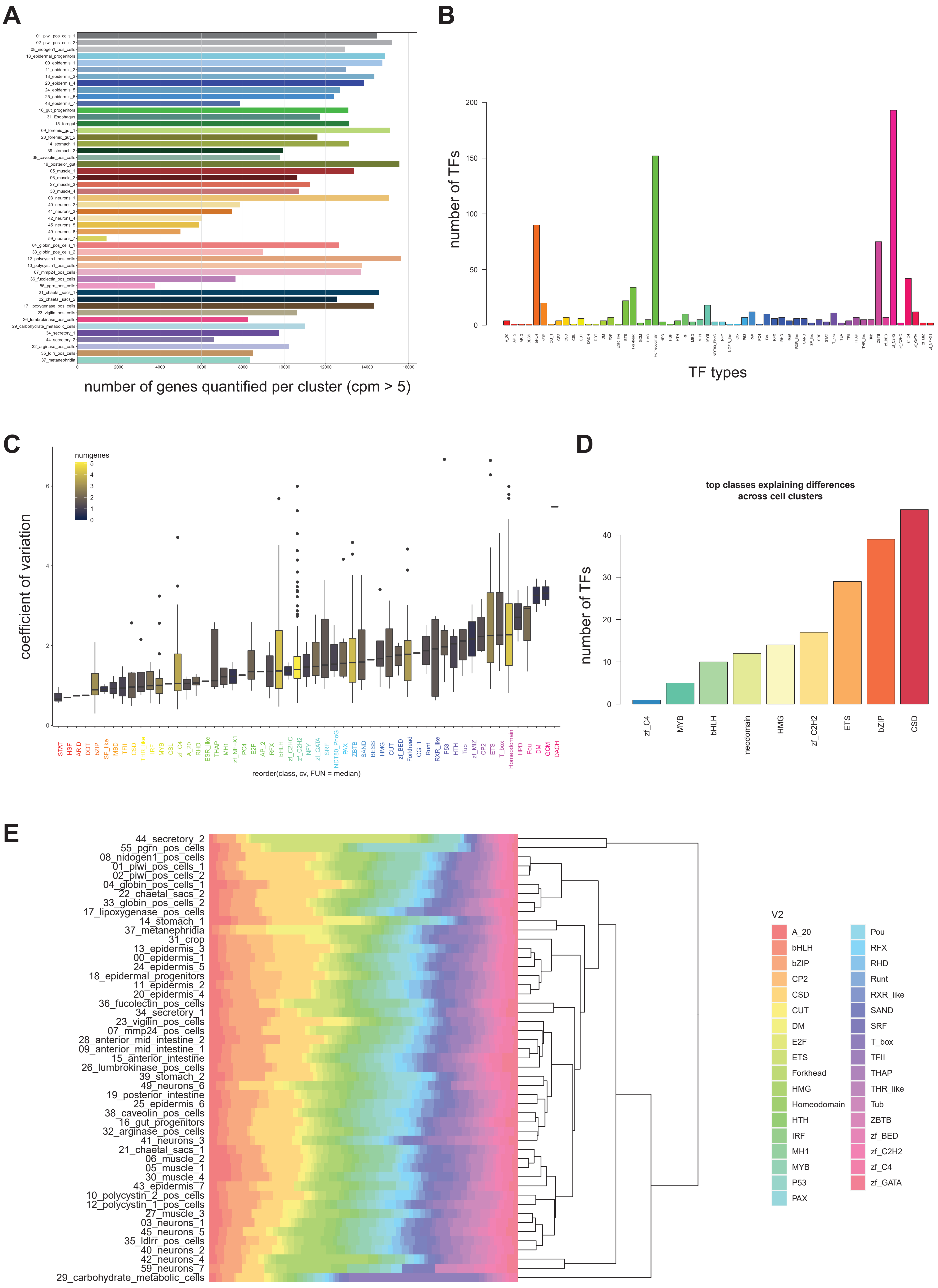

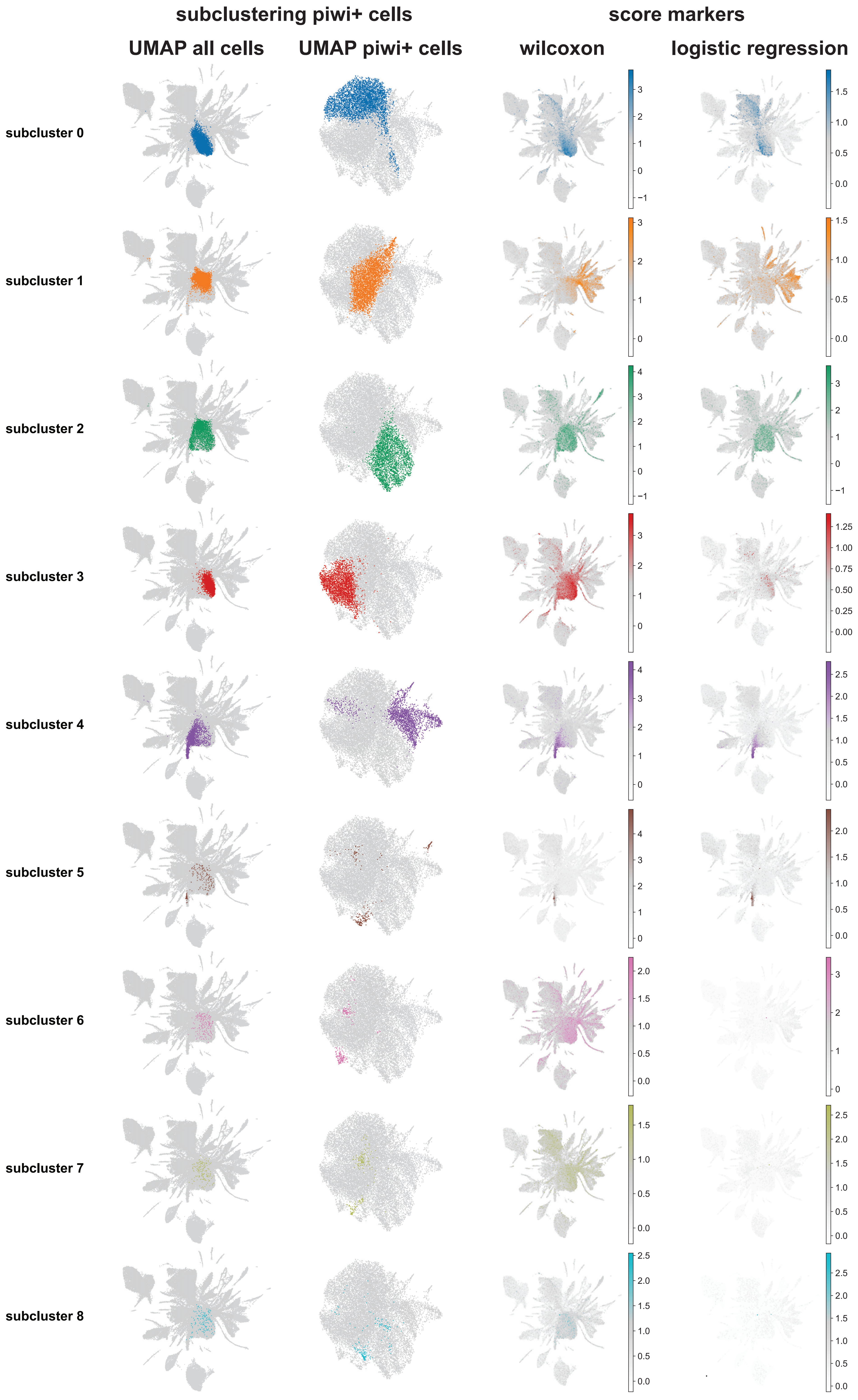
