## Supplementary material for "Annelid adult cell type diversity and their pluripotent cellular origins": Supp File 2.pdf

### minimum gene counts

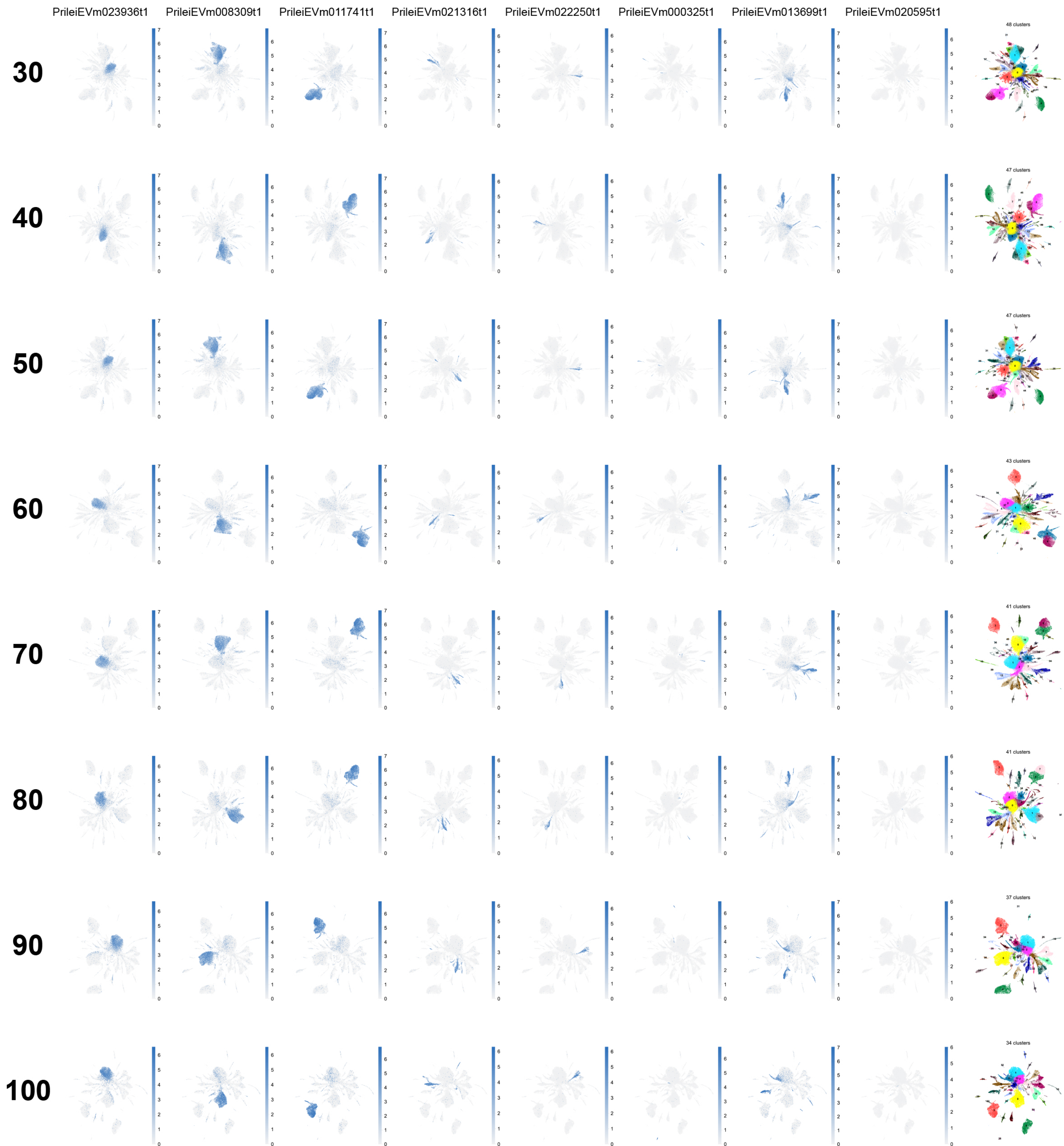

### maximum number of genes

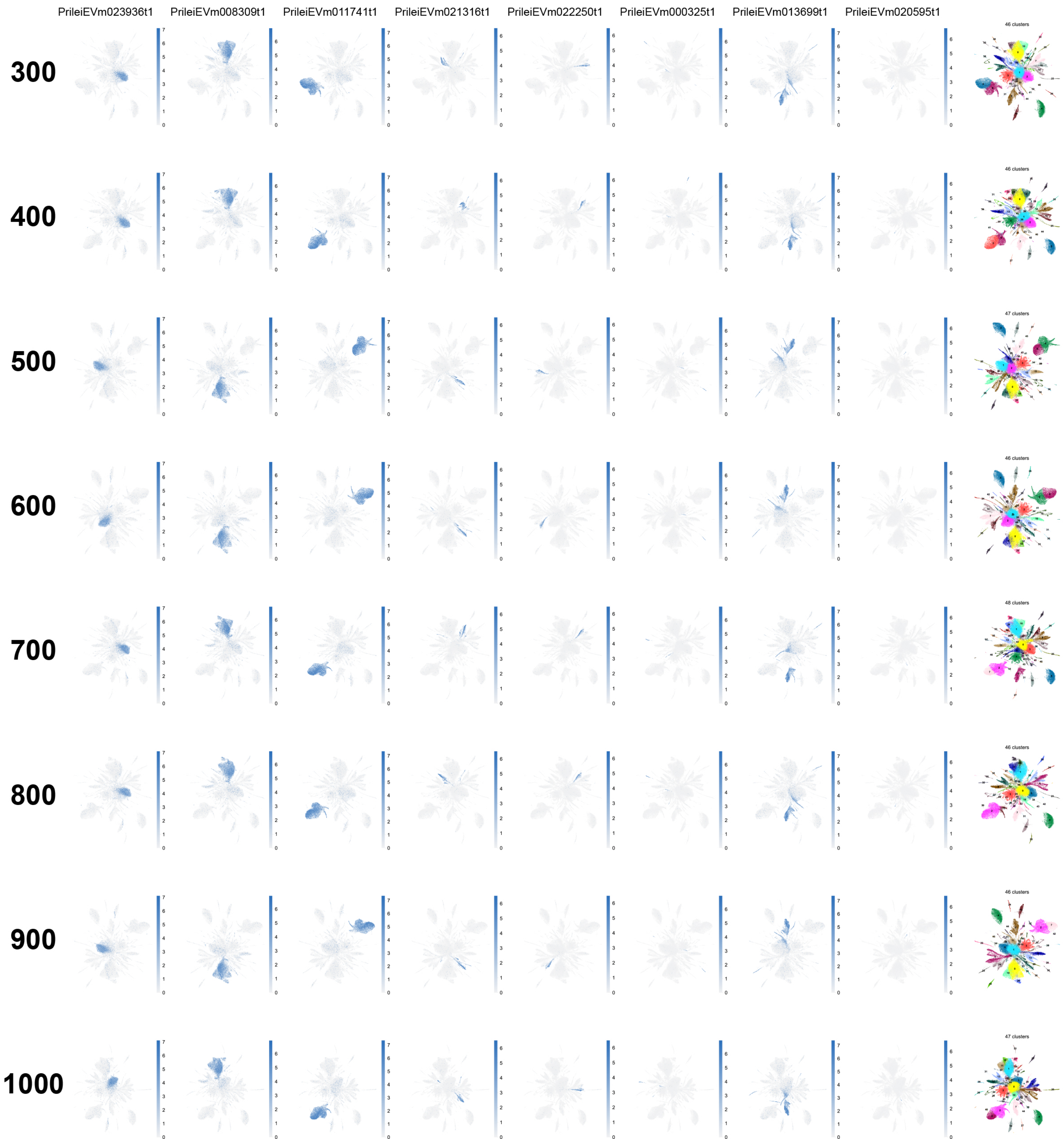

### maximum number of counts

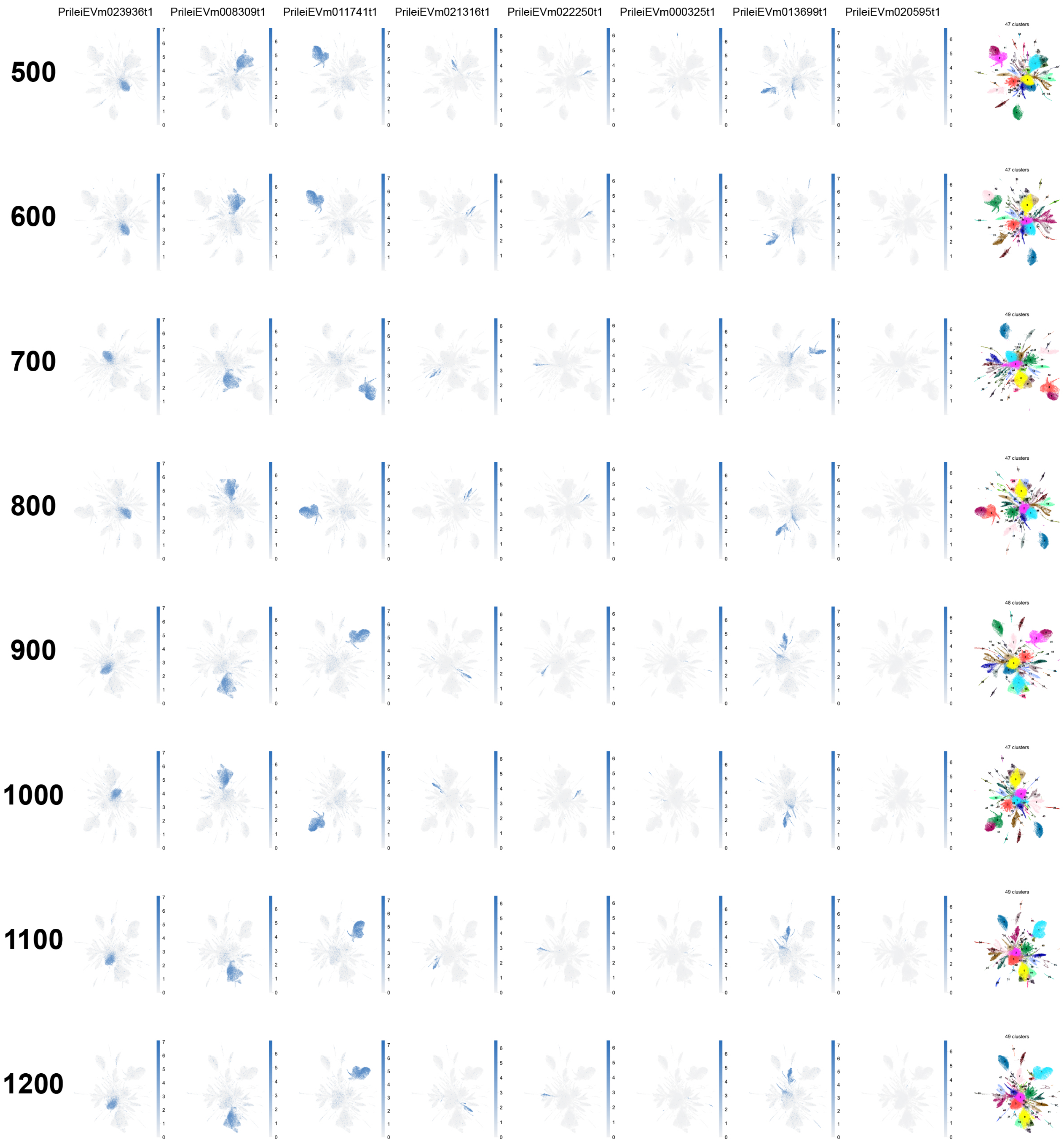

### number of top highly variable genes

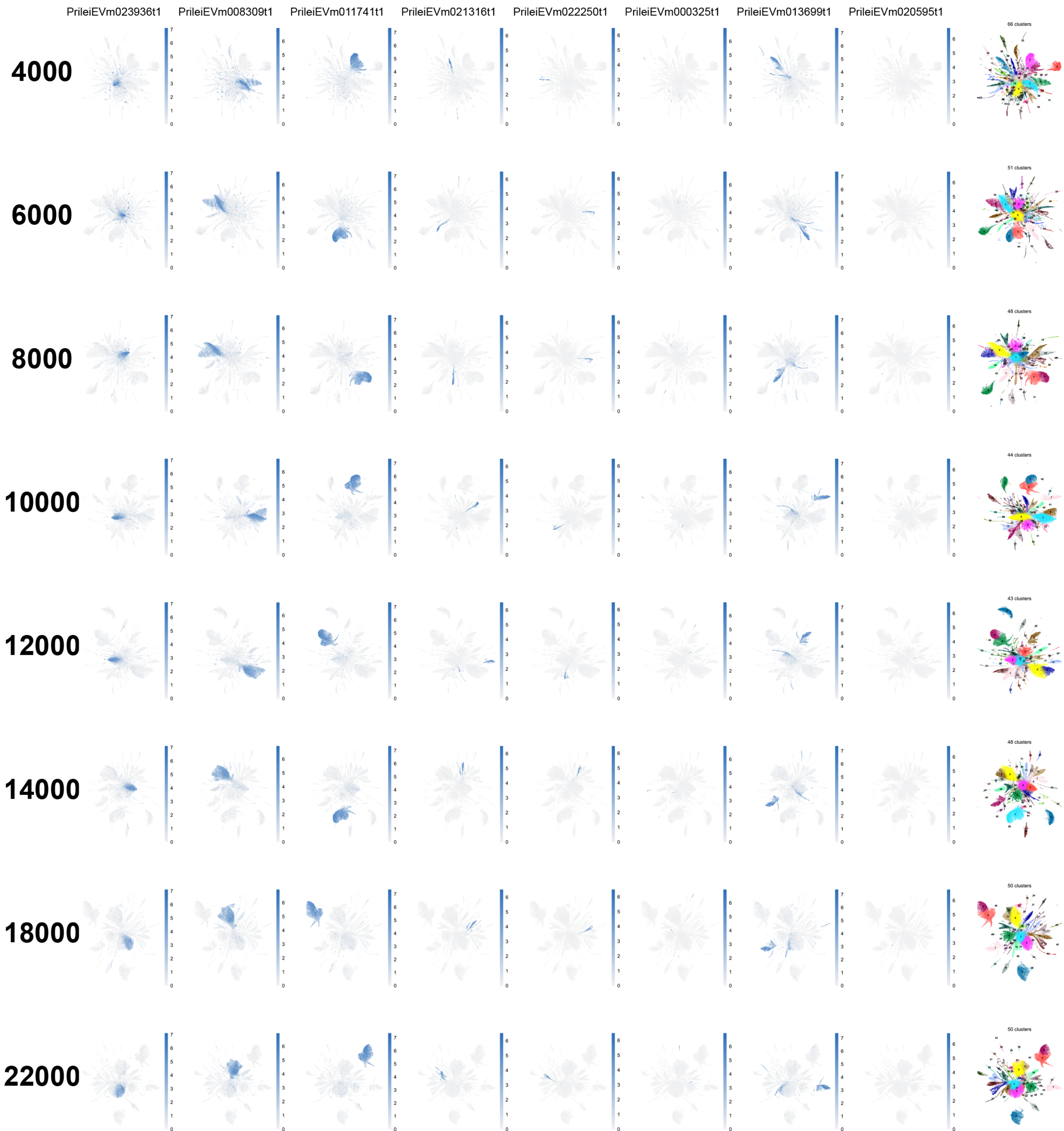

### number of neighbours

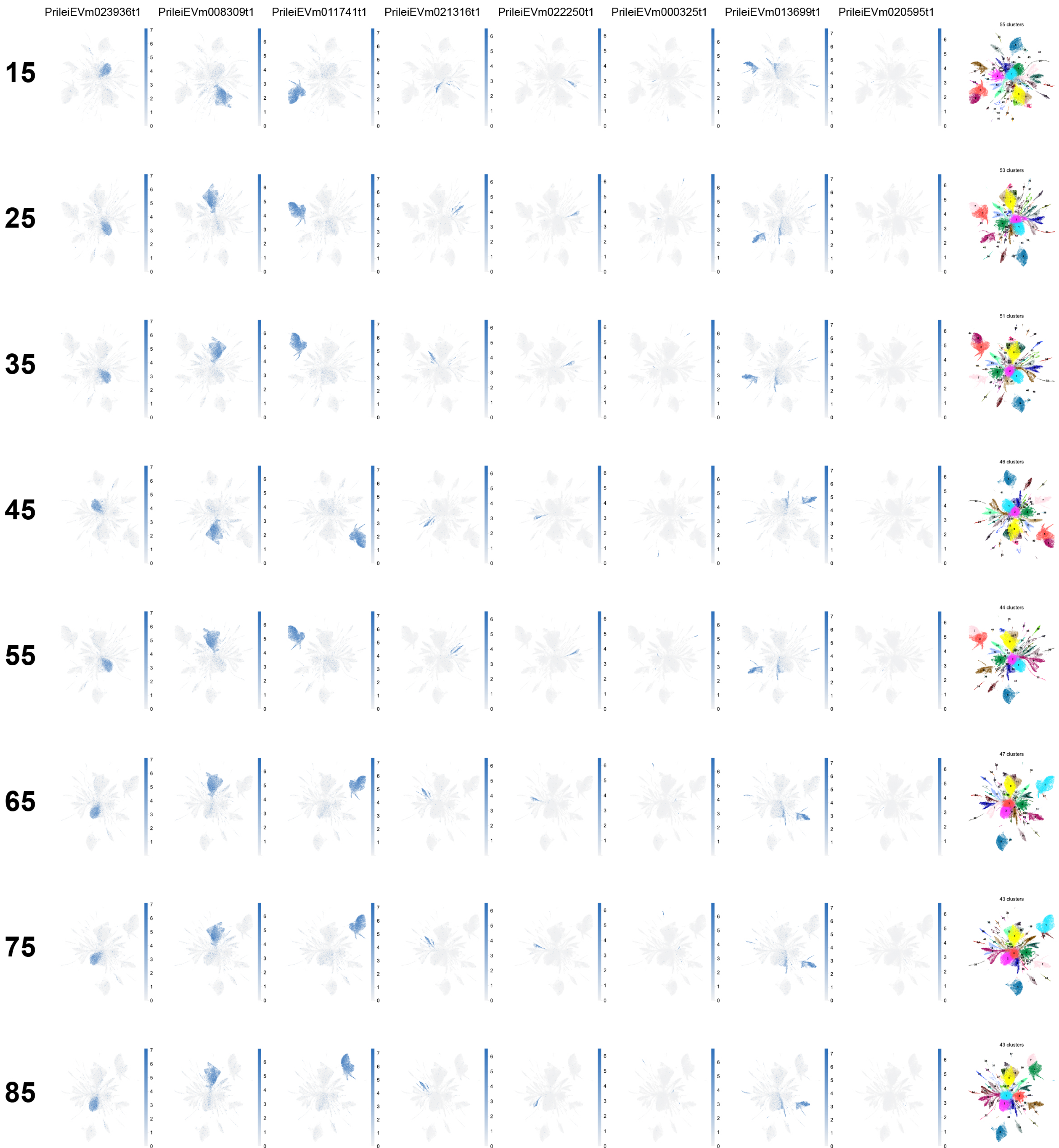

**number of principal components**

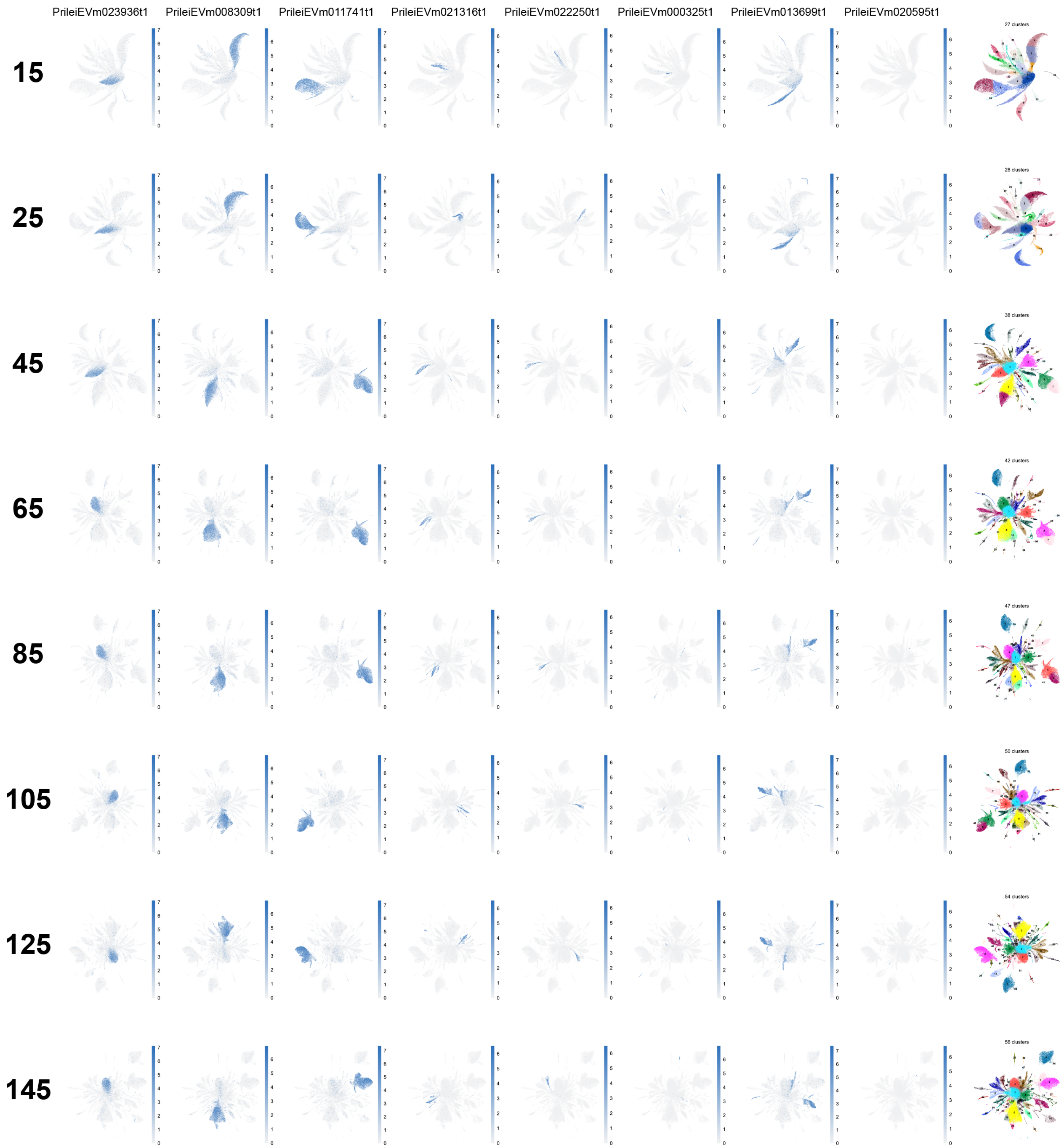
