## Supplementary material for "Annelid adult cell type diversity and their pluripotent cellular origins": Supp File 5.pdf

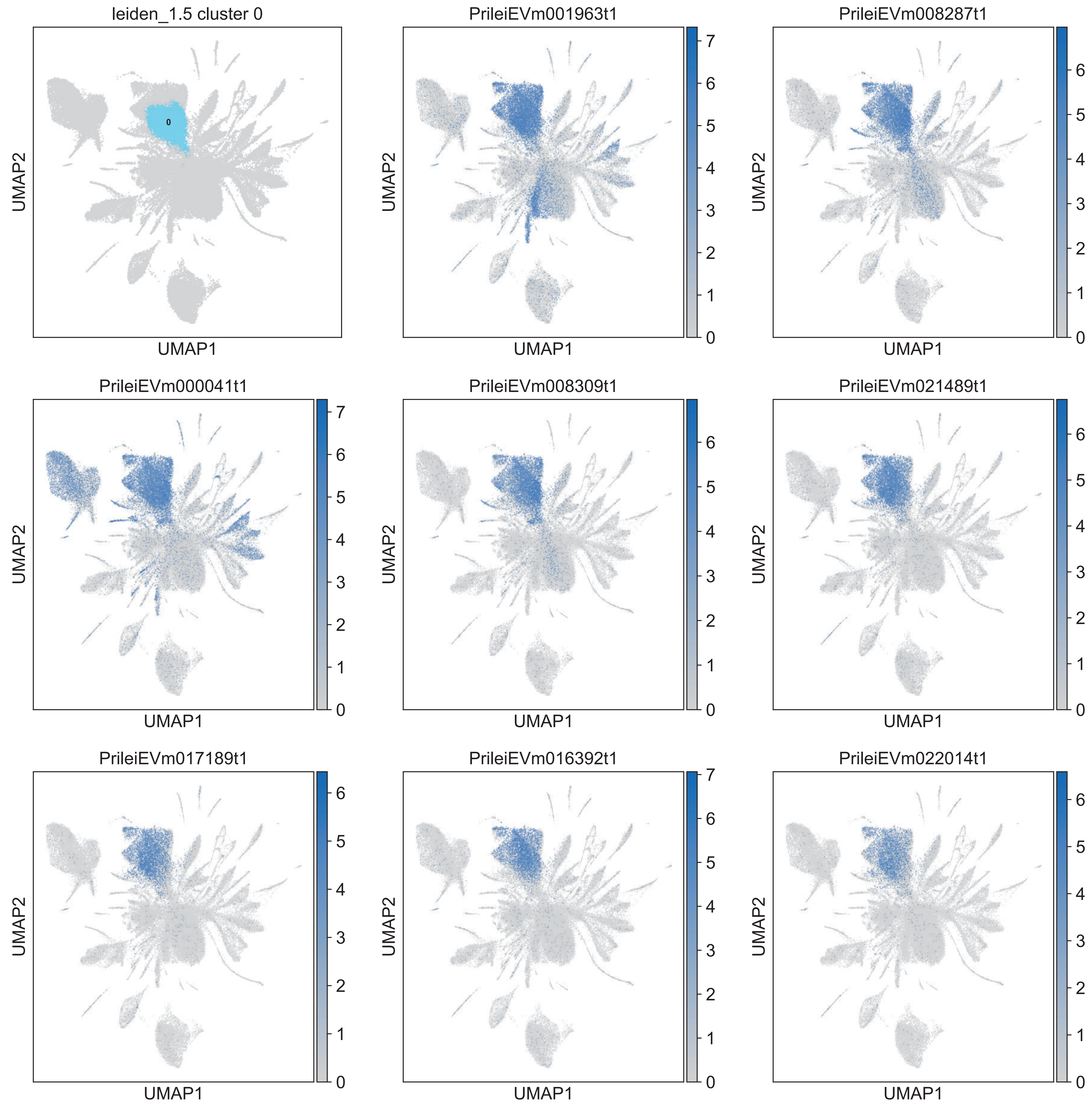

leiden_1.5 cluster 0
0
PrileiEVm001963t1
PrileiEVm008287t1
PrileiEVm000041t1
PrileiEVm008309t1
PrileiEVm021489t1
PrileiEVm017189t1
PrileiEVm016392t1
PrileiEVm022014t1
UMAP1
UMAP2
0
1
2
3
4
5
6
7

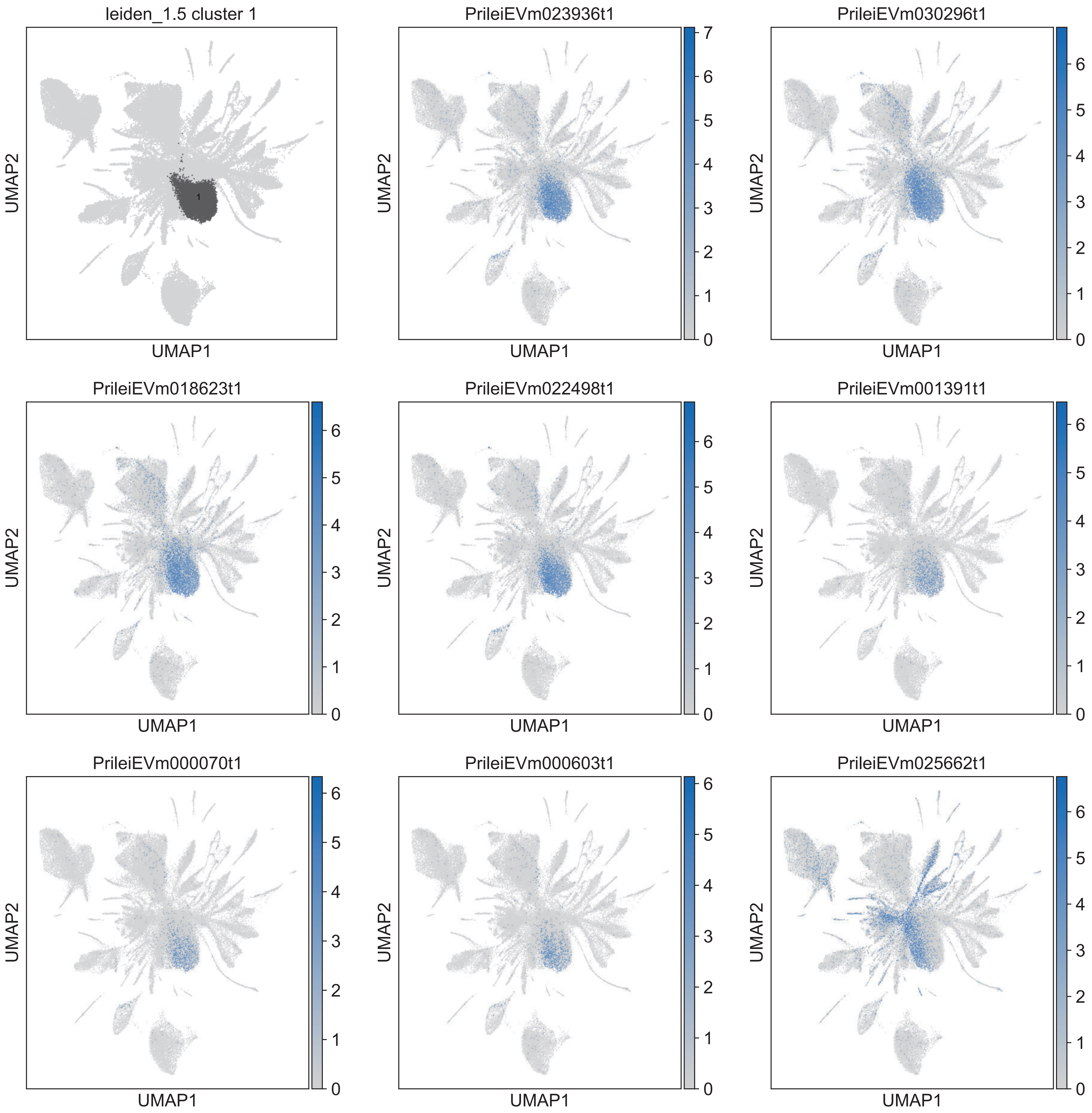

leiden_1.5 cluster 1
PrileiEVm023936t1
PrileiEVm030296t1
PrileiEVm018623t1
PrileiEVm022498t1
PrileiEVm001391t1
PrileiEVm000070t1
PrileiEVm000603t1
PrileiEVm025662t1
UMAP1
UMAP2

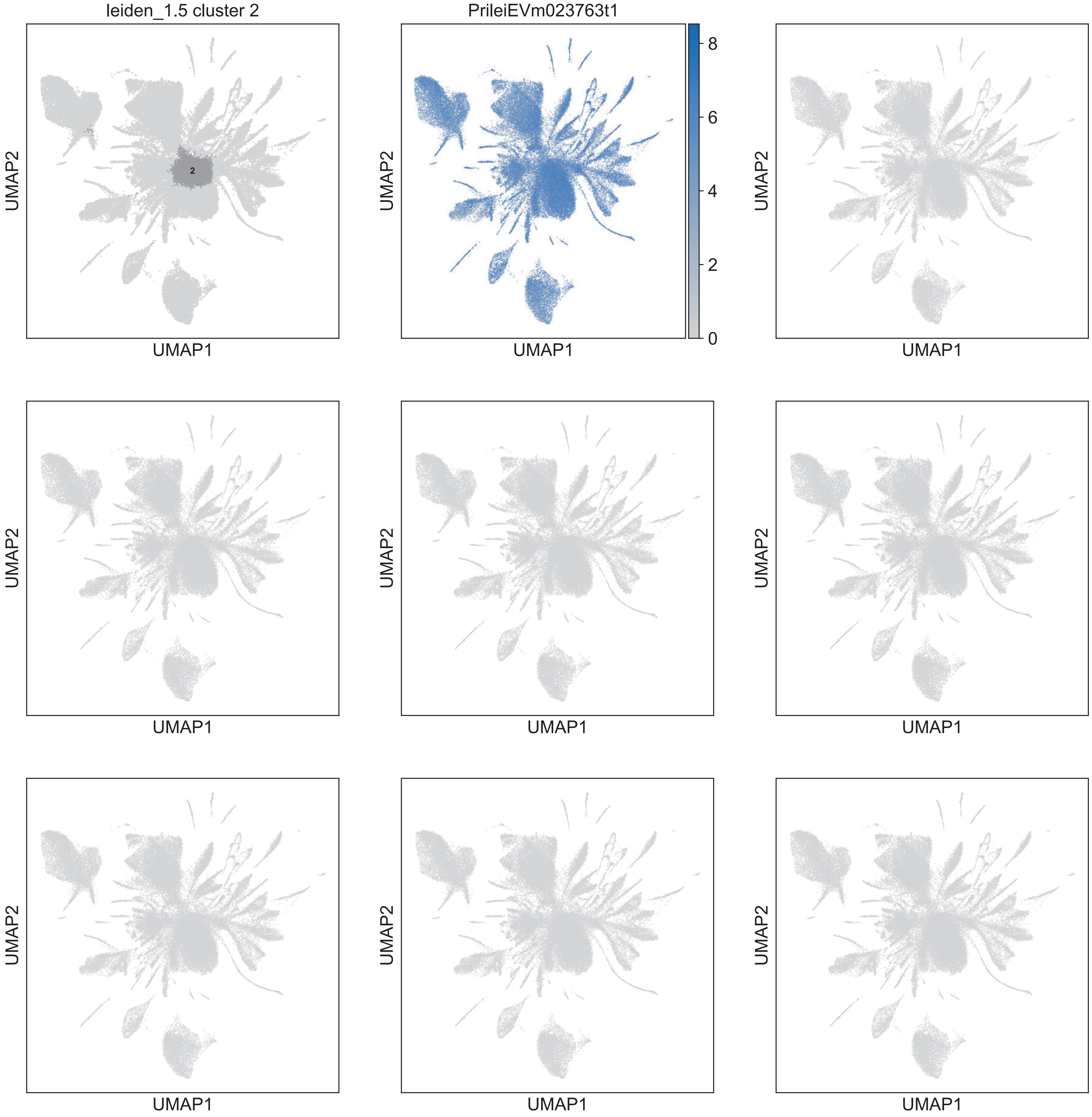
leiden_1.5 cluster 2
2
PrileiEVm023763t1
8
6
4
2
0
UMAP1
UMAP2

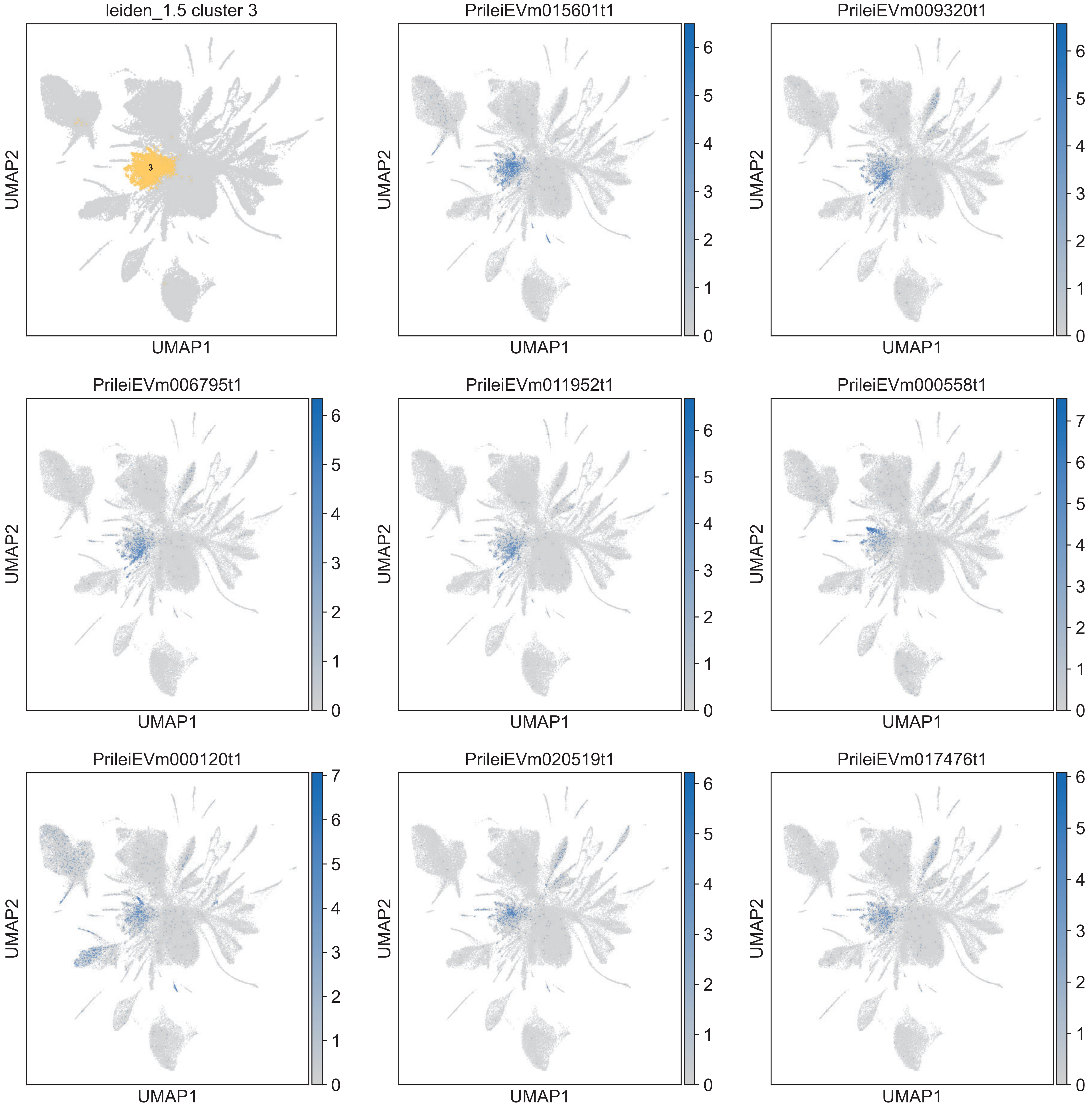
leiden_1.5 cluster 3
3
PrileiEVm015601t1
PrileiEVm009320t1
PrileiEVm006795t1
PrileiEVm011952t1
PrileiEVm000558t1
PrileiEVm000120t1
PrileiEVm020519t1
PrileiEVm017476t1
UMAP1
UMAP2

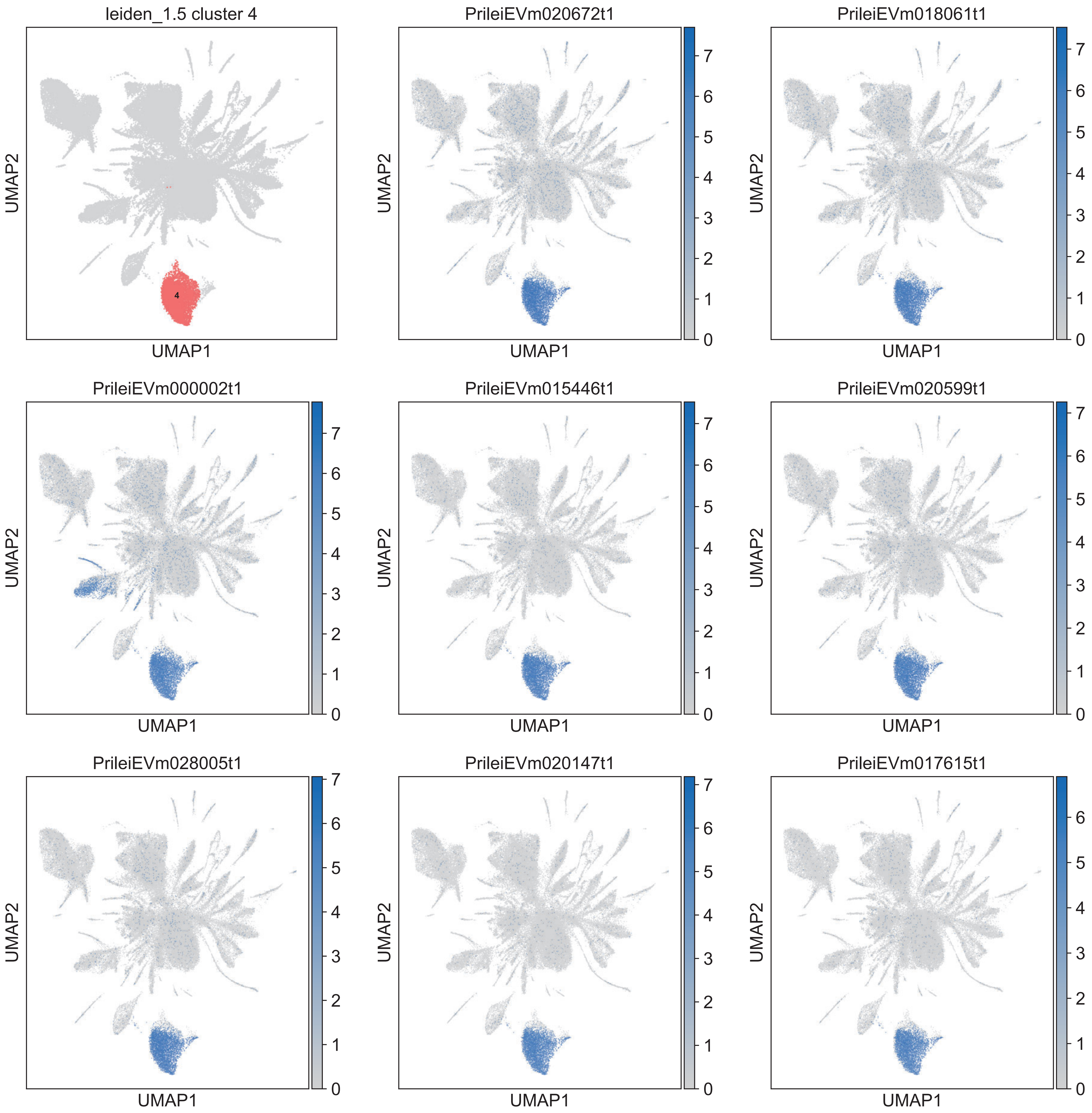
leiden_1.5 cluster 4
PrileiEVm020672t1
PrileiEVm018061t1
PrileiEVm000002t1
PrileiEVm015446t1
PrileiEVm020599t1
PrileiEVm028005t1
PrileiEVm020147t1
PrileiEVm017615t1
UMAP1
UMAP2

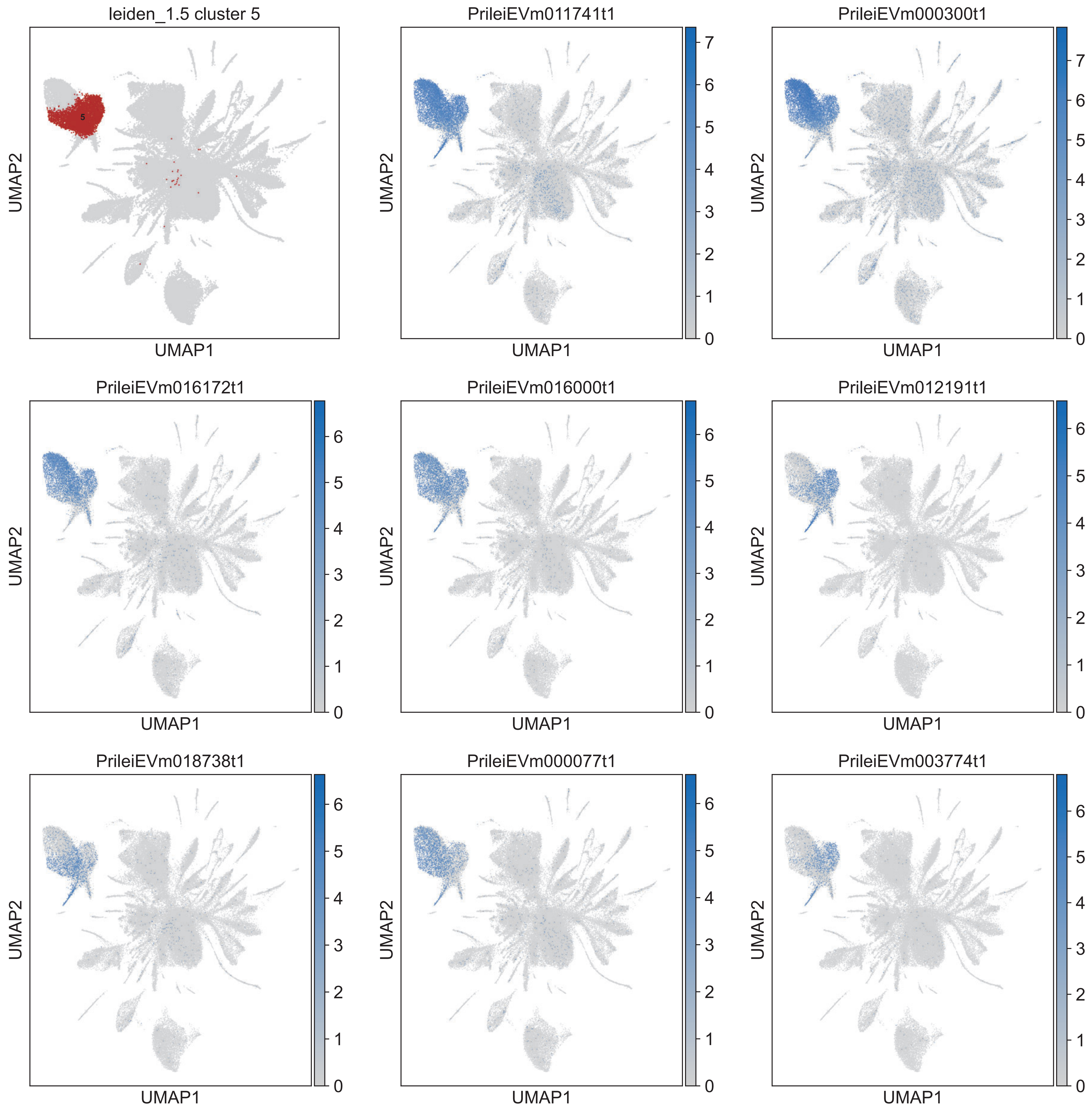
leiden_1.5 cluster 5
PrileiEVm011741t1
PrileiEVm000300t1
PrileiEVm016172t1
PrileiEVm016000t1
PrileiEVm012191t1
PrileiEVm018738t1
PrileiEVm000077t1
PrileiEVm003774t1
UMAP1
UMAP2

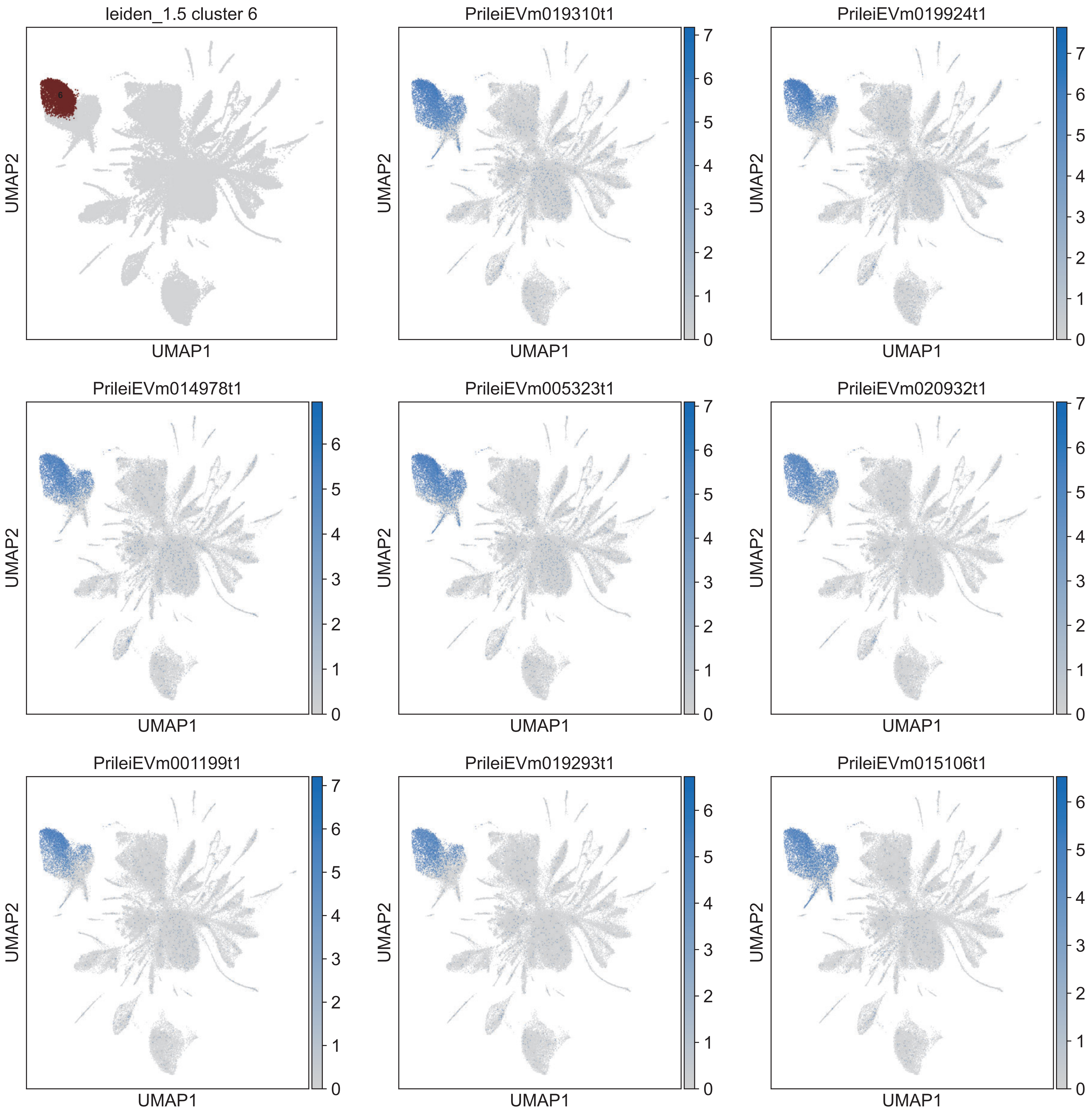
leiden_1.5 cluster 6
PrileiEVm019310t1
PrileiEVm019924t1
PrileiEVm014978t1
PrileiEVm005323t1
PrileiEVm020932t1
PrileiEVm001199t1
PrileiEVm019293t1
PrileiEVm015106t1
UMAP1
UMAP2

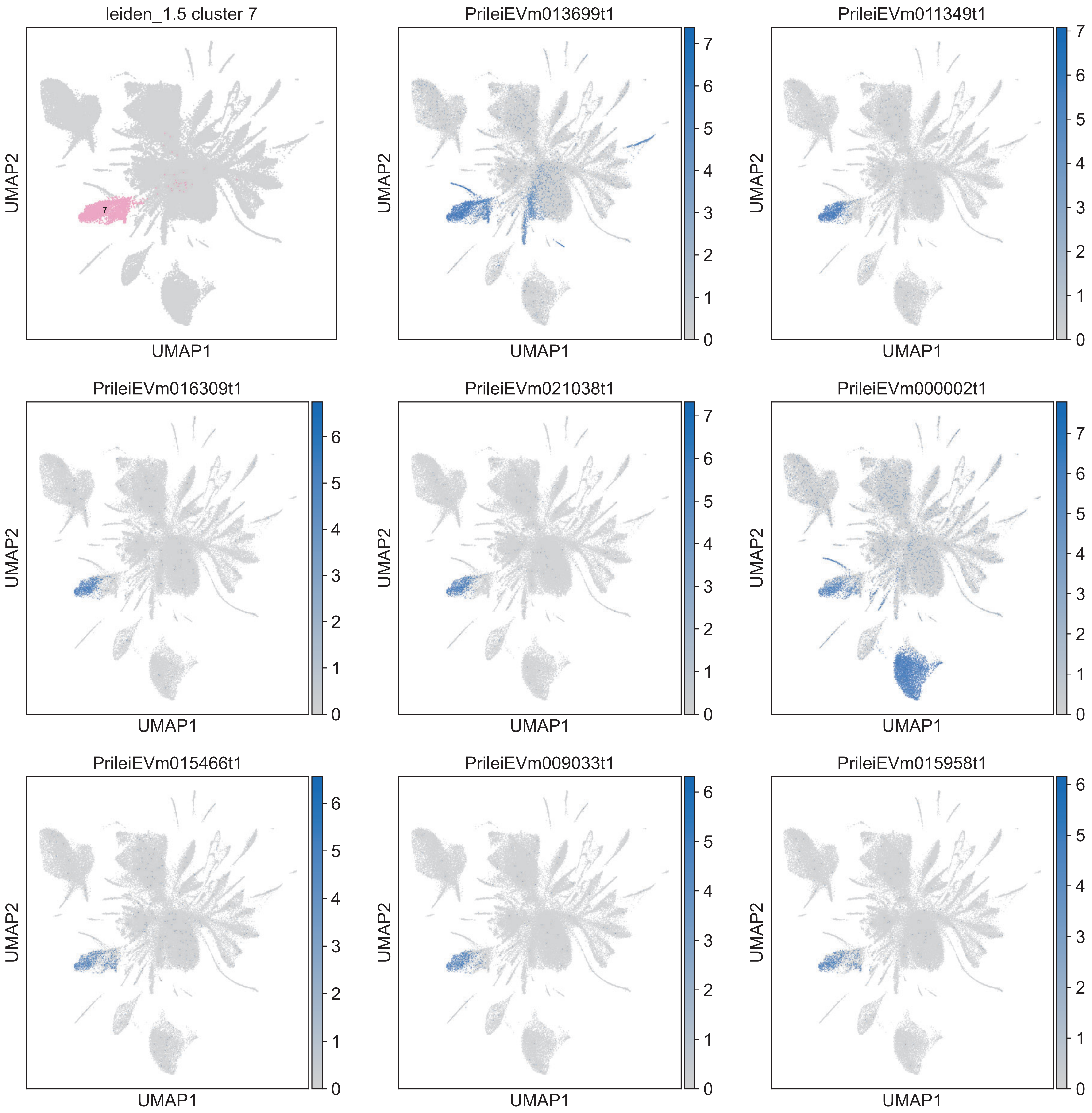
leiden_1.5 cluster 7
7
PrileiEVm013699t1
PrileiEVm011349t1
PrileiEVm016309t1
PrileiEVm021038t1
PrileiEVm000002t1
PrileiEVm015466t1
PrileiEVm009033t1
PrileiEVm015958t1
UMAP1
UMAP2

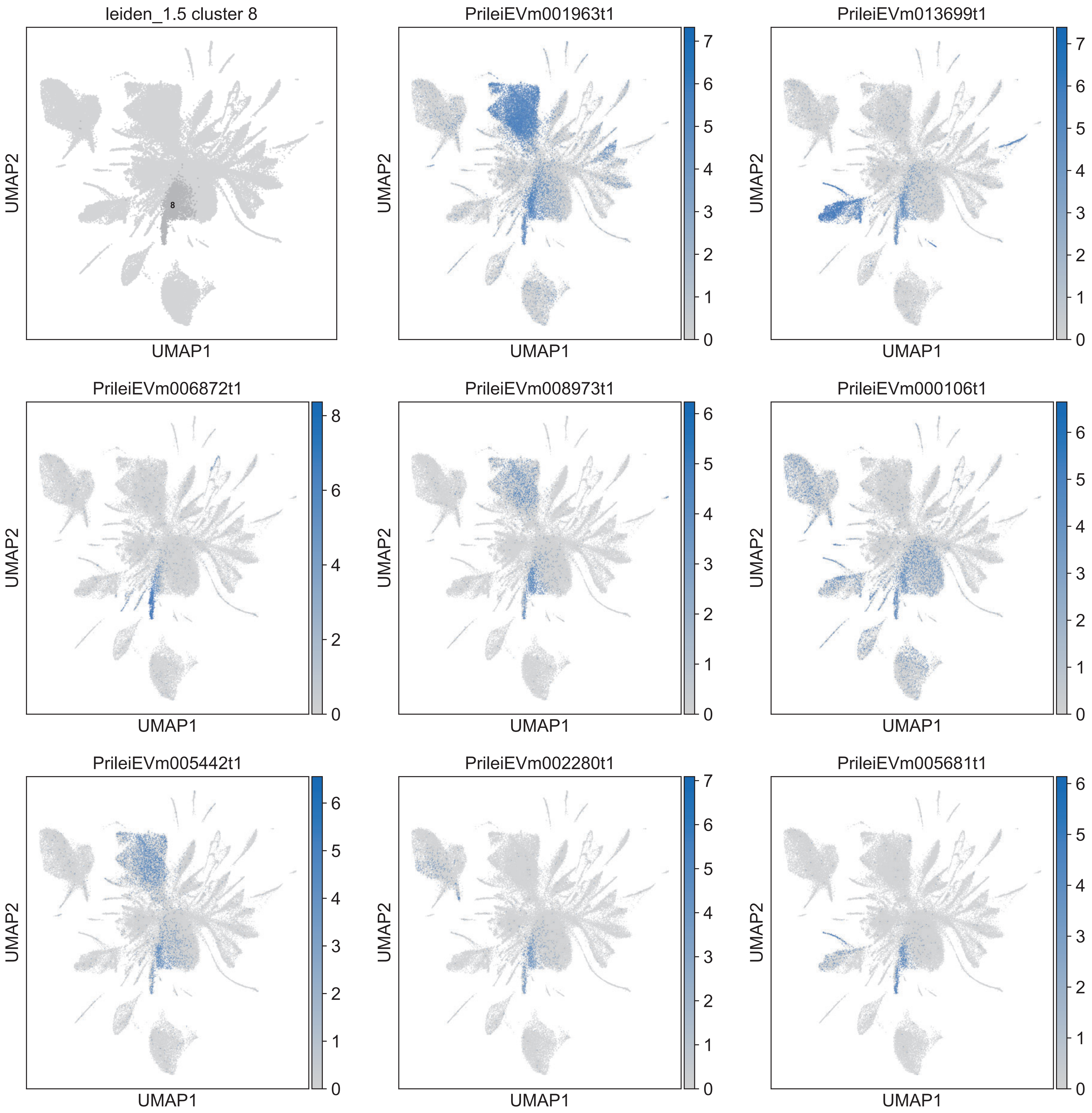
leiden_1.5 cluster 8
PrileiEVm001963t1
PrileiEVm013699t1
PrileiEVm006872t1
PrileiEVm008973t1
PrileiEVm000106t1
PrileiEVm005442t1
PrileiEVm002280t1
PrileiEVm005681t1
UMAP1
UMAP2

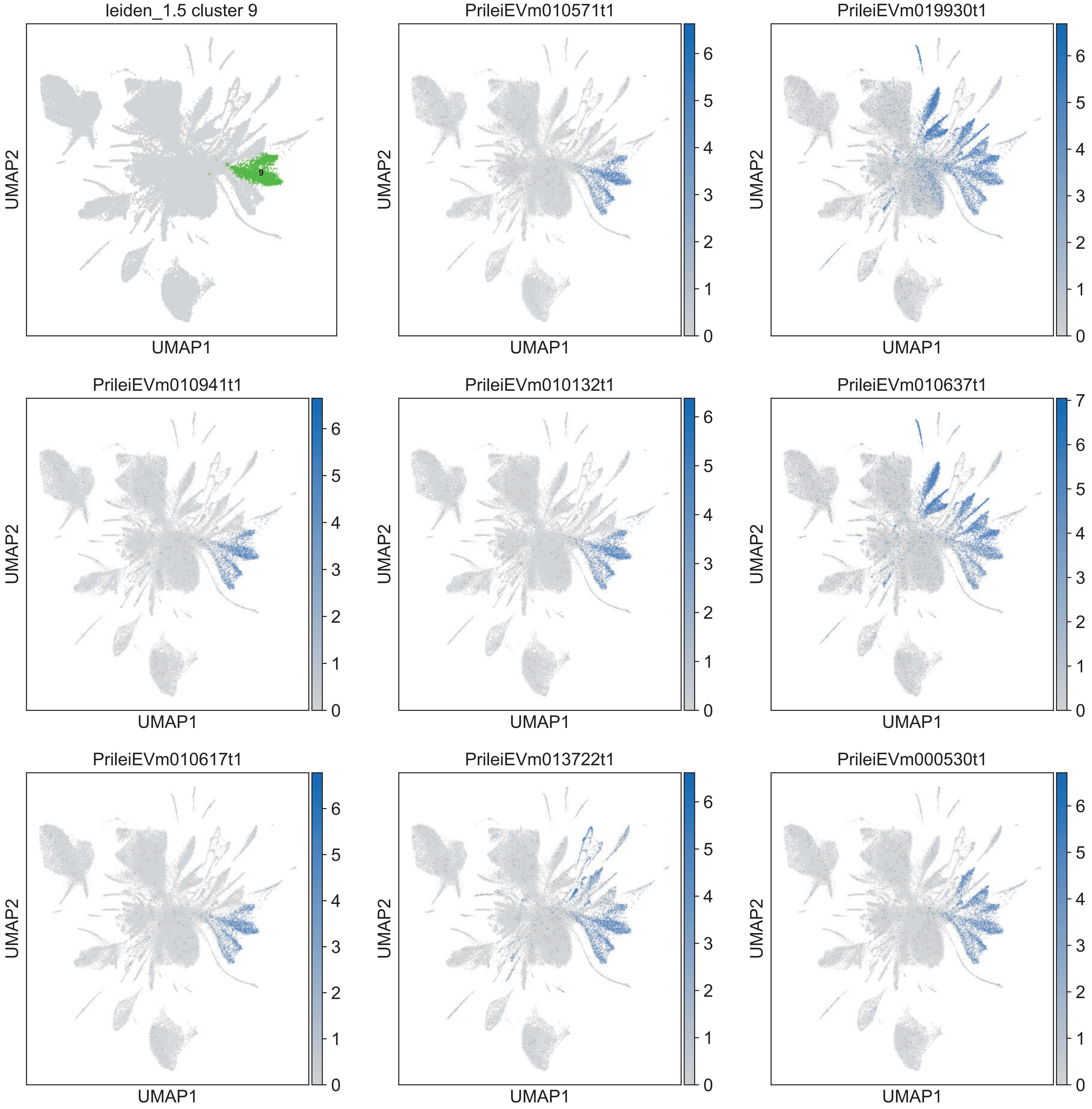
leiden_1.5 cluster 9
9
PrileiEVm010571t1
PrileiEVm019930t1
PrileiEVm010941t1
PrileiEVm010132t1
PrileiEVm010637t1
PrileiEVm010617t1
PrileiEVm013722t1
PrileiEVm000530t1
UMAP1
UMAP2

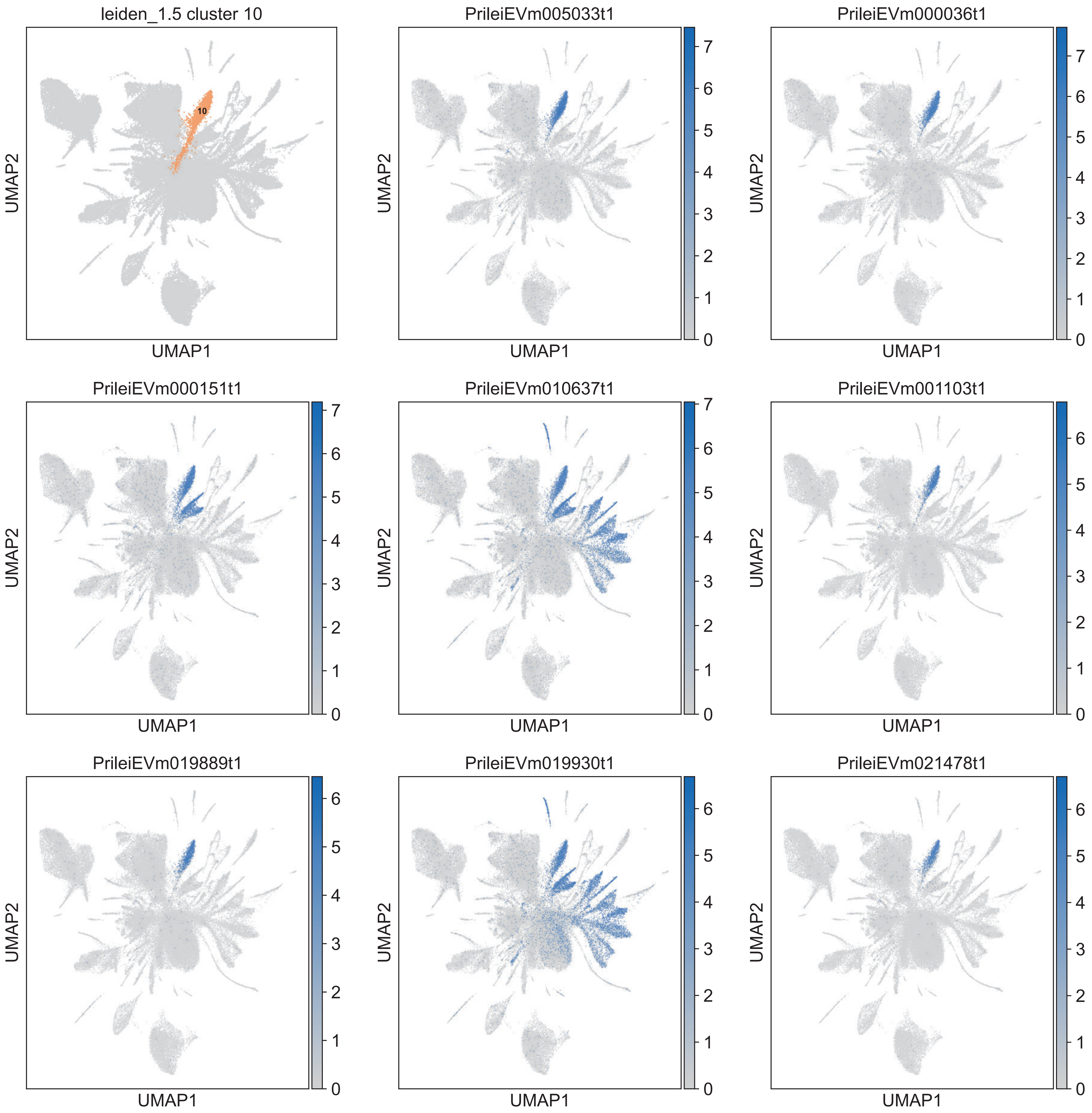
leiden_1.5 cluster 10
10
PrileiEVm005033t1
PrileiEVm000036t1
PrileiEVm000151t1
PrileiEVm010637t1
PrileiEVm001103t1
PrileiEVm019889t1
PrileiEVm019930t1
PrileiEVm021478t1
UMAP1
UMAP2

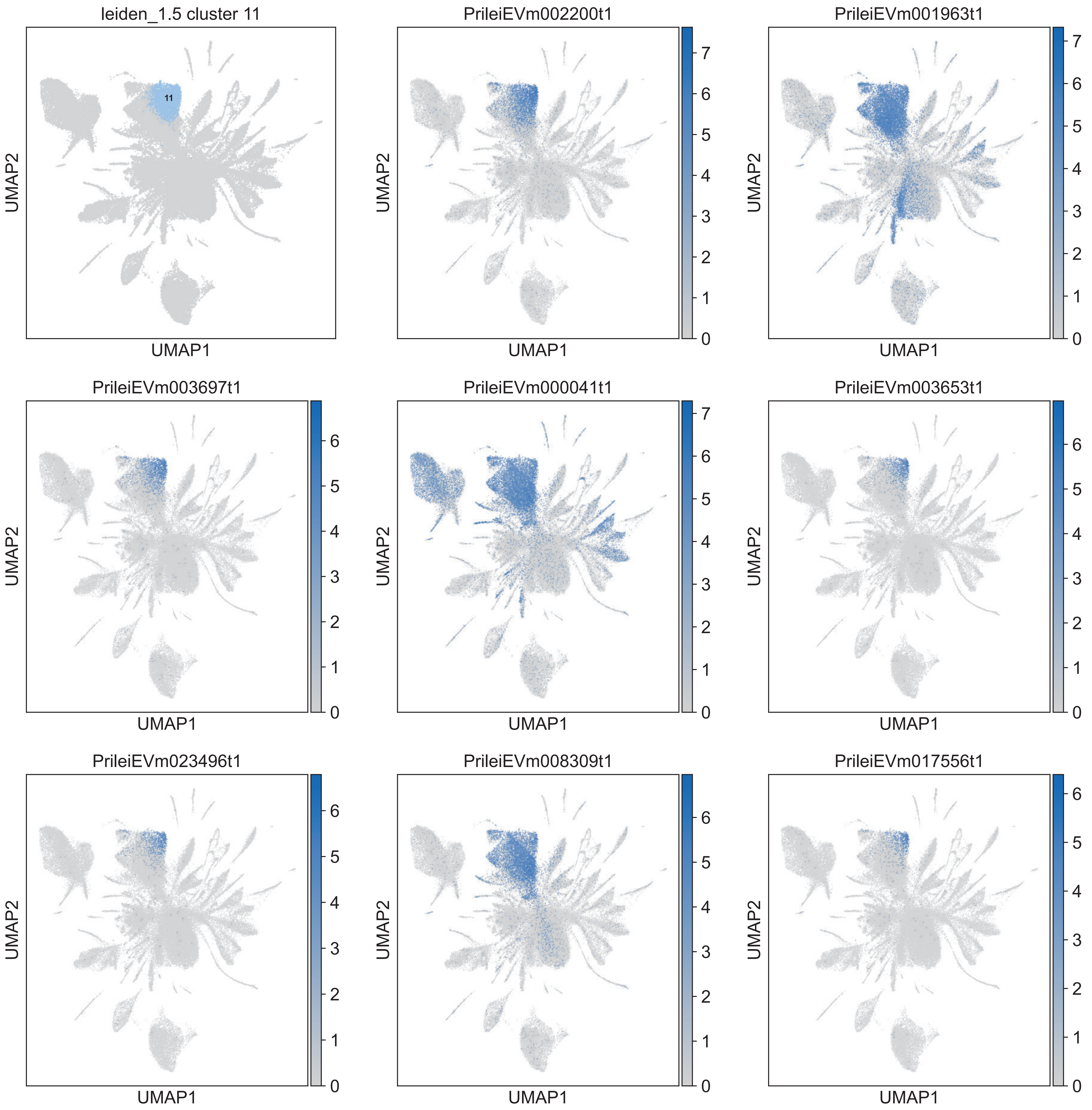
leiden_1.5 cluster 11
11
PrileiEVm002200t1
PrileiEVm001963t1
PrileiEVm003697t1
PrileiEVm000041t1
PrileiEVm003653t1
PrileiEVm023496t1
PrileiEVm008309t1
PrileiEVm017556t1
UMAP1
UMAP2

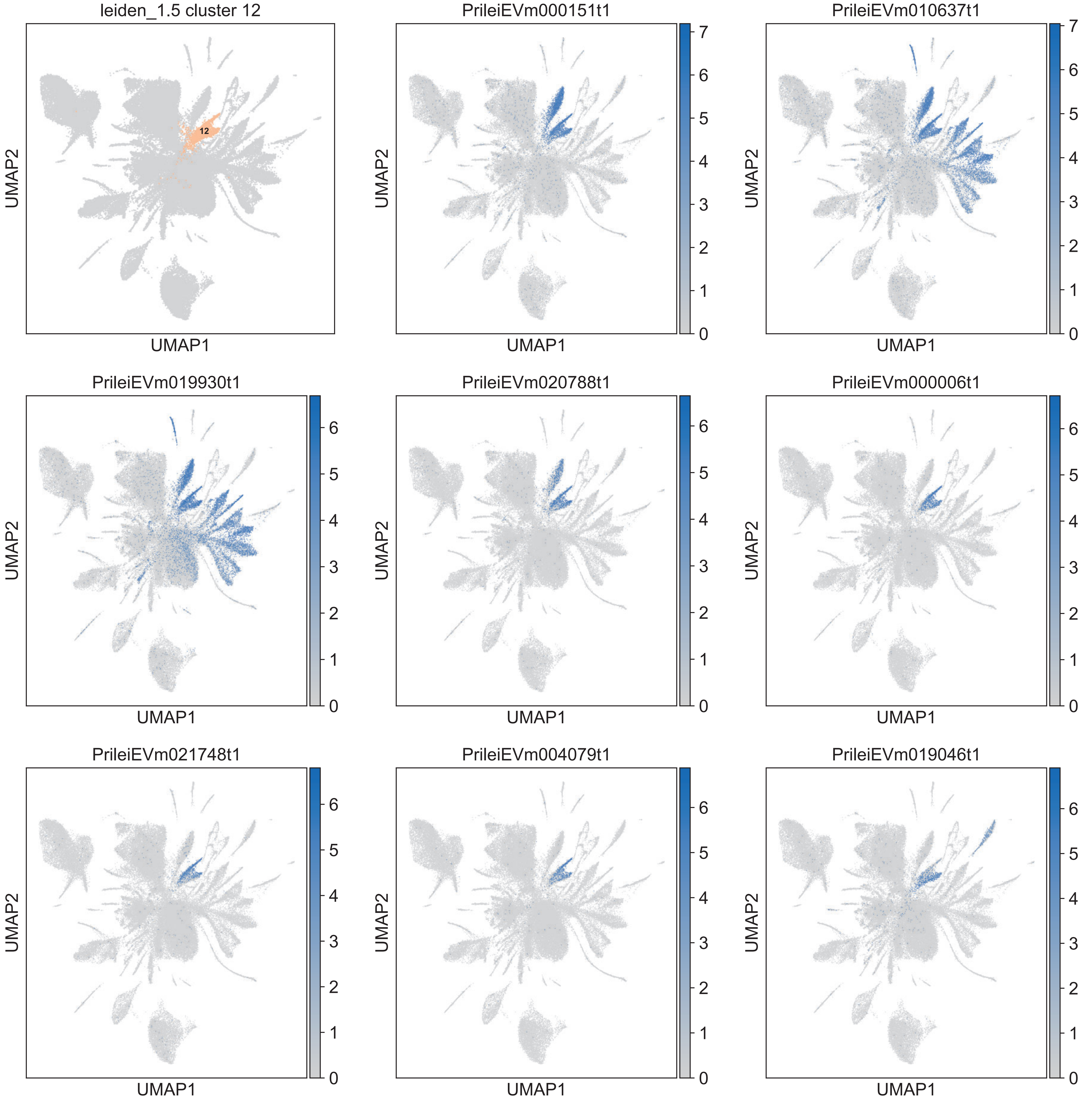
leiden_1.5 cluster 12
12
PrileiEVm000151t1
PrileiEVm010637t1
PrileiEVm019930t1
PrileiEVm020788t1
PrileiEVm000006t1
PrileiEVm021748t1
PrileiEVm004079t1
PrileiEVm019046t1
UMAP1
UMAP2

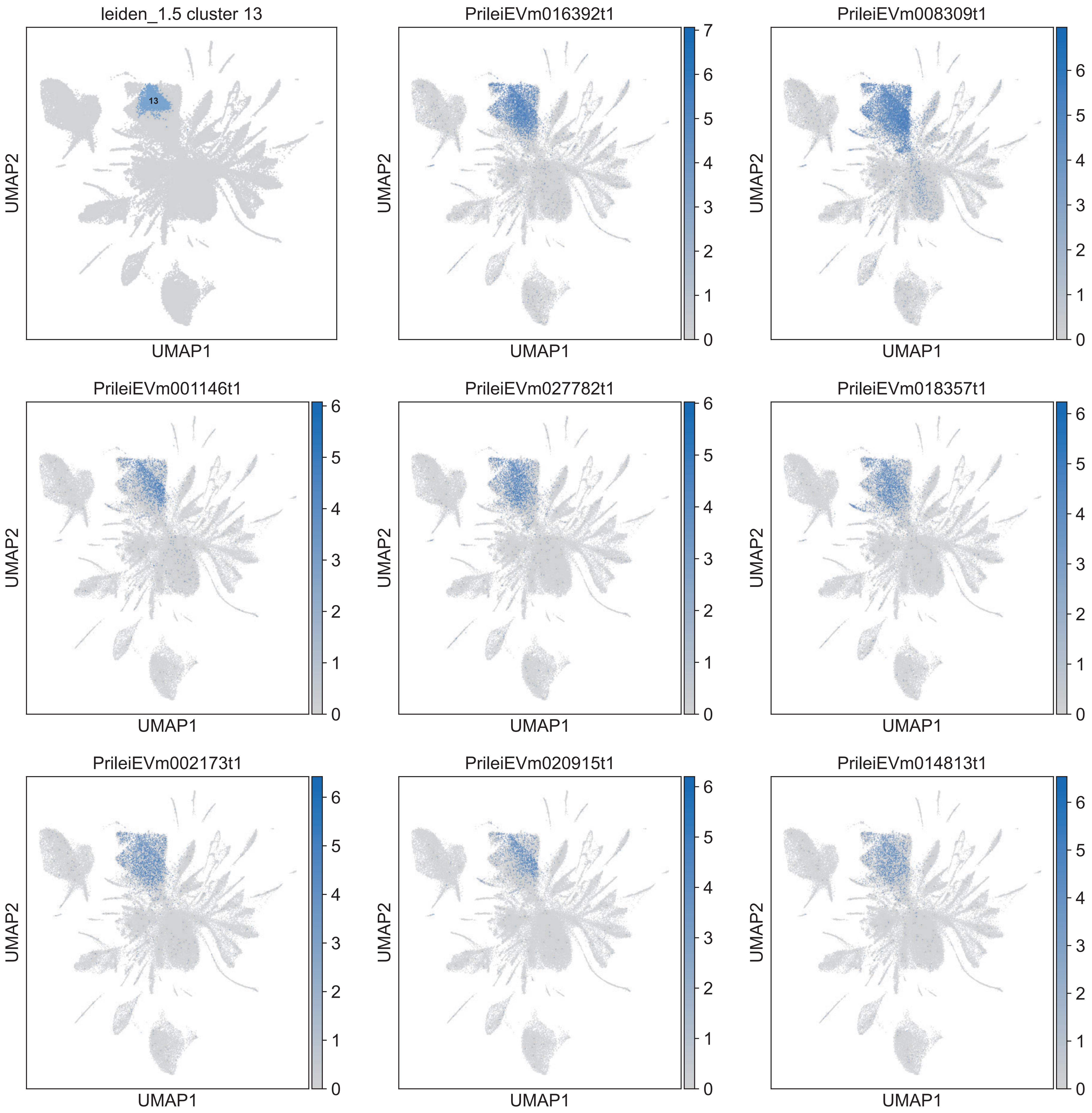
leiden_1.5 cluster 13
13
PrileiEVm016392t1
PrileiEVm008309t1
PrileiEVm001146t1
PrileiEVm027782t1
PrileiEVm018357t1
PrileiEVm002173t1
PrileiEVm020915t1
PrileiEVm014813t1
UMAP1
UMAP2

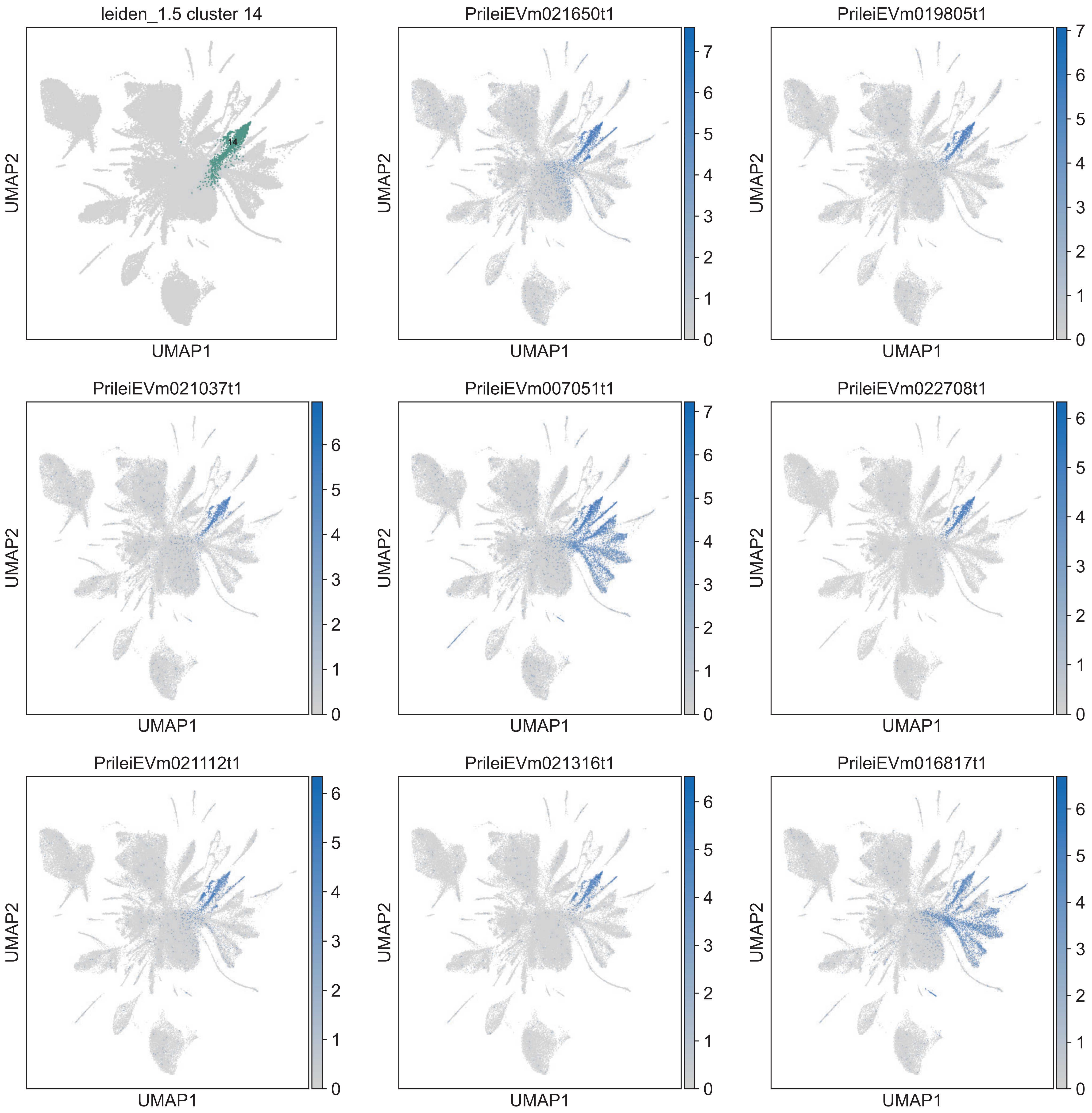
leiden_1.5 cluster 14
14
PrileiEVm021650t1
PrileiEVm019805t1
PrileiEVm021037t1
PrileiEVm007051t1
PrileiEVm022708t1
PrileiEVm021112t1
PrileiEVm021316t1
PrileiEVm016817t1
UMAP1
UMAP2

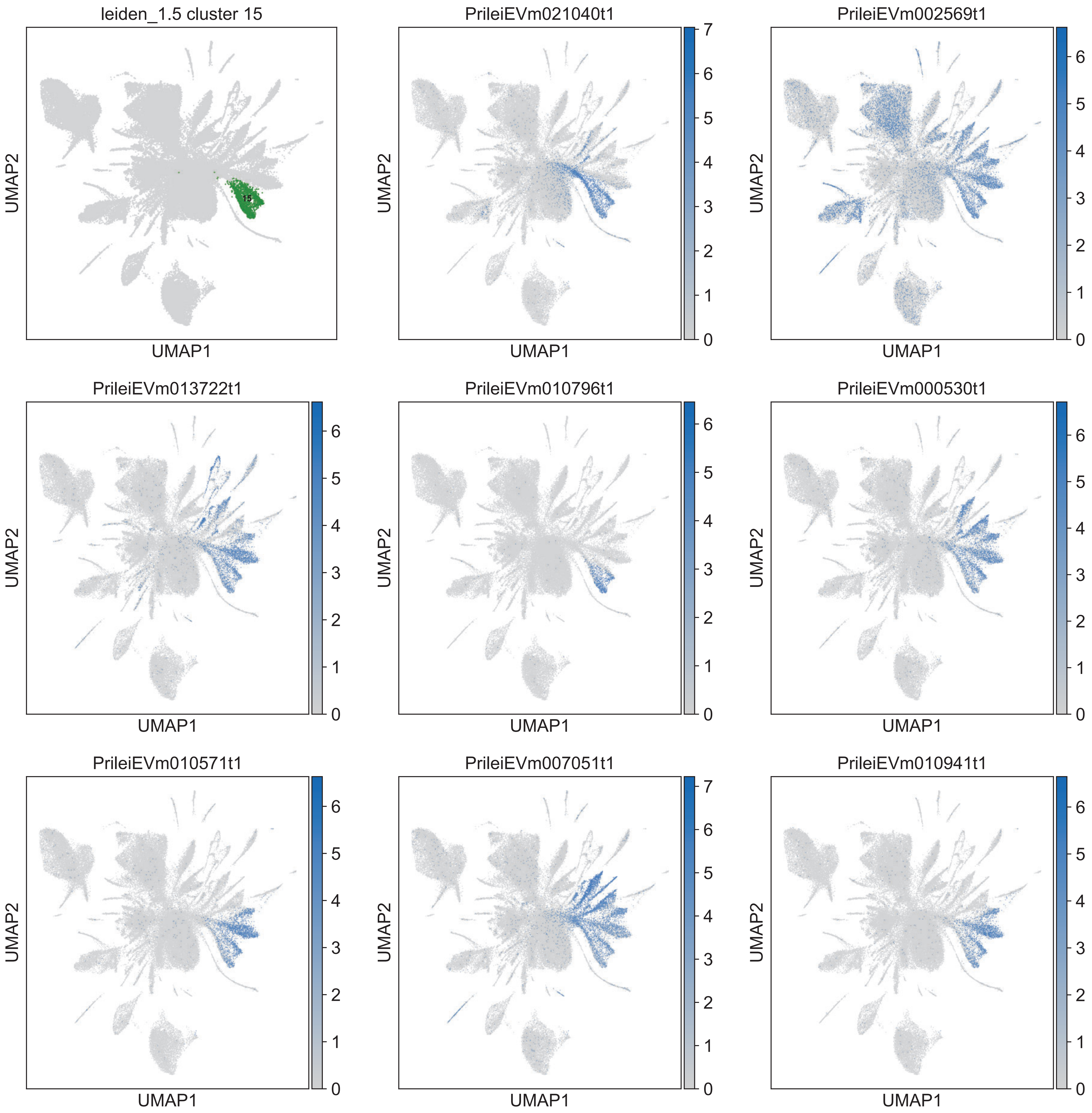
leiden_1.5 cluster 15
15
PrileiEVm021040t1
PrileiEVm002569t1
PrileiEVm013722t1
PrileiEVm010796t1
PrileiEVm000530t1
PrileiEVm010571t1
PrileiEVm007051t1
PrileiEVm010941t1
UMAP1
UMAP2

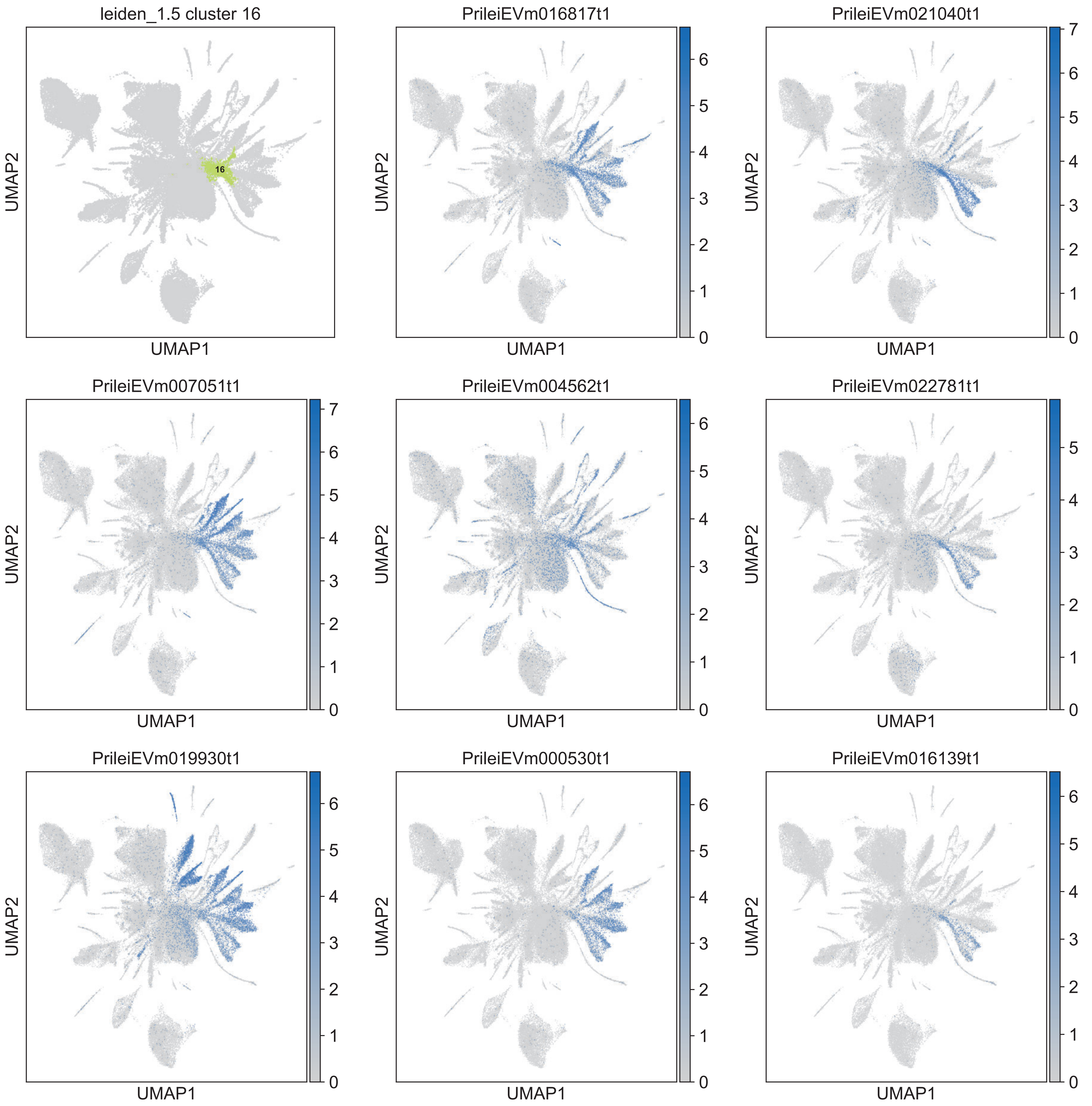

leiden_1.5 cluster 16
16
PrileiEVm016817t1
PrileiEVm021040t1
PrileiEVm007051t1
PrileiEVm004562t1
PrileiEVm022781t1
PrileiEVm019930t1
PrileiEVm000530t1
PrileiEVm016139t1
UMAP1
UMAP2

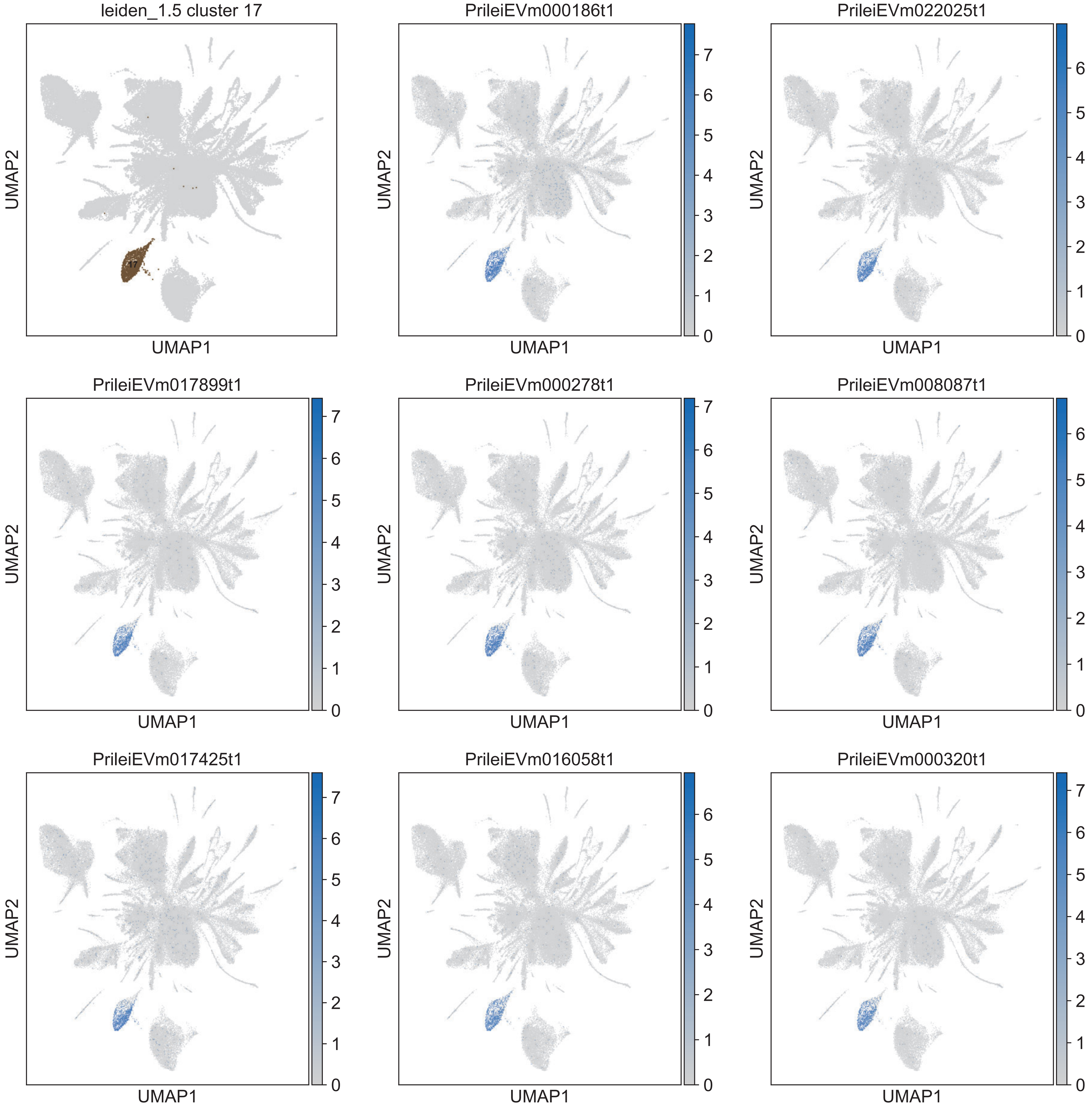

leiden_1.5 cluster 17
17
PrileiEVm000186t1
PrileiEVm022025t1
PrileiEVm017899t1
PrileiEVm000278t1
PrileiEVm008087t1
PrileiEVm017425t1
PrileiEVm016058t1
PrileiEVm000320t1
UMAP1
UMAP2

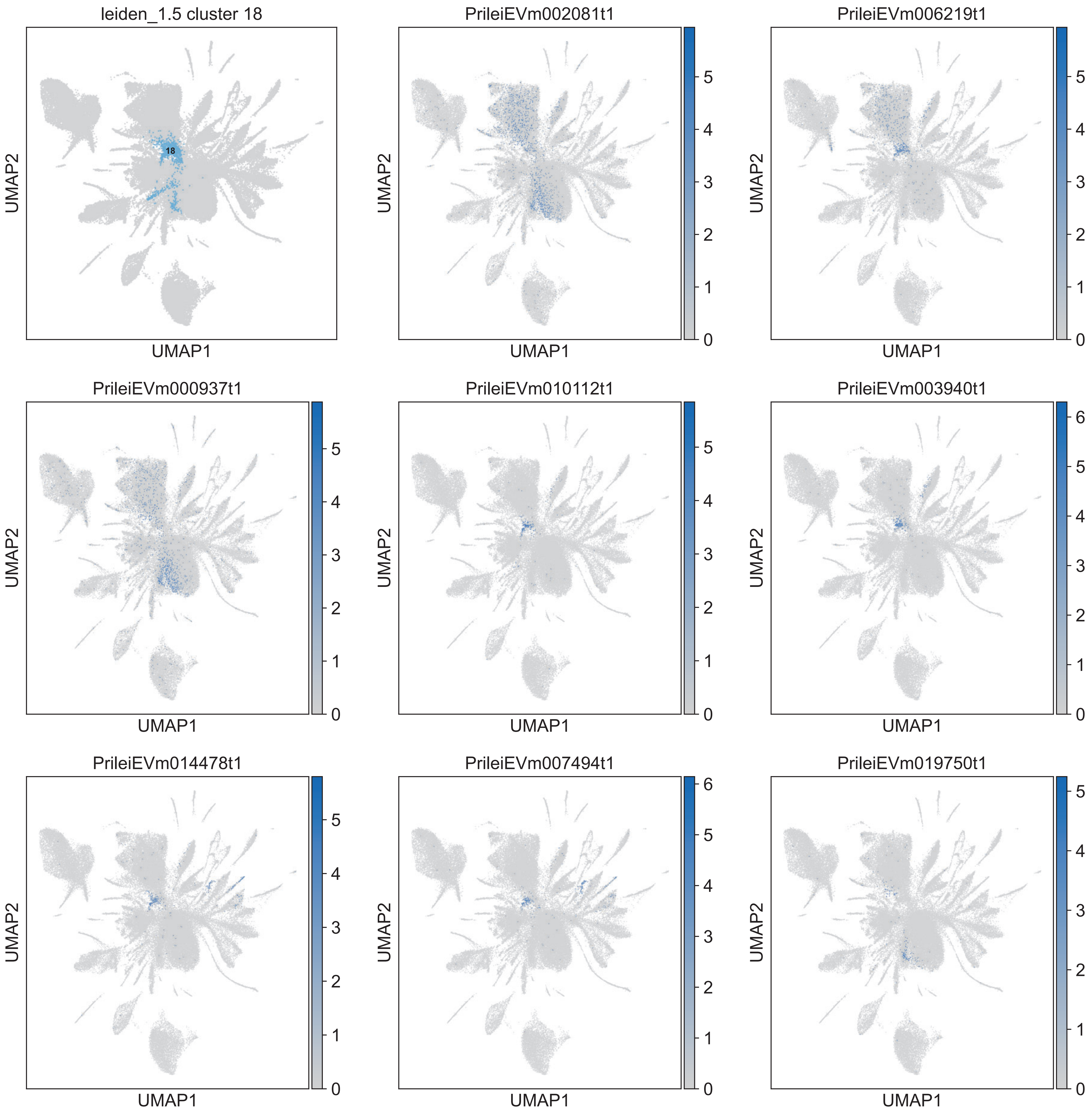

leiden_1.5 cluster 18
18
PrileiEVm002081t1
PrileiEVm006219t1
PrileiEVm000937t1
PrileiEVm010112t1
PrileiEVm003940t1
PrileiEVm014478t1
PrileiEVm007494t1
PrileiEVm019750t1
UMAP1
UMAP2

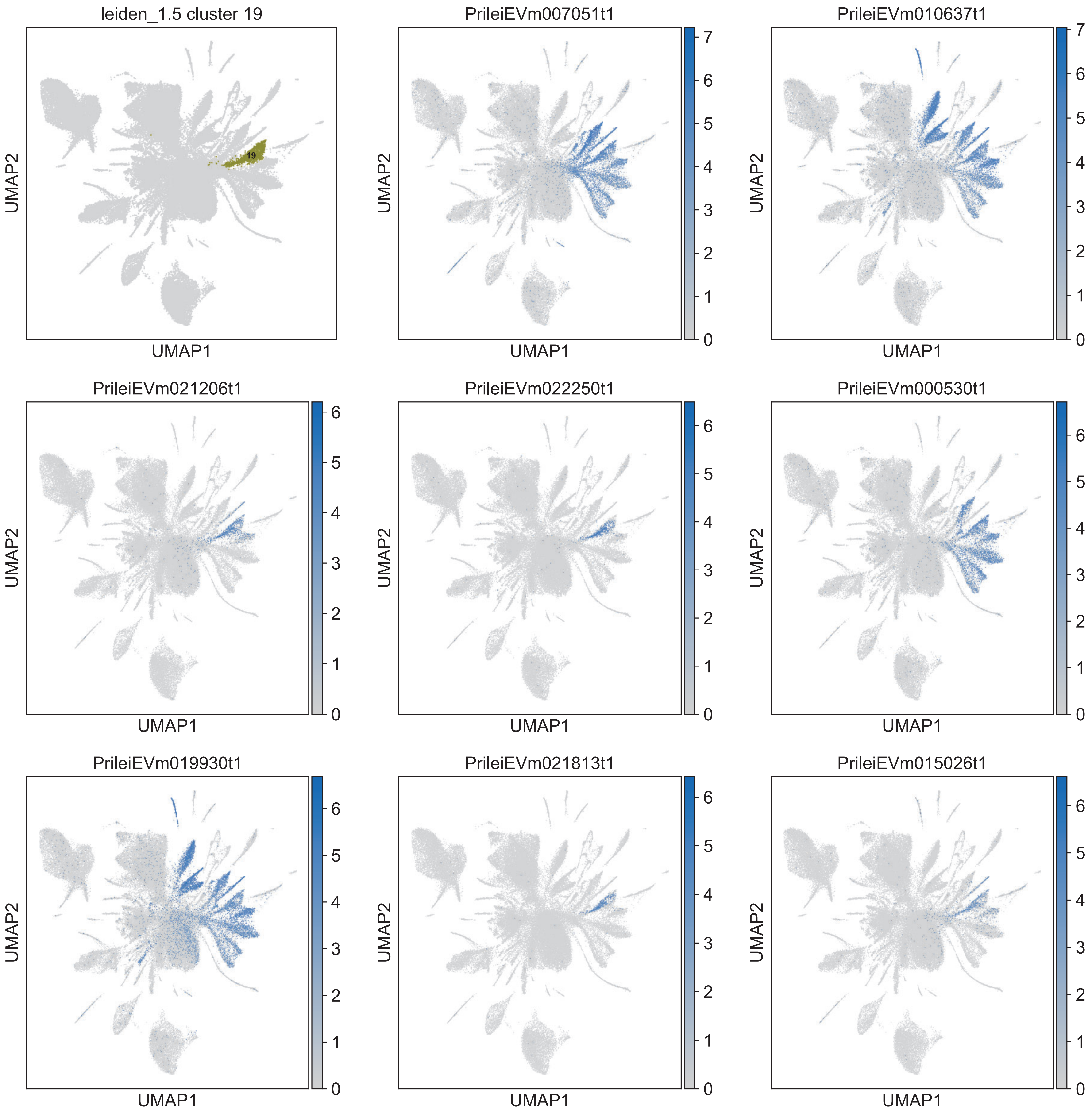

leiden_1.5 cluster 19
19
PrileiEVm007051t1
PrileiEVm010637t1
PrileiEVm021206t1
PrileiEVm022250t1
PrileiEVm000530t1
PrileiEVm019930t1
PrileiEVm021813t1
PrileiEVm015026t1
UMAP1
UMAP2

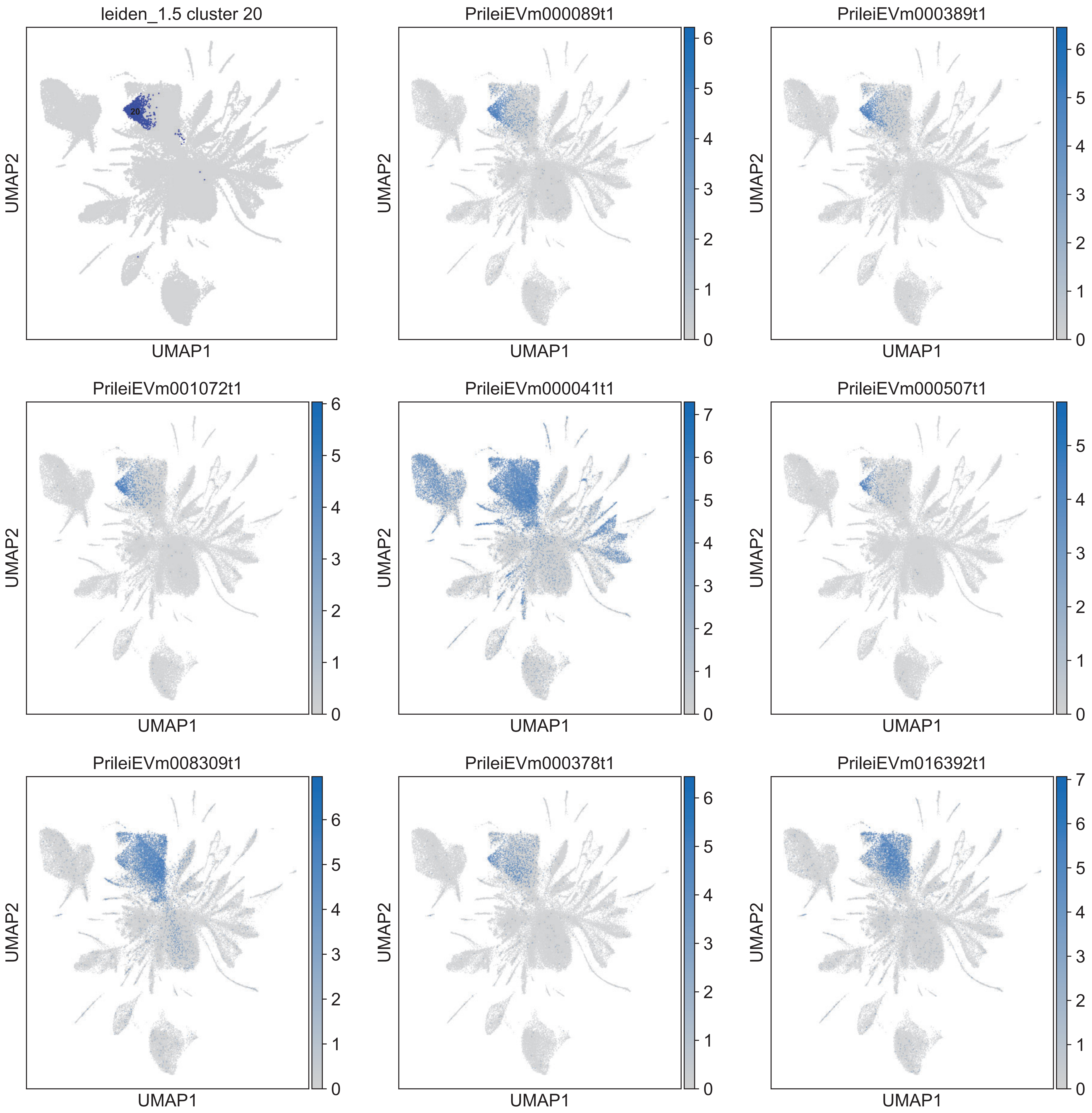

leiden_1.5 cluster 20
20
PrileiEVm000089t1
PrileiEVm000389t1
PrileiEVm001072t1
PrileiEVm000041t1
PrileiEVm000507t1
PrileiEVm008309t1
PrileiEVm000378t1
PrileiEVm016392t1
UMAP1
UMAP2

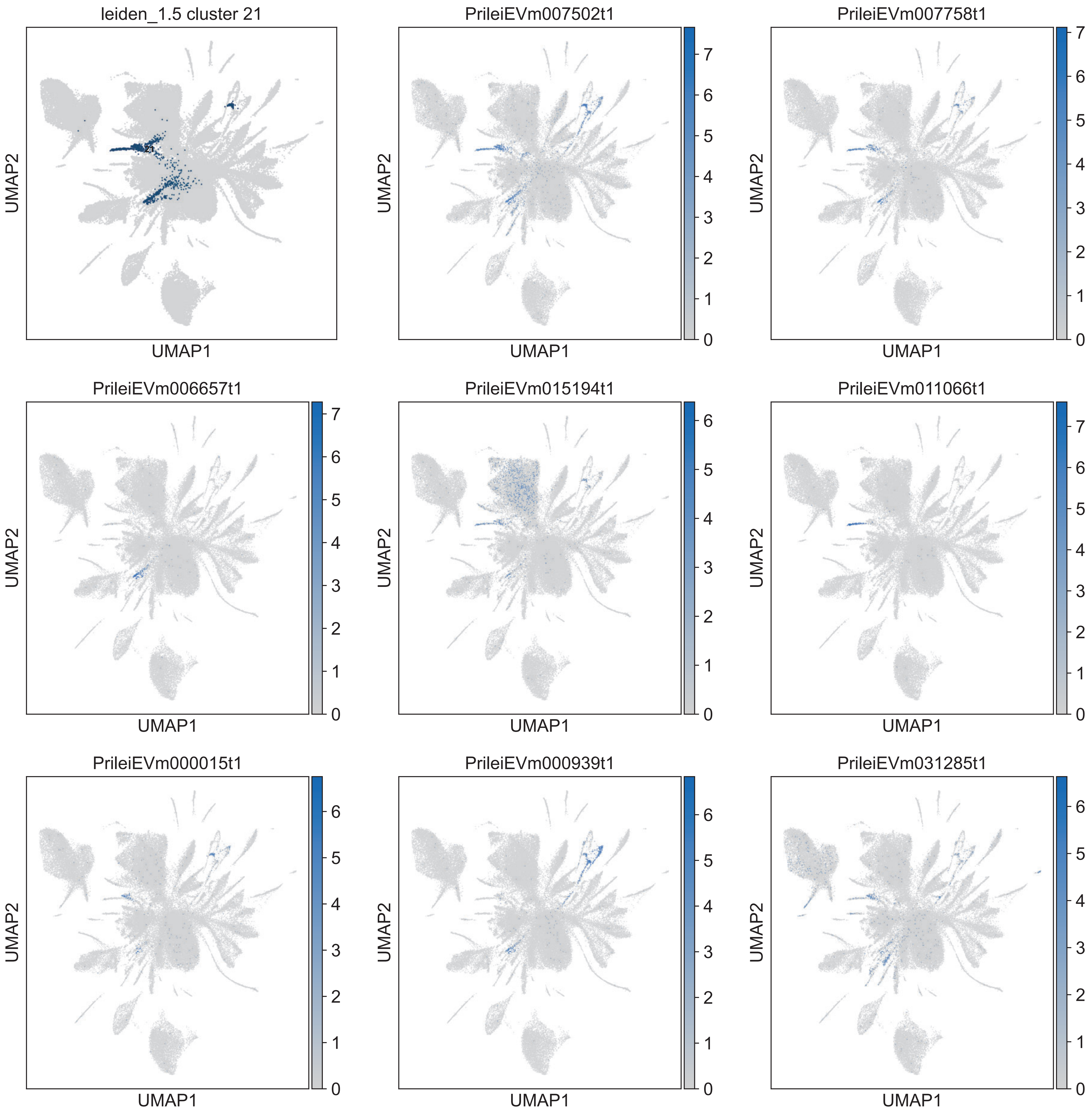
leiden_1.5 cluster 21
PrileiEVm007502t1
PrileiEVm007758t1
PrileiEVm006657t1
PrileiEVm015194t1
PrileiEVm011066t1
PrileiEVm000015t1
PrileiEVm000939t1
PrileiEVm031285t1
UMAP1
UMAP2

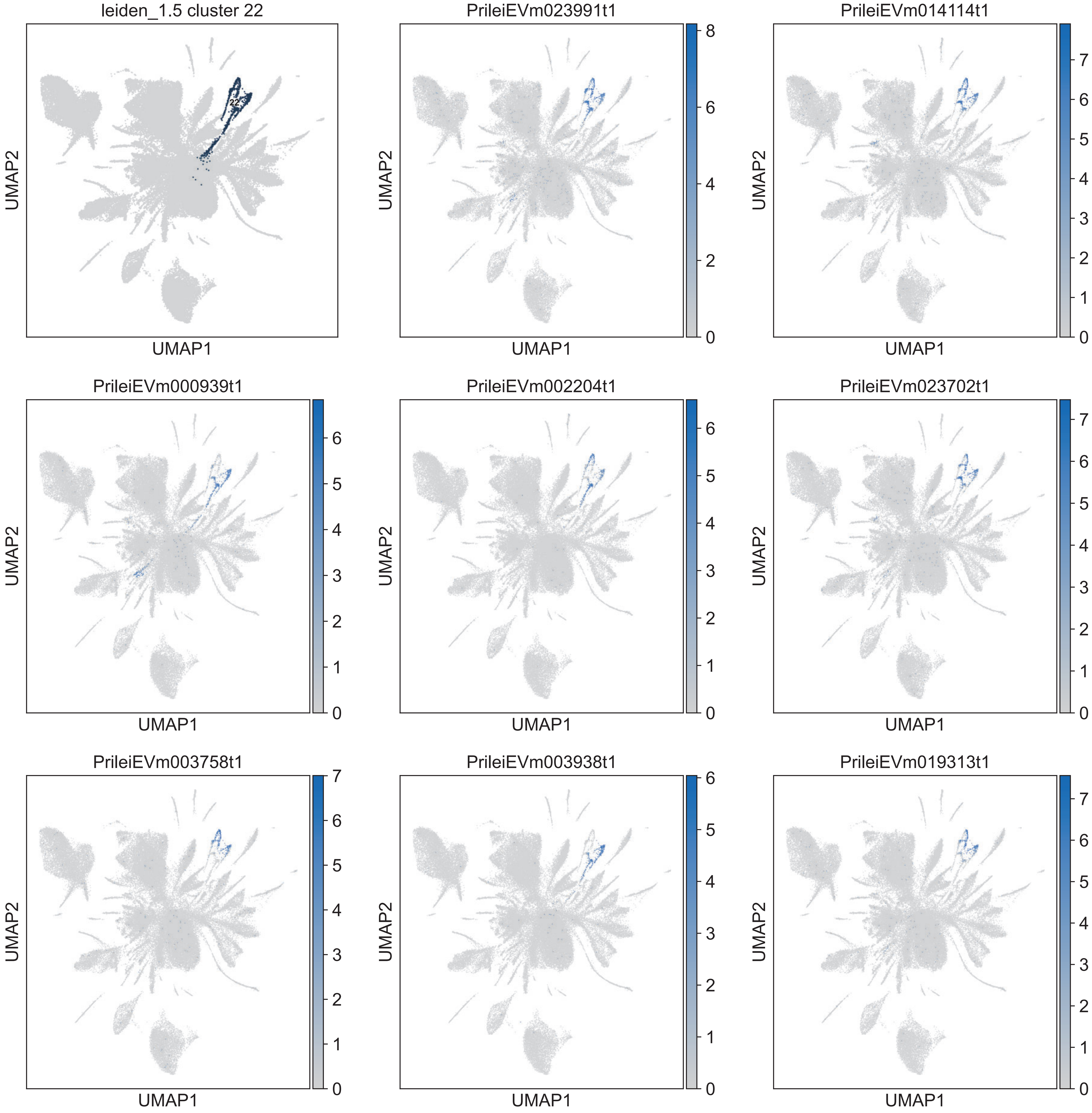

leiden_1.5 cluster 22
22
PrileiEVm023991t1
PrileiEVm014114t1
PrileiEVm000939t1
PrileiEVm002204t1
PrileiEVm023702t1
PrileiEVm003758t1
PrileiEVm003938t1
PrileiEVm019313t1
UMAP1
UMAP2

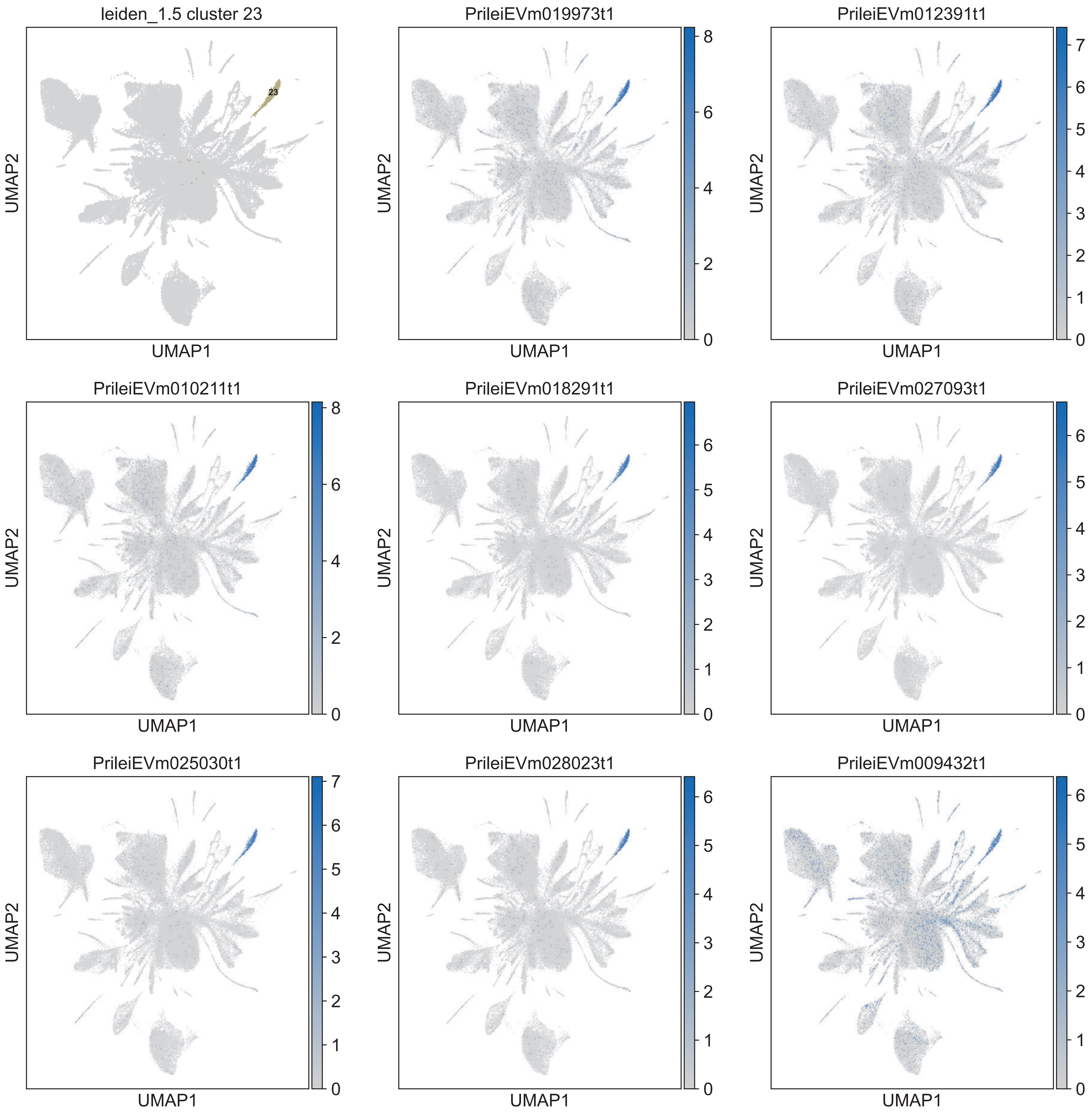
leiden_1.5 cluster 23
23
PrileiEVm019973t1
PrileiEVm012391t1
PrileiEVm010211t1
PrileiEVm018291t1
PrileiEVm027093t1
PrileiEVm025030t1
PrileiEVm028023t1
PrileiEVm009432t1
UMAP1
UMAP2

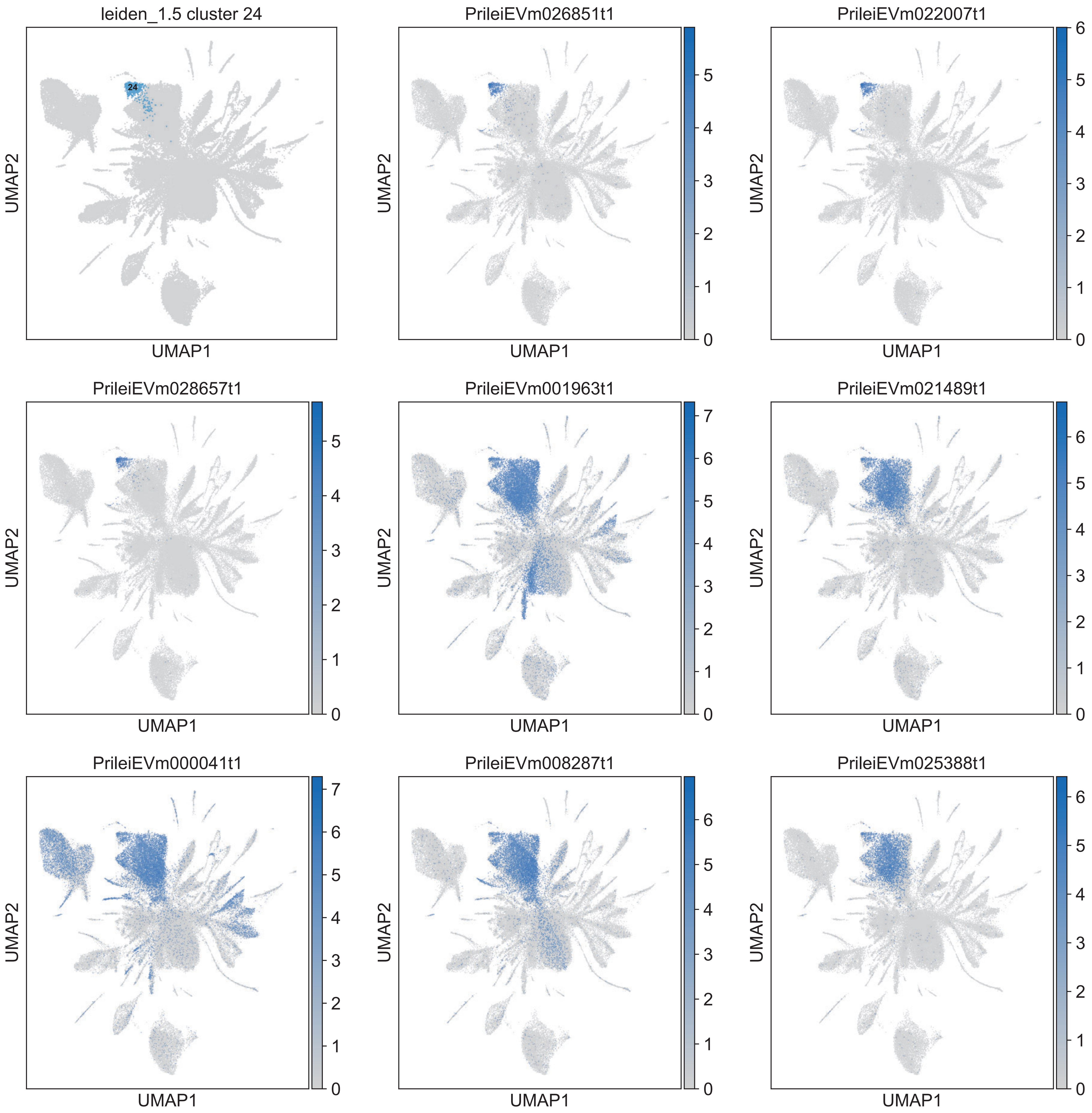

leiden_1.5 cluster 24
24
UMAP1
UMAP2
PrileiEVm026851t1
PrileiEVm022007t1
PrileiEVm028657t1
PrileiEVm001963t1
PrileiEVm021489t1
PrileiEVm000041t1
PrileiEVm008287t1
PrileiEVm025388t1

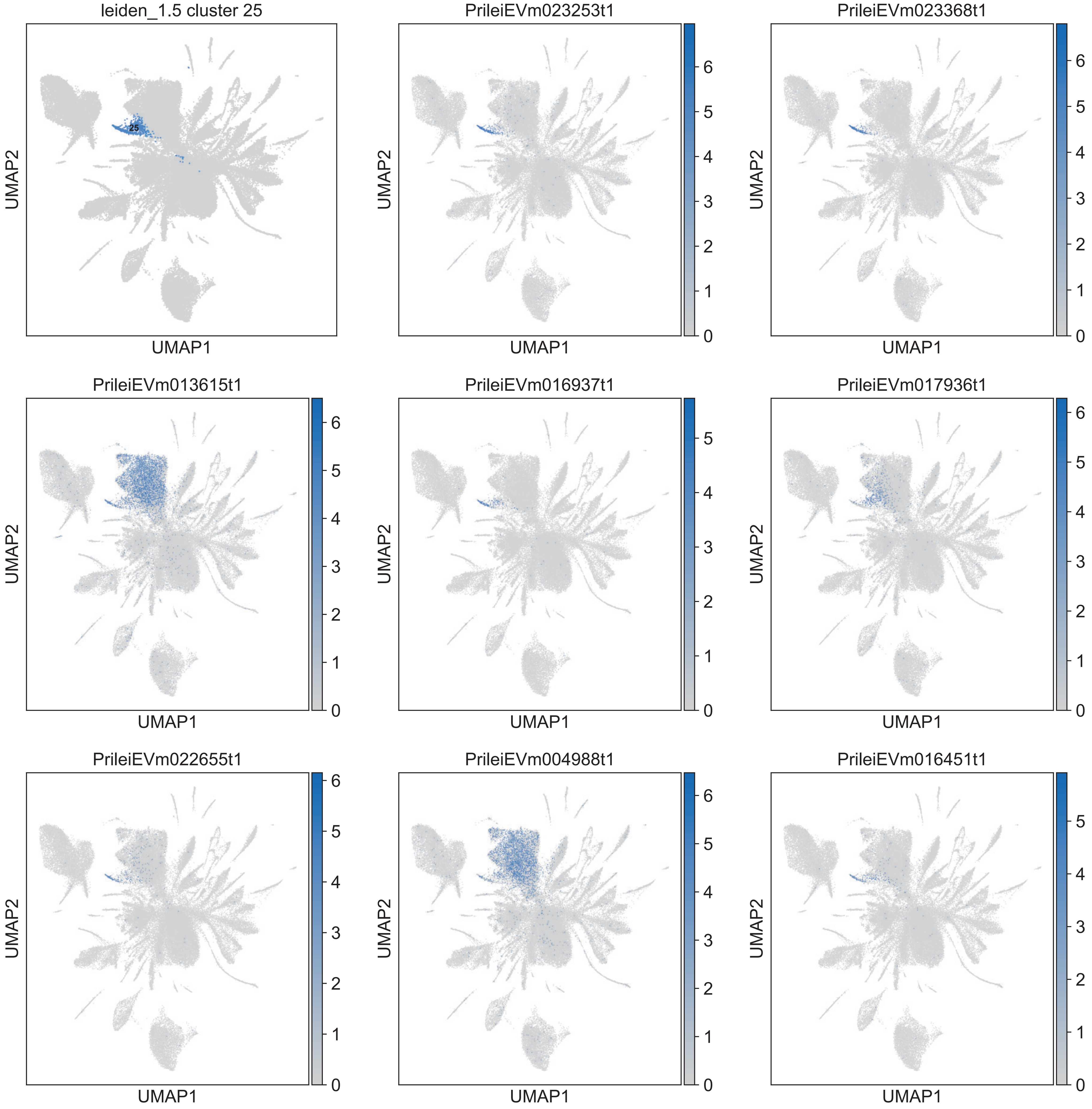
leiden_1.5 cluster 25
25
PrileiEVm023253t1
PrileiEVm023368t1
PrileiEVm013615t1
PrileiEVm016937t1
PrileiEVm017936t1
PrileiEVm022655t1
PrileiEVm004988t1
PrileiEVm016451t1
UMAP1
UMAP2

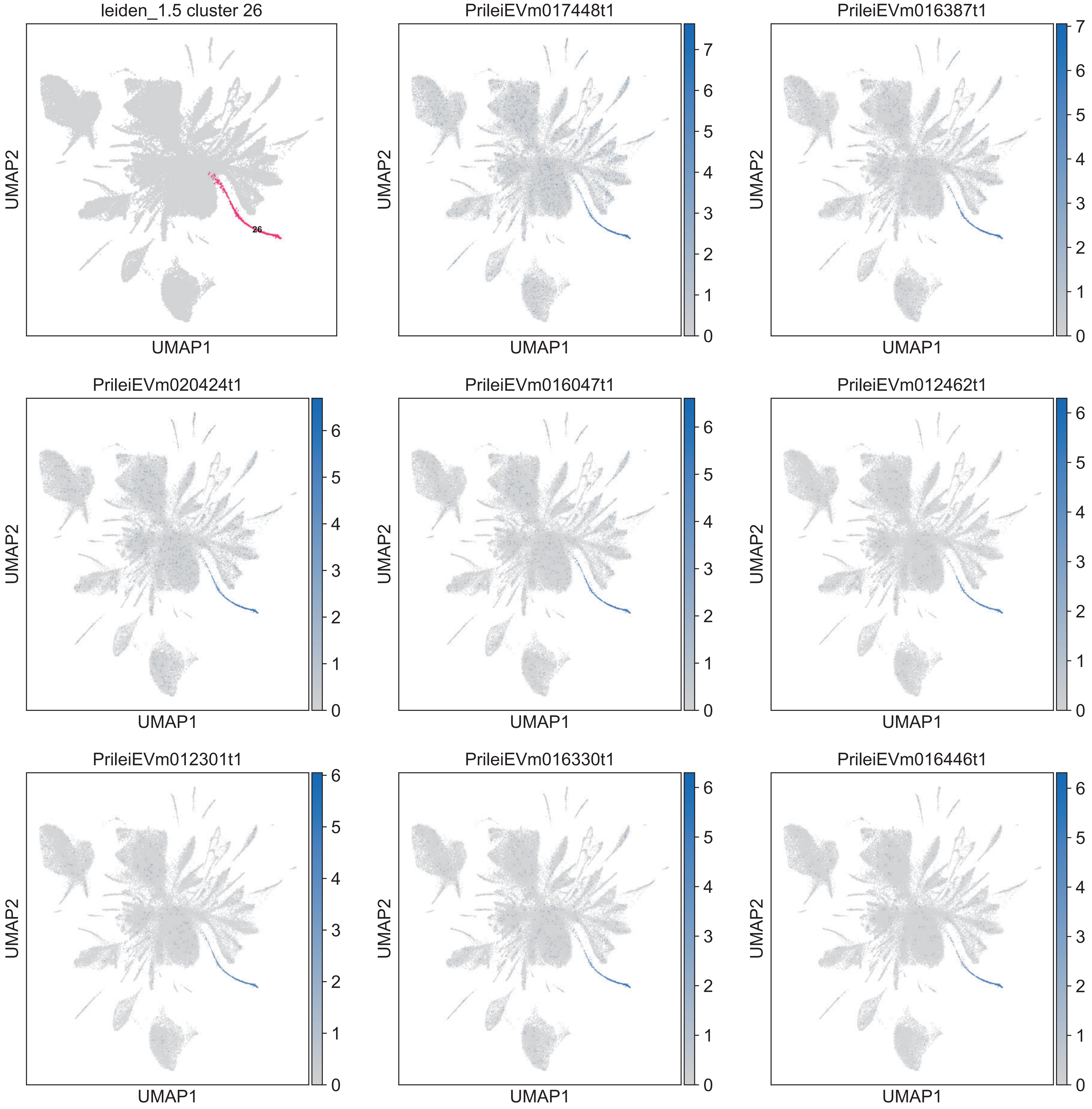
leiden_1.5 cluster 26
26
PrileiEVm017448t1
PrileiEVm016387t1
PrileiEVm020424t1
PrileiEVm016047t1
PrileiEVm012462t1
PrileiEVm012301t1
PrileiEVm016330t1
PrileiEVm016446t1
UMAP1
UMAP2

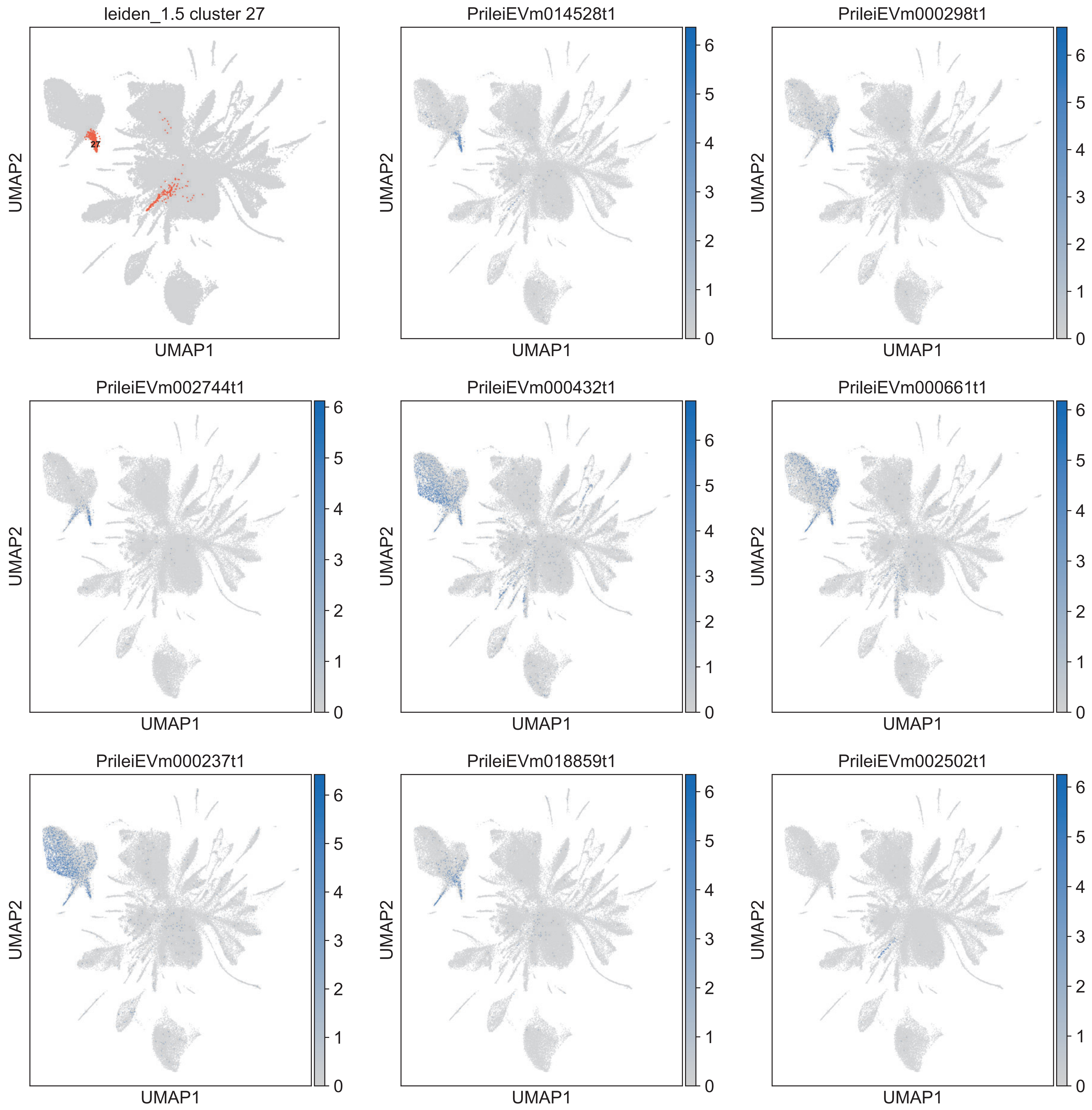

leiden_1.5 cluster 27
27
PrileiEVm014528t1
PrileiEVm000298t1
PrileiEVm002744t1
PrileiEVm000432t1
PrileiEVm000661t1
PrileiEVm000237t1
PrileiEVm018859t1
PrileiEVm002502t1
UMAP1
UMAP2
0
1
2
3
4
5
6

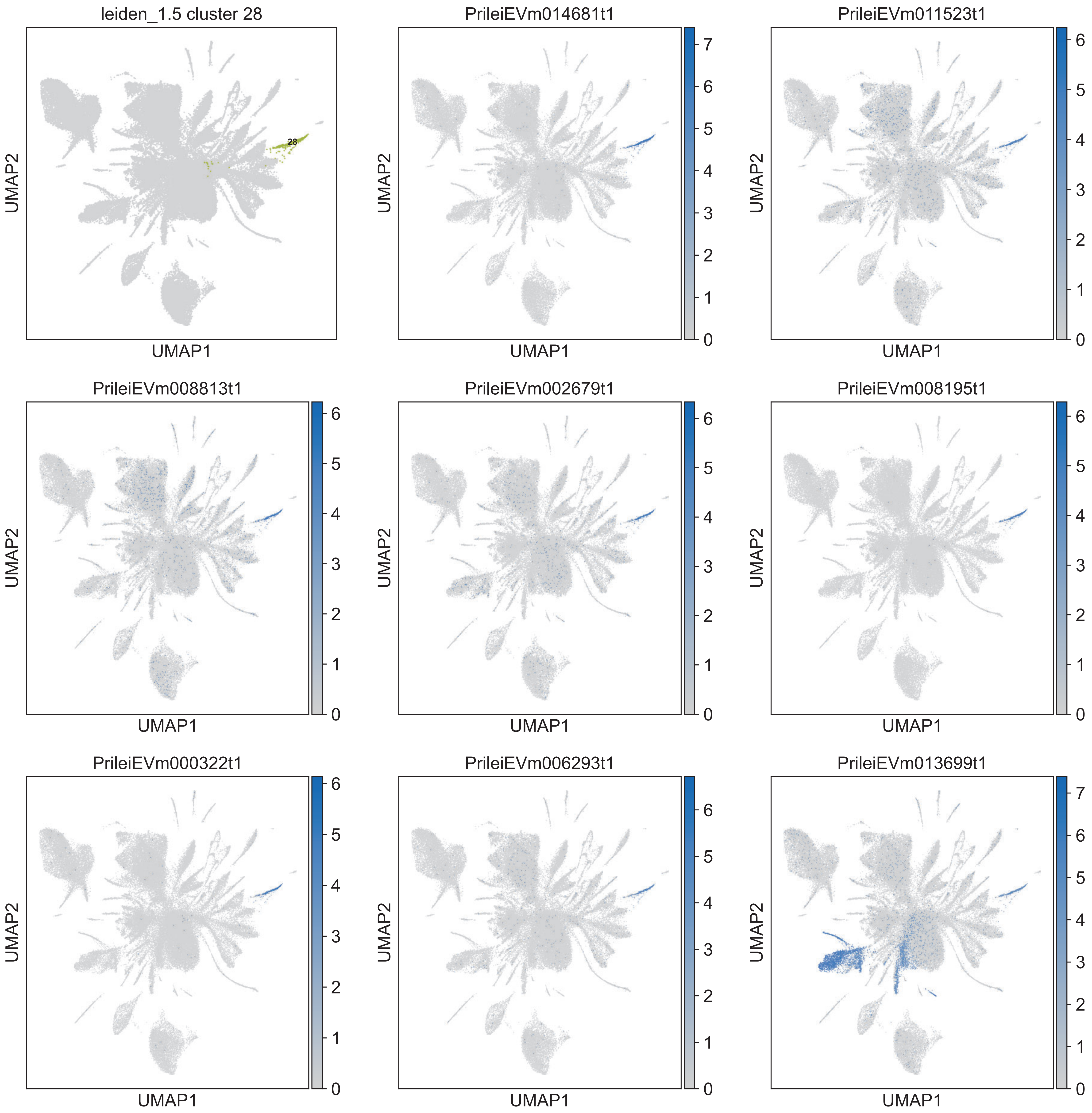

leiden_1.5 cluster 28
28
PrileiEVm014681t1
PrileiEVm011523t1
PrileiEVm008813t1
PrileiEVm002679t1
PrileiEVm008195t1
PrileiEVm000322t1
PrileiEVm006293t1
PrileiEVm013699t1
UMAP1
UMAP2

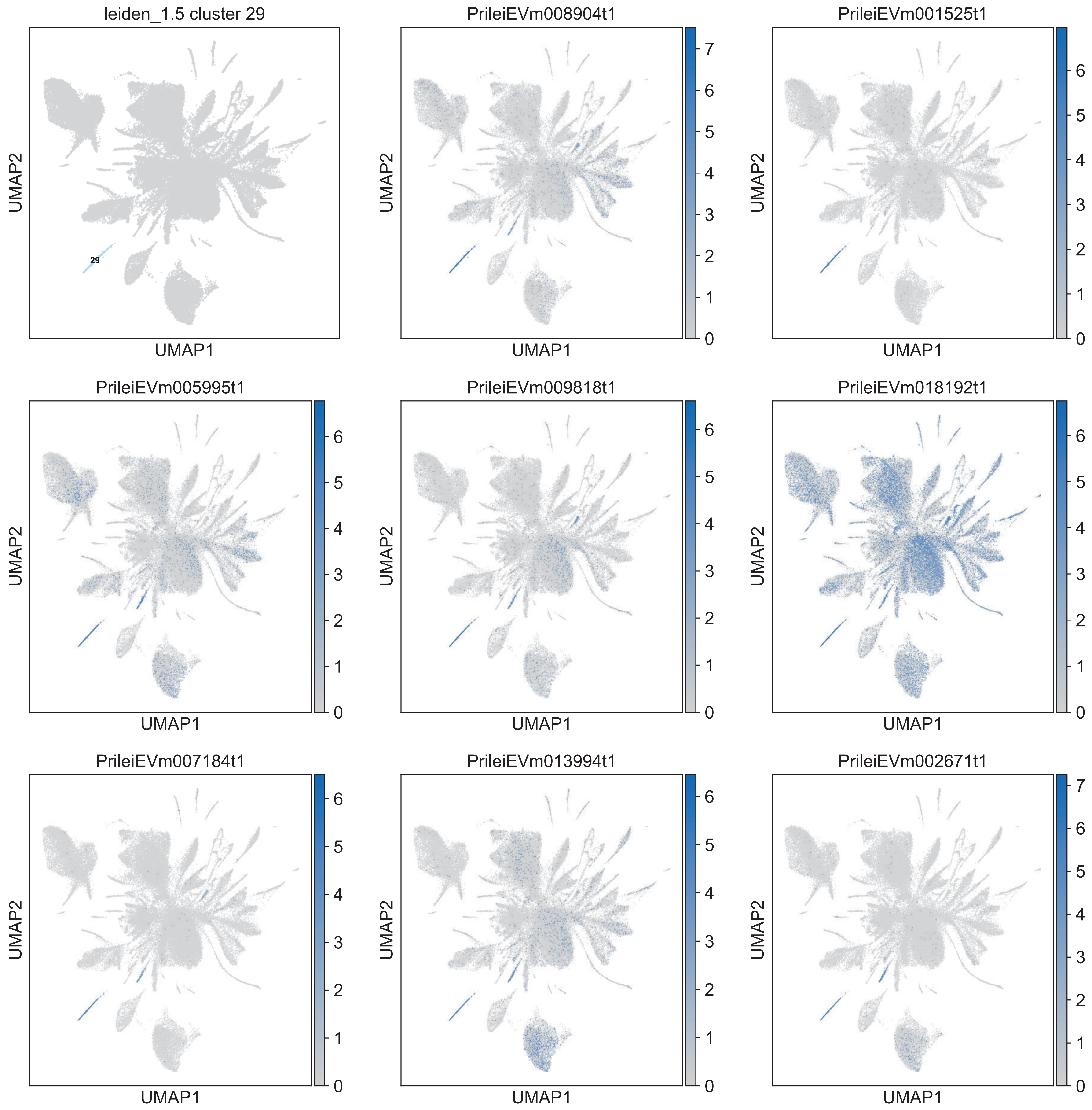
leiden_1.5 cluster 29
29
PrileiEVm008904t1
PrileiEVm001525t1
PrileiEVm005995t1
PrileiEVm009818t1
PrileiEVm018192t1
PrileiEVm007184t1
PrileiEVm013994t1
PrileiEVm002671t1
UMAP1
UMAP2

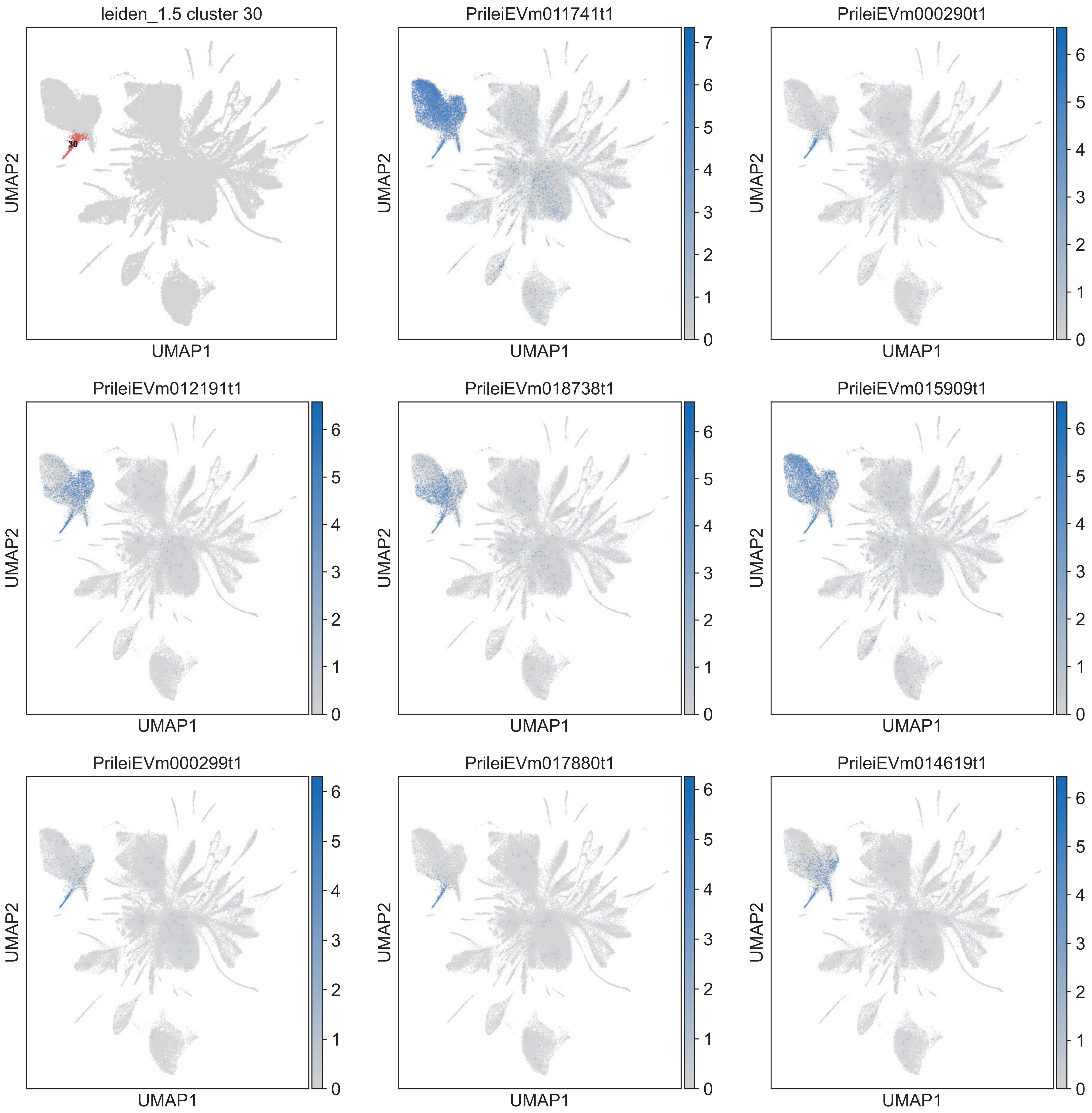

leiden_1.5 cluster 30
30
UMAP1
UMAP2
PrileiEVm011741t1
UMAP1
UMAP2
0 1 2 3 4 5 6 7
PrileiEVm000290t1
UMAP1
UMAP2
0 1 2 3 4 5 6
PrileiEVm012191t1
UMAP1
UMAP2
0 1 2 3 4 5 6
PrileiEVm018738t1
UMAP1
UMAP2
0 1 2 3 4 5 6
PrileiEVm015909t1
UMAP1
UMAP2
0 1 2 3 4 5 6
PrileiEVm000299t1
UMAP1
UMAP2
0 1 2 3 4 5 6
PrileiEVm017880t1
UMAP1
UMAP2
0 1 2 3 4 5 6
PrileiEVm014619t1
UMAP1
UMAP2
0 1 2 3 4 5 6

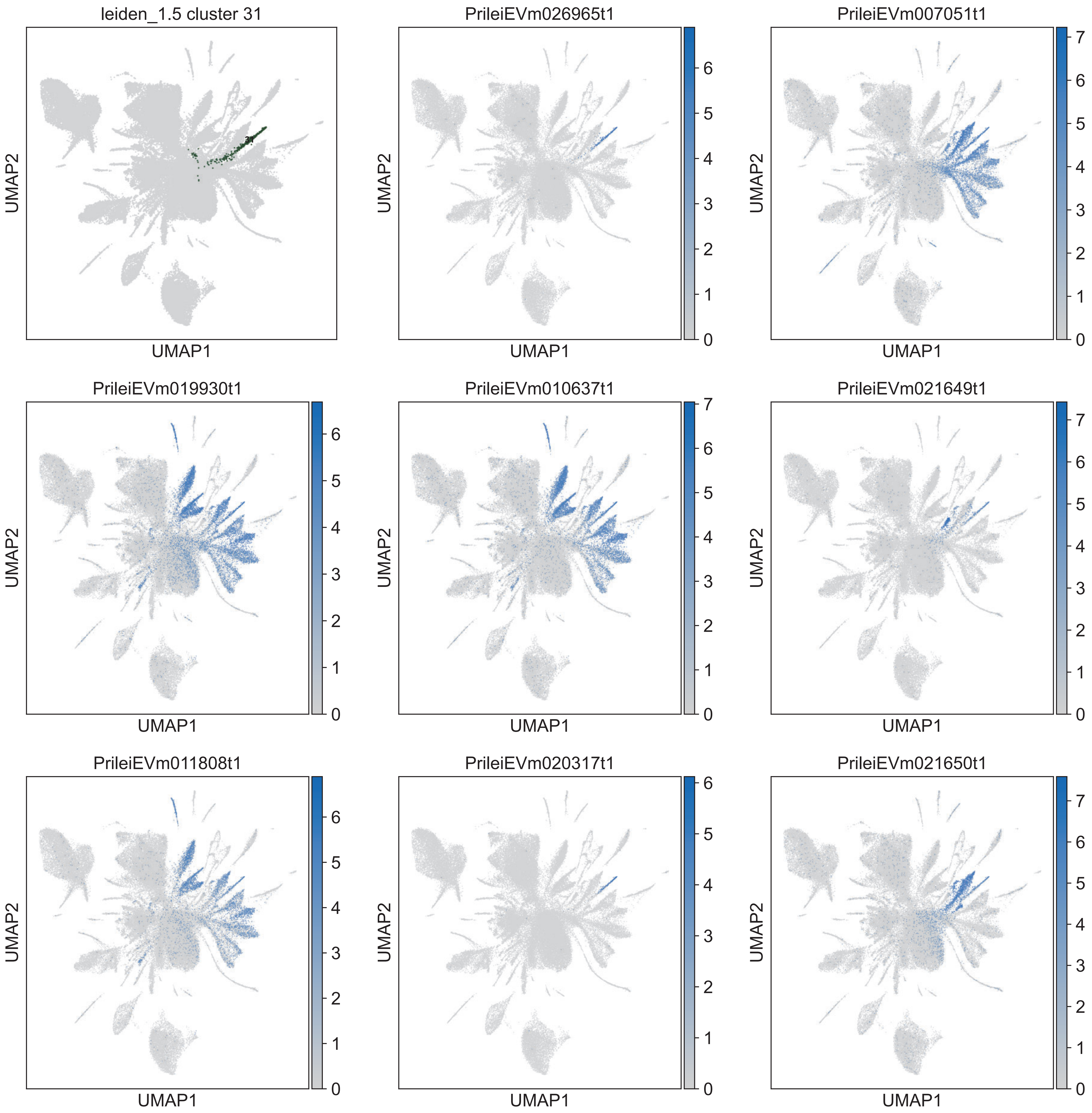
leiden_1.5 cluster 31
PrileiEVm026965t1
PrileiEVm007051t1
PrileiEVm019930t1
PrileiEVm010637t1
PrileiEVm021649t1
PrileiEVm011808t1
PrileiEVm020317t1
PrileiEVm021650t1
UMAP1
UMAP2

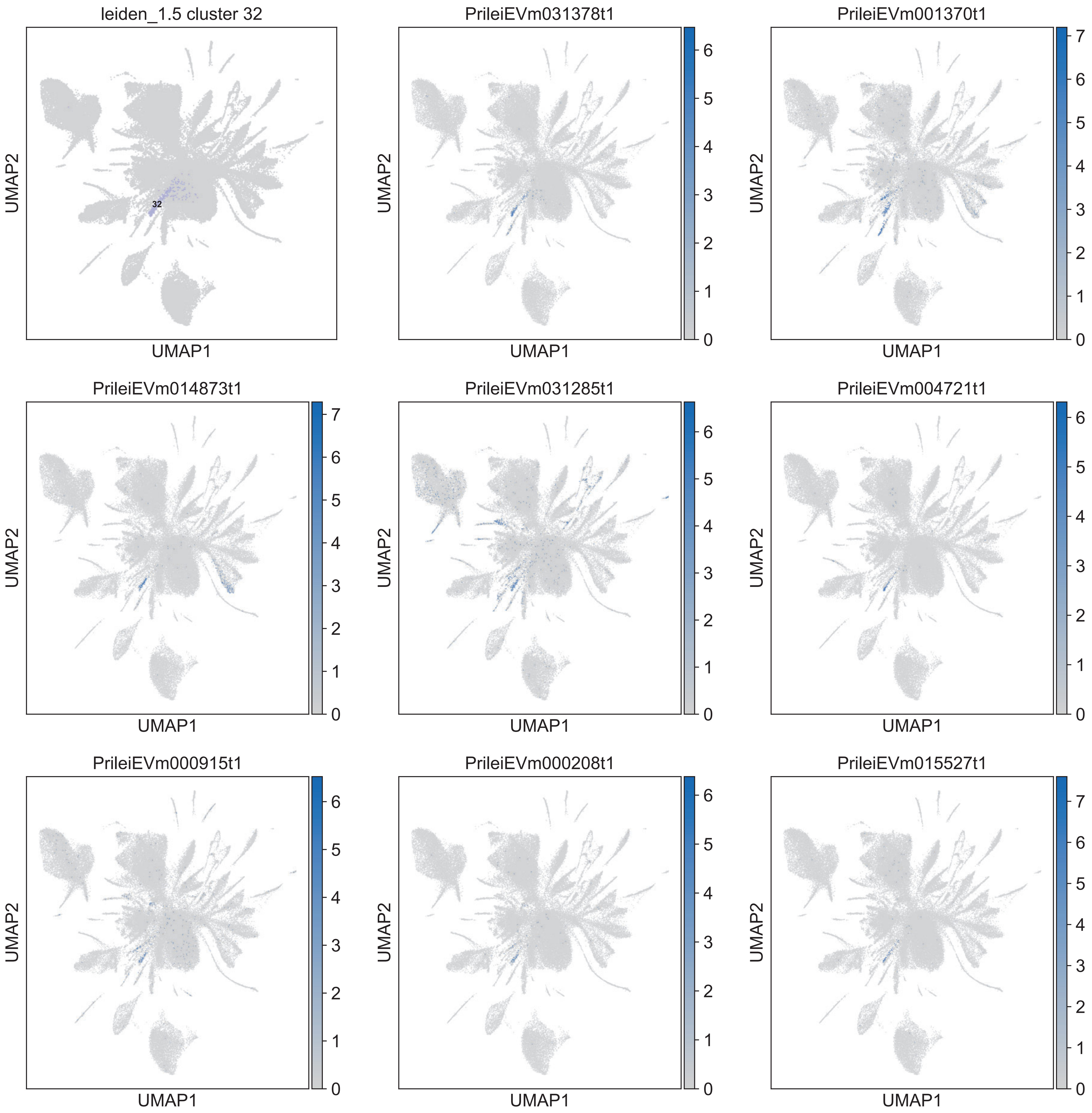

leiden_1.5 cluster 32
32
PrileiEVm031378t1
PrileiEVm001370t1
PrileiEVm014873t1
PrileiEVm031285t1
PrileiEVm004721t1
PrileiEVm000915t1
PrileiEVm000208t1
PrileiEVm015527t1
UMAP1
UMAP2

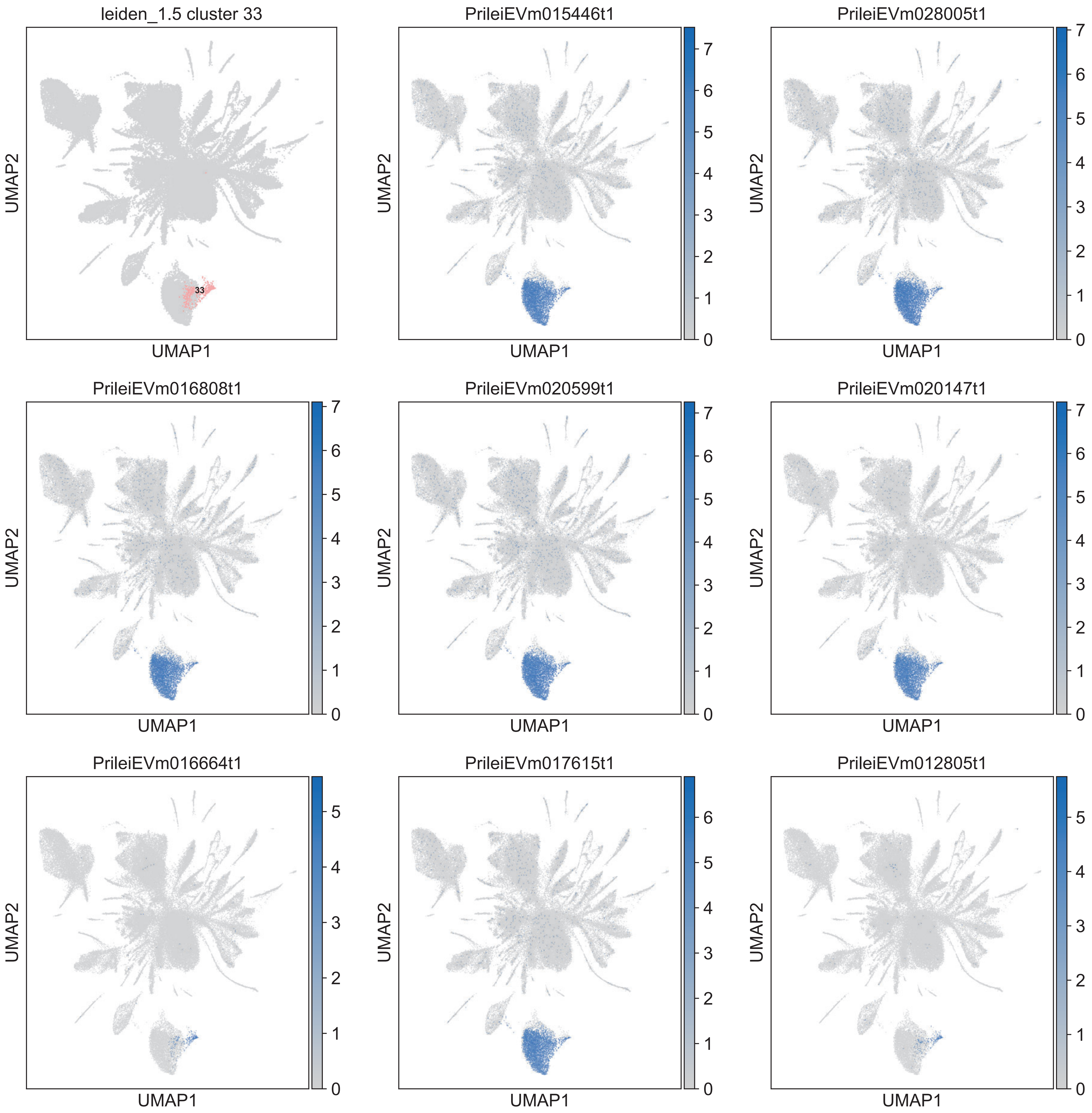

leiden_1.5 cluster 33
33
PrileiEVm015446t1
PrileiEVm028005t1
PrileiEVm016808t1
PrileiEVm020599t1
PrileiEVm020147t1
PrileiEVm016664t1
PrileiEVm017615t1
PrileiEVm012805t1
UMAP1
UMAP2

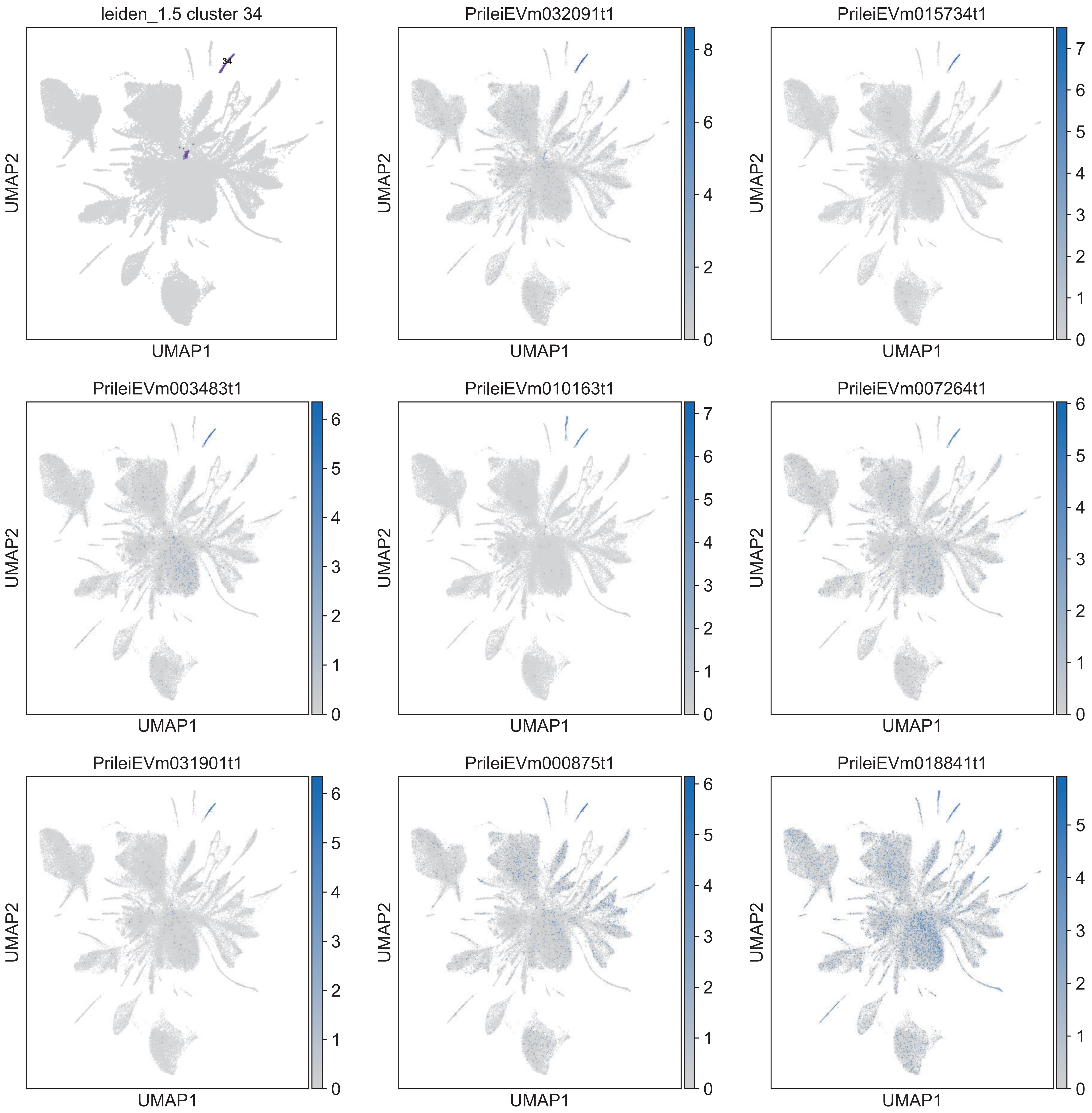

leiden_1.5 cluster 34
34
PrileiEVm032091t1
PrileiEVm015734t1
PrileiEVm003483t1
PrileiEVm010163t1
PrileiEVm007264t1
PrileiEVm031901t1
PrileiEVm000875t1
PrileiEVm018841t1
UMAP1
UMAP2

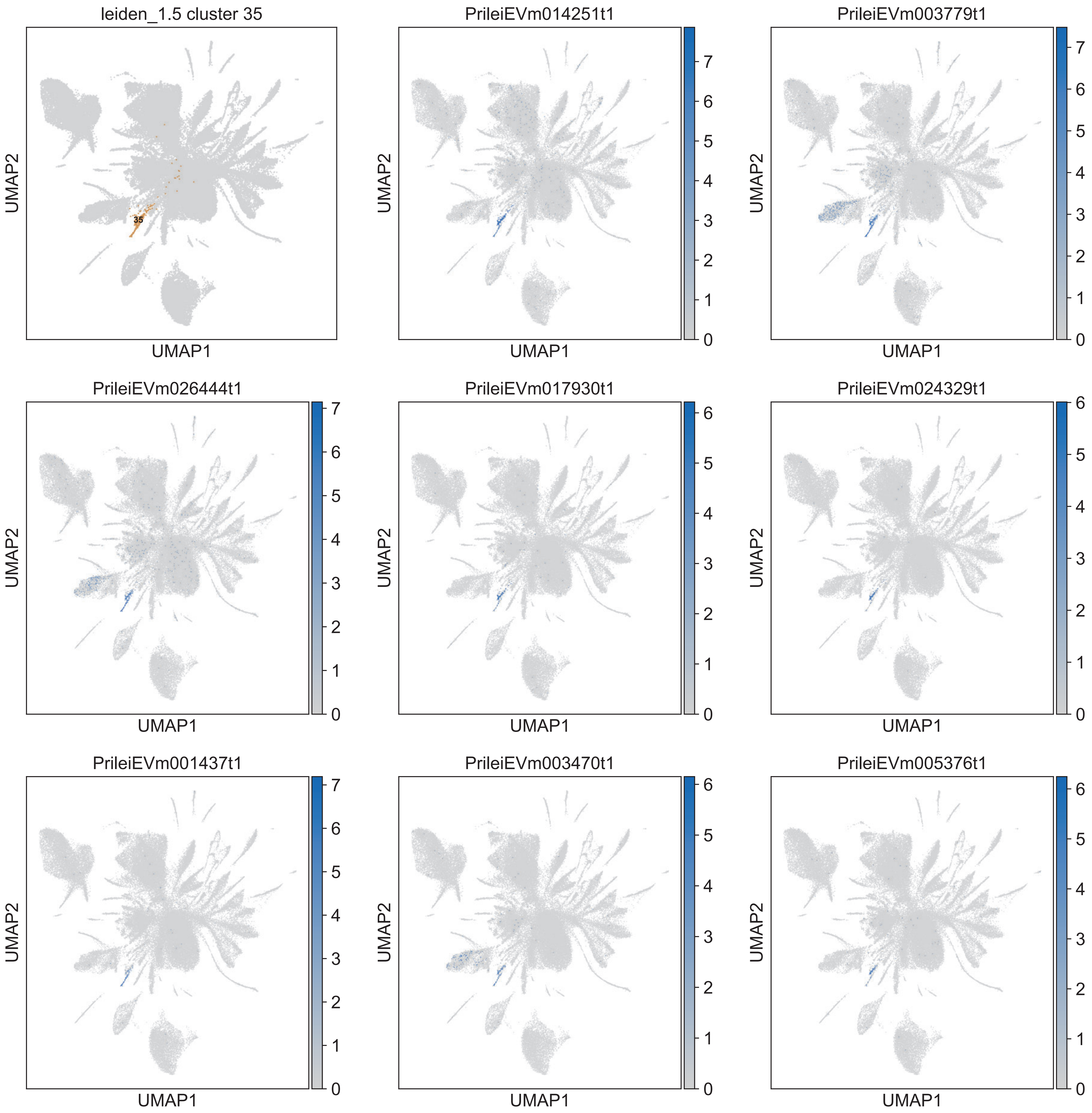

leiden_1.5 cluster 35
35
PrileiEVm014251t1
PrileiEVm003779t1
PrileiEVm026444t1
PrileiEVm017930t1
PrileiEVm024329t1
PrileiEVm001437t1
PrileiEVm003470t1
PrileiEVm005376t1
UMAP1
UMAP2

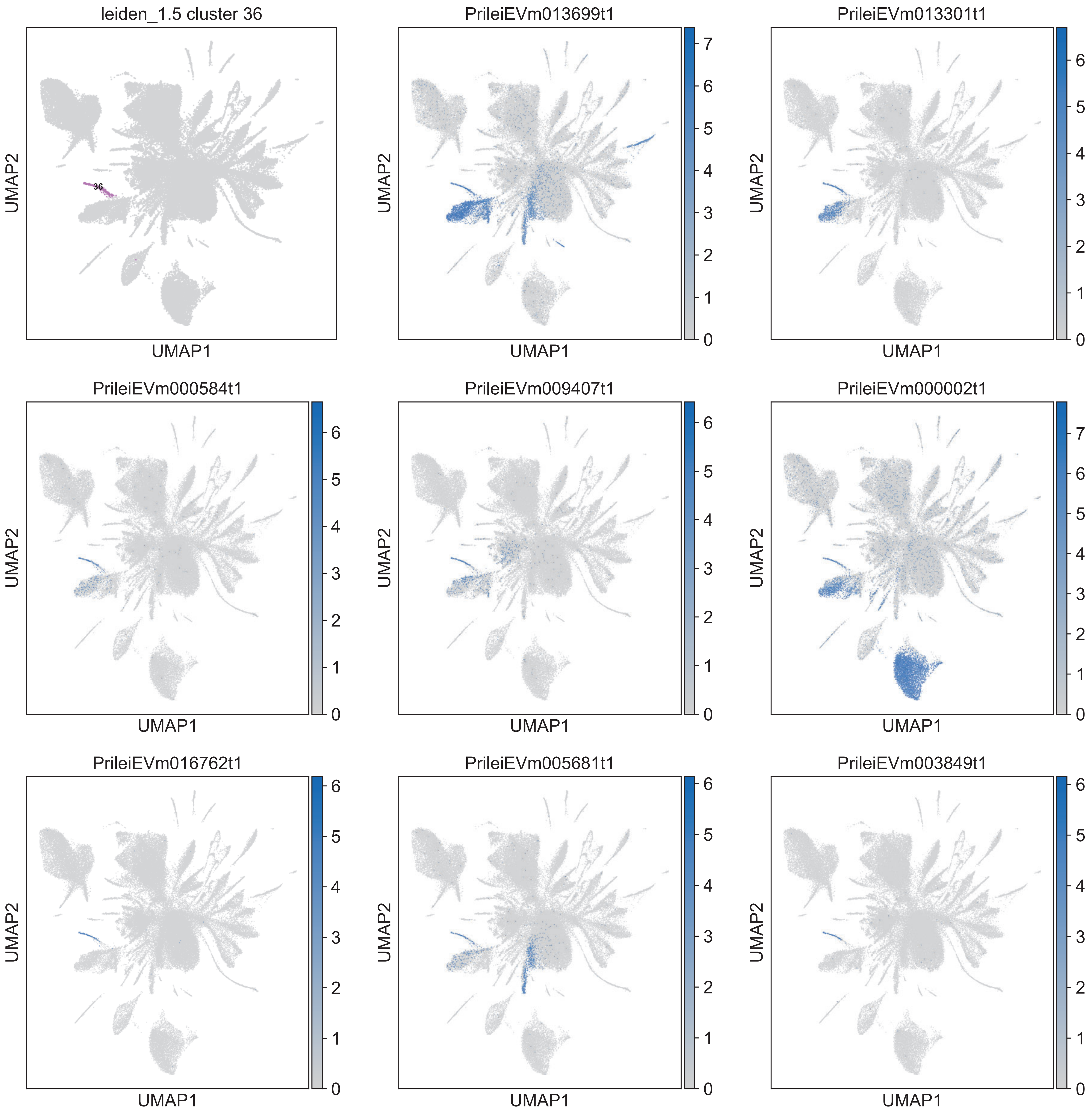

leiden_1.5 cluster 36
36
PrileiEVm013699t1
PrileiEVm013301t1
PrileiEVm000584t1
PrileiEVm009407t1
PrileiEVm000002t1
PrileiEVm016762t1
PrileiEVm005681t1
PrileiEVm003849t1
UMAP1
UMAP2

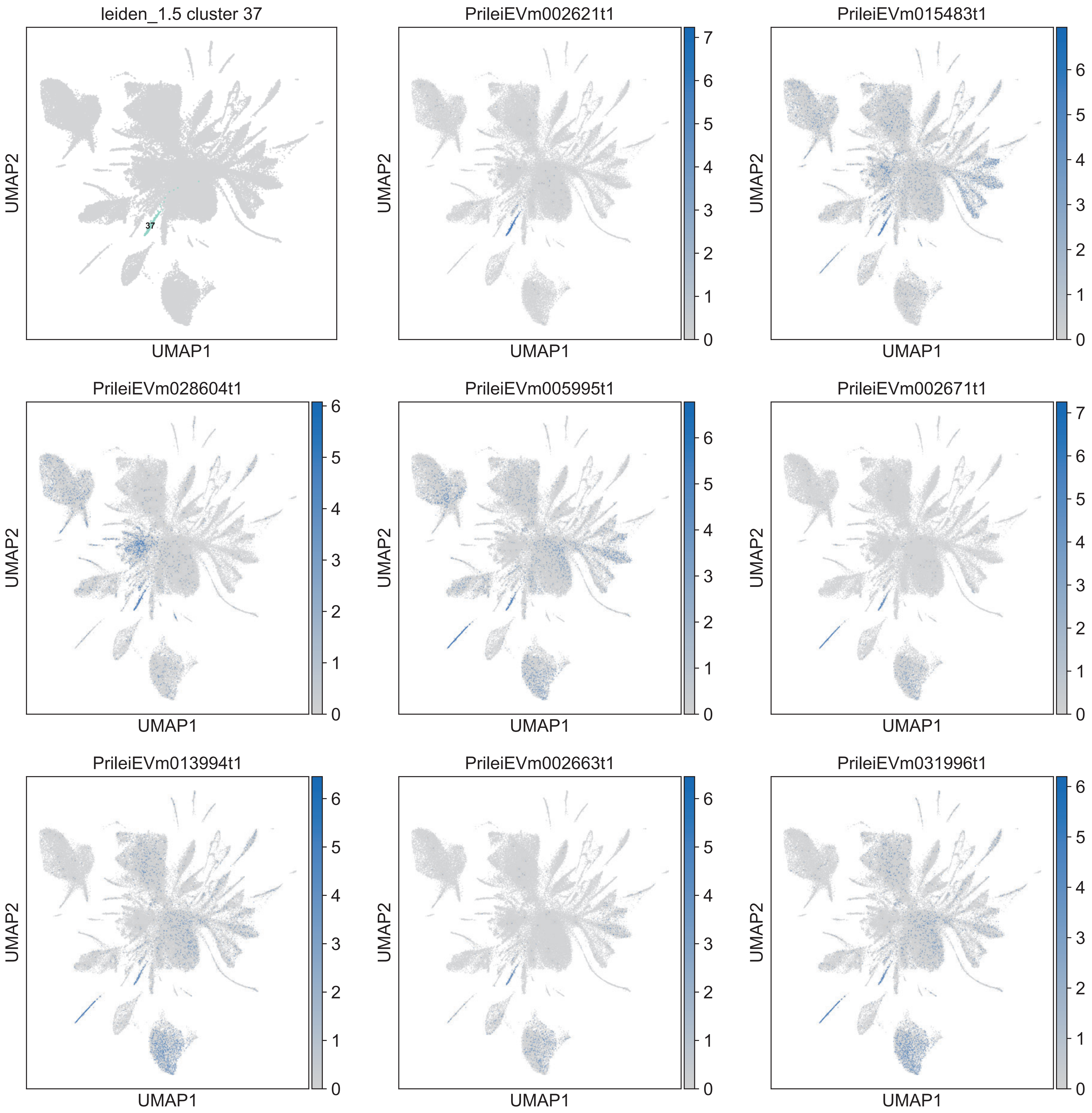

leiden_1.5 cluster 37
37
PrileiEVm002621t1
PrileiEVm015483t1
PrileiEVm028604t1
PrileiEVm005995t1
PrileiEVm002671t1
PrileiEVm013994t1
PrileiEVm002663t1
PrileiEVm031996t1
UMAP1
UMAP2

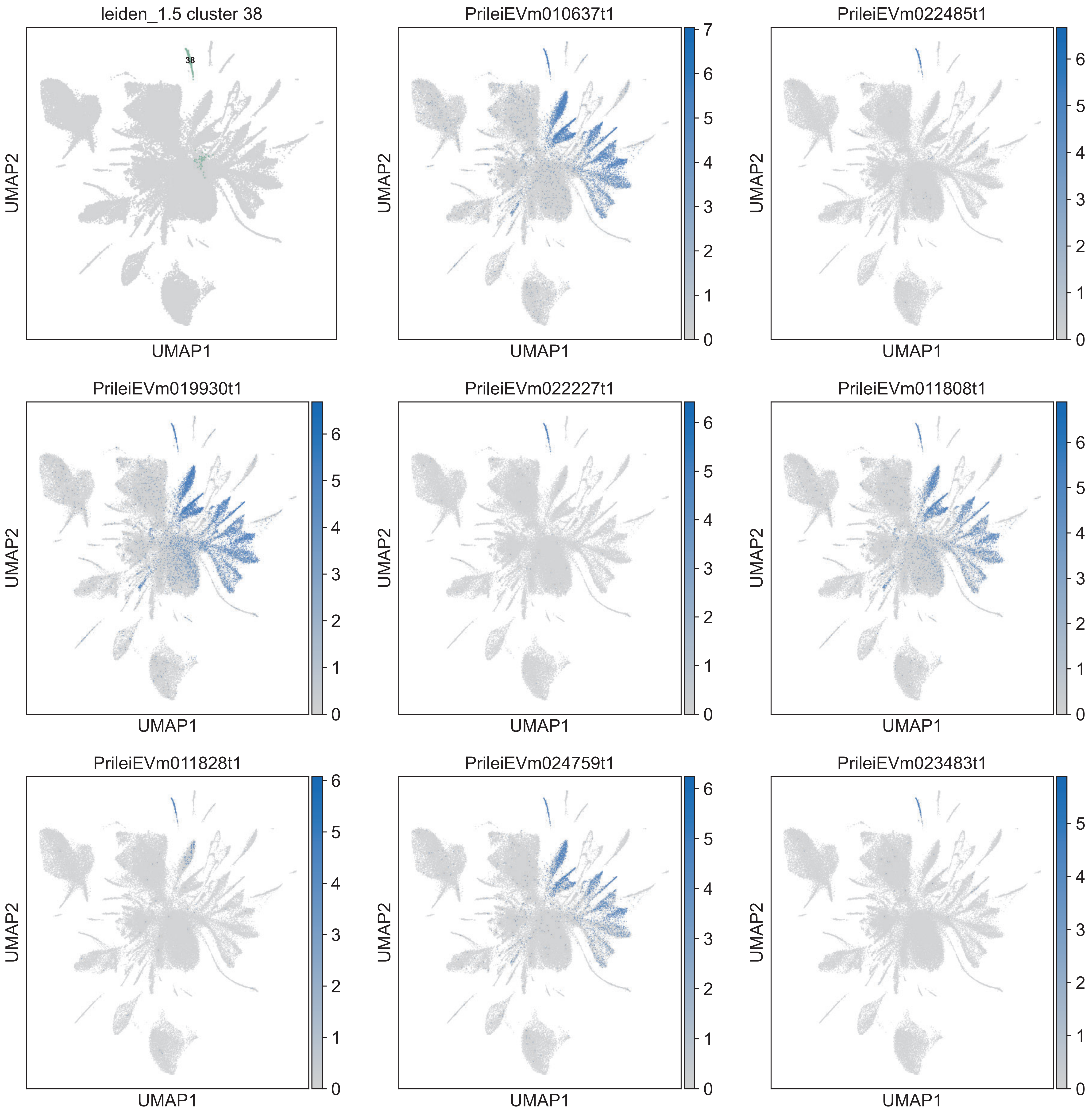

leiden_1.5 cluster 38
38
PrileiEVm010637t1
PrileiEVm022485t1
PrileiEVm019930t1
PrileiEVm022227t1
PrileiEVm011808t1
PrileiEVm011828t1
PrileiEVm024759t1
PrileiEVm023483t1
UMAP1
UMAP2

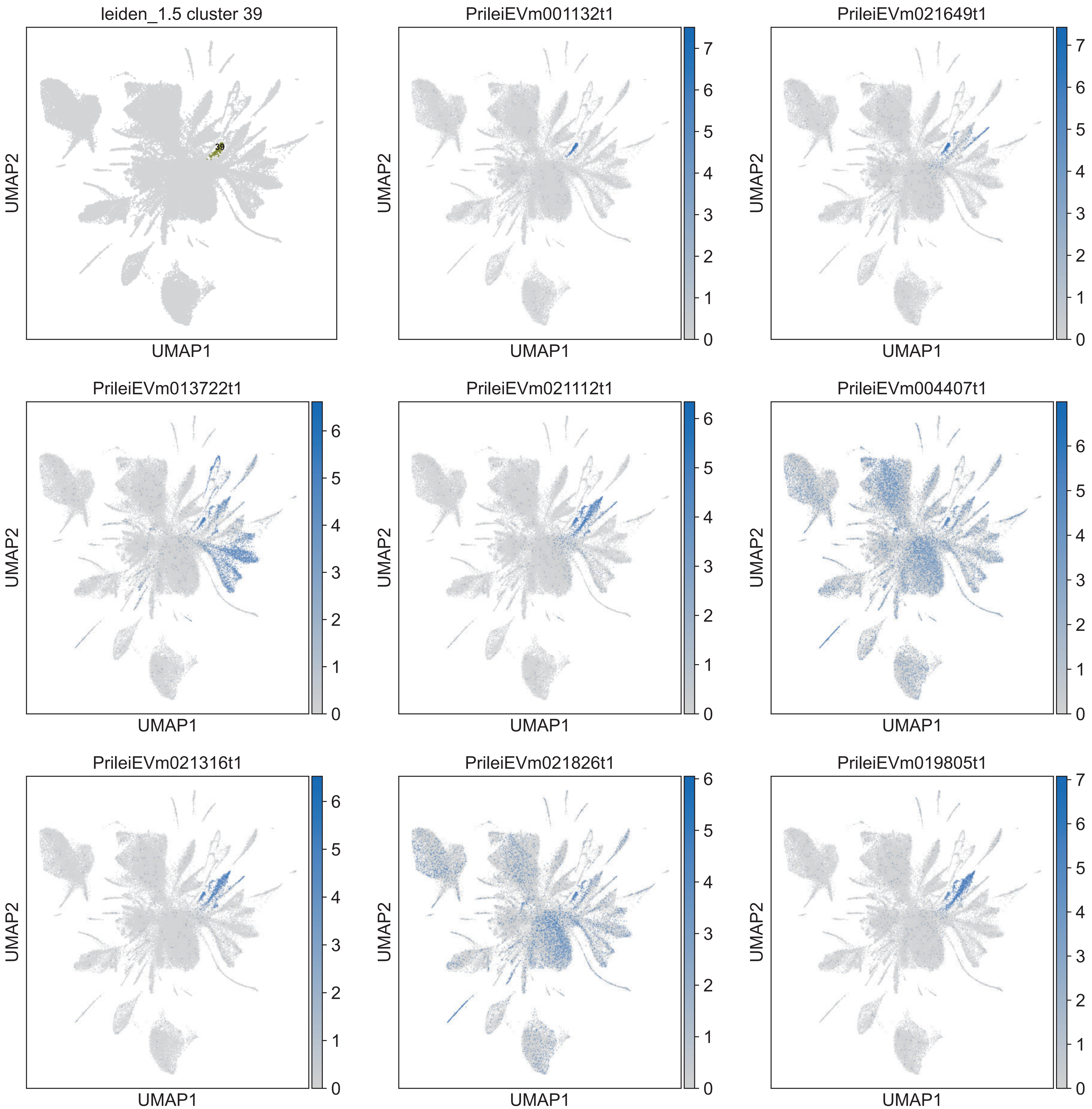

leiden_1.5 cluster 39
39
PrileiEVm001132t1
PrileiEVm021649t1
PrileiEVm013722t1
PrileiEVm021112t1
PrileiEVm004407t1
PrileiEVm021316t1
PrileiEVm021826t1
PrileiEVm019805t1
UMAP1
UMAP2

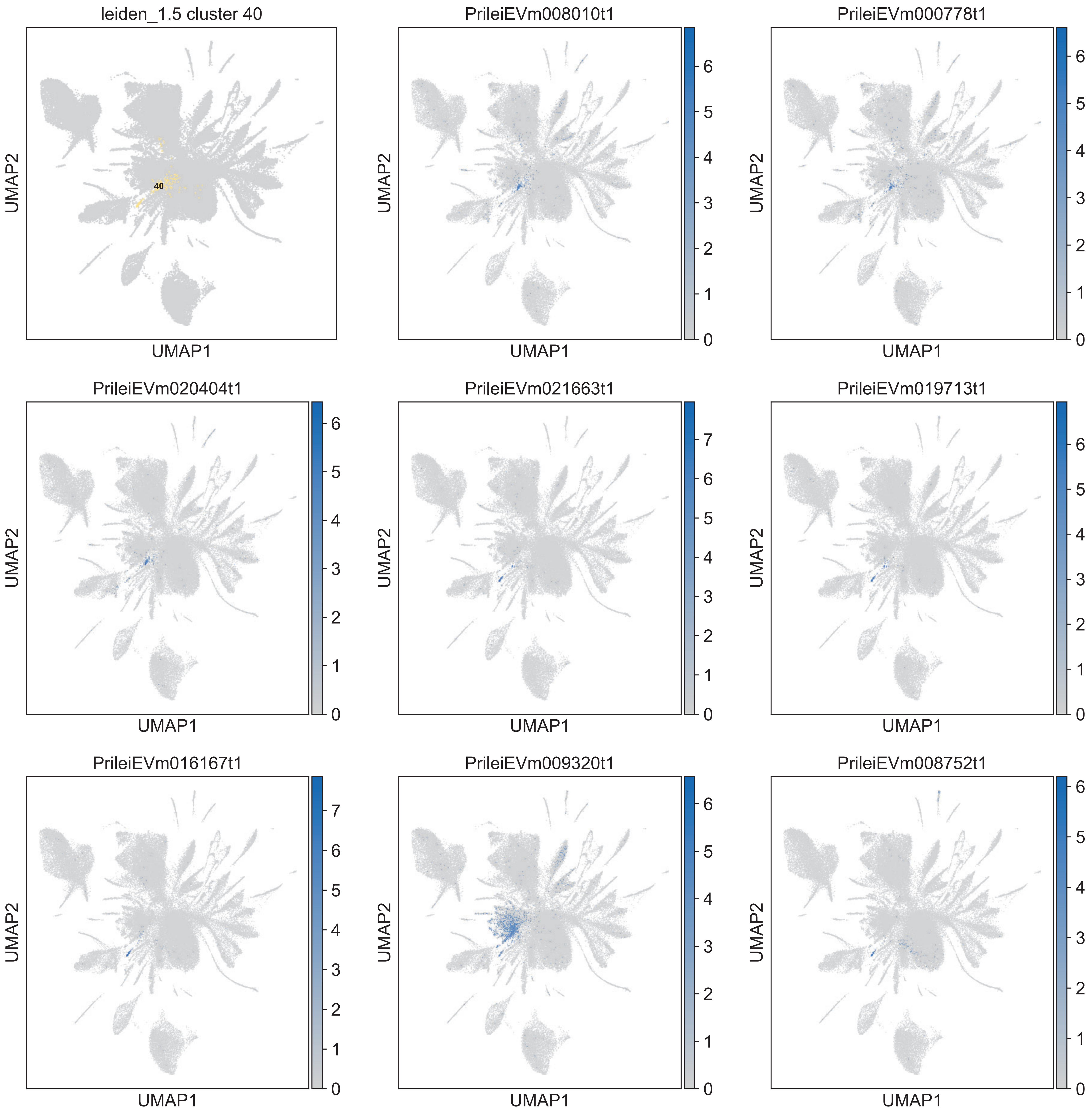

leiden_1.5 cluster 40
40
PrileiEVm008010t1
PrileiEVm000778t1
PrileiEVm020404t1
PrileiEVm021663t1
PrileiEVm019713t1
PrileiEVm016167t1
PrileiEVm009320t1
PrileiEVm008752t1
UMAP1
UMAP2

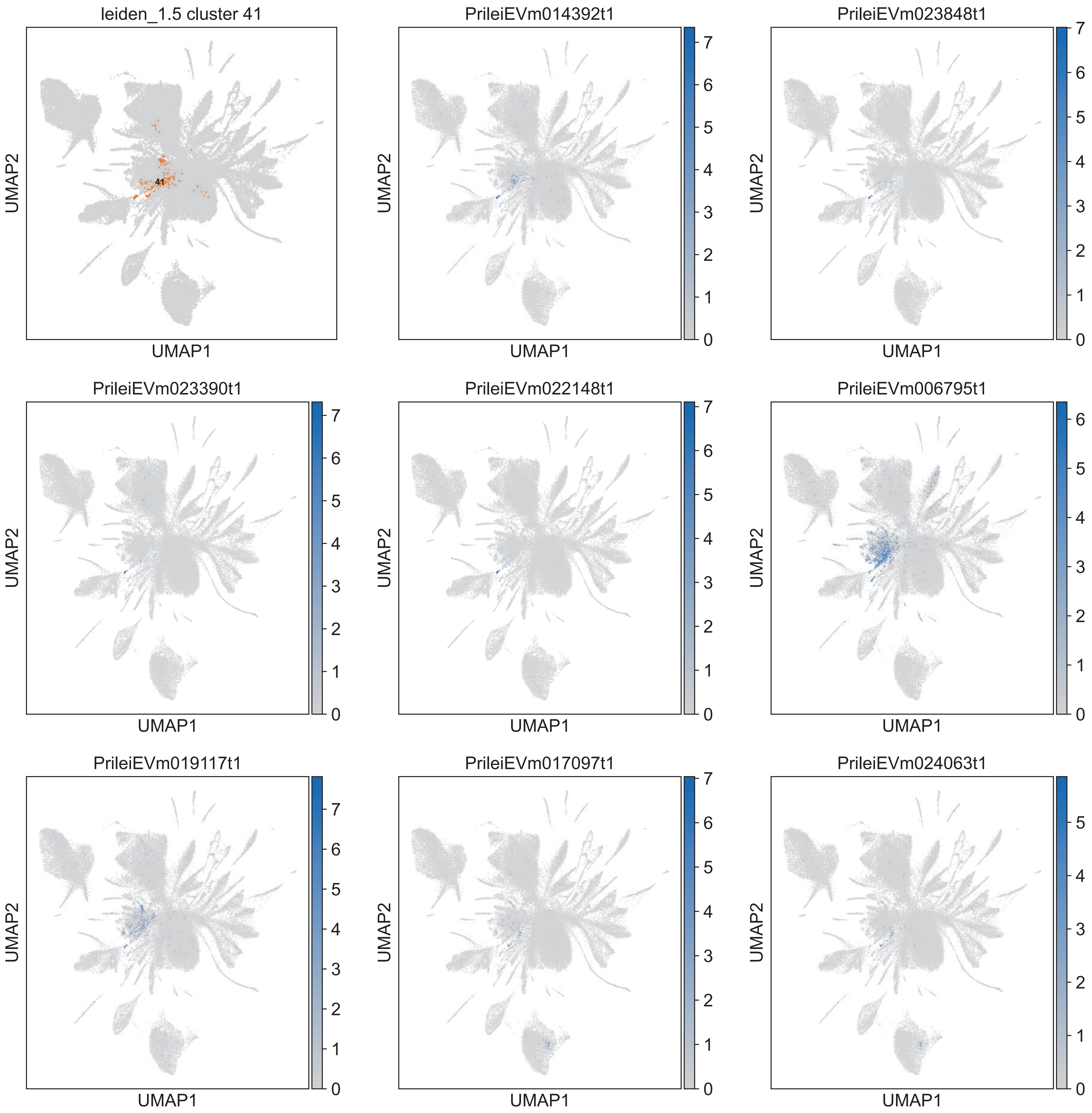
leiden_1.5 cluster 41
41
PrileiEVm014392t1
PrileiEVm023848t1
PrileiEVm023390t1
PrileiEVm022148t1
PrileiEVm006795t1
PrileiEVm019117t1
PrileiEVm017097t1
PrileiEVm024063t1
UMAP1
UMAP2

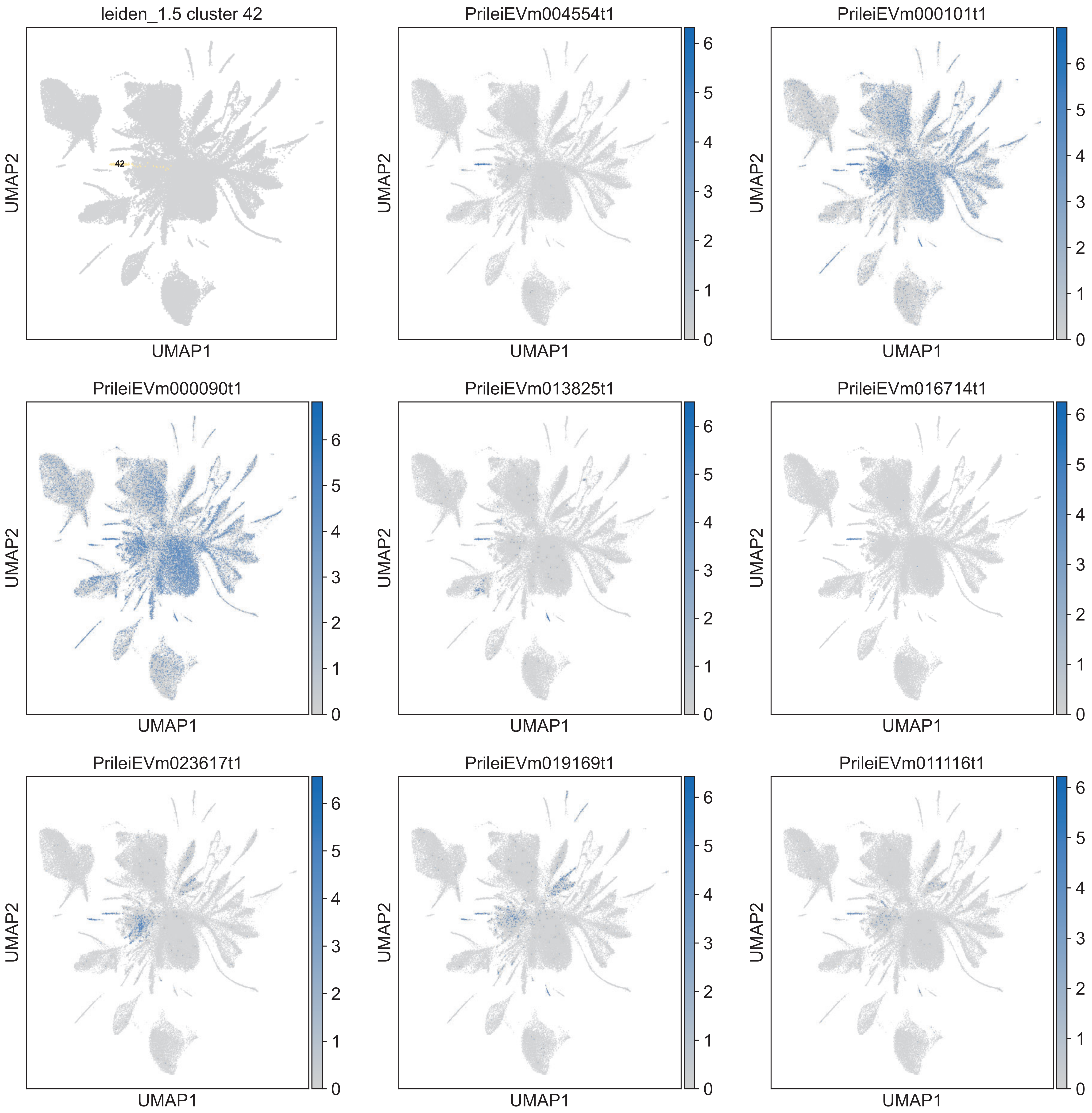
leiden_1.5 cluster 42
42
PrileiEVm004554t1
PrileiEVm000101t1
PrileiEVm000090t1
PrileiEVm013825t1
PrileiEVm016714t1
PrileiEVm023617t1
PrileiEVm019169t1
PrileiEVm011116t1
UMAP1
UMAP2
0
1
2
3
4
5
6

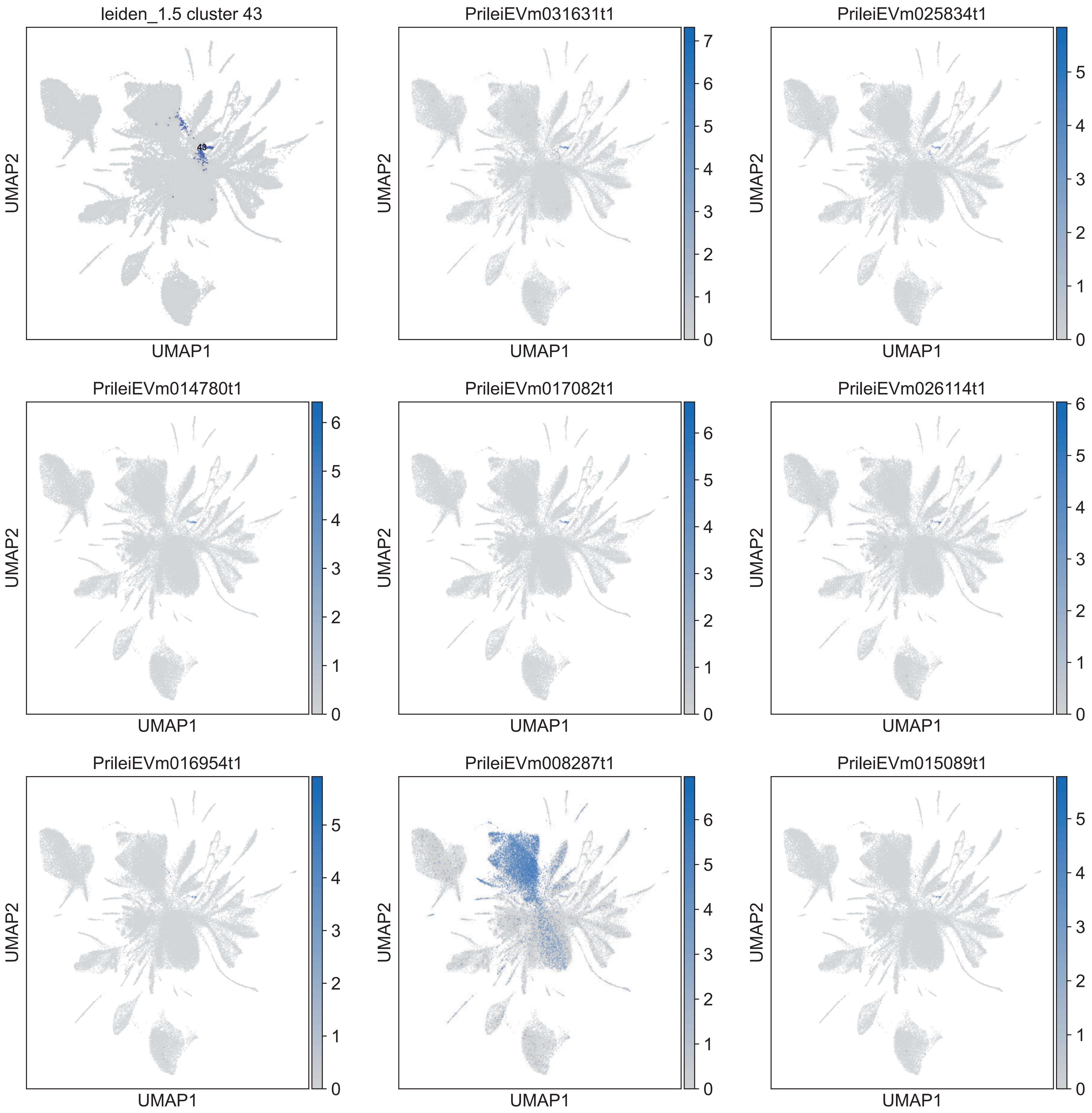

leiden_1.5 cluster 43
43
PrileiEVm031631t1
PrileiEVm025834t1
PrileiEVm014780t1
PrileiEVm017082t1
PrileiEVm026114t1
PrileiEVm016954t1
PrileiEVm008287t1
PrileiEVm015089t1
UMAP1
UMAP2

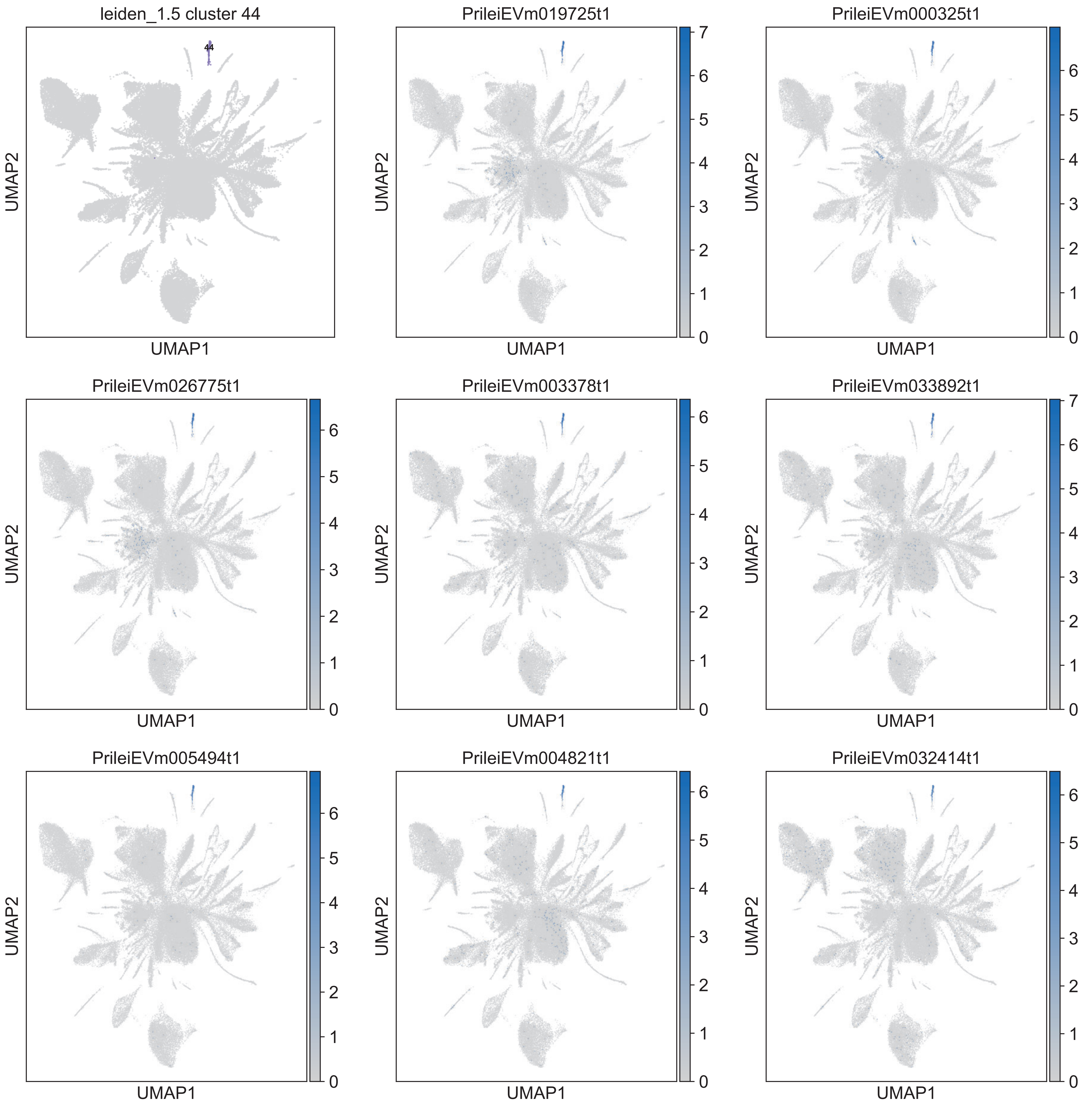

leiden_1.5 cluster 44
44
PrileiEVm019725t1
PrileiEVm000325t1
PrileiEVm026775t1
PrileiEVm003378t1
PrileiEVm033892t1
PrileiEVm005494t1
PrileiEVm004821t1
PrileiEVm032414t1
UMAP1
UMAP2

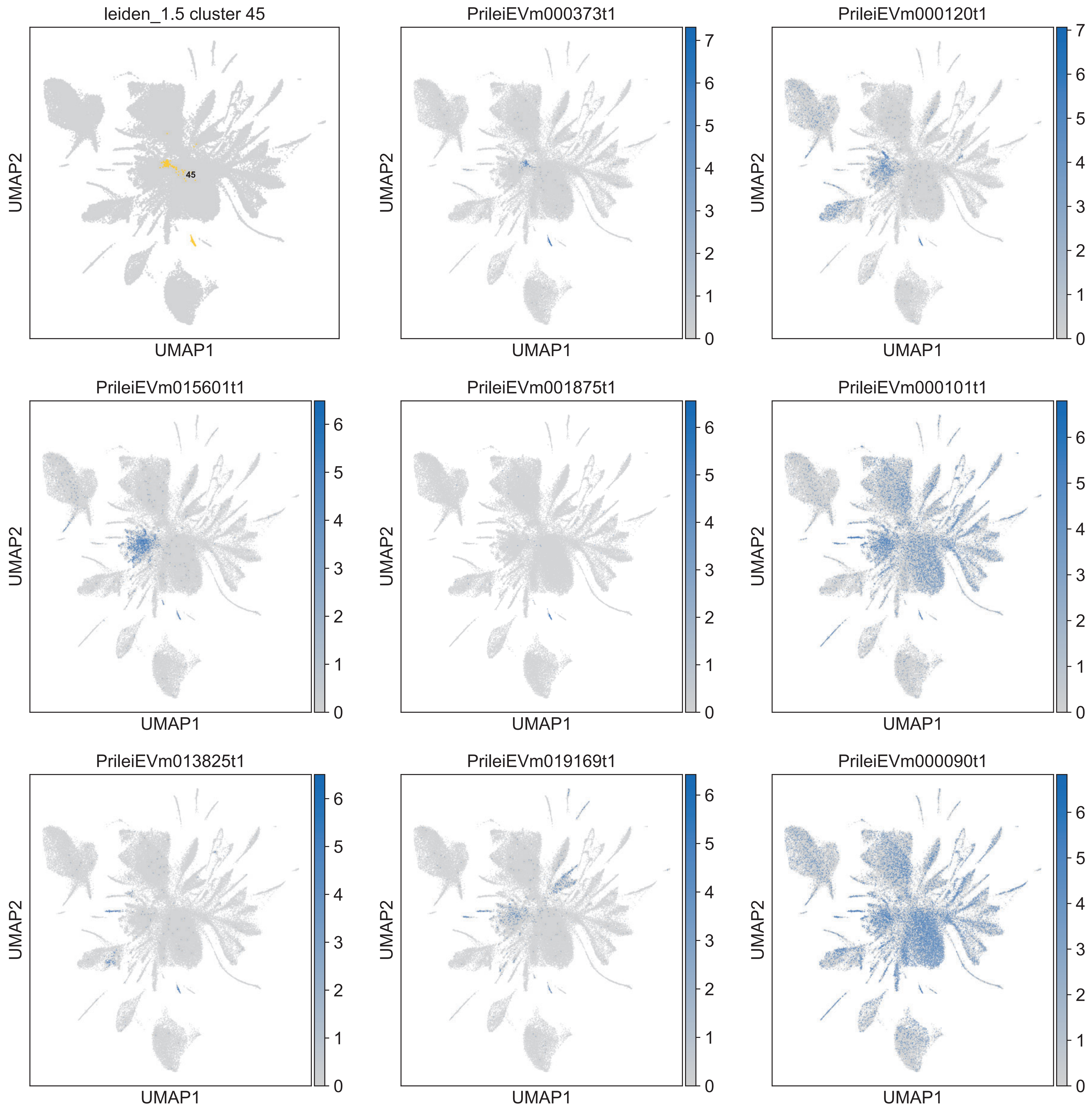
leiden_1.5 cluster 45
45
PrileiEVm000373t1
PrileiEVm000120t1
PrileiEVm015601t1
PrileiEVm001875t1
PrileiEVm000101t1
PrileiEVm013825t1
PrileiEVm019169t1
PrileiEVm000090t1
UMAP1
UMAP2

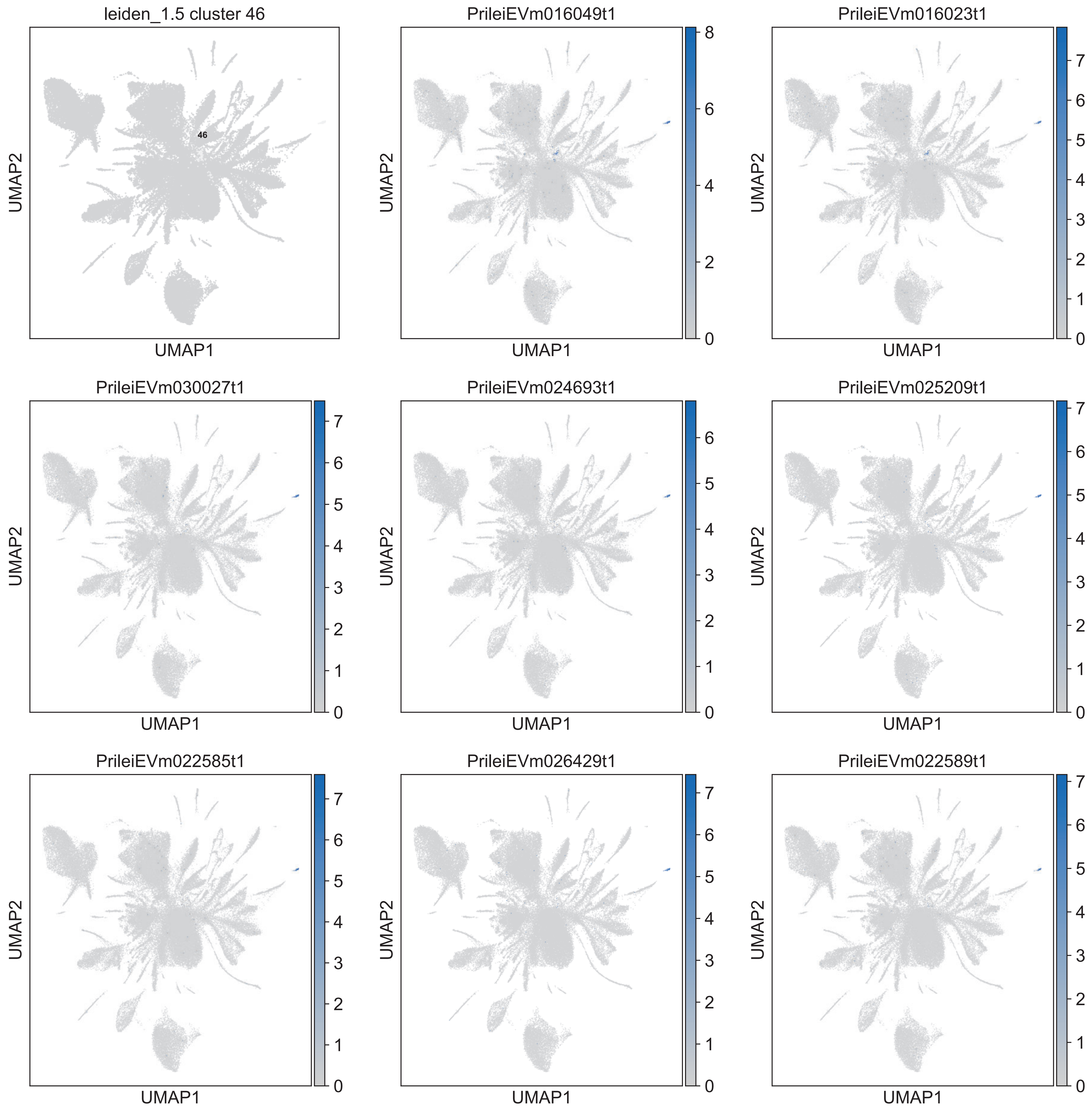

leiden_1.5 cluster 46
46
UMAP2
UMAP1
PrileiEVm016049t1
8
6
4
2
0
UMAP2
UMAP1
PrileiEVm016023t1
7
6
5
4
3
2
1
0
UMAP2
UMAP1
PrileiEVm030027t1
7
6
5
4
3
2
1
0
UMAP2
UMAP1
PrileiEVm024693t1
6
5
4
3
2
1
0
UMAP2
UMAP1
PrileiEVm025209t1
7
6
5
4
3
2
1
0
UMAP2
UMAP1
PrileiEVm022585t1
7
6
5
4
3
2
1
0
UMAP2
UMAP1
PrileiEVm026429t1
7
6
5
4
3
2
1
0
UMAP2
UMAP1
PrileiEVm022589t1
7
6
5
4
3
2
1
0
UMAP2
UMAP1

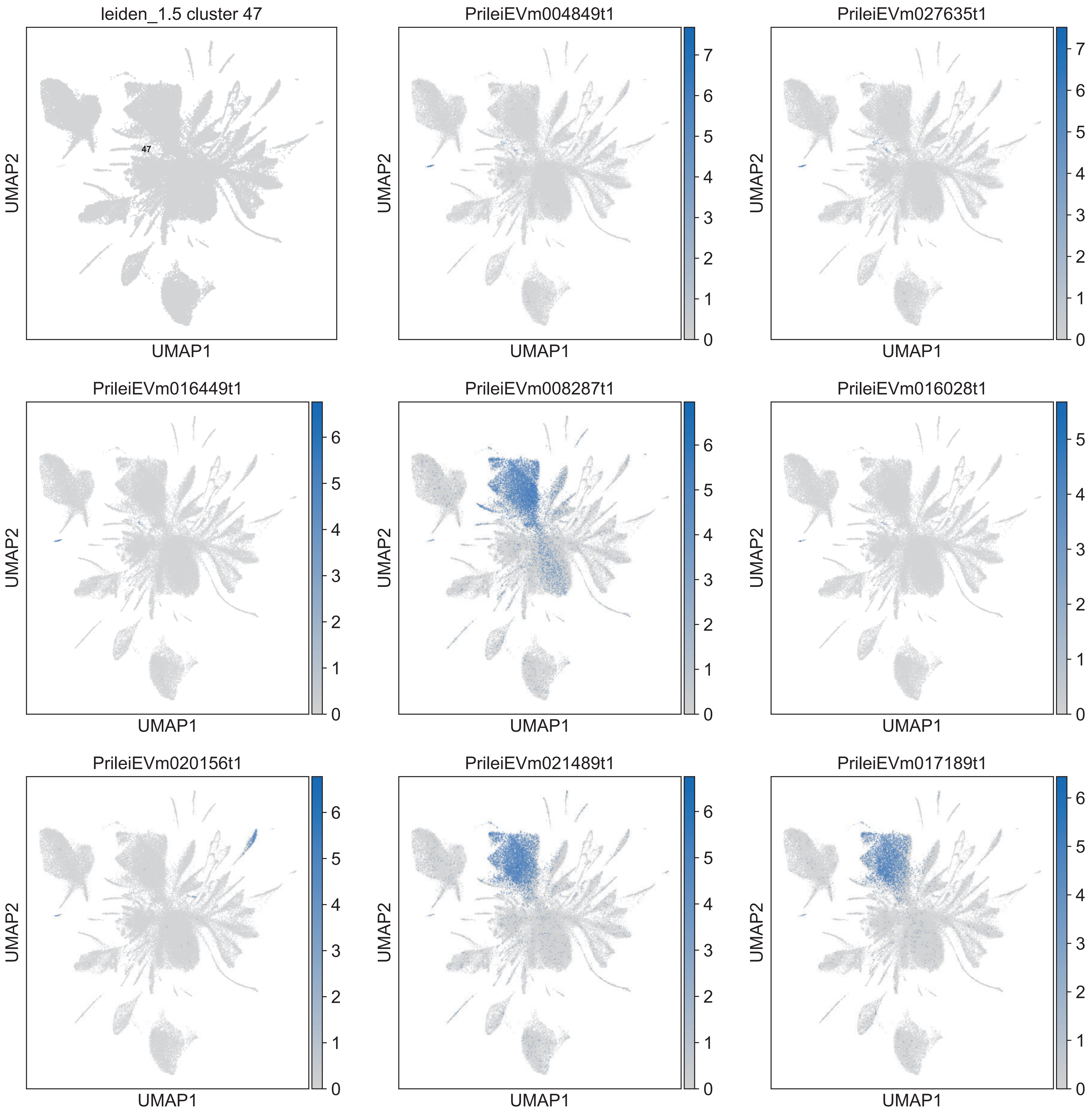
leiden_1.5 cluster 47
47
PrileiEVm004849t1
PrileiEVm027635t1
PrileiEVm016449t1
PrileiEVm008287t1
PrileiEVm016028t1
PrileiEVm020156t1
PrileiEVm021489t1
PrileiEVm017189t1
UMAP1
UMAP2

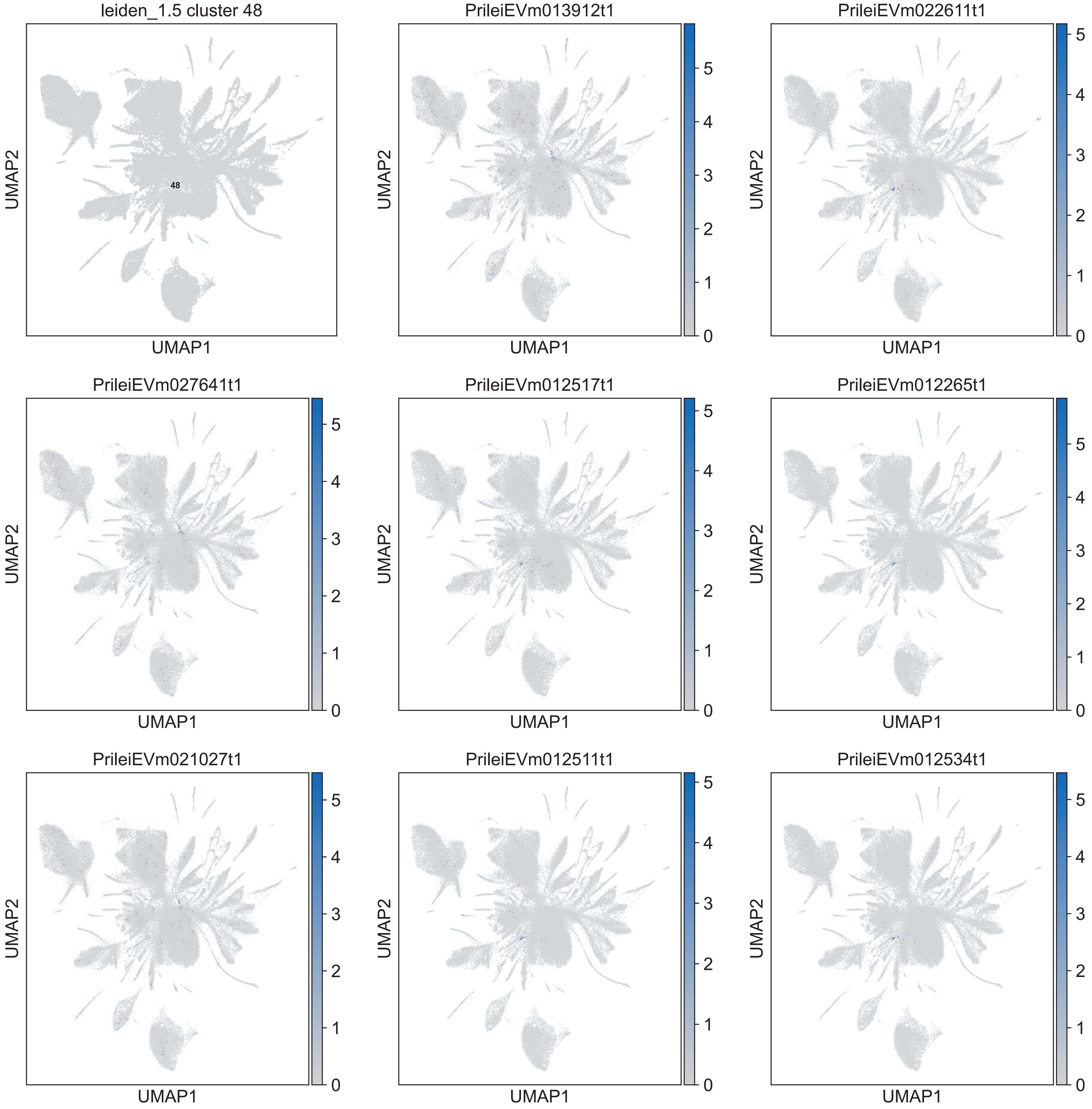
leiden_1.5 cluster 48
48
PrileiEVm013912t1
PrileiEVm022611t1
PrileiEVm027641t1
PrileiEVm012517t1
PrileiEVm012265t1
PrileiEVm021027t1
PrileiEVm012511t1
PrileiEVm012534t1
UMAP1
UMAP2

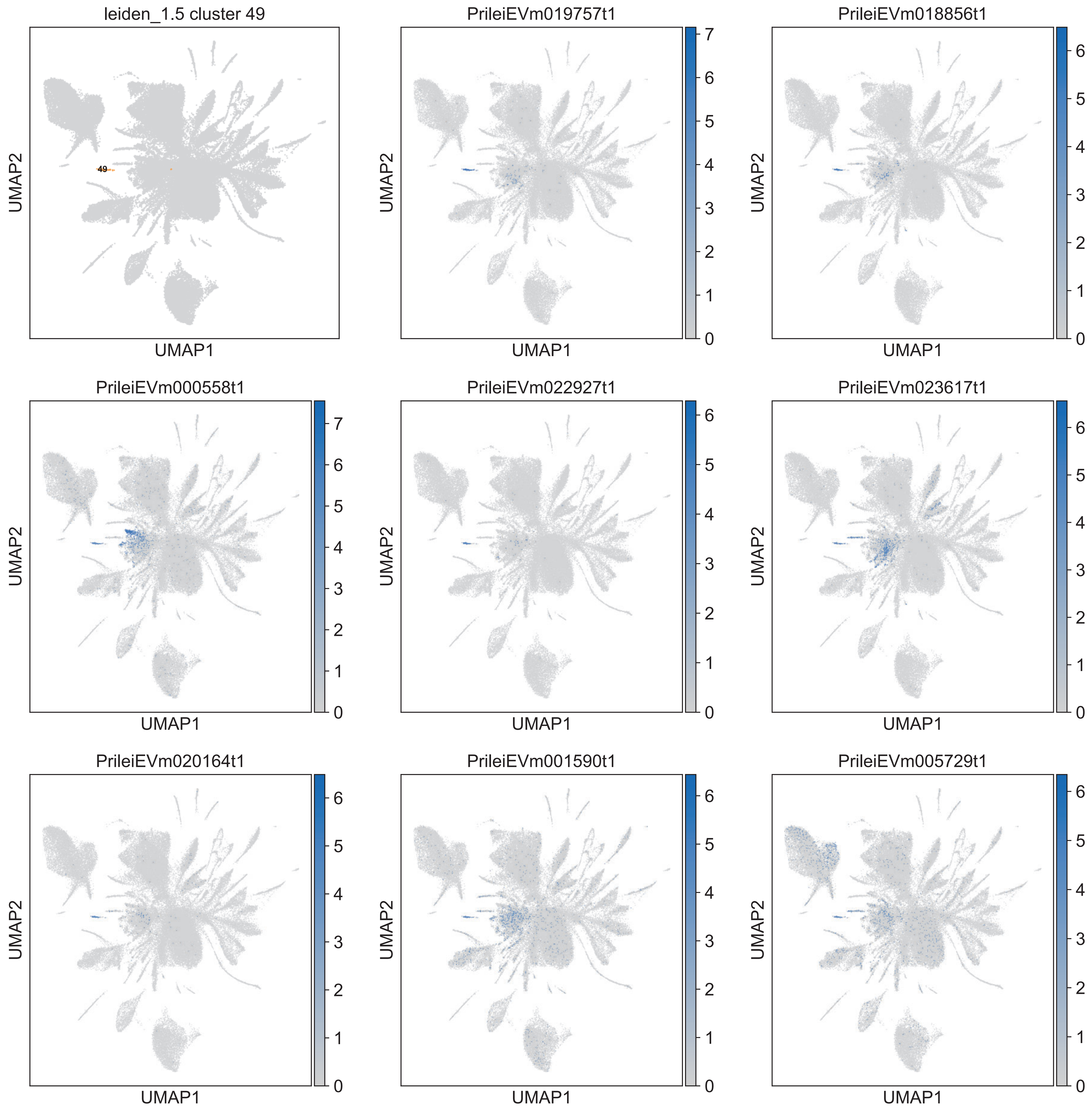

leiden_1.5 cluster 49
49
PrileiEVm019757t1
PrileiEVm018856t1
PrileiEVm000558t1
PrileiEVm022927t1
PrileiEVm023617t1
PrileiEVm020164t1
PrileiEVm001590t1
PrileiEVm005729t1
UMAP1
UMAP2

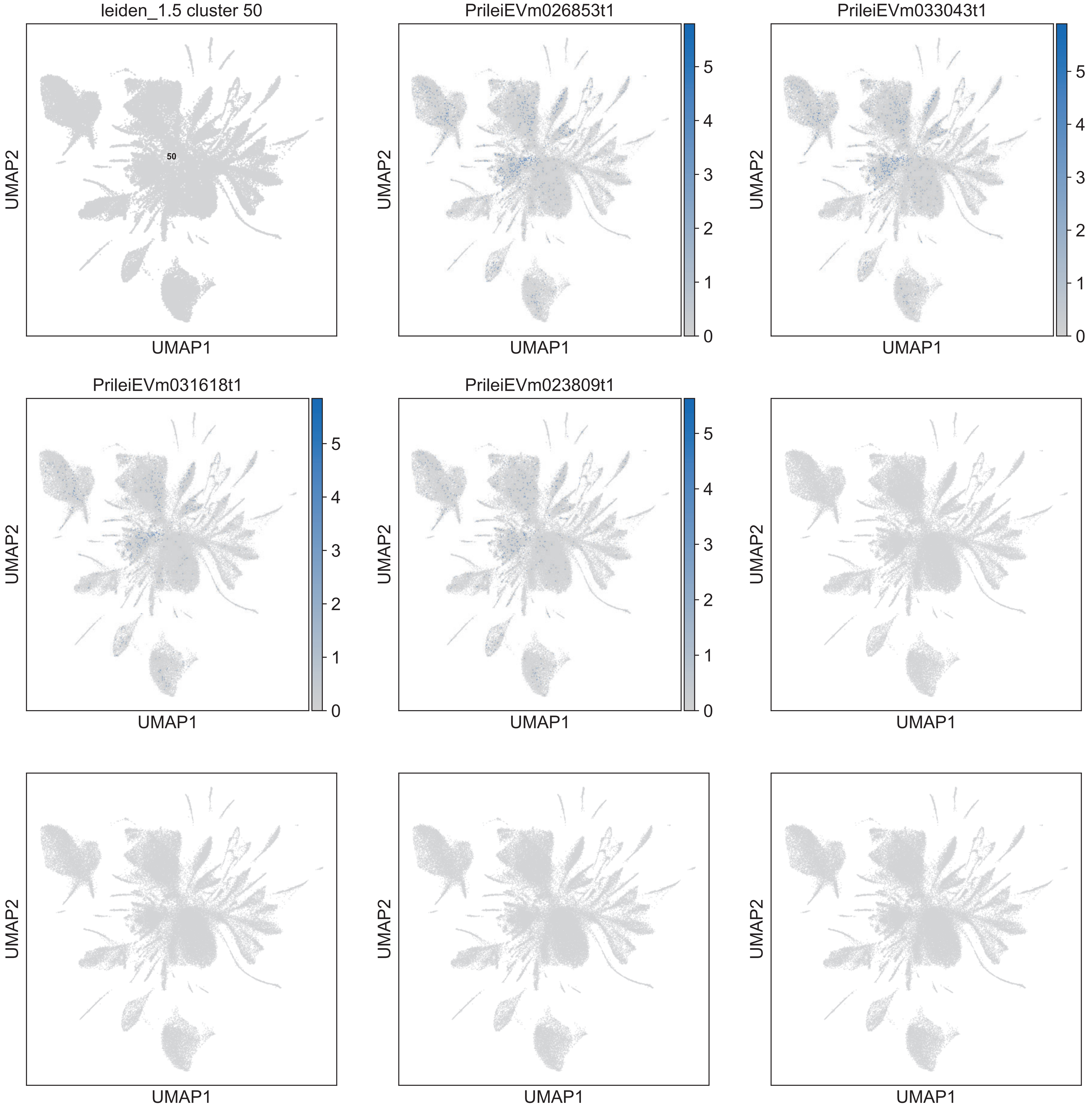

leiden_1.5 cluster 50
50
PrileiEVm026853t1
PrileiEVm033043t1
PrileiEVm031618t1
PrileiEVm023809t1
UMAP1
UMAP2
0
1
2
3
4
5

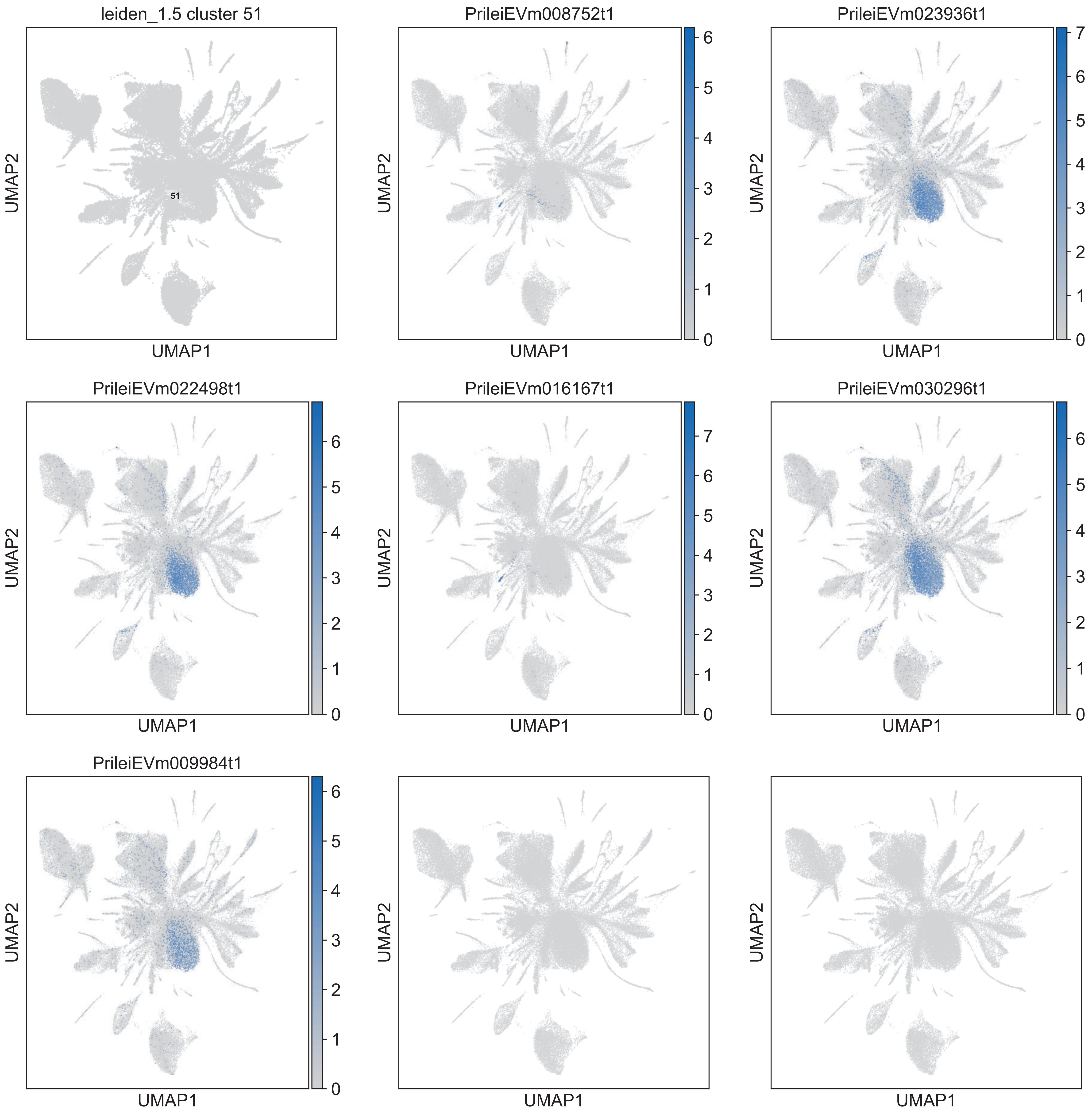
leiden_1.5 cluster 51
51
PrileiEVm008752t1
PrileiEVm023936t1
PrileiEVm022498t1
PrileiEVm016167t1
PrileiEVm030296t1
PrileiEVm009984t1
UMAP1
UMAP2
0
1
2
3
4
5
6
7

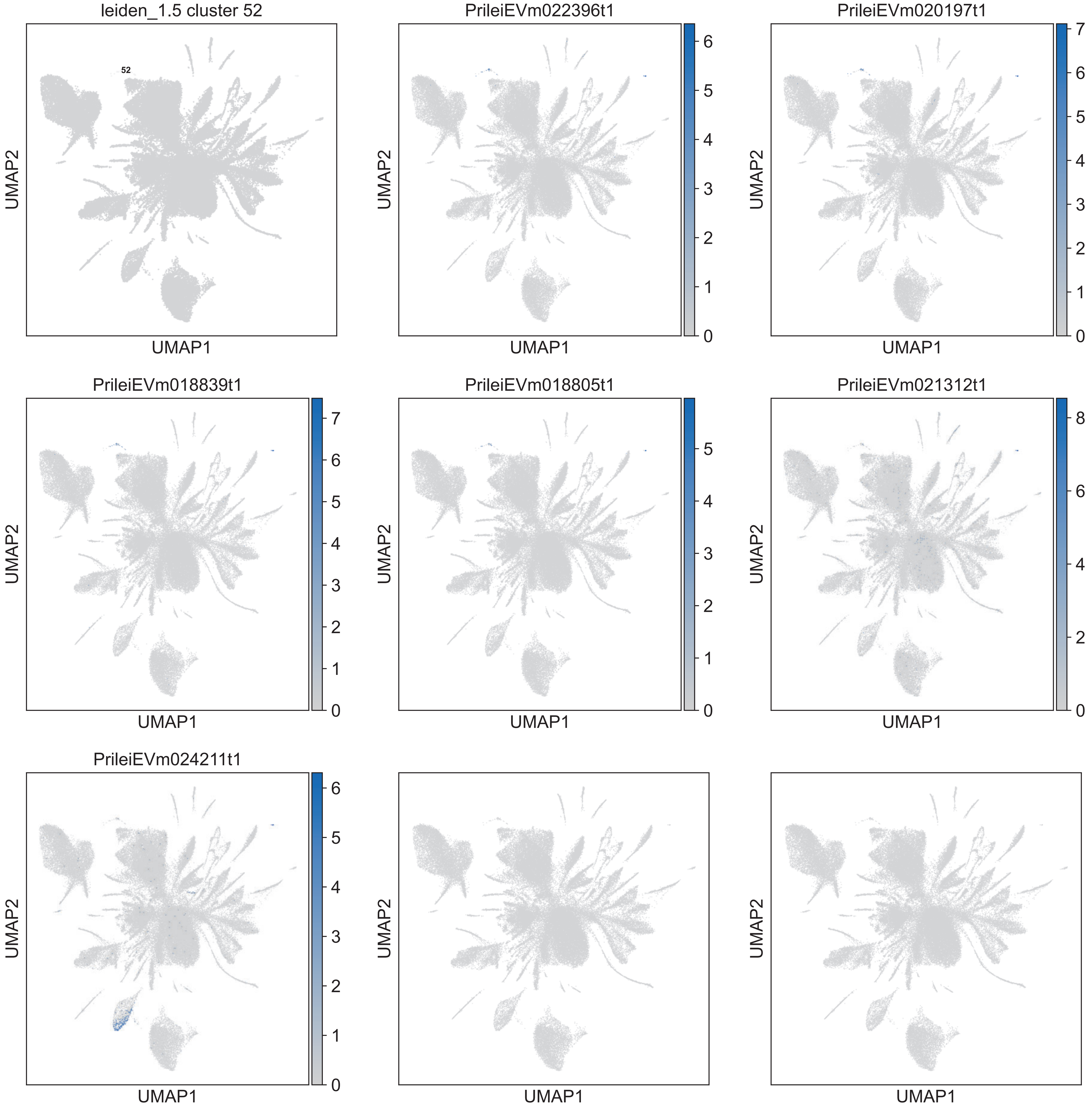
leiden_1.5 cluster 52
52
PrileiEVm022396t1
PrileiEVm020197t1
PrileiEVm018839t1
PrileiEVm018805t1
PrileiEVm021312t1
PrileiEVm024211t1
UMAP1
UMAP2

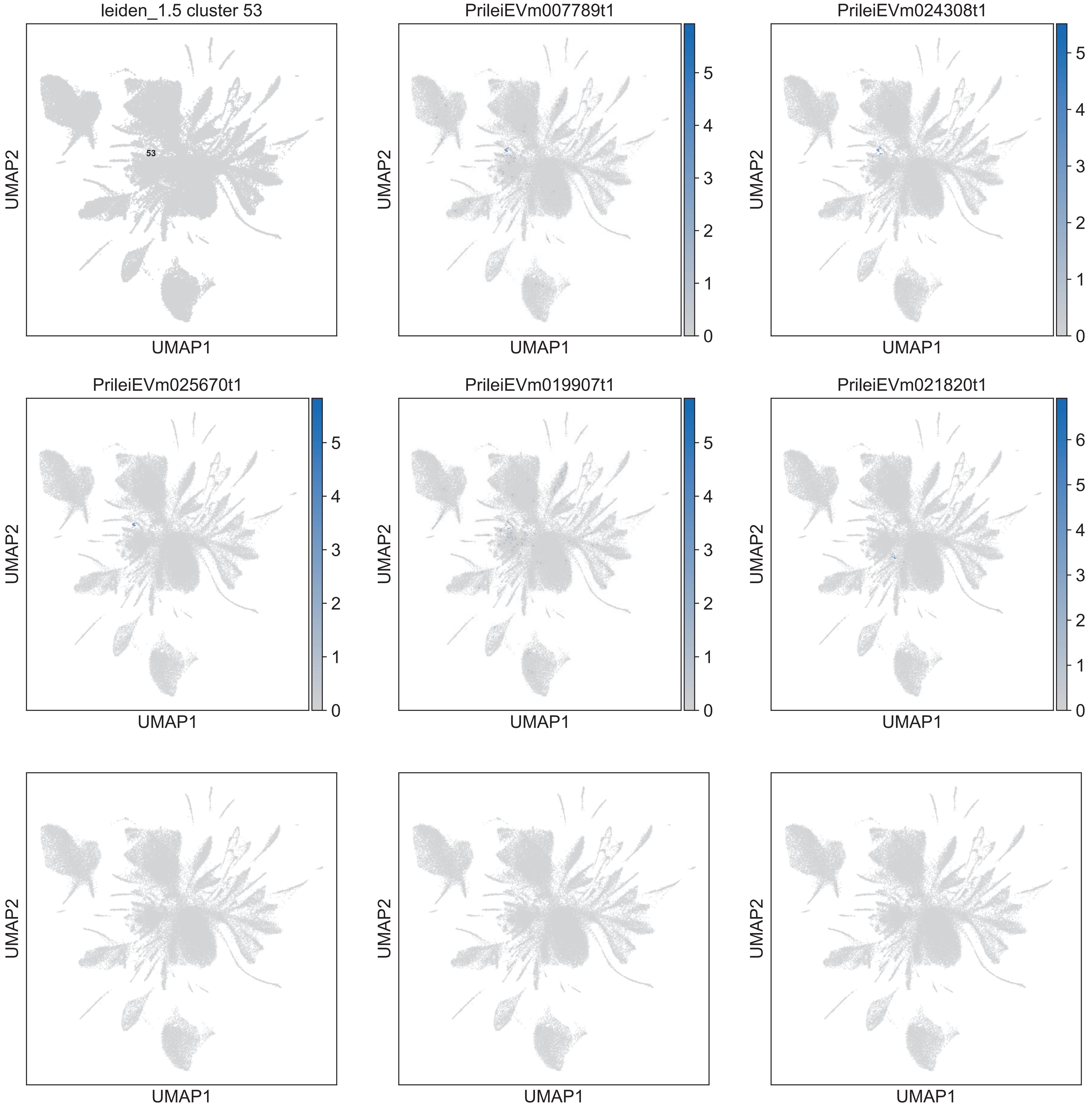
leiden_1.5 cluster 53
53
PrileiEVm007789t1
PrileiEVm024308t1
PrileiEVm025670t1
PrileiEVm019907t1
PrileiEVm021820t1
UMAP1
UMAP2

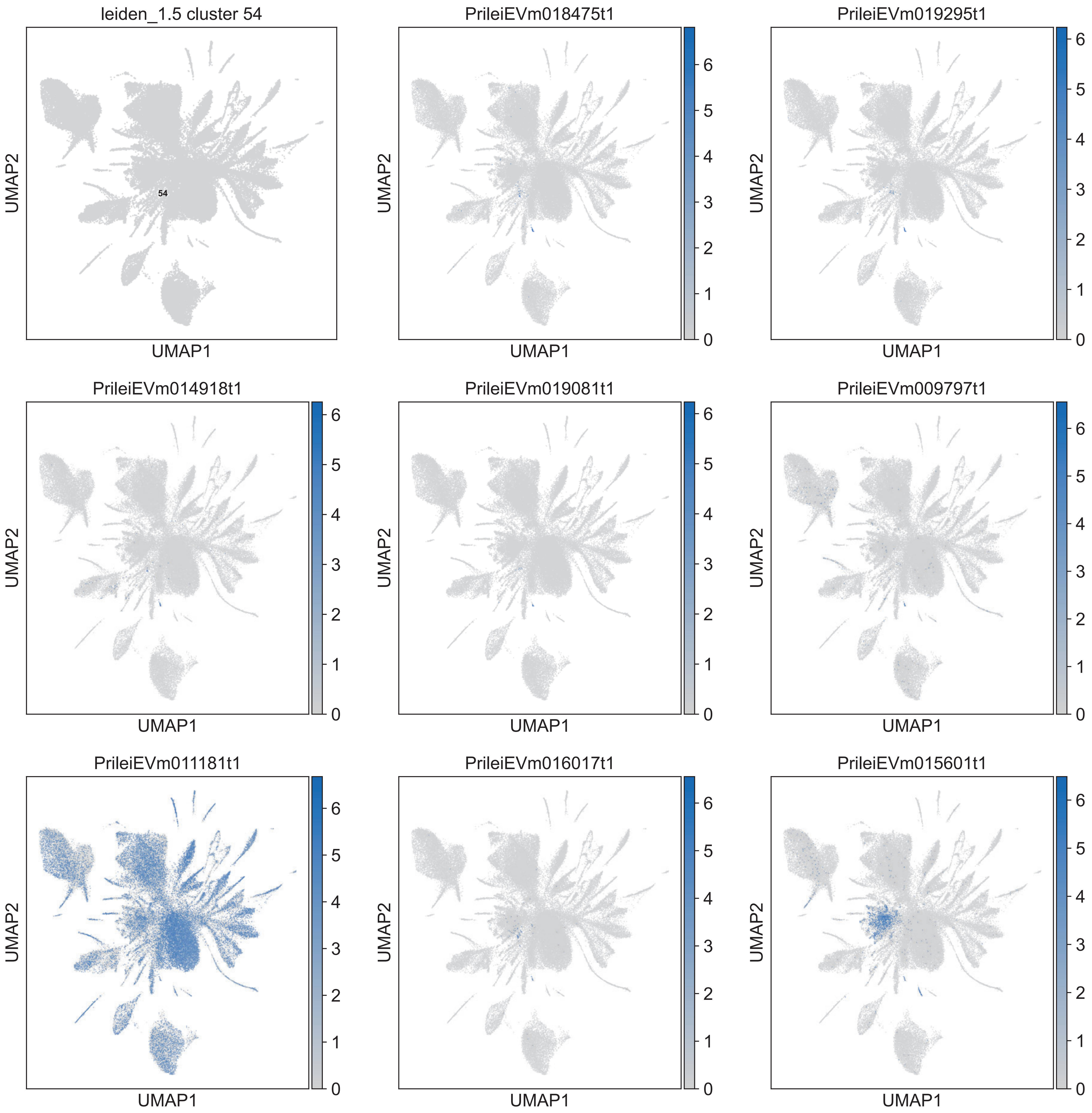
leiden_1.5 cluster 54
54
PrileiEVm018475t1
PrileiEVm019295t1
PrileiEVm014918t1
PrileiEVm019081t1
PrileiEVm009797t1
PrileiEVm011181t1
PrileiEVm016017t1
PrileiEVm015601t1
UMAP1
UMAP2
0
1
2
3
4
5
6

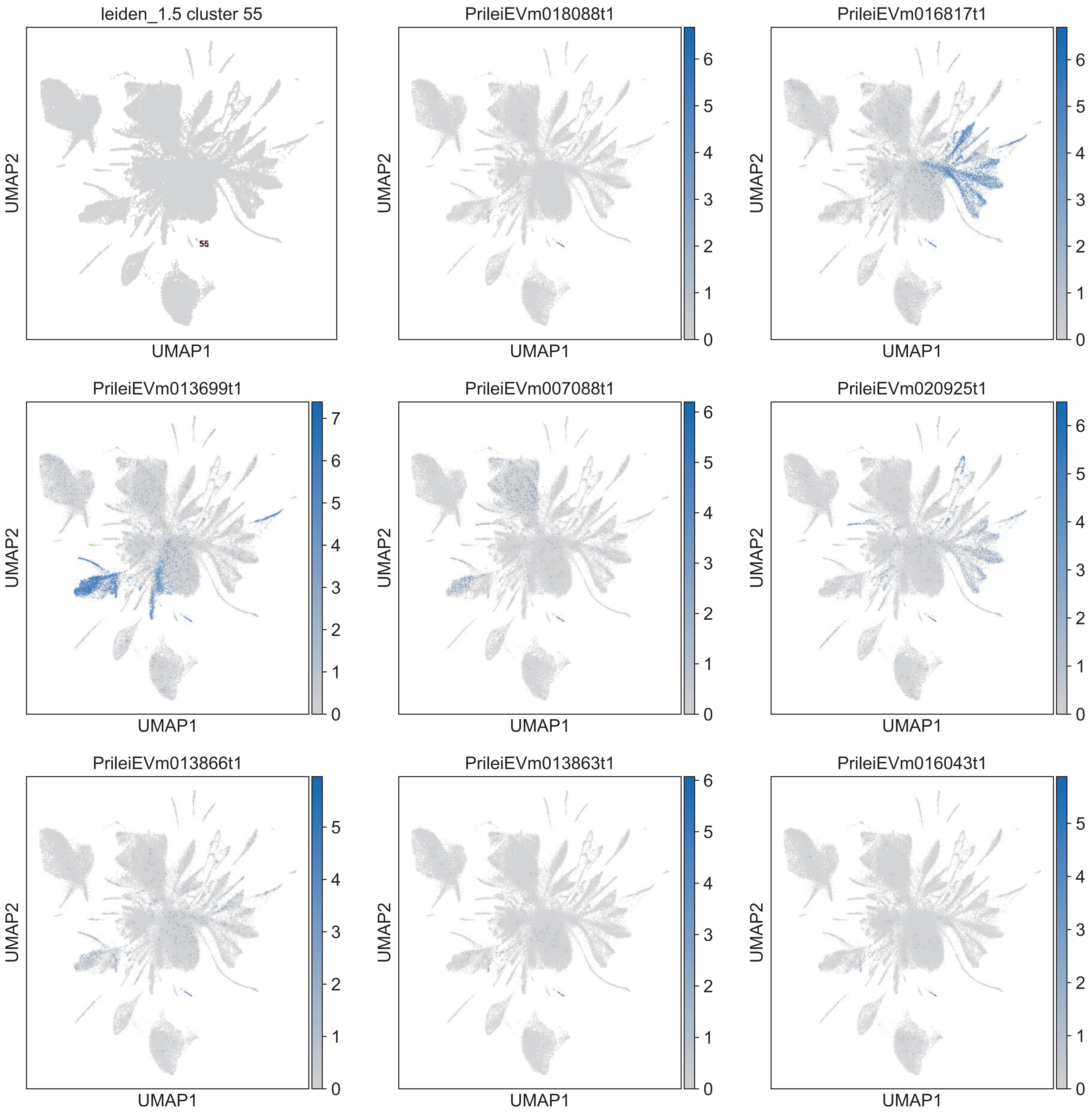
leiden_1.5 cluster 55
55
PrileiEVm018088t1
PrileiEVm016817t1
PrileiEVm013699t1
PrileiEVm007088t1
PrileiEVm020925t1
PrileiEVm013866t1
PrileiEVm013863t1
PrileiEVm016043t1
UMAP1
UMAP2
0
1
2
3
4
5
6
7

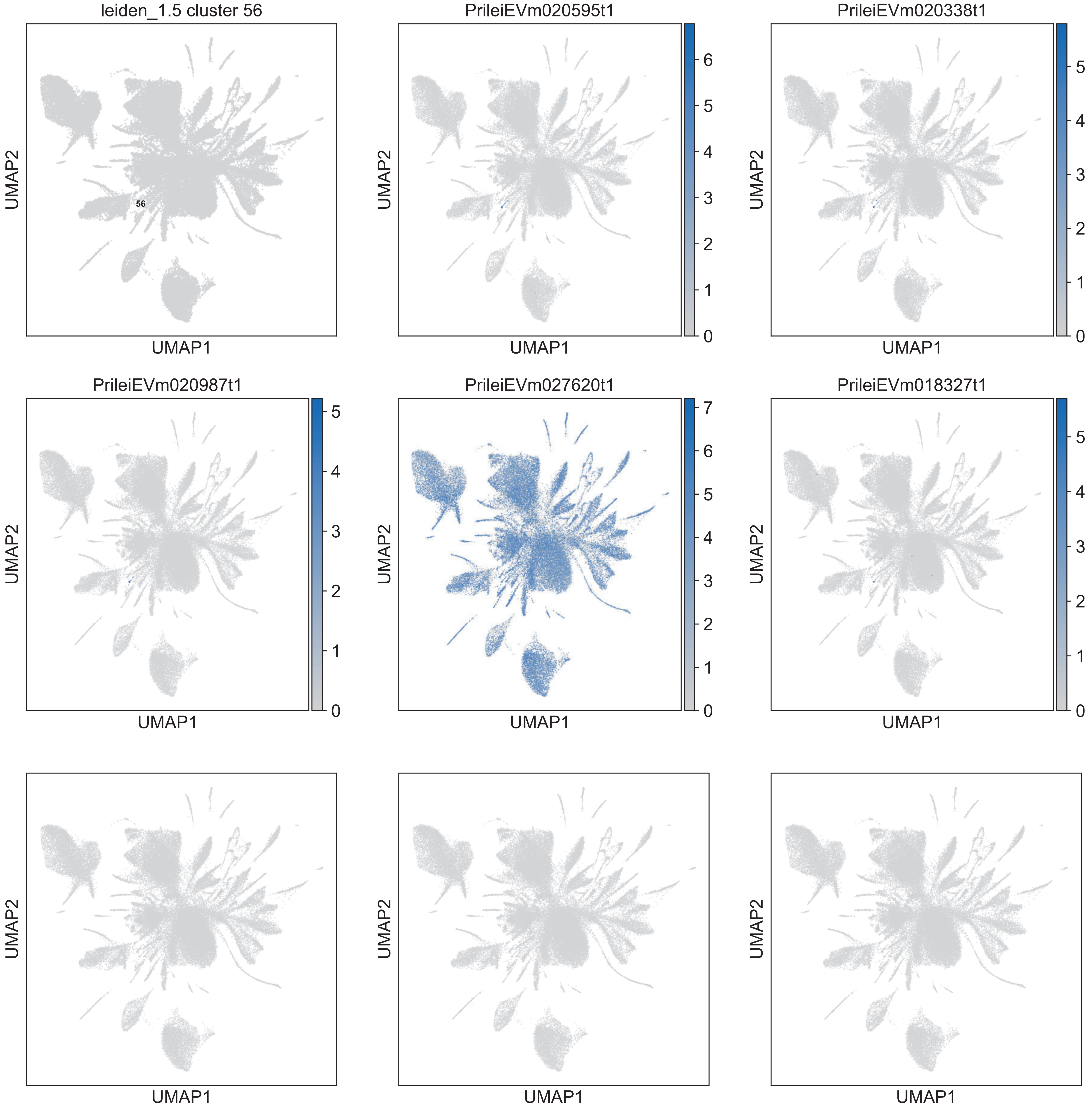

leiden_1.5 cluster 56
56
PrileiEVm020595t1
PrileiEVm020338t1
PrileiEVm020987t1
PrileiEVm027620t1
PrileiEVm018327t1
UMAP1
UMAP2

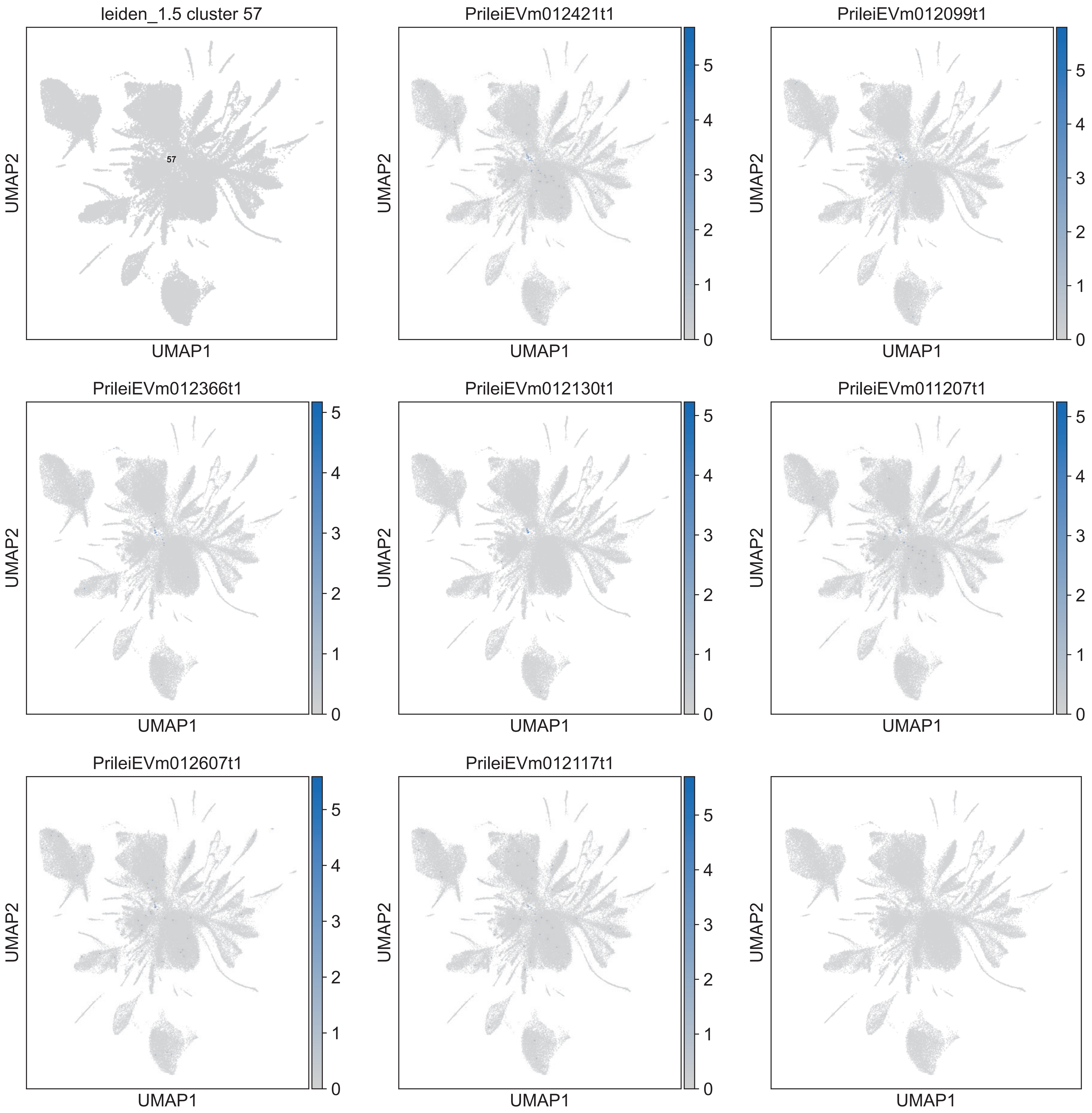
leiden_1.5 cluster 57
57
PrileiEVm012421t1
PrileiEVm012099t1
PrileiEVm012366t1
PrileiEVm012130t1
PrileiEVm011207t1
PrileiEVm012607t1
PrileiEVm012117t1
UMAP1
UMAP2
0
1
2
3
4
5

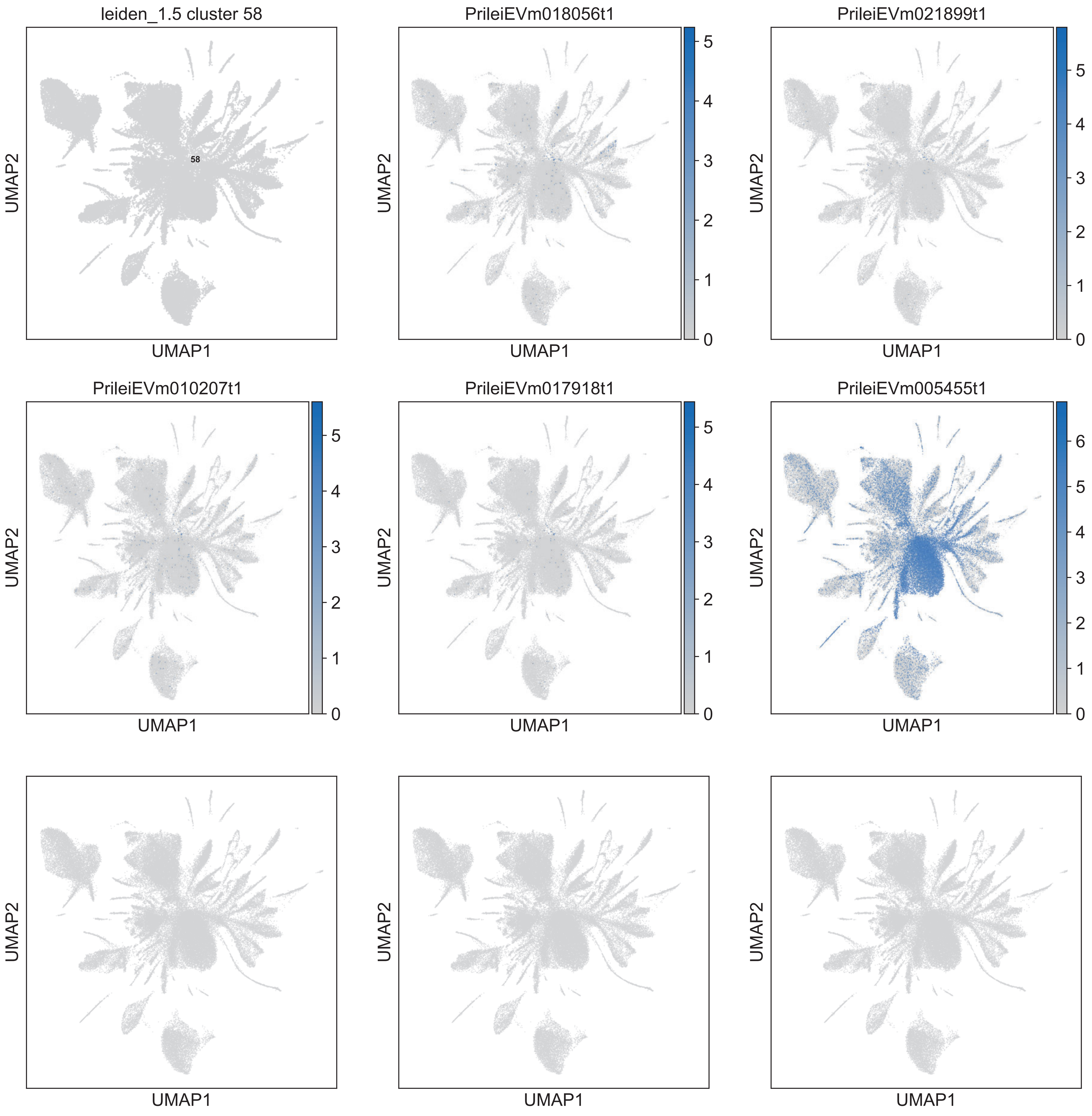
leiden_1.5 cluster 58
58
PrileiEVm018056t1
PrileiEVm021899t1
PrileiEVm010207t1
PrileiEVm017918t1
PrileiEVm005455t1
UMAP1
UMAP2

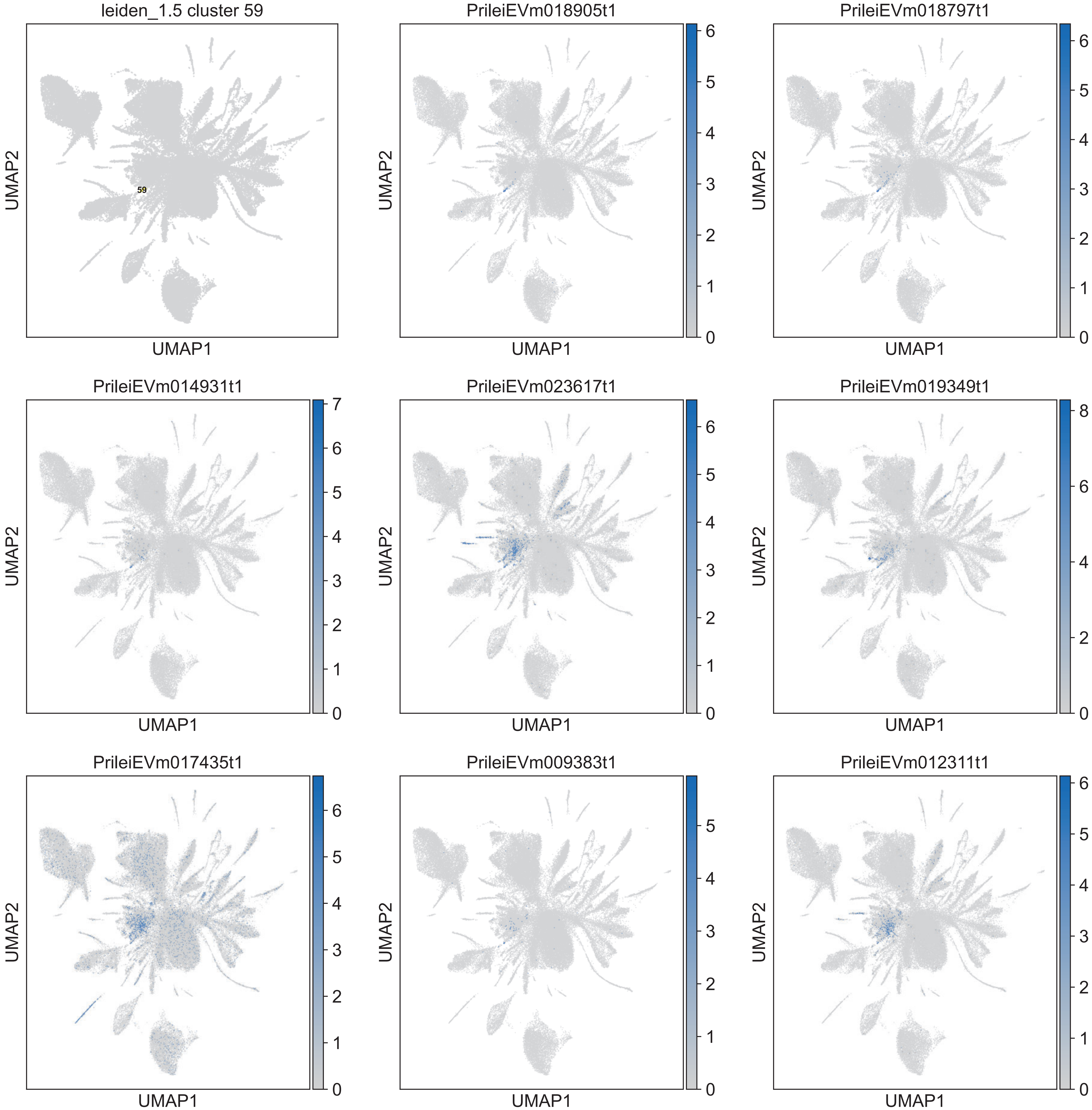
leiden_1.5 cluster 59
59
PrileiEVm018905t1
PrileiEVm018797t1
PrileiEVm014931t1
PrileiEVm023617t1
PrileiEVm019349t1
PrileiEVm017435t1
PrileiEVm009383t1
PrileiEVm012311t1
UMAP1
UMAP2
